# Ribosomal RNA operons define a central functional compartment in the *Streptomyces* chromosome

**DOI:** 10.1101/2022.06.23.497307

**Authors:** Jean-Noël Lorenzi, Annabelle Thibessard, Virginia S. Lioy, Frédéric Boccard, Pierre Leblond, Jean-Luc Pernodet, Stéphanie Bury-Moné

## Abstract

*Streptomyces* are prolific producers of specialized metabolites with applications in medicine and agriculture. These bacteria possess a large linear chromosome genetically compartmentalized: core genes are grouped in the central part, while terminal regions are populated by poorly conserved genes and define the chromosomal arms. In exponentially growing cells, chromosome conformation capture unveiled sharp boundaries formed by ribosomal RNA (*rrn*) operons that segment the chromosome into multiple domains. The first and last *rrn* operons delimit the highly expressed central compartment and the rather transcriptionally silent terminal compartments. Here we further explore the link between the genetic distribution of *rrn* operons and *Streptomyces* genomic compartmentalization. A large panel of genomes of species representative of the genus diversity revealed that *rrn* operons and core genes form a central skeleton, the former being identifiable from their core gene environment. We implemented a new nomenclature for *Streptomyces* genomes and trace their *rrn*-based evolutionary history. Remarkably, *rrn* operons are close to pericentric inversions. Moreover, the central compartment delimited by *rrn* operons has a very dense, nearly invariant core gene content. Finally, this compartment harbors genes with the highest expression levels, regardless of gene persistence and distance to the origin of replication. Our results highlight that *rrn* operons define the structural boundaries of a central functional compartment prone to transcription in *Streptomyces*.

## Introduction

*Streptomyces* are bacteria of great biotechnological interest due to the production of antibiotics and many other bioactive compounds (Berdy 2012). Remarkably for a bacterium, they have a linear chromosome and terminal inverted repeats (TIRs) capped by telomere-like sequences. Their genome is amongst the largest in bacteria (6-15 Mb), with an extreme GC content (circa 72%). In addition, the *Streptomyces* chromosome presents a partition, termed ‘genetic compartmentalization’, into a core region harboring genes shared by all *Streptomyces* and more variable extremities or ‘arms’ enriched in specialized metabolite biosynthetic gene clusters (Redenbach et al. 1996; Omura et al. 2001; Karoonuthaisiri et al. 2005; Ikeda et al. 2003; Choulet et al. 2006; Bentley et al. 2002; Lorenzi et al. 2021; Virginia S. Lioy et al. 2021). Consistently, DNA rearrangements and recombination events are more frequently fixed in the terminal arms than in the central region (Choulet et al. 2006; Fischer et al. 1998; Hoff et al. 2018; Tidjani et al. 2019; Hopwood 2006; Zhang et al. 2020). It has been proposed that strong evolutionary constraints shaped the distribution of genes along the chromosome owing to their potential benefit at the individual or population level (Lorenzi et al. 2021): genes encoding ‘private goods’ essential for vegetative growth are maintained in the central part of the genome, whereas social genes encoding ‘public goods’ of strong adaptive value for the colony (e.g. antibiotics) are located in the variable part of the genome, which may favor their rapid diversification. The mechanisms that govern the structure and function of these compartmentalized genomes remain mostly unknown.

We recently demonstrated that the genetic compartmentalization of *Streptomyces ambofaciens* ATCC 23877 correlates with chromosome architecture and gene expression in exponential phase (Virginia S. Lioy et al. 2021). During vegetative growth, the distal ribosomal RNA (*rrn*) operons delimit a highly structured and expressed region termed ‘central compartment’, presenting structural features distinct from those of the terminal compartments which are almost transcriptionally quiescent (Virginia S. Lioy et al. 2021). This led us to propose that these distal *rrn* operons may constitute some kind of barrier contributing to the evolution of *Streptomyces* genomes towards a compartmentalized organization (Virginia S. Lioy et al. 2021).

The number of *rrn* copies is thought to be a determinant of bacterial fitness, with the optimal number depending on the environmental and biological context in which the species evolve (Stevenson et Schmidt 2004; Roller, Stoddard, et Schmidt 2016; Espejo et Plaza 2018; Fleurier et al. 2022). Although 16S RNA sequences are the classical ‘chronometer’ for phylogenetic classification, their impact on genome evolution *per se* has rarely been considered (Espejo et Plaza 2018). Interestingly, the *rrn* operons, including 16S, 23S, 5S and internal transcribed spacer regions, coincide with sharp boundaries in the chromosome 3D-organization of bacteria with linear (Virginia S. Lioy et al. 2021) as well as circular (V. S. Lioy et al. 2018; Le et Laub 2016; Böhm et al. 2020; Marbouty et al. 2015; Wang et al. 2015) genomes. The formation of these boundaries correlates with a very high level of transcription, but does not require translation (Le et Laub 2016). Moreover, RNA polymerase is spatially organized into dense clusters engaged in ribosomal RNA synthesis when bacteria are grown in rich medium (D. J. Jin et Cabrera 2006; Cabrera et Jin 2006; Mata Martin et al. 2018; Weng et al. 2019). It has been proposed that *rrn* operons might form a bacterial equivalent of the nucleolus (Gaal et al. 2016), although these results remain controversial (Mata Martin et al. 2018). Altogether, these observations open the possibility that the *rrn* operons could play a role in genome evolution by coupling transcription and genome spatial conformation.

Guided by this hypothesis, we took advantage of the large number of sequenced *Streptomyces* genomes to explore the correlation between *rrn* operon dynamics (number, position) and chromosome organization in a panel of species representative of *Streptomyces* diversity. We notably observed that *rrn* operons coevolved with the core region and can be identified from their core gene environment. We set-up an *rrn*-based nomenclature for *Streptomyces* genome organization that we used to trace its evolutionary history. Pericentric recombination frequently occurred at the vicinity of *rrn* operons located close to the origin of replication. Moreover, we observed that the most external *rrn* operons, designated ‘distal *rrn* operons’, delimit the central compartment, whose size and content are highly correlated with the core genome dynamics. Genes within this central compartment are expressed at a higher level than in the terminal compartments, regardless of gene persistence and the distance to the origin of replication. Altogether, our results highlight that distal *rrn* operons may be considered as ‘structural limits’ that delineate a functional compartment in the linear genome of *Streptomyces*.

## Results

### New nomenclature of *Streptomyces* genomes based on *rrn* operons and *dnaA* gene orientation

We first characterized the organization of *rrn* operons in 127 *Streptomyces* genomes from a previously characterized panel of *Streptomyces* species representative of the genus diversity (Lorenzi et al. 2021; Virginia S. Lioy et al. 2021) (**Supplemental Table S1**). In this panel, most genomes (>85 %) share an average nucleotide identity based on BLAST+ (ANIb) lower than 95 %, a threshold used to distinguish species (Richter et Rosselló-Móra 2009) (**Supplemental Table S2**). We included several strains for eight species (*e.g. S. ambofaciens*, *Streptomyces venezuelae*) to access intra-species evolution and include strains for which –omics data were available for further analyses. We re-annotated all genomes and detected orthologous genes, as previously described (Lorenzi et al. 2021). This allowed the identification of 1,017 ortholog genes associated with best reciprocal matches between coding sequences present in all 127 genomes, further defined as the ‘core genome’. Interestingly, 943 of these genes (92.7 %) are included in the soft-core recently identified on a partially overlapping panel of *Streptomyces* genomes by Caicedo-Montoya *et al*. (Roary method) (Caicedo-Montoya, Manzo-Ruiz, et Ríos-Estepa 2021). We used the position of the most external genes of the core genome as limits between the ‘arms’ and the ‘core region’ (that therefore includes all the core genes and some non-core genes, **Fig. 1A**).

**Figure 1:**
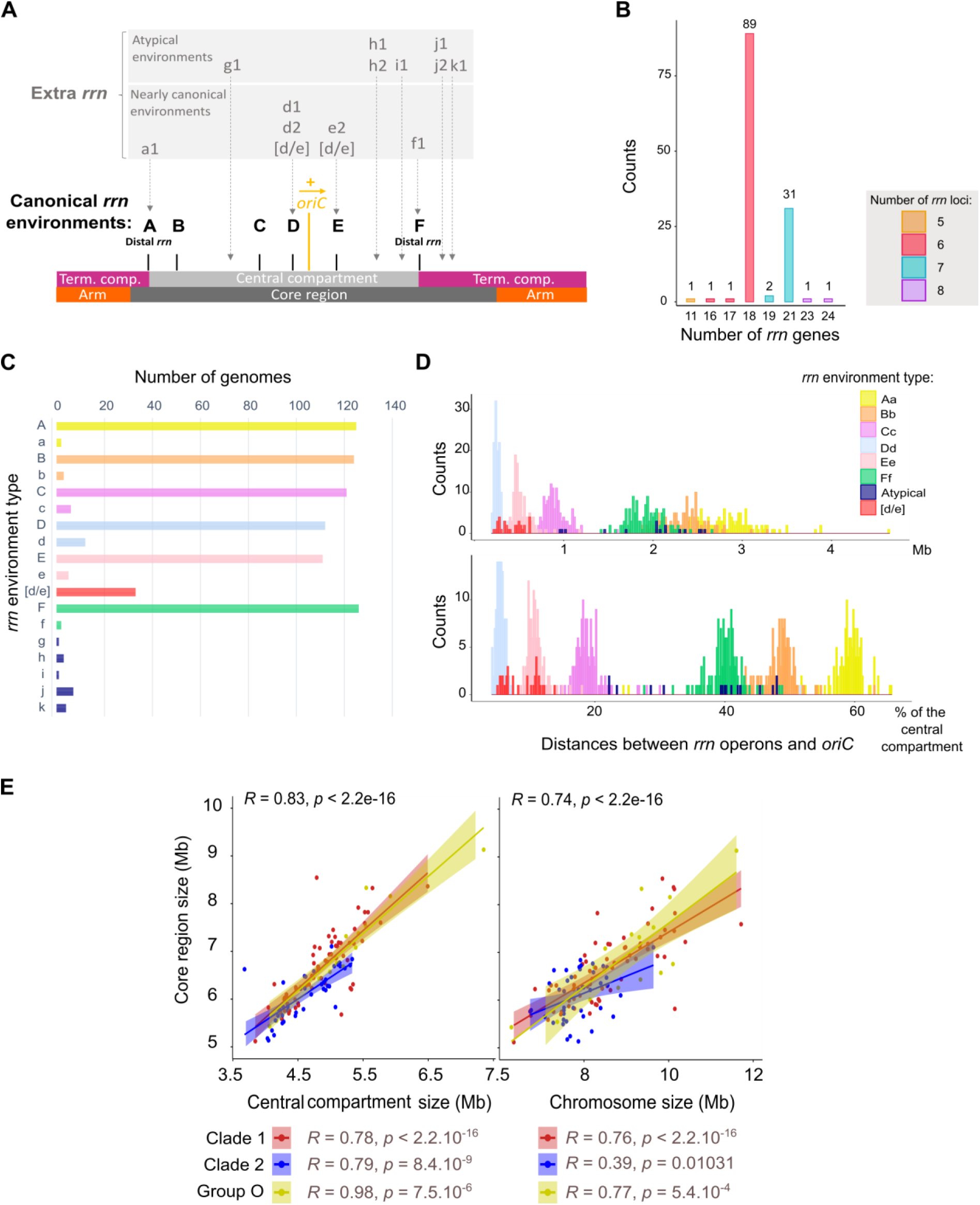
*Streptomyces rrn* operon genetic distribution and link with the core genome. **A. Schematic representation of the location of *rrn* operons in the *Streptomyces* genome.** The schematic representation of the chromosome is shown to scale using *S. ambofaciens* ATCC 23877 as reference. The origin of replication (*oriC*) was defined regarding the position of the *dnaA* gene, the yellow arrow representing the orientation of this gene. The detailed nomenclature describing each *rrn* environment is available in **Supplemental Figure S2**. Abbreviation: ‘Term. comp.’ = Terminal compartment. **B. Number of *rrn* genes in the panel of 127 genomes.** The bars are filled according to the number of *rrn* loci (corresponding to complete or incomplete operons) in each genome. The values above each box correspond to the number of genomes. **C. Frequency of the different *rrn* core gene environments in the panel of 127 genomes.** **D. Distribution of the distances between each *rrn* operon and the origin of replication (*oriC*) in the panel of 127 genomes.** The distance is expressed in Mb (top) or as the percentage of the size of the ‘central compartment’ (bottom). The results are presented separately for each *rrn* category as defined in **Supplemental Figure S2**. **E. Scatter plots presenting the correlation between the core region size and the size of the central compartment or the chromosome**. The rho coefficients (*R*) and *p* values of Spearman’s rank correlations were calculated with the whole set of genomes (*n* = 127) as well as within each clade and group (*n*_Clade 1_ = 67, *n*_Clade 2_ =43, *n*_Group O_ =17).

Most *Streptomyces* genomes from our panel (70.1 %) harbor six *rrn* operons encoding all three 16S, 23S, 5S ribosomal RNAs (**Fig. 1B**). About a quarter of genomes (24.4 %) contain seven complete *rrn* operons, eight complete operons being quite exceptional (only *Streptomyces hundungensis* BH38). These results are in accordance with the number of 16S *rrn* genes per strain reported in the *rrn*DB database (Roller, Stoddard, et Schmidt 2016) in a panel of 265 *Streptomyces* genomes (74.0 % and 22.3 % with six and seven 16S *rrn* genes, respectively - https://rrndb.umms.med.umich.edu/, version 5.7). Thus, sampling biases seem negligible when comparing data from our panel and an independent set of genomes.

Pairwise comparison of core genomes revealed that synteny of core genes is strong between all strains, highlighting a ‘core skeleton’ with a rather stable core gene order in *Streptomyces* (**Supplemental Fig. S1**). In the middle of the genome, we confirmed the existence of a region at the origin of replication in which core gene synteny is perfectly conserved between all strains, as previously described for a smaller set of strains (Algora-Gallardo et al. 2021). We then determined a consensus order of core genes by assigning them their most frequent rank in a panel of genomes representative of the most frequent global core skeleton organization. Interestingly, five core genomes of the panel (*e.g. Streptomyces viridosporus* T7A ATCC 39115) present exactly this consensus organization, and twelve (e.g. *S. ambofaciens* both strains in the clade 1, *Streptomyces ficellus* NRRL 8067 in the clade 2) differ only by the order of 2 genes of the core genome owing to a local inversion (**Table S3**).

In accordance with the existence of a core skeleton, we noticed that six *rrn* operons almost always have the same core gene environment in all the strains (**Fig. 1A**). These six *rrn* core gene neighborhoods, hereinafter referred to as ‘canonical’ and designated from ‘A’ to ‘F’ in capital letters (**Supplemental Fig. S2**), are exactly conserved in 87.4 to 99.2 % of the genomes (**Fig. 1C**). In this nomenclature, the same letter is kept when at least one core gene is in common between two *rrn* genetic environments. An asterisk (**Supplemental Fig. S1, Supplemental Table S1**) has been added to indicate an identical environment but in reverse orientation to that shown in the **Supplemental Figure S2**. We also identified recombination between ‘D’ and ‘E’ *rrn* operons, leading to [d/e] hybrid core gene environments in some strains (**Fig. 1A**). On the contrary, the 7^th^ and 8^th^ *rrn* operons (when present) can be located in various core gene environments, named ‘g’ through ‘k’ in lower case with a number, to indicate their non-canonical nature (**Fig. 1C**). Based on these observations, we proposed that the ancestor of all *Streptomyces* had 6 *rrn* operons, the 7^th^ and 8^th^ *rrn* operons emerging from *rrn* operon duplication/acquisition in the vicinity of a canonical *rrn* operon (e.g. ‘e2’ *rrn* operon in *S. venezuelae* ATCC 10712) or at an ectopic position (e.g. ‘k1’ *rrn* operon in *Streptomyces albidoflavus* strains and *S. hundungensis* BH38). Accordingly, the presence of only five *rrn* operons [*Streptomyces asterosporus* (synonym: *calvus*) DSM 41452] likely corresponds to the loss of an *rrn* operon.

Each genome was thus classified according to the orientation of the *rrn* genetic environments and the *dnaA* gene (as a proxy of the origin of replication orientation) (**Supplemental Table S1**).

In the panel, 40.9 % of the genomes harbor the *rrn* operons in the same order along the core genome, the major shared configuration being ‘*rrn* ABCDEF *dnaA*+’ (*e.g. S. ambofaciens* ATCC 23877), which was subsequently considered canonical (**Supplemental Table S1**). The second most frequent configuration represents 10.2% of cases and corresponds to genomes harboring a pericentric inversion [‘*rrn* ABCE*D*F *dnaA*-‘, *e.g. Streptomyces coelicolor* A3(2)] (**Supplemental Table S1**). Not taking into account extra copies of *rrn* or small local variations of the *rrn* environments, 50.4 % and 18.1 % of the strains have these two configurations: ‘*rrn* Aa Bb Cc Dd Ee Ff, *dnaA*+ (± extra *rrn*)’, and ‘*rrn* Aa Bb Cc (Ee)* (Dd)* Ff, *dnaA*- (± extra *rrn*)’, respectively (**Supplemental Table S1; Figure 2**). These results confirm that the proposed nomenclature allows a fairly general description of the organization of *Streptomyces* genomes, and that the ancestor of this genus probably had an ‘*rrn* ABCDEF *dnaA*+’-type genome.

**Figure 2:**
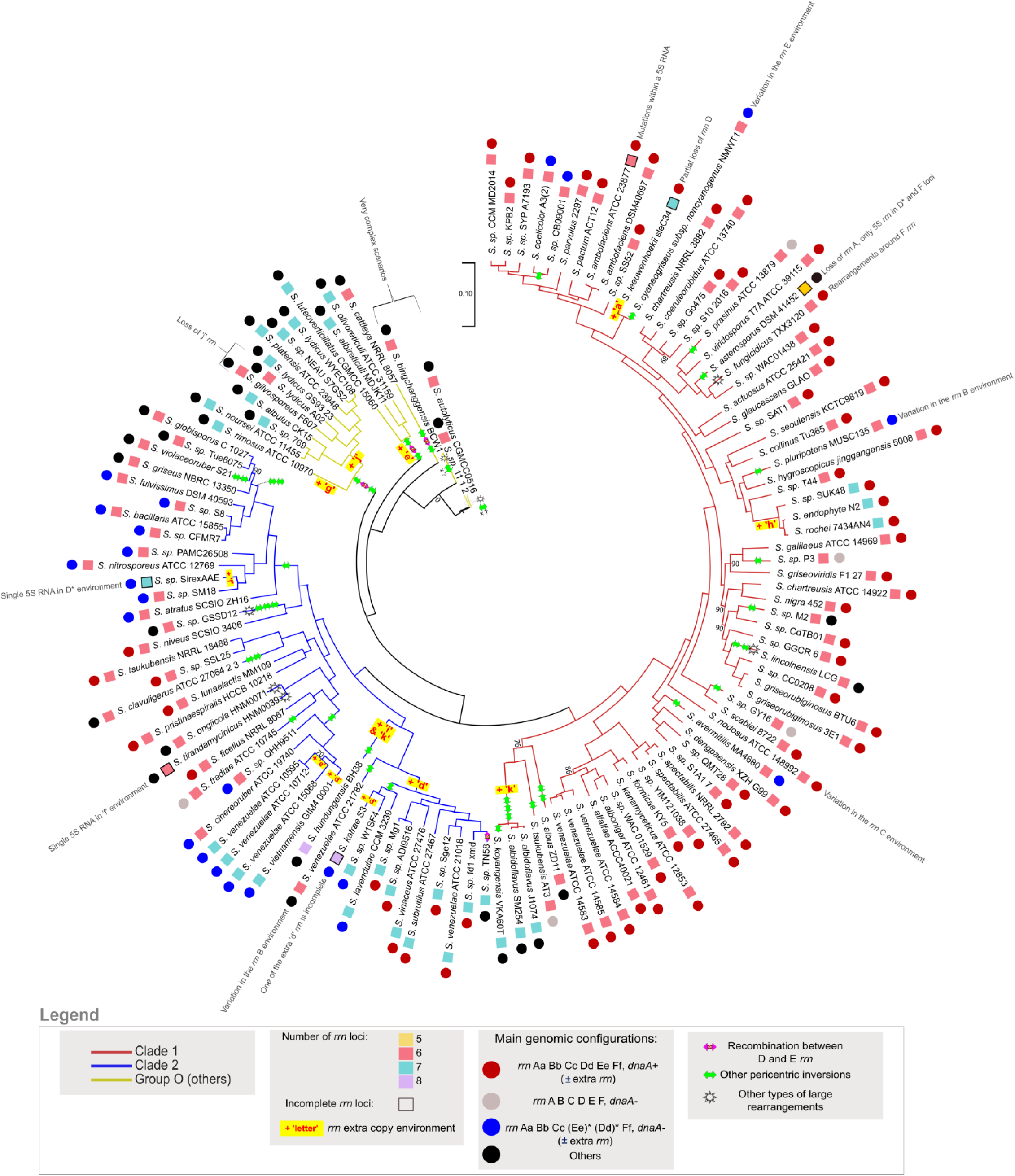
Core genome phylogenetic tree and proposed model of *Streptomyces* chromosome evolution regarding *rrn* operons and pericentric inversions. The core genome phylogenetic tree was constructed using the 1017 core genes. The bootstrap values inferior to 95 % are indicated. Branch colors represent the two clades (1 and 2) and other lineages (group ‘O’) of *Streptomyces* previously reported (McDonald et Currie 2017; Lorenzi et al. 2021; Caicedo-Montoya, Manzo-Ruiz, et Ríos-Estepa 2021). The number and completeness of *rrn* loci as well as *rrn* configuration and main intra-chromosomal rearrangements are indicated for each strain as detailed in the legend panel. Some specific events are indicated next to the relevant strains/species. The most parsimonious scenario is proposed, but in some cases (indicated by a sun), complex rearrangements in the central compartment make it difficult to develop robust evolutionary scenarios. The **supplemental** Figures S1 and S3 present pairwise comparisons of the core genomes that support this model. Interestingly, the pairwise comparison of the core genome order of the strains *Streptomyces* sp. 11 1 2, *S. autolyticus* CGMCC0516 and *S. bingchenggensis* BCW 1 suggests that they probably have a common ancestor (*S.* sp. 11 1 2 and *S. autolyticus* CGMCC0516 having almost the same core gene order), which the core-based phylogenetic tree fails to resolve clearly (**Supplemental Fig. S3.I**). The *rrn* configuration of each strain/species is detailed in **Supplemental Table S1**. The relative position of the events described (inversion, loss/acquisition of *rrn*, complex rearrangements) is arbitrary and does not predict the order in which the events occurred. The sign “x ?” indicates that there have been several pericentric inversions, their exact number being difficult to determine due to the highly rearranged organization of the genomes.

Consistent with this conservation of the *rrn* genomic core environment, the distribution of the *rrn* operons along the chromosome seems rather conserved for each *rrn* category (except for the atypical *rrn* operons) and evenly spaced from the origin of replication (*oriC*) (**Fig. 1D**). This phenomenon is particularly visible if considering the distance between the *rrn* operons and *oriC* relative to the total size of the central compartment (rather than in bp) (**Fig. 1D**), suggesting that the central compartment appears to be an entity within which the distances of *rrn* operons to the origin co-evolve. Moreover, the distribution of *rrn* operons on either side of the origin of replication is asymmetric (2/3 on one side and 1/3 on the other) as are the distances of the A and F *rrn* operons from the origin (**Fig. 1A & D**), leading to an imbalance in terminal compartment sizes.

Finally, we observed a strong correlation between the size of this core region and the size of the central compartment (**Fig.1E**). Remarkably, these correlations are stronger than between the sizes of the core region and the whole chromosome, this latter correlation being not even statistically significant in clade 2 (**Fig. 1E**).

Altogether, these observations give rise to a vision of the *Streptomyces* chromosome organized around a conserved skeleton constituted by both the core and the *rrn* genes.

### Evolutionary history of the *Streptomyces* genome in relation to *rrn* dynamics

The core genome was used to reconstruct a phylogenetic tree which recapitulates the previously described (McDonald et Currie 2017; Lorenzi et al. 2021; Caicedo-Montoya, Manzo-Ruiz, et Ríos-Estepa 2021) division of the *Streptomyces* genus into two main monophyletic clades (clades ‘1’ and ‘2’) and other lineages further referred to as group ‘O’ (for ‘others’) (**Figure 2**). By crossing this tree with genome nomenclature based on *rrn* categories and *dnaA* orientation, we propose a parsimonious scenario explaining the diversity observed in the panel of 127 analyzed genomes. According to this model, recombination between ‘D’ and ‘E’ *rrn* operons, and duplication/acquisition of *rrn* operons occurred at least 4 and 13 times, respectively (**Figure 2**). Notably, *rrn* duplications/acquisition occurred or were fixed more frequently in clade 2 and the group ‘O’ than in clade 1 (odds ratio respectively of 5.5 and 14.9, *p* values respectively of 6.7.10^-4^ and 1.2.10^-5^, Fisher’s Exact Test for Count Data) (**Figure 2, Supplemental Table S1**). The two strains harboring 8 *rrn* operons both belong to clade 2. Remarkably, none of the genomes in the group ‘O’ show any of the most frequent *rrn* configurations, highlighting the complex evolutionary history of these species (**Supplemental Table S1, Figure 2**).

We also identify two events of complete *rrn* operon loss by analyzing the phylogeny of *S. asterosporus* DSM 41452, *Streptomyces lydicus* A02 and *Streptomyces gilvosporeus* F607 strains (**Figure 2**). Moreover, a few strains harbor incomplete *rrn* loci, devoid of functional 5S (*S. ambofaciens* ATCC 23877) or 16S (*Streptomyces katrae* S3) *rrn* genes, or in most cases, composed of a single 5S *rrn* (*Streptomyces* sp. Sirex AA-E, *Streptomyces leeuwenhoekii* C34, *Streptomyces tirandamycinicus* HNM0039, *S. asterosporus* DSM 41452). To note, in the incomplete *rrn* operon of *S. ambofaciens* ATCC 23877, the sequence encoding the 5S *rrn* gene is present but has accumulated mutations (**Supplemental Figure S2.B**). Remarkably, *S. asterosporus* DSM 41452 harbors only three complete *rrn* operons, its two other *rrn* loci corresponding to single 5S *rrn* genes whose sequences differ from those present within the complete operons. Theoretically, single 5S *rrn* loci may result either from the loss of the 23S and 16S *rrn* genes or from a partial duplication/acquisition of an *rrn* operon, the distinction between these scenarios not always being possible (**Figure 2**). Altogether, these incomplete *rrn* loci or *rrn* loss remain a minority (frequency < 7 % of the 127 strains analyzed).

The vast majority of the *rrn* operons (99.6 %) is oriented in the direction of the continuous replication, with very few cases of lagging strand orientation (*Streptomyces lincolnensis* LC G, *Streptomyces bingchenggensis* BCW1) (**Supplemental Table S4**). This could either illustrate i) a strong bias introduced by *rrn* expression on chromosome organization (to avoid polymerase collisions (Sinha et al. 2017)), as previously proposed (Lim, Furuta, et Kobayashi 2012), or ii) a positive selection of the genomic organization that limits large genomic deletions in case of recombination between *rrn* operons.

Altogether these observations indicated that *rrn* duplication/acquisition is the most frequently fixed phenomena directly involving *rrn* operons. Taking into account evolutionary events involving *rrn* operons (gain, loss, mutation) enriches the overall picture of *Streptomyces* chromosome evolution.

### Large pericentric inversions located in the vicinity of *rrn* operons

Driven by the observation of recombinant ‘[d/e]’ *rrn* operons (**Fig. 1D, Supplemental Fig. S2.A**), we examined the possible link between *rrn* operons and large genome rearrangements. We identified large rearrangements in the *Streptomyces* chromosome by comparing the order of genes from the core genome of each strain to that of *S. viridosporus* T7A ATCC 39115 (exactly ordered as the consensus). Most of them correspond to pericentric inversions, with only a few other cases (*e.g. S. lincolnensis* LC G, *Streptomyces ongiicola* HNM0071, *S. tirandamycinicus* HNM0039, *Streptomyces fungicidicus* TXX3120, *Streptomyces* sp. 11 1 2, *Streptomyces autolyticus* CGMCC0516 and *S. bingchenggensis* BCW 1) highlighting complex evolutionary scenarios (**Fig. 2**). Indeed, the group ‘O’ contains the species with the largest number of rearrangement events. Given the sparse distribution of genomes in this class, this may reflect missing steps in the proposed evolutionary scenario and this ‘O’ group may actually contain several clades. Accordingly, three strains of this group (*Streptomyces* sp. 11 1 2, *S. autolyticus* CGMCC0516 and *S. bingchenggensis* BCW 1) were considered too ambiguous to be included in the analysis presented below.

Core genome pairwise comparisons allowed the identification of 49 large rearrangements (> 200 kb) within the central compartment of 60 genomes, some of which likely occurred in the common ancestor of certain strains (**Fig. 2**, **Fig. 3**, **Supplemental Table S5**, **Supplemental Fig. S1 & S3**). We thereafter calculated the distances of each rearrangement end to the closest *rrn* operon (**Fig.3, Supplemental Table S5**). Although this method has a resolution limit related to the distance of core genes to *rrn* and rearrangement ends, 6 of these large rearrangements (env. 12 %) occurred less than 10 kb from an *rrn* locus, four of them corresponding to independent events of recombination between D and E *rrn* operons (**Supplemental Fig. S2**). Indeed, the distal core genes of these large rearrangements are located (at least on one side) at a median of less than 93 kb from an *rrn* operon (mainly belonging to Cc, Dd, Ee or [d/e] *rrn* categories, table of **Fig. 3.B**), a distance which represents 1.1 % of the mean genome size. Taken together, these results suggest that *rrn* operons, and especially those located around the origin of replication, constitute and/or are frequently close to recombination sites. This observation raises the question of mechanisms (other than homologous recombination between D and E *rrn* operons) by which *rrn* environments could favor the occurrence and/or fixation of pericentric inversions (see discussion).

**Figure 3:**
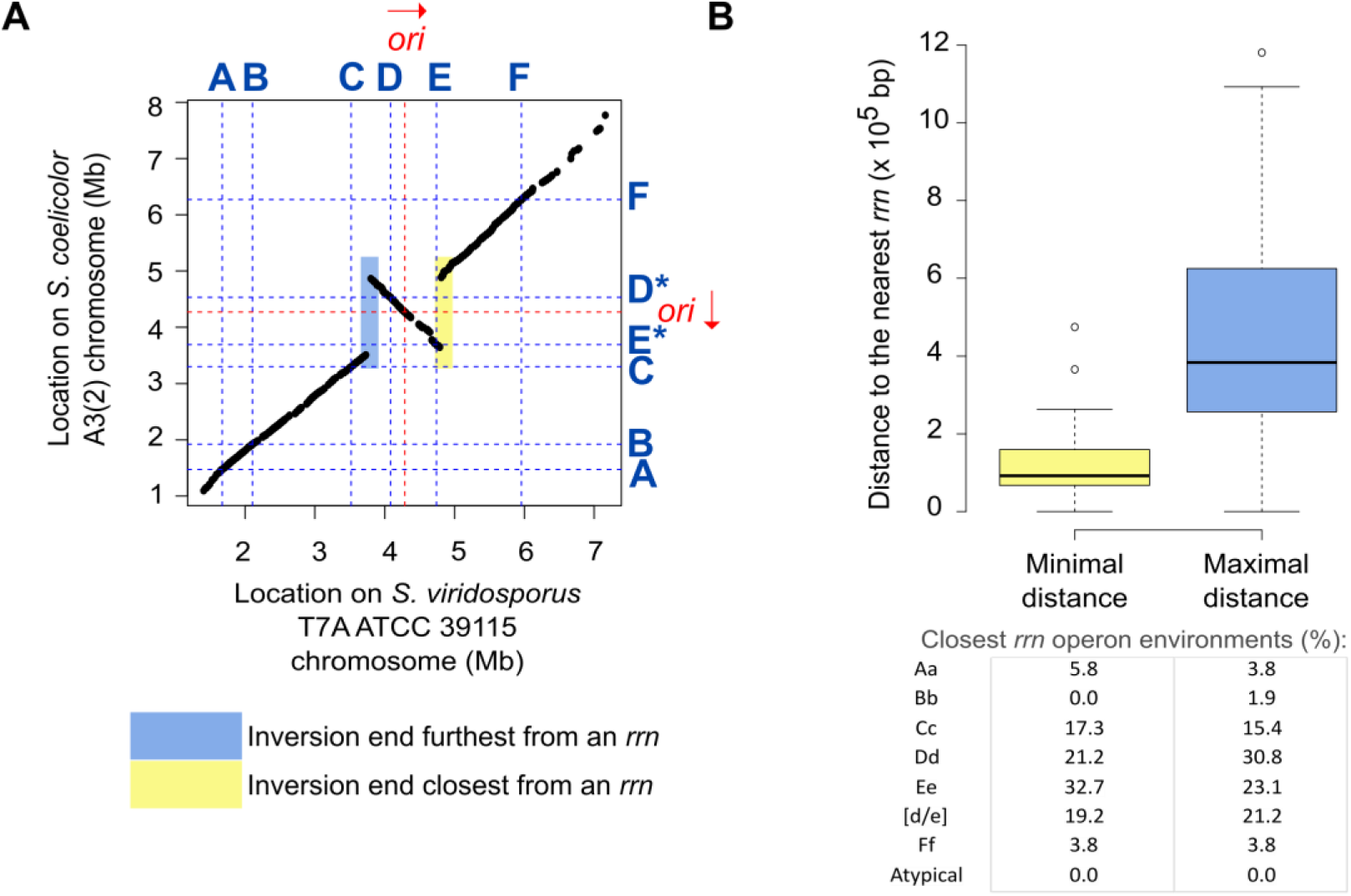
Distance from *rrn* loci of large rearrangements occurring in the central compartment. **A. Pairwise comparison of the core genomes of *S. coelicolor* A3(2) and *S. viridosporus* T7A ATCC 39115**, used as a reference for the consensus core genome order in *Streptomyces*. The identity of each *rrn* locus is specified using the nomenclature proposed in this study. The origin of replication (*oriC*) was defined regarding the position of the *dnaA* gene, the red arrow representing the orientation of this gene. The regions colored yellow and blue indicate the position of the closest and farthest ends from an *rrn* operon, respectively. These were subsequently used to calculate the minimum and maximum distances of the rearrangement ends to an *rrn* operon, with the resolution limit of the distance of these elements to the core genes. **B. Boxplot of minimal and maximal distance of the intra-chromosomal rearrangements to *rrn* loci.** When the same event was shared by several strains, the mean values (distances of both ends to the nearest *rrn* loci) were calculated so that each event (*n* = 49) is considered only once. The frequency of the nearest *rrn* operon category is indicated for both sides. The boxplots of both panels represent the first quartile, median and third quartile. The upper whisker extends from the hinge to the largest value no further than 1.5 * the inter-quartile range (IQR, *i.e*. distance between the first and third quartiles) from the hinge. The lower whisker extends from the hinge to the smallest value at most 1.5 * IQR of the hinge.

### Link between *rrn* operons and density in core genes

Driven by the observation that the sizes of the central compartment and the core region are highly correlated (**Fig. 1E**), we fit general linear models to model the interplay between core and *rrn* gene dynamics. We explored the predictability of the core region size depending notably on the location and number or *rrn* genes. Other possible explanatory variables were evaluated such as the distance to the origin of replication, the size of the chromosome and of the region encompassing all tRNA encoding genes (‘tDNA region’, **Supplemental Fig. S4A**), as well as the phylogenetic origin. We conducted both forward and backward regression approaches to select the best predictors. The best fitting model includes as explanatory variables: the sizes of the central compartment and of the chromosome, the maximal distance between the distal *rrn* operons and the origin of replication (‘d_max_*rrn*_ori’, **Fig.4A**), as well as the number of *rrn* genes (**Supplemental Fig. S4**). This overall model is statistically significant (*p* < 2.2.10^-16^) and suggests that the four explanatory variables included in the model explain approximatively 86 % of the core region variability (**Supplemental Fig. S4.B**). Accordingly the correlation between the observed size of the core region and the value predicted by this ANOVA model is very strong (*R* = 0.92, *p* < 2.2.10^-16^, Pearson correlation, **Fig.4A**), supporting the existence of an evolutionary relationship between the position of distal *rrn* genes and the core region.

**Figure 4:**
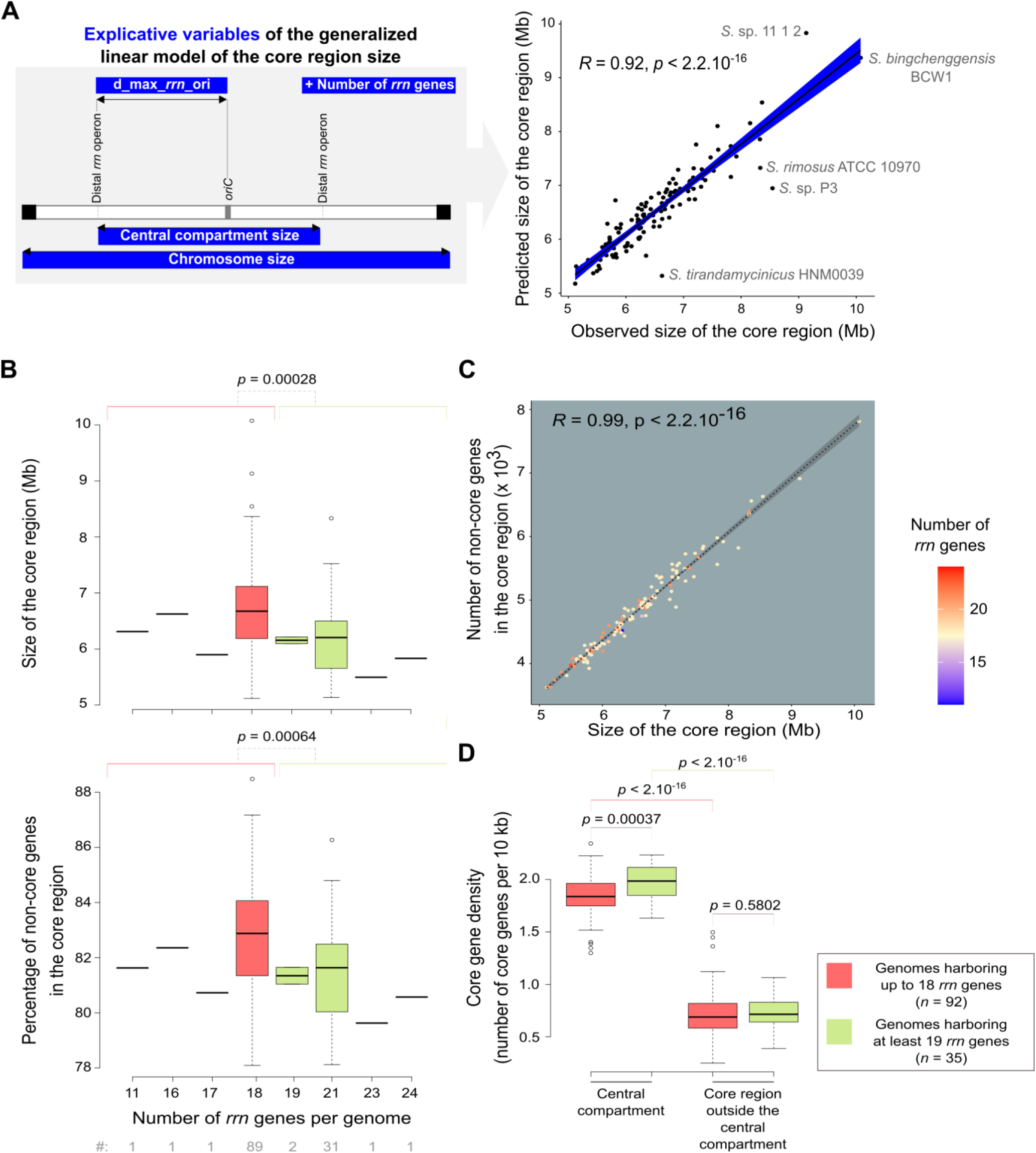
Interplay between *rrn* operons and core region dynamics. **A. Correlation between the predicted and observed core region sizes in the panel of 127 genomes of interest.** The explanatory variables are represented in the left panel. The *R* coefficient and *p* value of Pearson’s correlation tests were calculated with the whole set of genomes (*n* = 127). The names of the species are indicated for the genomes that present an unusual pattern in the diagnostic tests (**Supplemental Fig.S4**). In fact, these genomes belong to the ’O’ group, except for Streptomyces sp. P3 genome which has the most asymmetric organization (**Supplemental Table S1**). This suggests that the prediction model has limitations in the case of rather complex evolutionary scenarios or atypical genomic organizations and/or could help to identify them. **B. Boxplots presenting the size of the core region and its percentage of non-core genes depending on the number of *rrn* genes.** The boxplots represent the first quartile, median and third quartile. The upper whisker extends from the hinge to the largest value no further than 1.5 * the inter-quartile range (IQR, i.e. distance between the first and third quartiles) from the hinge. The lower whisker extends from the hinge to the smallest value at most 1.5 * IQR of the hinge. Outliers are represented (dots). The *p* values of two-sided Wilcoxon rank sum tests with continuity correction comparing the values observed in genomes harboring up to 18 *rrn* genes (red) in genomes harboring at least 19 *rrn* genes (green) are indicated. The number of genomes in each category (‘#’) is indicated below the graphs. **C. Scatter plot presenting the correlation between the core region size and the number of non-core genes**. The rho coefficients and *p* values of Spearman’s rank test were calculated with the whole set of genomes (*n* = 127). Each point corresponds to a genome colored according to the number of *rrn* operons it contains. **D. Boxplot presenting the core gene density depending on the number of *rrn* genes and the location inside or outside the central compartment.** The boxplot represents the same parameters as in panel B. The core gene density expressed as the number of core genes per 10 kb was calculated in the central compartment and in the core region located outside the central compartment (‘delta_core_*rrn*’ in the **Supplemental Fig.S4A**). The *p* values of two-sided Wilcoxon rank sum tests with continuity correction are presented.

The size of the tDNA region was not among the best predictors of the size of the core region (**Supplemental Fig. S4A**), emphasizing the importance of *rrn*-defined limits *per se*, independently of their role in the translation process. Moreover, this analysis supports the fact that the evolution of the core region size is determined by the number of *rrn* genes rather than their phylogenetic origin.

Remarkably, the increase in the number of *rrn* operons is correlated with a decrease in the core region size. This result highlights some kind of core region ‘densification’ (*i.e.* fewer non-core genes in the core region) correlated to the increase in the number of *rrn* operons (**Fig.4B & C**). This effect is in fact limited to the central compartment which harbors slightly more core genes per kb in genomes containing at least 19 *rrn* genes than in those containing up to 18 *rrn* genes (**Fig.4D**). The central compartment *per se* is 2.7-fold more dense in core genes (per size unit) than the core region located between the distal *rrn* and the last core genes (‘delta_core_*rrn*’ in the **Supplemental Fig.S4A**), as illustrated for *S. coelicolor* in **Figure 5A**. Overall, these results indicate that *rrn* operons define a central compartment characterized by a high density of core genes, a feature that tends to be more pronounced as the number of *rrn* genes increases.

**Figure 5:**
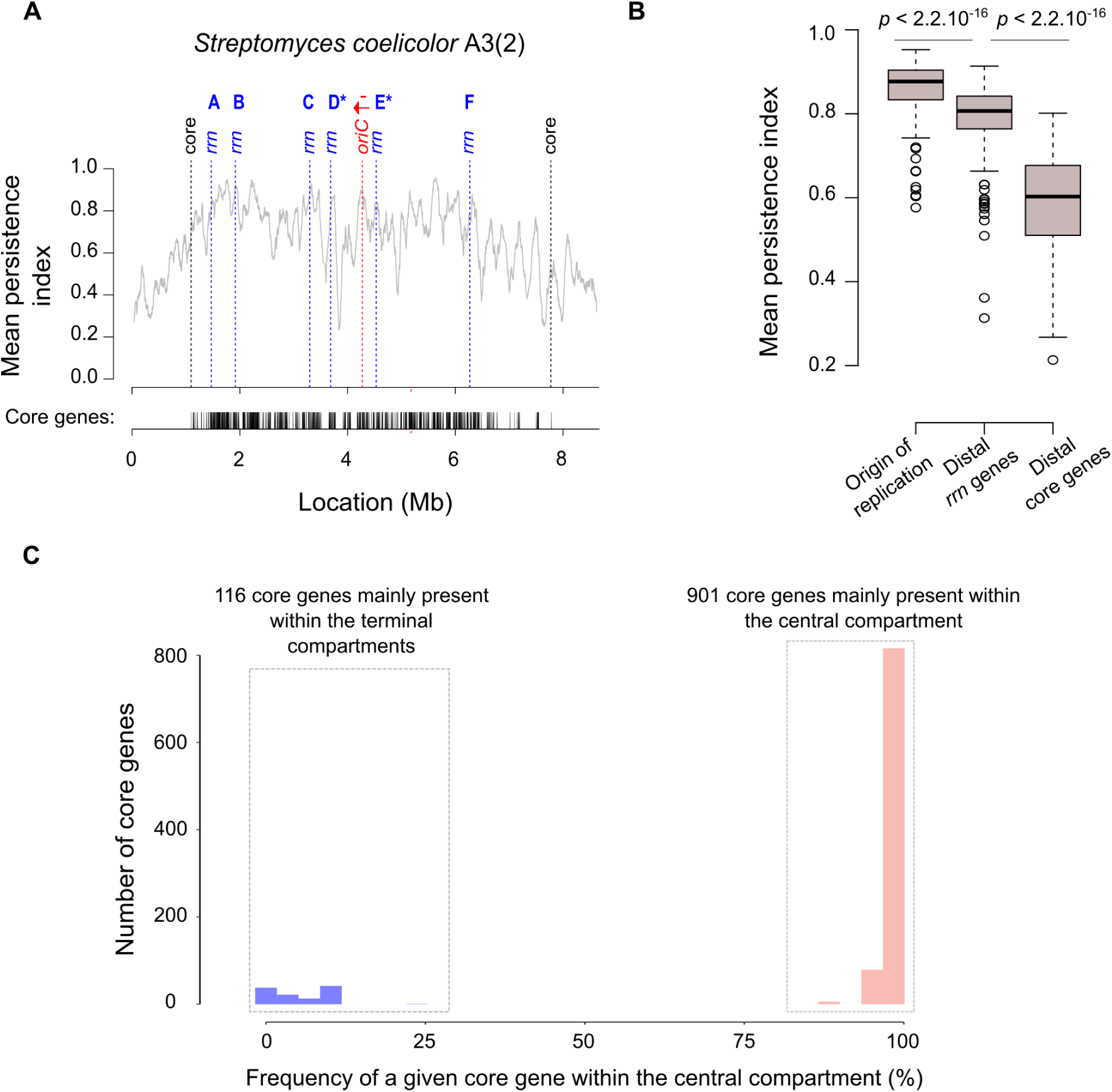
Gene persistence and core gene content within the central compartment. **A. Gene persistence along the chromosome of *S. coelicolor* A3(2).** The level of persistence along the chromosome is represented using a sliding window (81 coding sequences (CDSs), with 1 CDS steps). The positions of distal core genes (‘core’) and of all *rrn* operons are indicated by dashed lines. The density of the core genes is indicated below the graph. The identity of each *rrn* operon is specified using the nomenclature as proposed in this study (**Supplemental Figure S2**). The origin of replication (*oriC*) was defined regarding the position of the *dnaA* gene, the red arrow representing the orientation of this gene. **B. Mean gene persistence in the regions surrounding the origin of replication, the distal *rrn* and core genes.** The mean persistence was calculated within a window of 81 CDS centered on the genomic feature of interest. The boxplot is built as in Fig.4B. The *p* values of a two-sided Wilcoxon rank sum test with continuity correction comparing the mean persistence index at the vicinity of the origin of replication (*n* = 127), and of the distal *rrn* (*n* = 254) or core (*n* = 254) genes, are indicated. **C. Distribution of the core genes within and outside the central compartment.**

### The gene content within the central compartment is remarkably stable

We then explored the qualitative gene content in the central compartment. We previously reported the interest of using the gene persistence index to evaluate the level of gene conservation along the *S. ambofaciens* ATCC 23877 chromosome (Virginia S. Lioy et al. 2021). This index corresponds to the frequency of a given gene in a set of complete genomes of interest. In the present study, we enlarged this analysis to all the genomes of our panel (**Supplementary Fig. S5**). As illustrated by the representative example of the *S. coelicolor* A3(2) chromosome (**Fig. 5A**), beyond the distal *rrn* operons, there are generally a few core genes and then gene persistence decreases sharply. More precisely, the gene persistence index fluctuates along the genome, reaching the highest levels within the central compartment, especially near the origin of replication. The *rrn* operons, especially the ones located in a canonical core gene environment, usually localize with a persistence peak superior to 0.8, except for the ‘D’ *rrn* category (**Supplemental Fig. 6**). Another notable exception involves the *rrn* operons in the lagging orientation in *S. bingchenggensis* BCW1 (**Supplemental Fig. S5**).

**Figure 6:**
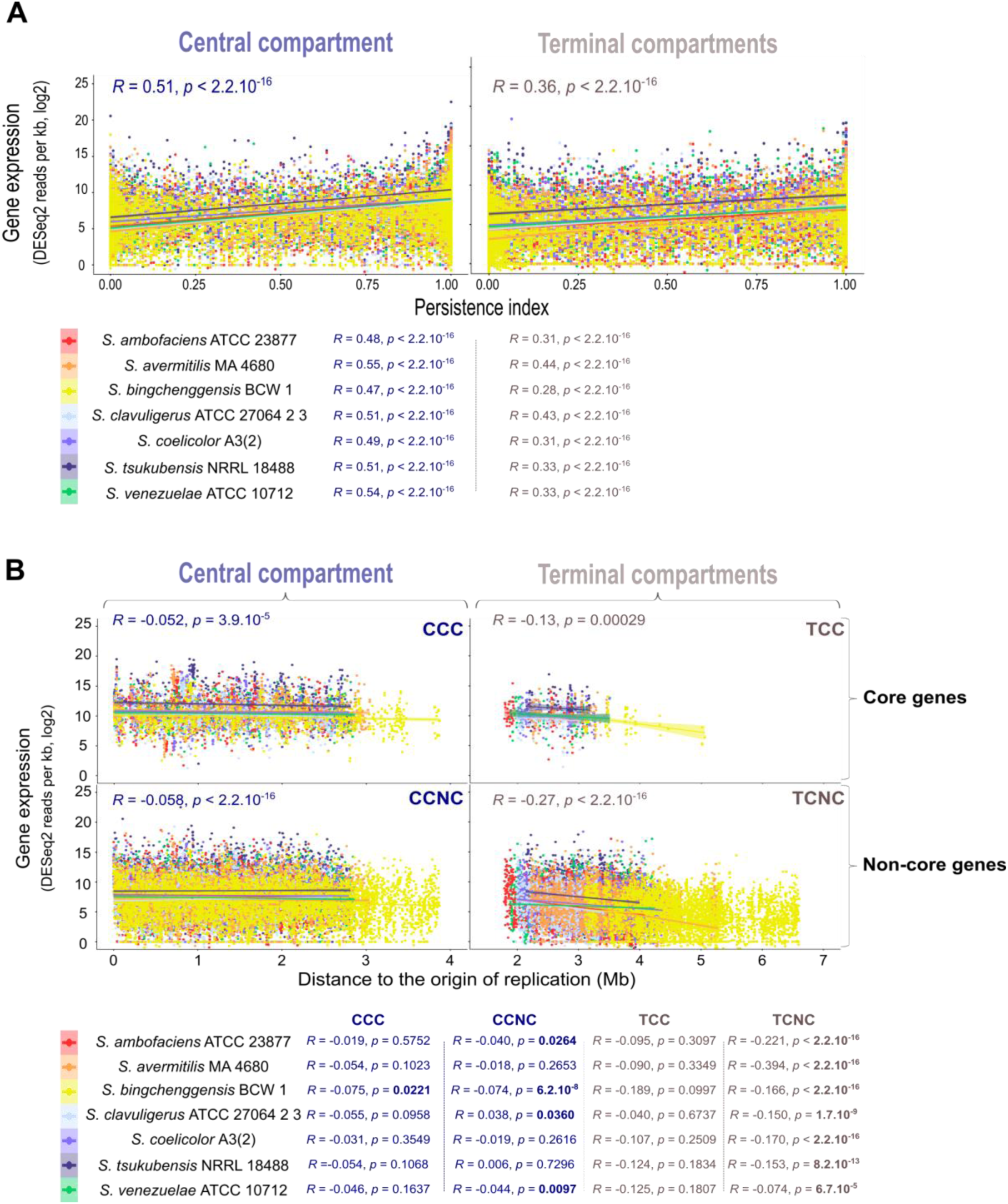
Gene expression level depending on the location inside or outside the central compartment. **A. Correlation between gene expression and gene persistence in the central (left) and terminal (right) compartments.** Gene transcription (in sense orientation) during the trophophase corresponds to the log 2 of the number of DESeq2 normalized reads per kb. The correlations were analyzed by a Spearman’s rank correlation test, performed on the whole transcriptomes (values indicated on the graph) or for each species individually (values indicated below the graph). **B. Correlation between gene expression and the distance to the origin of replication in the central (left) and terminal (right) compartments.** Core (top) and non-core (bottom) gene transcription (in sense orientation) were measured during the trophophase and expressed as the log 2 of the number of DESeq2 normalized reads per kb. The correlations were analyzed by a Spearman’s rank correlation test, performed on the whole transcriptomes (values indicated on the graph) or for each species individually (values indicated below the graph). Statistically significant *p*-values are written in bold. Abbreviations: CCC (central compartment core genes); CCNC (central compartment non-core genes); TCC (terminal compartment core genes); TCNC (terminal compartment non-core genes).

Importantly, gene persistence around the distal *rrn* operons is in general higher than at the limits of the core region (**Fig. 5B**). This result is in accordance with the lower core gene density observed in the core region outside the central compartment (**Fig.4D**).

We previously reported (Virginia S. Lioy et al. 2021) that whereas the size of the central compartment represents a little more than half of the entire chromosome (updated values: 57.7 ± 5.7 %, standard deviation, *n* = 127), the percentage of core genes within the central compartment is remarkably high and stable (updated values: 88.5 ± 3.0 %, standard deviation, *n* = 127). In this study, we have extended this observation by analyzing the qualitative composition of the central compartment in core genes. Interestingly, a set of 901 core genes is almost always located in the central compartment of the *Streptomyces* genomes we analyzed (**Fig. 5C**). This set of genes, further named CCC genes for ‘central compartment core’ genes, are enriched in genes encoding key cellular processes (‘private goods’) related, for instance, to central metabolism and translation (**Supplemental Figure S7.A**). The genes of the core genome that are generally located in the terminal domains, further termed TCC genes for ‘terminal compartment core’ genes, are enriched in only a few functional categories, related mostly to lipid metabolism (**Supplemental Figure S7.B**).

Altogether, these results indicate that the central compartment constitutes a specific evolutionary entity and suggest that the distal *rrn* operons constitute pertinent limits to describe a functional central compartment in the *Streptomyces* genome.

### High levels of transcription in the central compartment of *Streptomyces* genomes

To examine the central compartment from a functional point of view, we compared gene expression inside and outside this region over growth. We thus analyzed the available transcriptome data during metabolic differentiation of seven *Streptomyces* species form the clade 1 [*S. ambofaciens* ATCC 23877(Virginia S. Lioy et al. 2021), *S. avermitilis* MA 4680 (Kim et al. 2020), *S. coelicolor* A3(2) (Jeong et al. 2016)], clade 2 [*S. clavuligerus* ATCC 27064 2 3 (Kim et al. 2020)*, S. tsukubensis* NRRL 18488 (Kim et al. 2020), *S. venezuelae* ATCC 10712 (Gehrke et al. 2019)], and group ‘O’ [S*. bingchenggensis* BCW 1/BC-101-4 (P. Jin et al. 2020)].

For all species, we observed a positive correlation between gene persistence and expression (**Fig.6A**, **Supplemental Fig. S8**), as previously reported in *S. ambofaciens* ATCC 23877 (Virginia S. Lioy et al. 2021) and other bacteria (Acevedo-Rocha et al. 2013). As expected, this positive correlation is the highest during the trophophase, *i.e.* during vegetative growth which is associated with the lowest expression of variable regions belonging to the specialized metabolite biosynthetic gene clusters (SMBGCs) (Virginia S. Lioy et al. 2021) (**Supplemental Fig. S9**). We then compared the strength of this correlation as a function of whether the genes were located inside or outside the central compartment. Interestingly, for all strains, the positive correlation between gene persistence and transcription, measured by the Rho Spearman coefficient, is higher (≍ + 30 %) in the central compartment than in the terminal compartments (**Fig. 6A**). Moreover, during the trophophase, genes are more expressed in the central compartment, regardless of their category (core, non-core or SMBGC genes), in most strains (**Supplemental Fig. S9**). Remarkably, this higher expression in the central compartment is conserved in all strains for non-core genes and SMBGCs during the idiophase (*i.e.* after the metabolic differentiation leading to specialized metabolite/idiolyte production) (**Supplemental Fig. S9**). Altogether these results indicate that the central compartment delimitates a region associated with increased transcription (and/or RNA stability) compared to the rest of the genome.

We therefore consider the possibility that this effect could be related to a dose effect, gene copy number being higher close to the origin of replication in actively replicated chromosomes. We thus calculated the correlation between the level of expression and the distance to the origin, according to the localization of the genes inside or outside the central compartment (**Fig. 6B**). Remarkably, this dose effect is negligible in the central compartment, whereas the distance from the origin of replication is associated with a decrease in the quantity of transcripts produced from the terminal compartments, especially from non-core genes (**Fig. 6B**).

These results indicate that the central compartment is associated with a higher level of gene expression, regardless of either gene persistence or distance to the origin of replication in the seven strains we analyzed, even in *S. bingchenggensis* BCW 1/BC-101-4 which presents an atypical central compartment core gene content as a result of extensive chromosomal rearrangement (‘*rrn* a2 F* c2 d3 E b1* *dnaA*+’ configuration, **Supplemental Fig.S1**). Indeed, 85 core genes usually located in the terminal compartments are present within the central compartment of *S. bingchenggensis* BCW 1, 46 core genes being relocated from the central to the terminal compartments in this strain. The pattern of correlations between gene expression and distance to the origin is globally the same in *S. bingchenggensis* BCW 1 as in the other strains examined (**Fig.6B**). These data thus strongly suggest that the physical location of genes in the central compartment *per se* is a major determinant of their higher expression.

## Discussion

This study reports for the first time the in-depth analysis of *rrn* operon dynamics in a panel of 127 *Streptomyces* species. We gather a series of observations supporting that *rrn* operons are part of a core skeleton and can be distinguished based on their core gene environment. This allows us to propose a new genome nomenclature based on *rrn* operons and *dnaA* orientation. On this basis, we defined a canonical organization (’*rrn* ABCDEF *dnaA+*’) carried by 42% of species, and a consensus order of genes from the core genome, perfectly conserved in some contemporary strains such as *S. viridosporus* T7A ATCC 39115. The pairwise comparison of the genes of the core genome and *rrn* organization of this species to other strains/species of our panel (**Supplemental Fig. S2 & S3**) allow us to propose an evolutionary history of the central compartment of the *Streptomyces* chromosome (**Fig. 2**).

Interestingly, *rrn* operons, especially centrally located (*rrn* C, D and E, **Fig. 7**), tend to be close to rearrangement borders (**Fig. 3**), some being directly involved in homologous recombination (**Fig.2**). This suggests that recombination events in the vicinity of the pericentric *rrn* operons are more fixed and/or more frequent. We recently published *S. ambofaciens*’ chromosome conformation during metabolic differentiation, and we showed that, in exponential phase, these *rrn* operons form sharp boundaries (Virginia S. Lioy et al. 2021), reflecting their high transcription. Moreover, they are localized in a central region that appears to be enriched in contacts around the origin (Virginia S. Lioy et al. 2021) (**Fig.7**).

**Figure 7:**
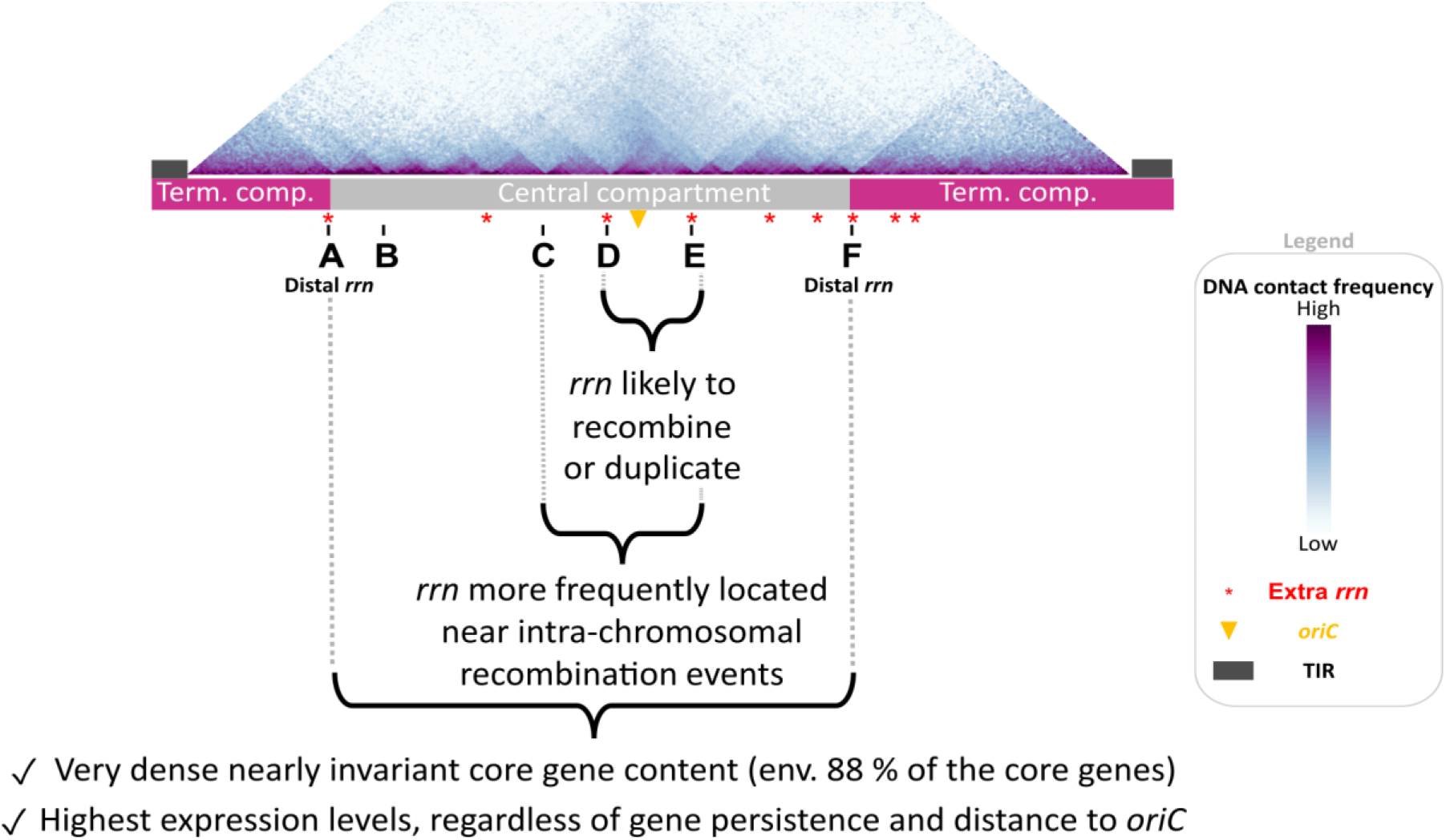
Main properties of the *Streptomyces* chromosome in relation to *rrn* operons. The schematic representation of the chromosome is shown to scale using *S. ambofaciens* ATCC 23877 as reference. The DNA contacts along its chromosome in exponential phase have been previously reported (Virginia S. Lioy et al. 2021). The relative positions of extra *rrn* copies in other *Streptomyces* genomes are indicated in red. Abbreviations: ‘term. comp.’ = terminal compartment; TIR = terminal inverted repeats.

Inter-arm contacts are favored by the SMC machineries in the *S. venezuelae* ATCC 10712 chromosome (Szafran et al. 2021). Thus we speculate that the intra-chromosomal recombination observed in these regions could perhaps result from the spatial proximity of these regions and/or the occurrence of DNA breaks related to the strong structural tensions exerted in this genomic region (boundaries related to strong transcription, loop exclusion by SMC near the origin, and/or DNA replication progression in actively replicating cells).

Interestingly, Fleurier *et al*. (Fleurier et al. 2022) recently demonstrated that transcription-dependent DNA replication blockages at overexpressed *rrn* operons can result in DNA breakage and cell death. Consequently, double strand breaks may be more frequent at the vicinity of *rrn* operons and stimulate pericentric inversions. This raises the question of whether this process is widespread in bacteria since recombination between *rrn* operons (Jumas-Bilak et al. 1998; Klockgether et al. 2010; Sato et Miyazaki 2017; Irvine et al. 2019; Gifford, Dasgupta, et Barrick 2021) or domain boundaries at the *rrn* operons (V. S. Lioy et al. 2018; Le et Laub 2016; Böhm et al. 2020; Marbouty et al. 2015; Wang et al. 2015) have also been reported in other bacteria.

Overall, our study provides a model of the core genome size based on predictors that can be easily collected from the knowledge of the genome sequence (chromosome size, position and number of *rrn* genes, position of the origin of replication - **Fig.4A, Supplemental Fig. S4**). Moreover, the location of the distal *rrn* operons can be easily implemented in dedicated software as an additional criterion for predicting regions of interest in the search for antibiotic-encoding SMBGCs, which tend to be acquired by horizontal gene transfer and enriched in the terminal and variable regions.

Our results confirm that *rrn* operons co-evolve closely with the core genome, their number being an important determinant to explain its dynamics (**Fig. 4**). The ancestor of the *Streptomyces* genus probably had 6 *rrn* operons, the acquisition of additional *rrn* being fixed at least 13 times independently during genus evolution (**Fig. 2**). Interestingly, the acquisition of an extra copy of *rrn* leads to a decreased propensity of the core region to contain non-core genes (**Fig. 4**). This could reflect less acquisition of foreign DNA and/or the displacement of poorly conserved genes towards the ends. Overall, these results are consistent with the observation that the central region of the *Streptomyces* chromosome is more constrained than the arms, gene flux and shuffling operating more intensively in the latter (Lorenzi et al. 2021).

In parallel, we show that the *rrn* are close to highly persistent gene environments and constitute approximate limits beyond which the persistence of genes tends to decrease rapidly (**Fig.5, Supplemental Fig. S5**). In fact, while the concept of genomic compartmentalization of the *Streptomyces* chromosome is not new, defining the exact barriers has remained a challenge. Here, we propose to consider distal *rrn* as structural limits since they delimitate a highly conserved and expressed region, harboring 88.5 % of the core genome, almost always composed of the same set of core genes.

This central compartment has functional consequences. Indeed, the correlation between gene persistence and expression is stronger within than outside this region. Higher transcription propensity at the vicinity of *rrn* operons has also been reported in *Escherichia coli* (Scholz et al. 2019). Moreover, the expression of the genes located within the compartment is independent of a dose effect, *i.e.* it is not correlated with the distance to the origin of replication. These observations suggest that the central compartment may constitute a specific molecular environment.

HiC experiments performed in eukaryotes and some archaea revealed the existence of two compartments, namely A/B type associated to high and low transcription, respectively (Takemata et Bell 2021; Lieberman-Aiden et al. 2009). Remarkably, a hub-like structure with colocalized genes involved in ribosome biogenesis has been identified in the genome of some archaea (Takemata et Bell 2021). At present, bacteria with circular genomes are considered to lack such compartments. The present study supports the existence of a bacterial nucleolus-like environment, constituting a molecular environment/compartment prone to transcription.

Collectively, these results indicate a link between evolutionary processes, including genome compartmentalization and *rrn* operon dynamics in *Streptomyces*. Our study raises the question of whether *rrn* operons are directly involved in genome compartmentalization, for example by protecting the core from terminal recombination, or whether they are just proxies for the evolution of a core skeleton. We believe this study brings new insights into the rules governing chromosome spatial organization, expression, recombination and evolution.

## Methods

### Genome annotation and orthology assignment

The set of genomes used in this study consists of 125 genomes whose selection was previously described (Lorenzi et al. 2021), to which we added 2 genomes of model strains (*S. venezuelae* ATCC 10712 and *S. albidoflavus* J1074) for which genomic data are available. All genomes (**Supplemental Table S1**) were automatically annotated on the RAST server (Aziz et al. 2008; Overbeek et al. 2014) using the RAST Classic pipeline (FIGfam version: release 70) to standardize annotation protocols, a key step for the subsequent assignment of orthology relationships. For each pair of genomes, orthologs were identified by BLASTp reciprocal best hits (BBH) (Fang et al. 2010; Tatusov, Koonin, et Lipman 1997; Overbeek et al. 1999) with at least 40% identity, 70% coverage (based on the shortest sequence) and an E-value of less than 10^-10^. Each orthologous group was identified by a number using a graph approach based on a simple linkage method (**Supplemental Figure S10**). The core-genome corresponds to the set of orthologs (1017) present in all the genomes of our dataset and forming a clique (**Supplemental Figure S10**). The annotation of the whole genomes is available in **Supplemental Table S6**. The SMBGCs and prophages were predicted using AntiSMASH5.0 (Blin et al. 2019) and PHASTER (Arndt et al. 2016), respectively.

### Phylogenetic analysis

For each strain, the protein sequences of the 1017 genes of the core-genome were retrieved. The sequences were concatenated and aligned with MAFFT (Katoh 2002; Katoh et Standley 2013) (v7.490). The multiple alignment (441,390 positions) was then subjected to RAxML-NG (Kozlov et al. 2019) with the LG substitution model for maximum-likelihood-based tree inference. Fifty bootstrap replications were performed. The phylogenetic tree was represented using MEGA X software (Kumar et al. 2018).

### Average nucleotide identity (ANIb) computation

The average nucleotide identity between query and reference genomes was calculated by using the BLASTn algorithm (ANIb) (Goris et al. 2007). First, the query genome was fragmented into 1,000 consecutive parts, which were then each aligned to the reference genome sequence using BLASTn (v2.11.0+) (Altschul 1997). The ANIb score is the average value of the percentages of nucleotide identity of the query fragments with a positive match to the reference genome (alignment greater than 70 % with at least 30 % of nucleotide identity) (Goris et al. 2007). Because the ANIb score is not reciprocal (i.e. the ANIb score of genome A *versus* genome B may be slightly different from the ANIb score of genome B *versus* genome A), we used the average of the two reciprocal values as the final score.

### Core gene consensus order building

The consensus order of core genes was determined from the analysis of the 52 *Streptomyces* strains that harbor the most frequent *rrn* configuration, termed canonical (‘*rrn* ABCDEF *dnaA*+’). A rank (from 1 to 1017) was assigned to each gene within each strain. The most frequent rank was attributed to each core gene. An ambiguity between ranks 498 and 499 required a dedicated analysis of the most frequent gene order on the corresponding area. The scripts used to conduct this analysis are available in **Supplemental File 1**.

### The *rrn* nomenclature rules

The core gene neighborhood of each *rrn* operon (nearest core gene and its previous and next core genes) was determined for all genomes in the panel (detailed in **Supplemental Fig. S2**). The order of the consensus core genes was used to determine the *rrn* neighborhood order described as ‘sense’. If the order of the genes was in the other orientation, an asterisk was added to represent the ‘antisense’ orientation. The six most frequent *rrn* core gene environments were designated from ‘A’ to ‘F’, whereas the other *rrn* core gene environments were named from ‘g’ through ‘k’ in lower case with a number, to indicate their non-canonical nature. The same letter is kept when at least one core gene is in common between two *rrn* environments. Finally, the *dnaA* gene orientation was included in the nomenclature, as a proxy for the orientation of the replication origin. In this context, ‘*dnaA*+’ and ‘*dnaA*-’ refer to the sense (start codon located before the stop codon) or antisense orientation of the *dnaA* gene, respectively. Some sequences released from the databases were oriented in an inverted manner with respect to the consensus core order determined in this study. We conserved the orientation provided by the databases for the analyses presented in this paper, but considered the genome configuration in the appropriate orientation (*e.g. ’rrn* F*E*D*C*B*A* *dnaA*-‘, considered as equivalent to ‘*rrn* ABCDEF *dnaA*+’). To decide whether a sequence from the database is in the same orientation as the reference consensus used in this study (*Streptomyces viridosporus* T7A ATCC 39115), it is necessary to consider the results of pairwise comparison presented in **Supplemental Figure S1**: when the diagonal starts at the bottom left and ends at the top right, it means that the genome sequence available in the databases is oriented as in the consensus, and that the *rrn* configuration shown below each graph can be directly transposed onto the graph. If not, the sequence is in the opposite direction. In this case, the consensus should be reversed when transposed on the graph (e.g. ’*rrn* ABCDEF *dnaA*+’ becomes ’*rrn* F*E*D*C*B*A* *dnaA*-’).

### Core gene-based identification of large genome rearrangements within the central compartment

The *Streptomyces viridosporus* T7A ATCC 39115 core genome was used as a reference in this analysis. The difference (‘delta_VIRO’, in bp) between the position of the core genes in the central compartment of each strain and the reference strain was calculated, and then the difference between the ‘delta_VIRO’ values of successive genes (‘delta_VIRO_delta’) within each strain. Positions for which the ‘delta_VIRO_delta’ values were greater than 200 kb were selected. Manual curation was performed based on pairwise comparisons of core genomes (**Supplemental Fig. S1 and S3**). Rearrangements identified in multiple strains sharing a common ancestor were considered only once, taking the average values of size and distance to the nearest *rrn* operons. In some cases, the exact position of the rearrangement was determined by comparison with a more closely related species (e.g. *S. koyakasensis* versus *S. albidoflavus*). In order to include only rearrangements whose identification was not ambiguous, three strains with too complex evolutionary scenario (*S.* sp. 11 1 2, *S. autolyticus* CGMCC0516 and *S. bingchenggensis* BCW 1) were excluded from this analysis. Thus, there were probably more rearrangements than proposed in the scenario (especially in the group ‘O’). All data are available in **Supplemental Table S5**.

### Modeling

We fit linear regression models using the ‘lm’ function of R software (R Core Team 2021) to explore the predictability of the core region size (*n* = 127) using as explanatory variables the sizes and distances represented in **Fig.4A** as well as the number of *rrn* genes and the phylogenetic origin of the strains. We conducted both forward and backward regression approaches to select the best predictors. The script associated with this approach is detailed in the **Supplemental File 1**. The best fitting model according to Akaike Information Criterion (AIC) was checked visually using diagnostic plots (residuals *vs*. fitted values, and QQ plots to check normality) (**Supplemental Fig.S4**).

### GO enrichment analysis

The GO enrichment analysis was performed on the CCC and TCC genes of *S. coelicolor* A3(2), which is the most studied and therefore annotated *Streptomyces* genome. The SCO and GO annotations of its core genome are detailed in the **Supplemental Table S9**. The g:Profiler g:GOst software (https://biit.cs.ut.ee/gprofiler/gost, version e105_eg52_p16_e84549f) was used on line after uploading a GMT file corresponding to the *S. coelicolor* A3(2) complete GO annotation (**Supplemental File 2**).

### Transcriptome analyses

RNA-seq data were retrieved from the NCBI Gene Expression Omnibus (GEO, https://www.ncbi.nlm.nih.gov/geo/) under the following accession codes: GSE162865 (*S. ambofaciens* ATCC 23877) (Virginia S. Lioy et al. 2021), GSE118597 (*S. avermitilis* MA 4680) (Kim et al. 2020), GSE147644 (*S. bingchenggensis* BCW1/BC-101-4) (P. Jin et al. 2020), GSE69350 [*S. coelicolor* A3(2)] (Jeong et al. 2016), GSE128216 (*S. clavuligerus* ATCC 27064 2 3) (Kim et al. 2020), GSE97637 (*S. tsukubensis* NRRL 18488) (Kim et al. 2020), GSE115439 (*S. venezuelae* ATCC 10712) (Gehrke et al. 2019). STAR software (Dobin et al. 2013) (v2.5.4) was used for mapping RNA-seq to the reference genome containing only one terminal inverted repeat (TIR). This avoids any biases with multiple mapping within the duplicated extremities of the genome (since the two TIR sequences are indistinguishable). We used the *featureCounts* program (Liao, Smyth, et Shi 2014) (v2.0.1) to quantify reads in the sense-orientation. SARTools (Statistical Analysis of RNA-Seq data Tools, v1.6.3) DESeq2-based R pipeline (Love, Huber, et Anders 2014; Varet et al. 2016) was used with default parameters for systematic quality controls, normalization and detection of differentially expressed genes in each strain considered independently. The first time point was used as the reference condition. The DESeq2 counts were normalized on gene size (DESeq2 reads per kb) in each growth condition (**Supplemental Table 7**). Only protein-coding genes were considered to generate the data presented in **Fig. 6** and **Supplemental Fig. 8**.

### Statistical procedure and codes

Statistical analyses were performed with R software (R Core Team 2021). The scripts used to annotate the genome (orthologous groups, core genome, etc.) and to calculate gene persistence and the ANIb are available on Github (Jnlorenzi 2022). The scripts used to conduct the data analyses are detailed in the **Supplemental File 1**.

### Data Access

The **Supplemental Table 1** contains a precise description of all the genomes (including accession numbers, *rrn* configuration, as well as values for distances or numbers of genes of interest used in this study). The ANIb scores are available in the **Supplemental Table 2**. The species closest to the consensus/ancestor in terms of core gene order are listed in the **Supplemental Table 3**. The **Supplemental Table 4** provides the annotation of *rrn* operons in all genomes. The **Supplemental Table 5** lists the large rearrangements identified in the central compartment in 62 genomes. The complete (core and non-core) CDS annotation of all genomes is detailed in the **Supplemental Table 6**. The RNA-seq raw data used in this study are available on the NCBI Gene Expression Omnibus (GEO, https://www.ncbi.nlm.nih.gov/geo/) under the following accession codes: GSE162865 (*S. ambofaciens* ATCC 23877) (Virginia S. Lioy et al. 2021), GSE118597 (*S. avermitilis* MA 4680) (Kim et al. 2020), GSE147644 (*S. bingchenggensis* BCW1/BC-101-4) (P. Jin et al. 2020), GSE69350 [*S. coelicolor* A3(2)] (Jeong et al. 2016), GSE128216 (*S. clavuligerus* ATCC 27064 2 3) (Kim et al. 2020), GSE97637 (*S. tsukubensis* NRRL 18488) (Kim et al. 2020), GSE115439 (*S. venezuelae* ATCC 10712) (Gehrke et al. 2019). The RNA-seq data normalized by DESeq2 are available in the **Supplemental Table 7**. The **Supplemental Table 8** contains additional data on *rrn* operons, useful for easily reproducing some of the analyses described in the **Supplemental File 1**. The **Supplemental Table 9** contains the core genome SCO and GO annotations. The **Supplemental File 2** is a GMT file required to perform GO enrichment analysis using g:Profiler.

## Author Contributions

Supervision & design of the analyses: SBM; Bioinformatic analyses and script development: JNL, SBM; Writing – Original draft: SBM; Writing – Reviewing and Editing: all authors; Funding acquisition: JLP, SBM.

## Acknowledgments

We acknowledge Hoda Jaffal for fruitful discussions and advice, Linda Sperling for proofreading the manuscript, and ANR (ANR-21-CE12-0044-01/STREPTOMICS) for funding. SBM acknowledges G. Lelandais, P. Poulain and B. Cosson for their valuable teaching and advice during the DUO degree. SBM also thanks C. Dillmann, T Mary-Huard and other teachers of the modeling course of the doctoral school ABIES.

