## Supplementary material for "Ribosomal RNA operons define a central functional compartment in the *Streptomyces* chromosome": Figure S1.pdf

Fig.S1 (1/15)

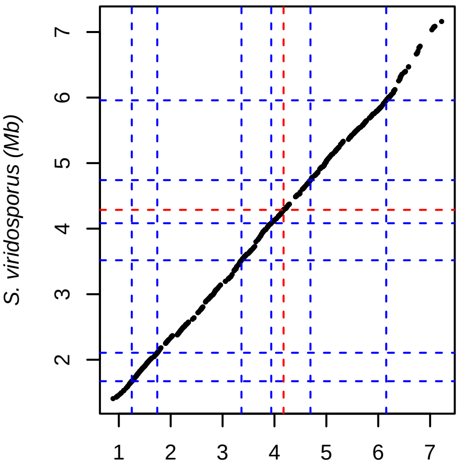


S. actuosus ATCC 25421 (Mb)
A B C D E F dnaA+
6 rrn loci, Clade_1

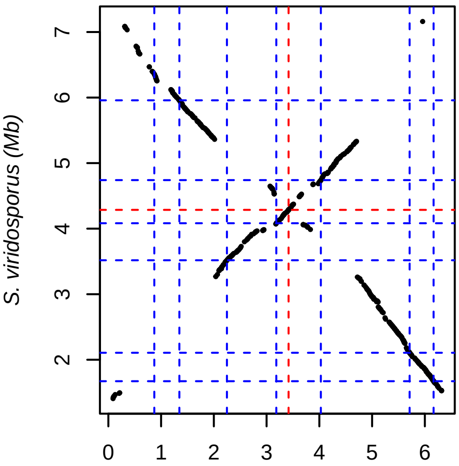


S. albidoflavus J1074 (bp)
A B E* D* C* F k1 dnaA−
7 rrn loci, Clade_1

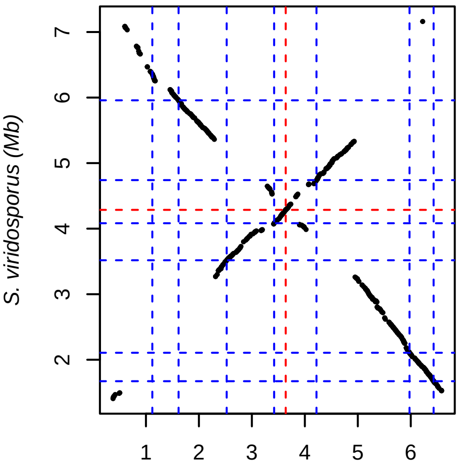


S. albidoflavus SM254 (bp)
A B E* D* C* F k1 dnaA−
7 rrn loci, Clade_1

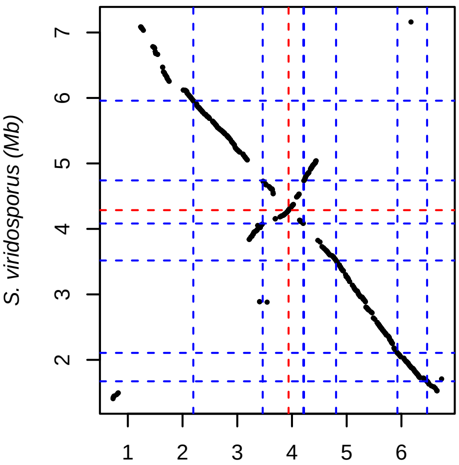


S. albireticuli MDJK11 (bp)
A B C [d/e2]* [d/e4] [d/e3]* F dnaA−
7 rrn loci, Group_O

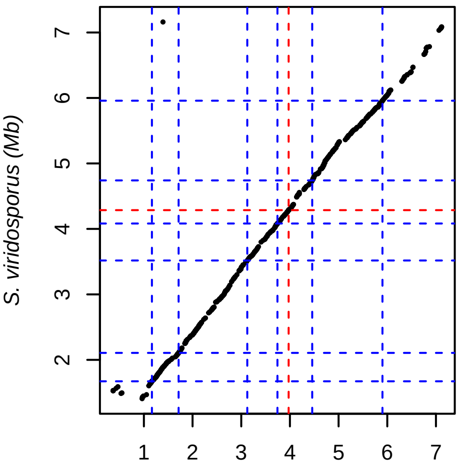


S. alboniger ATCC 12461 (Mb)
A B C D E F dnaA+
6 rrn loci, Clade_1

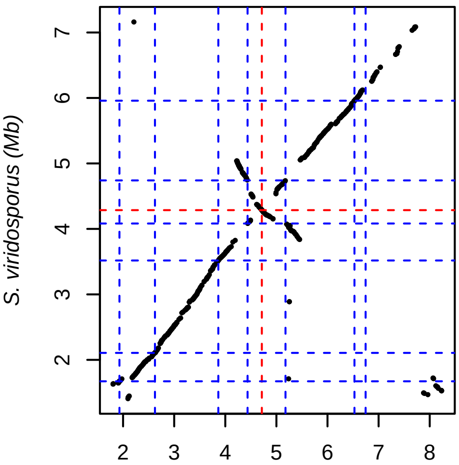


S. albulus CK 15 (bp)
A B C [d/e5] [d/e1]* F j1 dnaA−
7 rrn loci, Group_O

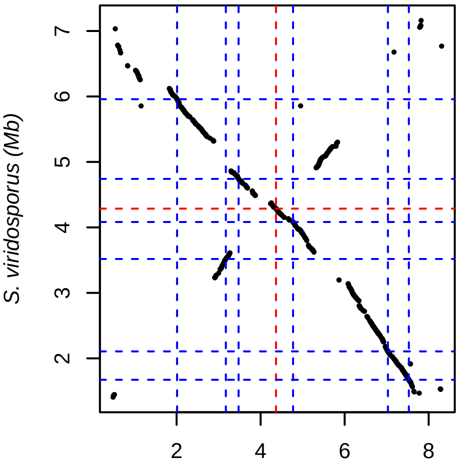


S. albus ZD11 (Mb)
A B D E C* F dnaA+
6 rrn loci, Clade_1

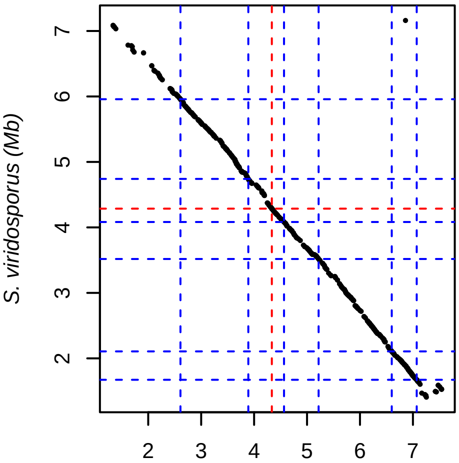


S. alfalfae ACCC40021 (Mb)
A B C D E F dnaA+
6 rrn loci, Clade_1

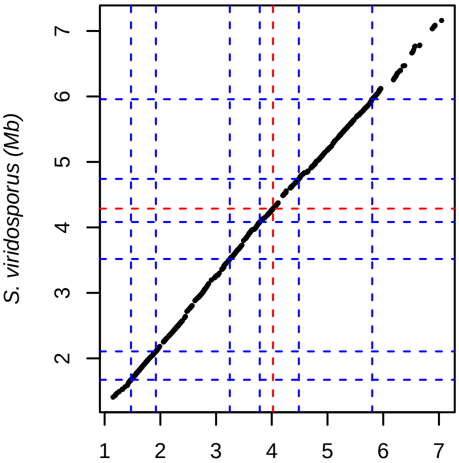


S. ambofaciens ATCC 23877 (Mb)
A B C D E F dnaA+
6 rrn loci, Clade_1

Fig.S1 (2/15)

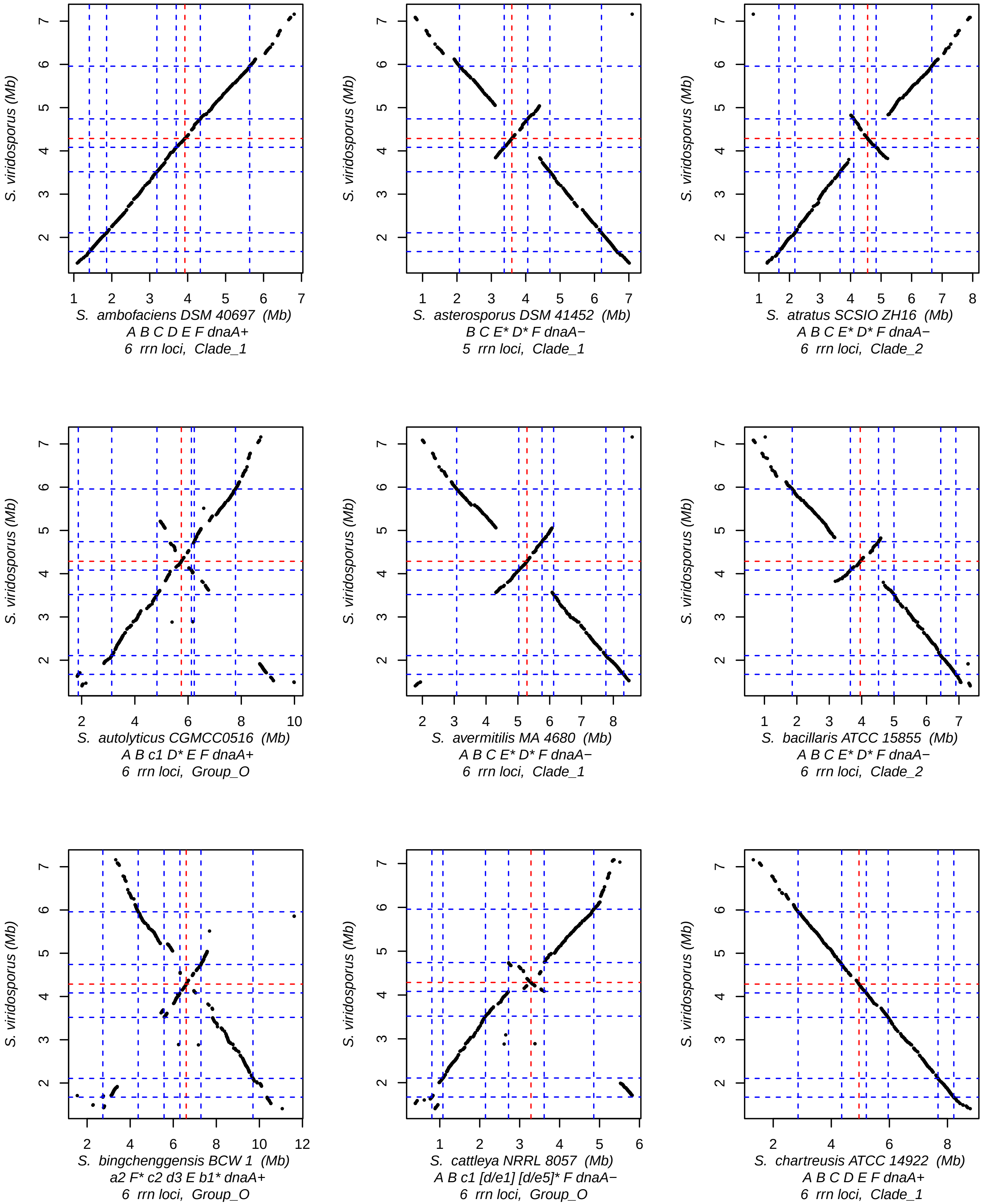
S. viridosporus (Mb)
S. ambofaciens DSM 40697 (Mb)
A B C D E F dnaA+
6 rrn loci, Clade_1
S. asterosporus DSM 41452 (Mb)
B C E* D* F dnaA−
5 rrn loci, Clade_1
S. atratus SCSIO ZH16 (Mb)
A B C E* D* F dnaA−
6 rrn loci, Clade_2
S. autolyticus CGMCC0516 (Mb)
A B c1 D* E F dnaA+
6 rrn loci, Group_O
S. avermitilis MA 4680 (Mb)
A B C E* D* F dnaA−
6 rrn loci, Clade_1
S. bacillaris ATCC 15855 (Mb)
A B C E* D* F dnaA−
6 rrn loci, Clade_2
S. bingchenggensis BCW 1 (Mb)
a2 F* c2 d3 E b1* dnaA+
6 rrn loci, Group_O
S. cattleya NRRL 8057 (Mb)
A B c1 [d/e1] [d/e5]* F dnaA−
6 rrn loci, Group_O
S. chartreusis ATCC 14922 (Mb)
A B C D E F dnaA+
6 rrn loci, Clade_1

Fig.S1 (3/15)

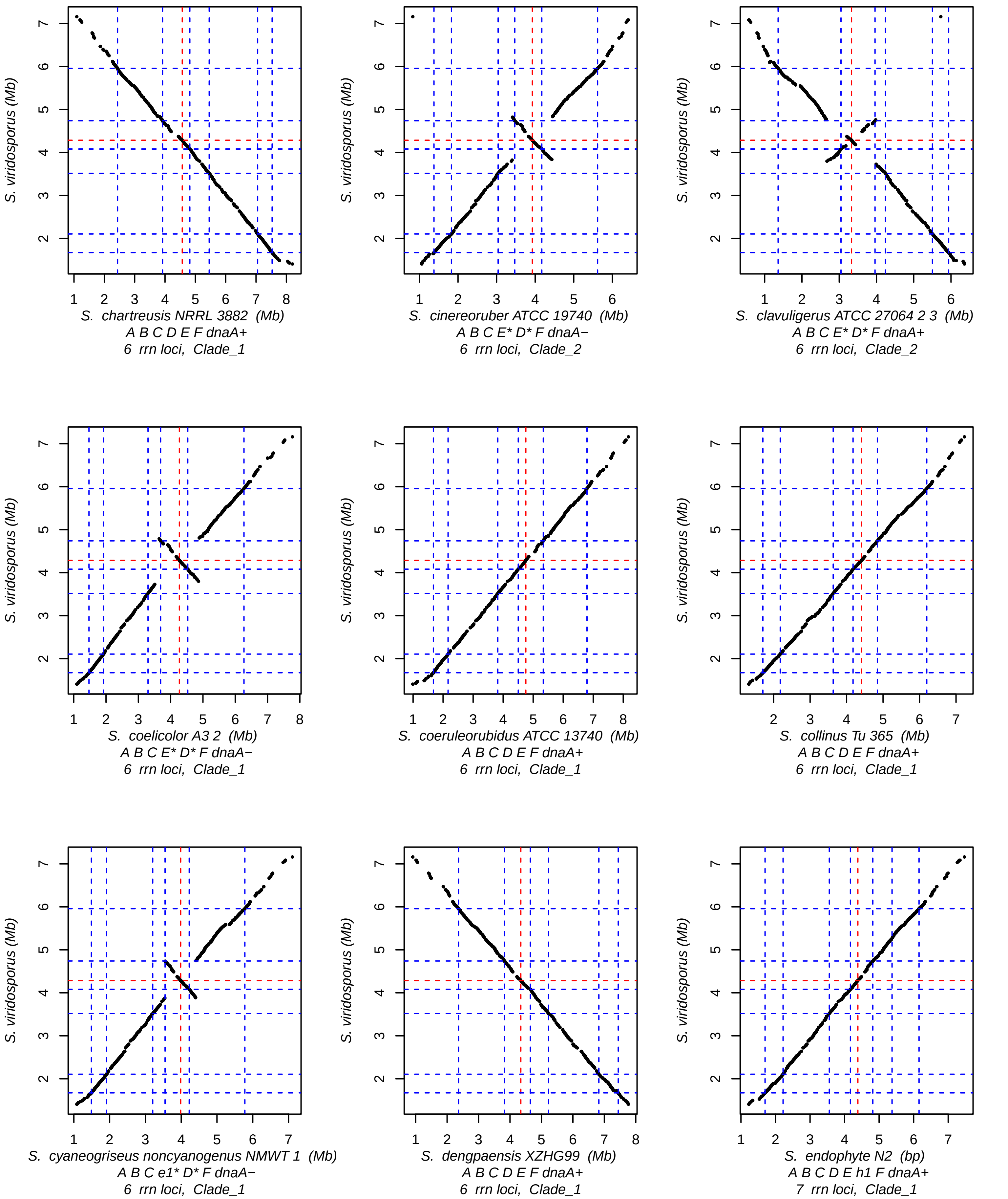
S. viridosporus (Mb)
S. chartreusis NRRL 3882 (Mb)
A B C D E F dnaA+
6 rrn loci, Clade_1
S. cinereoruber ATCC 19740 (Mb)
A B C E* D* F dnaA−
6 rrn loci, Clade_2
S. clavuligerus ATCC 27064 2 3 (Mb)
A B C E* D* F dnaA+
6 rrn loci, Clade_2
S. coelicolor A3 2 (Mb)
A B C E* D* F dnaA−
6 rrn loci, Clade_1
S. coeruleorubidus ATCC 13740 (Mb)
A B C D E F dnaA+
6 rrn loci, Clade_1
S. collinus Tu 365 (Mb)
A B C D E F dnaA+
6 rrn loci, Clade_1
S. cyaneogriseus noncyanogenus NMWT 1 (Mb)
A B C e1* D* F dnaA−
6 rrn loci, Clade_1
S. dengpaensis XZHG99 (Mb)
A B C D E F dnaA+
6 rrn loci, Clade_1
S. endophyte N2 (bp)
A B C D E h1 F dnaA+
7 rrn loci, Clade_1

Fig.S1 (4/15)

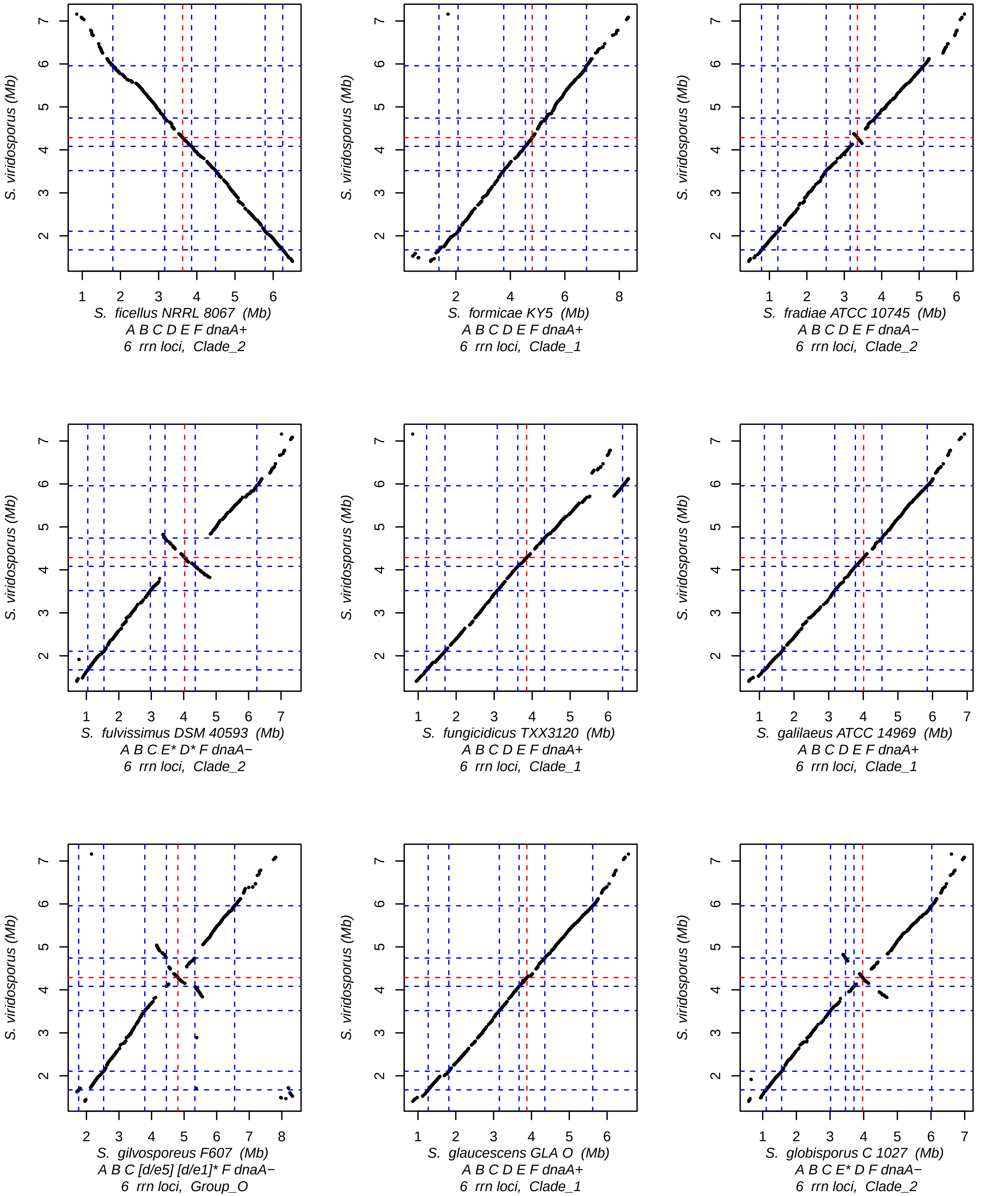
S. viridosporus (Mb)
S. ficellus NRRL 8067 (Mb)
A B C D E F dnaA+
6 rrn loci, Clade_2
S. formicae KY5 (Mb)
A B C D E F dnaA+
6 rrn loci, Clade_1
S. fradiae ATCC 10745 (Mb)
A B C D E F dnaA−
6 rrn loci, Clade_2
S. fulvissimus DSM 40593 (Mb)
A B C E* D* F dnaA−
6 rrn loci, Clade_2
S. fungicidicus TXX3120 (Mb)
A B C D E F dnaA+
6 rrn loci, Clade_1
S. galilaeus ATCC 14969 (Mb)
A B C D E F dnaA+
6 rrn loci, Clade_1
S. gilvosporeus F607 (Mb)
A B C [d/e5] [d/e1]* F dnaA−
6 rrn loci, Group_O
S. glaucescens GLA O (Mb)
A B C D E F dnaA+
6 rrn loci, Clade_1
S. globisporus C 1027 (Mb)
A B C E* D F dnaA−
6 rrn loci, Clade_2

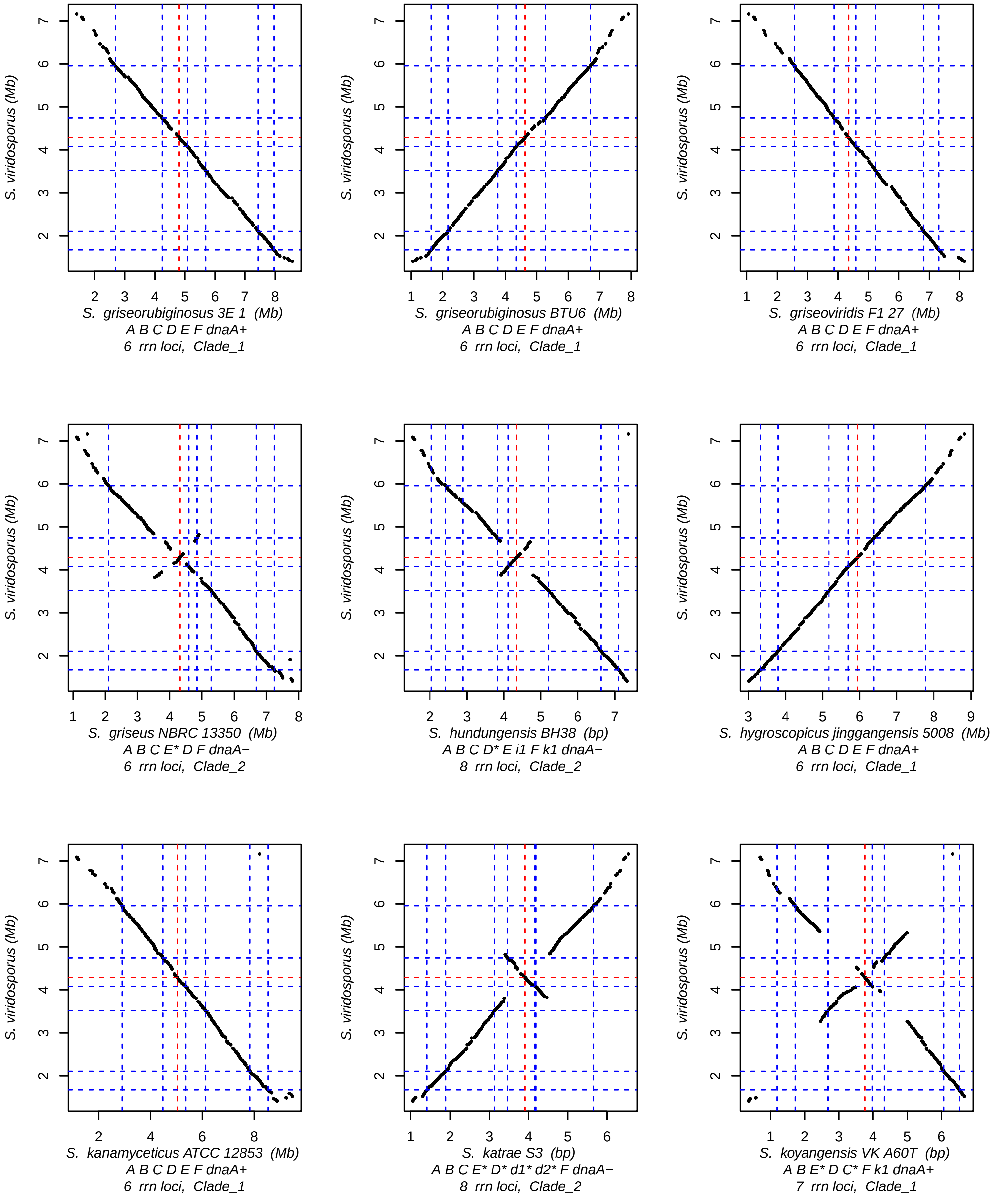
S. viridosporus (Mb)
S. griseorubiginosus 3E 1 (Mb)
A B C D E F dnaA+
6 rrn loci, Clade_1
S. griseorubiginosus BTU6 (Mb)
A B C D E F dnaA+
6 rrn loci, Clade_1
S. griseoviridis F1 27 (Mb)
A B C D E F dnaA+
6 rrn loci, Clade_1
S. griseus NBRC 13350 (Mb)
A B C E* D F dnaA−
6 rrn loci, Clade_2
S. hundungensis BH38 (bp)
A B C D* E i1 F k1 dnaA−
8 rrn loci, Clade_2
S. hygroscopicus jinggangensis 5008 (Mb)
A B C D E F dnaA+
6 rrn loci, Clade_1
S. kanamyceticus ATCC 12853 (Mb)
A B C D E F dnaA+
6 rrn loci, Clade_1
S. katrae S3 (bp)
A B C E* D* d1* d2* F dnaA−
8 rrn loci, Clade_2
S. koyangensis VK A60T (bp)
A B E* D C* F k1 dnaA+
7 rrn loci, Clade_1

Fig.S1 (6/15)

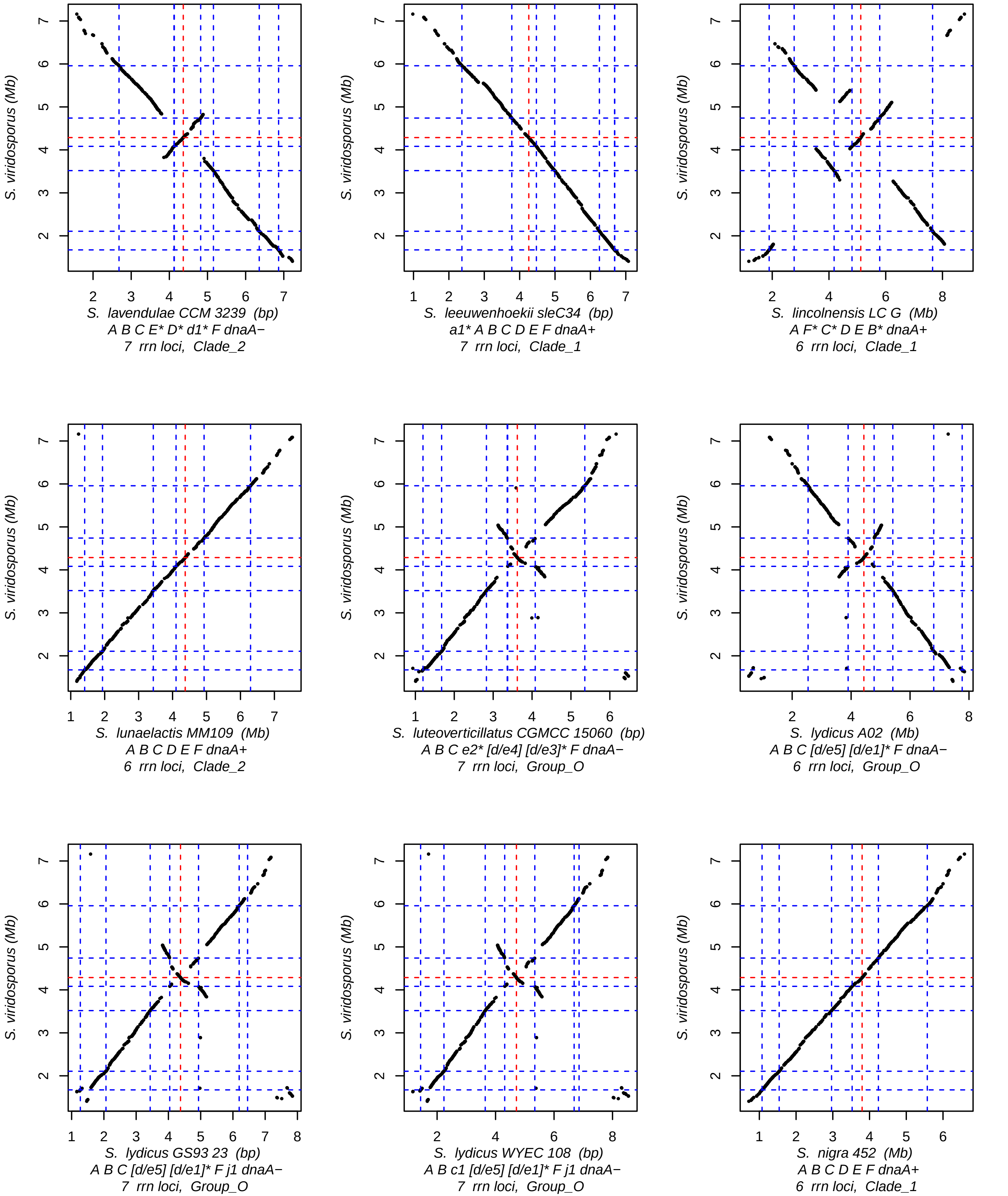
S. viridosporus (Mb)
S. lavendulae CCM 3239 (bp)
A B C E* D* d1* F dnaA−
7 rrn loci, Clade_2
S. leeuwenhoekii sleC34 (bp)
a1* A B C D E F dnaA+
7 rrn loci, Clade_1
S. lincolnensis LC G (Mb)
A F* C* D E B* dnaA+
6 rrn loci, Clade_1
S. lunaelactis MM109 (Mb)
A B C D E F dnaA+
6 rrn loci, Clade_2
S. luteoverticillatus CGMCC 15060 (bp)
A B C e2* [d/e4] [d/e3]* F dnaA−
7 rrn loci, Group_O
S. lydicus A02 (Mb)
A B C [d/e5] [d/e1]* F dnaA−
6 rrn loci, Group_O
S. lydicus GS93 23 (bp)
A B C [d/e5] [d/e1]* F j1 dnaA−
7 rrn loci, Group_O
S. lydicus WYEC 108 (bp)
A B c1 [d/e5] [d/e1]* F j1 dnaA−
7 rrn loci, Group_O
S. nigra 452 (Mb)
A B C D E F dnaA+
6 rrn loci, Clade_1

Fig.S1 (7/15)

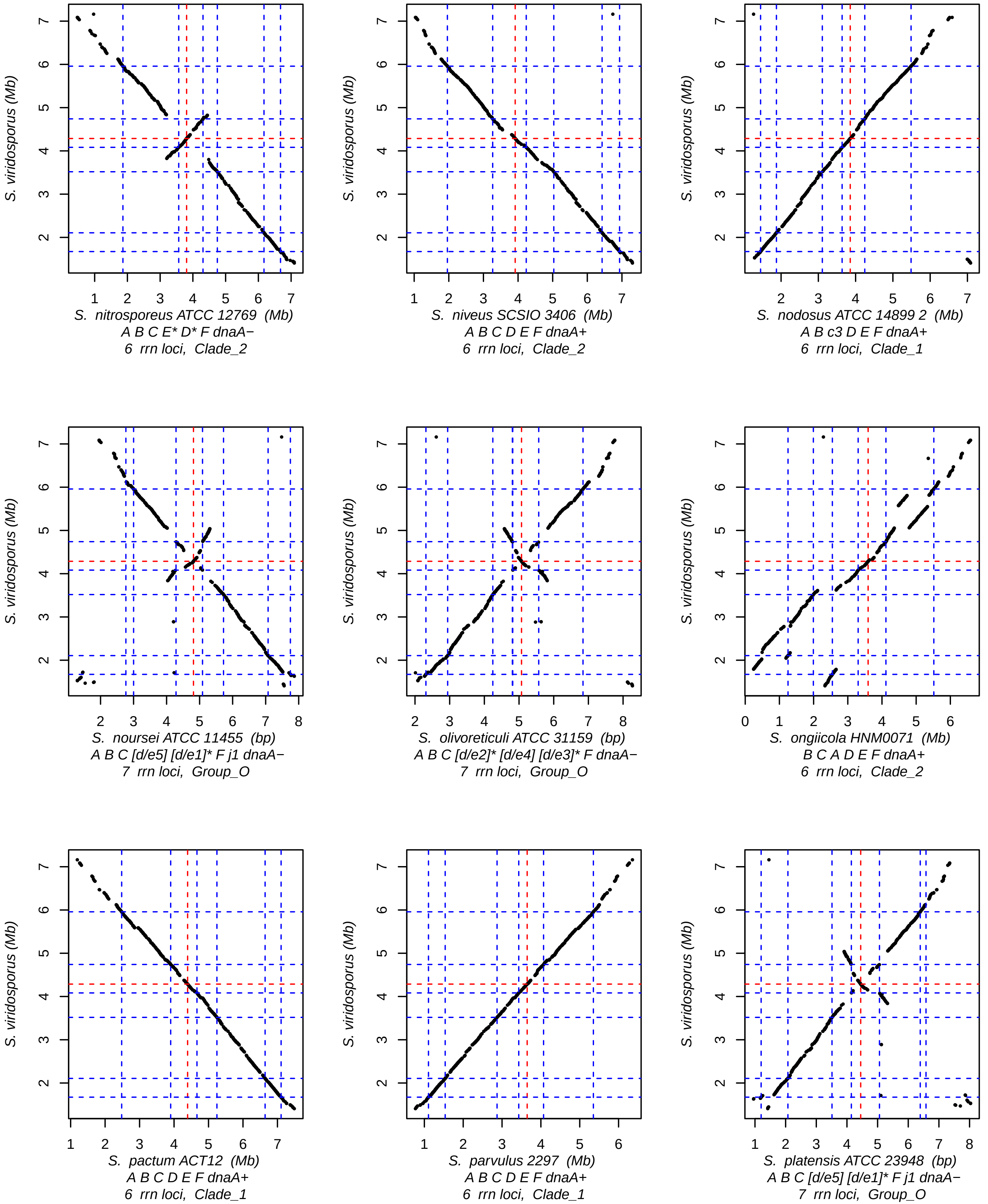
S. viridosporus (Mb)
S. nitrosporeus ATCC 12769 (Mb)
A B C E* D* F dnaA−
6 rrn loci, Clade_2
S. niveus SCSIO 3406 (Mb)
A B C D E F dnaA+
6 rrn loci, Clade_2
S. nodosus ATCC 14899 2 (Mb)
A B c3 D E F dnaA+
6 rrn loci, Clade_1
S. noursei ATCC 11455 (bp)
A B C [d/e5] [d/e1]* F j1 dnaA−
7 rrn loci, Group_O
S. olivoreticuli ATCC 31159 (bp)
A B C [d/e2]* [d/e4] [d/e3]* F dnaA−
7 rrn loci, Group_O
S. ongiicola HNM0071 (Mb)
B C A D E F dnaA+
6 rrn loci, Clade_2
S. pactum ACT12 (Mb)
A B C D E F dnaA+
6 rrn loci, Clade_1
S. parvulus 2297 (Mb)
A B C D E F dnaA+
6 rrn loci, Clade_1
S. platensis ATCC 23948 (bp)
A B C [d/e5] [d/e1]* F j1 dnaA−
7 rrn loci, Group_O

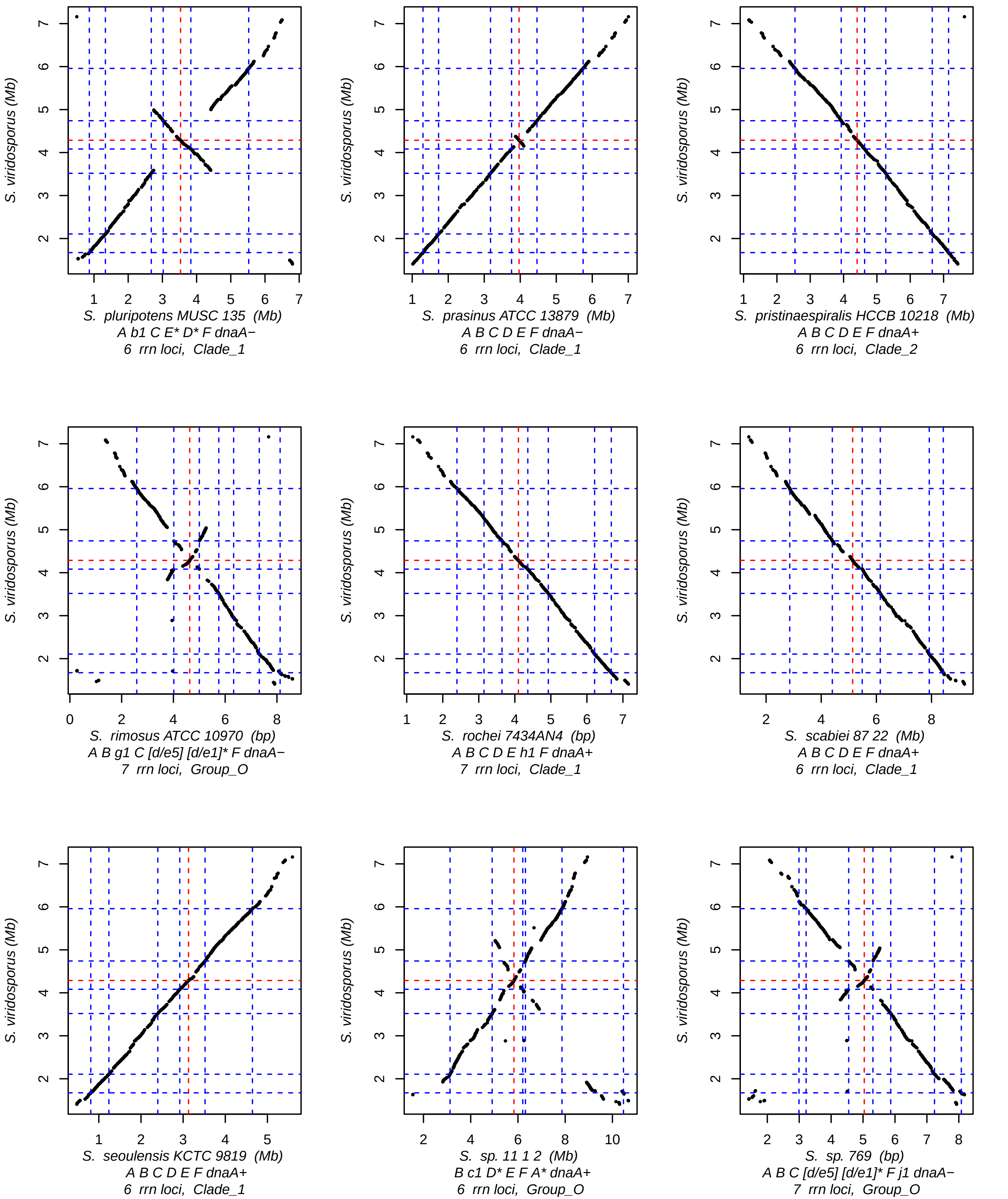
S. viridosporus (Mb)
S. pluripotens MUSC 135 (Mb)
A b1 C E* D* F dnaA−
6 rrn loci, Clade_1
S. prasinus ATCC 13879 (Mb)
A B C D E F dnaA−
6 rrn loci, Clade_1
S. pristinaespiralis HCCB 10218 (Mb)
A B C D E F dnaA+
6 rrn loci, Clade_2
S. rimosus ATCC 10970 (bp)
A B g1 C [d/e5] [d/e1]* F dnaA−
7 rrn loci, Group_O
S. rochei 7434AN4 (bp)
A B C D E h1 F dnaA+
7 rrn loci, Clade_1
S. scabiei 87 22 (Mb)
A B C D E F dnaA+
6 rrn loci, Clade_1
S. seoulensis KCTC 9819 (Mb)
A B C D E F dnaA+
6 rrn loci, Clade_1
S. sp. 11 1 2 (Mb)
B c1 D* E F A* dnaA+
6 rrn loci, Group_O
S. sp. 769 (bp)
A B C [d/e5] [d/e1]* F j1 dnaA−
7 rrn loci, Group_O

Fig.S1 (9/15)

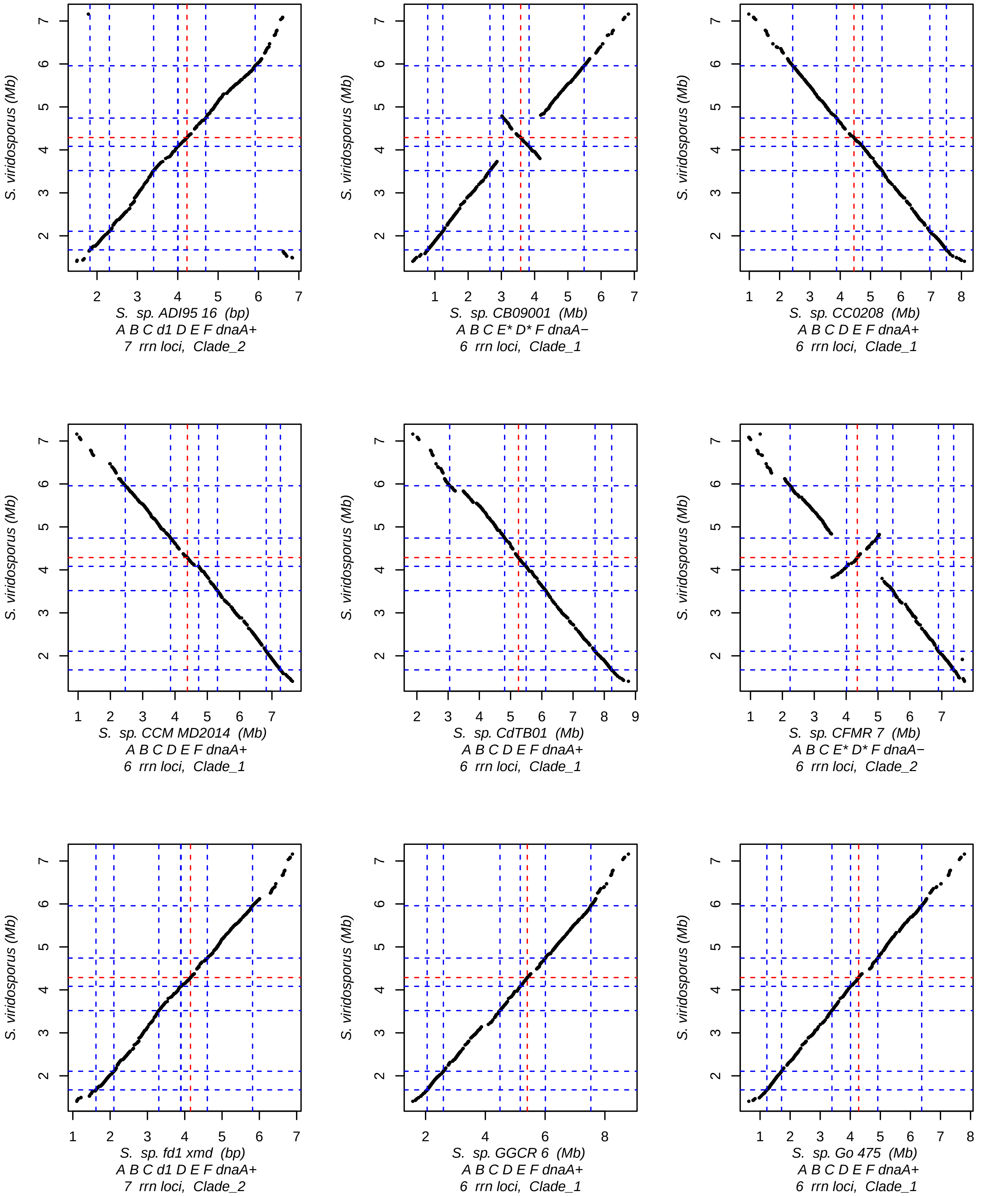
S. viridosporus (Mb)
S. sp. ADI95 16 (bp)
A B C d1 D E F dnaA+
7 rrn loci, Clade_2
S. viridosporus (Mb)
S. sp. CB09001 (Mb)
A B C E* D* F dnaA−
6 rrn loci, Clade_1
S. viridosporus (Mb)
S. sp. CC0208 (Mb)
A B C D E F dnaA+
6 rrn loci, Clade_1
S. viridosporus (Mb)
S. sp. CCM MD2014 (Mb)
A B C D E F dnaA+
6 rrn loci, Clade_1
S. viridosporus (Mb)
S. sp. CdTB01 (Mb)
A B C D E F dnaA+
6 rrn loci, Clade_1
S. viridosporus (Mb)
S. sp. CFMR 7 (Mb)
A B C E* D* F dnaA−
6 rrn loci, Clade_2
S. viridosporus (Mb)
S. sp. fd1 xmd (bp)
A B C d1 D E F dnaA+
7 rrn loci, Clade_2
S. viridosporus (Mb)
S. sp. GGCR 6 (Mb)
A B C D E F dnaA+
6 rrn loci, Clade_1
S. viridosporus (Mb)
S. sp. Go 475 (Mb)
A B C D E F dnaA+
6 rrn loci, Clade_1

Fig.S1 (10/15)

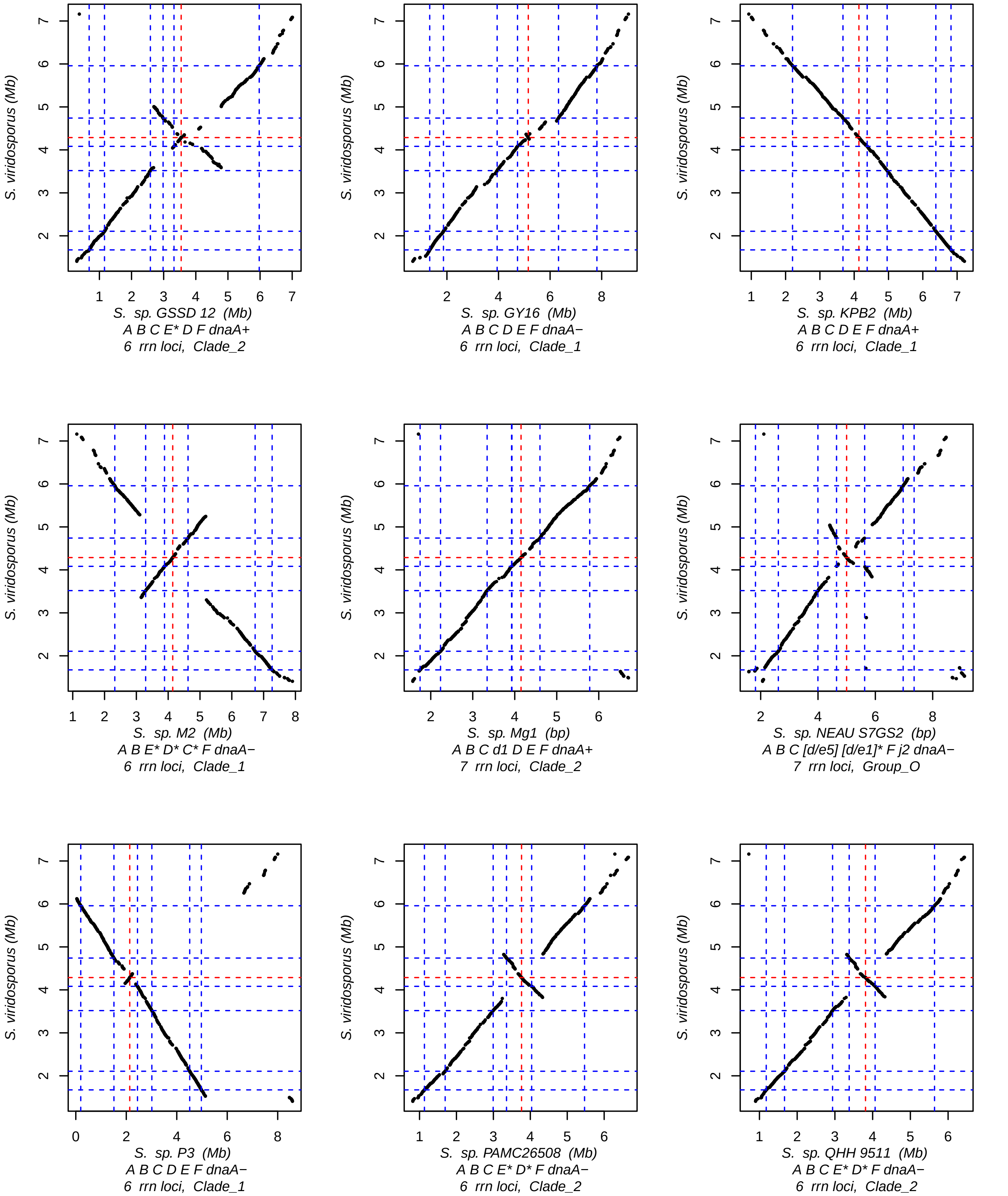
S. viridosporus (Mb)
S. sp. GSSD 12 (Mb)
A B C E* D F dnaA+
6 rrn loci, Clade_2
S. viridosporus (Mb)
S. sp. GY16 (Mb)
A B C D E F dnaA−
6 rrn loci, Clade_1
S. viridosporus (Mb)
S. sp. KPB2 (Mb)
A B C D E F dnaA+
6 rrn loci, Clade_1
S. viridosporus (Mb)
S. sp. M2 (Mb)
A B E* D* C* F dnaA−
6 rrn loci, Clade_1
S. viridosporus (Mb)
S. sp. Mg1 (bp)
A B C d1 D E F dnaA+
7 rrn loci, Clade_2
S. viridosporus (Mb)
S. sp. NEAU S7GS2 (bp)
A B C [d/e5] [d/e1]* F j2 dnaA−
7 rrn loci, Group_O
S. viridosporus (Mb)
S. sp. P3 (Mb)
A B C D E F dnaA−
6 rrn loci, Clade_1
S. viridosporus (Mb)
S. sp. PAMC26508 (Mb)
A B C E* D* F dnaA−
6 rrn loci, Clade_2
S. viridosporus (Mb)
S. sp. QHH 9511 (Mb)
A B C E* D* F dnaA−
6 rrn loci, Clade_2

Fig.S1 (11/15)

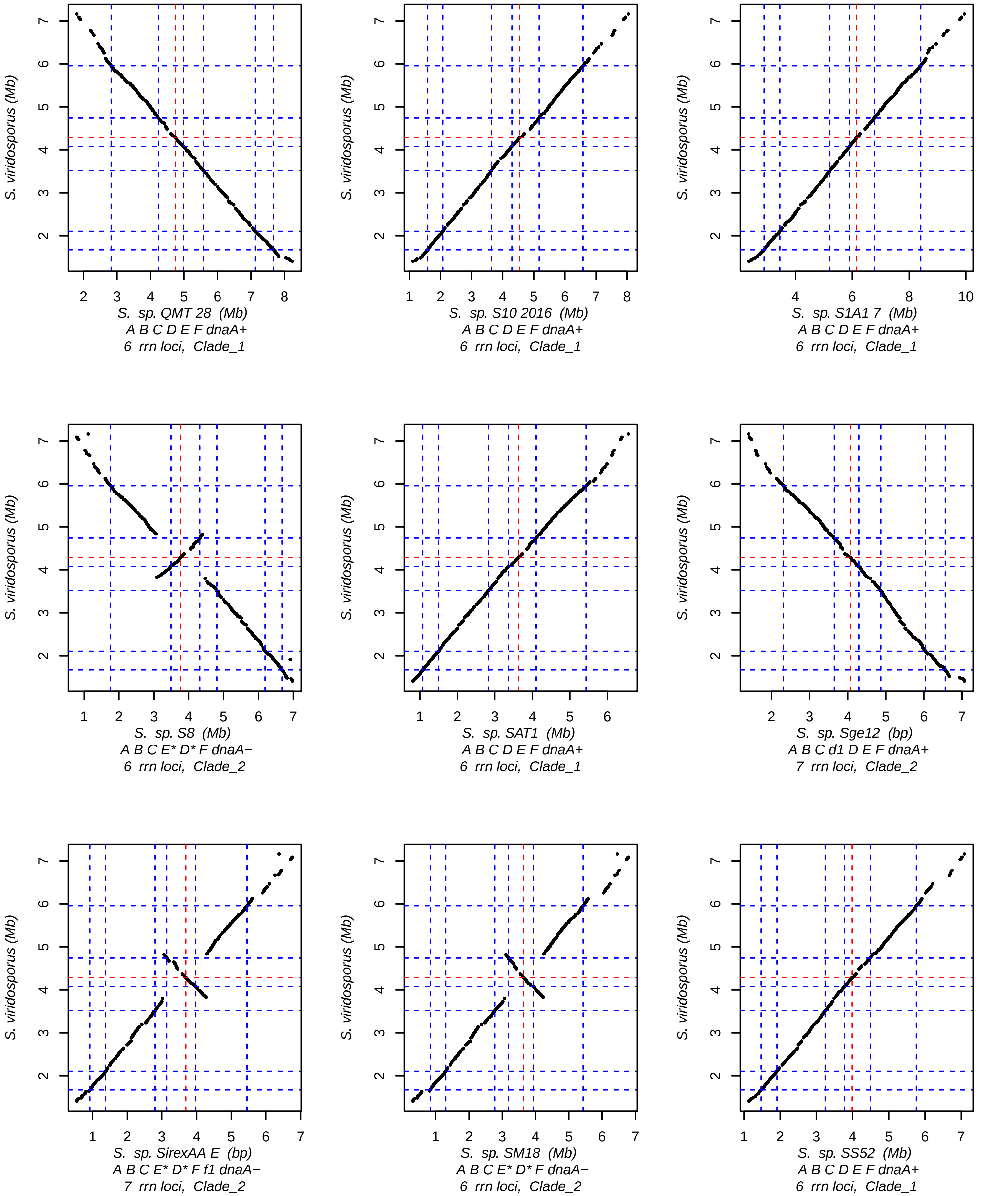
S. viridosporus (Mb)
S. sp. QMT 28 (Mb)
A B C D E F dnaA+
6 rrn loci, Clade_1
S. sp. S10 2016 (Mb)
A B C D E F dnaA+
6 rrn loci, Clade_1
S. sp. S1A1 7 (Mb)
A B C D E F dnaA+
6 rrn loci, Clade_1
S. sp. S8 (Mb)
A B C E* D* F dnaA−
6 rrn loci, Clade_2
S. sp. SAT1 (Mb)
A B C D E F dnaA+
6 rrn loci, Clade_1
S. sp. Sge12 (bp)
A B C d1 D E F dnaA+
7 rrn loci, Clade_2
S. sp. SirexAA E (bp)
A B C E* D* F f1 dnaA−
7 rrn loci, Clade_2
S. sp. SM18 (Mb)
A B C E* D* F dnaA−
6 rrn loci, Clade_2
S. sp. SS52 (Mb)
A B C D E F dnaA+
6 rrn loci, Clade_1

Fig.S1 (12/15)

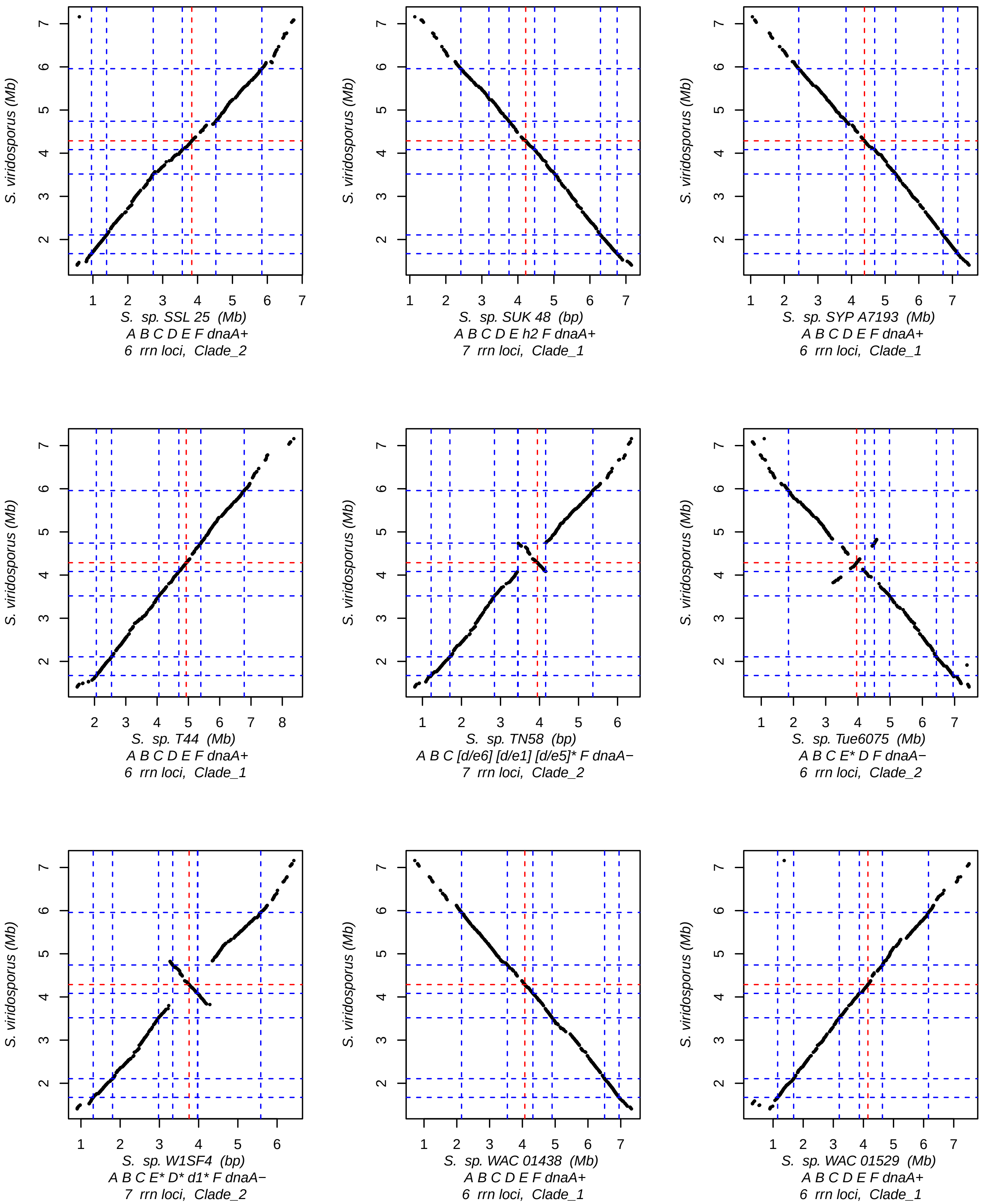
S. viridosporus (Mb)
S. sp. SSL 25 (Mb)
A B C D E F dnaA+
6 rrn loci, Clade_2
S. sp. SUK 48 (bp)
A B C D E h2 F dnaA+
7 rrn loci, Clade_1
S. sp. SYP A7193 (Mb)
A B C D E F dnaA+
6 rrn loci, Clade_1
S. sp. T44 (Mb)
A B C D E F dnaA+
6 rrn loci, Clade_1
S. sp. TN58 (bp)
A B C [d/e6] [d/e1] [d/e5]* F dnaA−
7 rrn loci, Clade_2
S. sp. Tue6075 (Mb)
A B C E* D F dnaA−
6 rrn loci, Clade_2
S. sp. W1SF4 (bp)
A B C E* D* d1* F dnaA−
7 rrn loci, Clade_2
S. sp. WAC 01438 (Mb)
A B C D E F dnaA+
6 rrn loci, Clade_1
S. sp. WAC 01529 (Mb)
A B C D E F dnaA+
6 rrn loci, Clade_1

Fig.S1 (13/15)

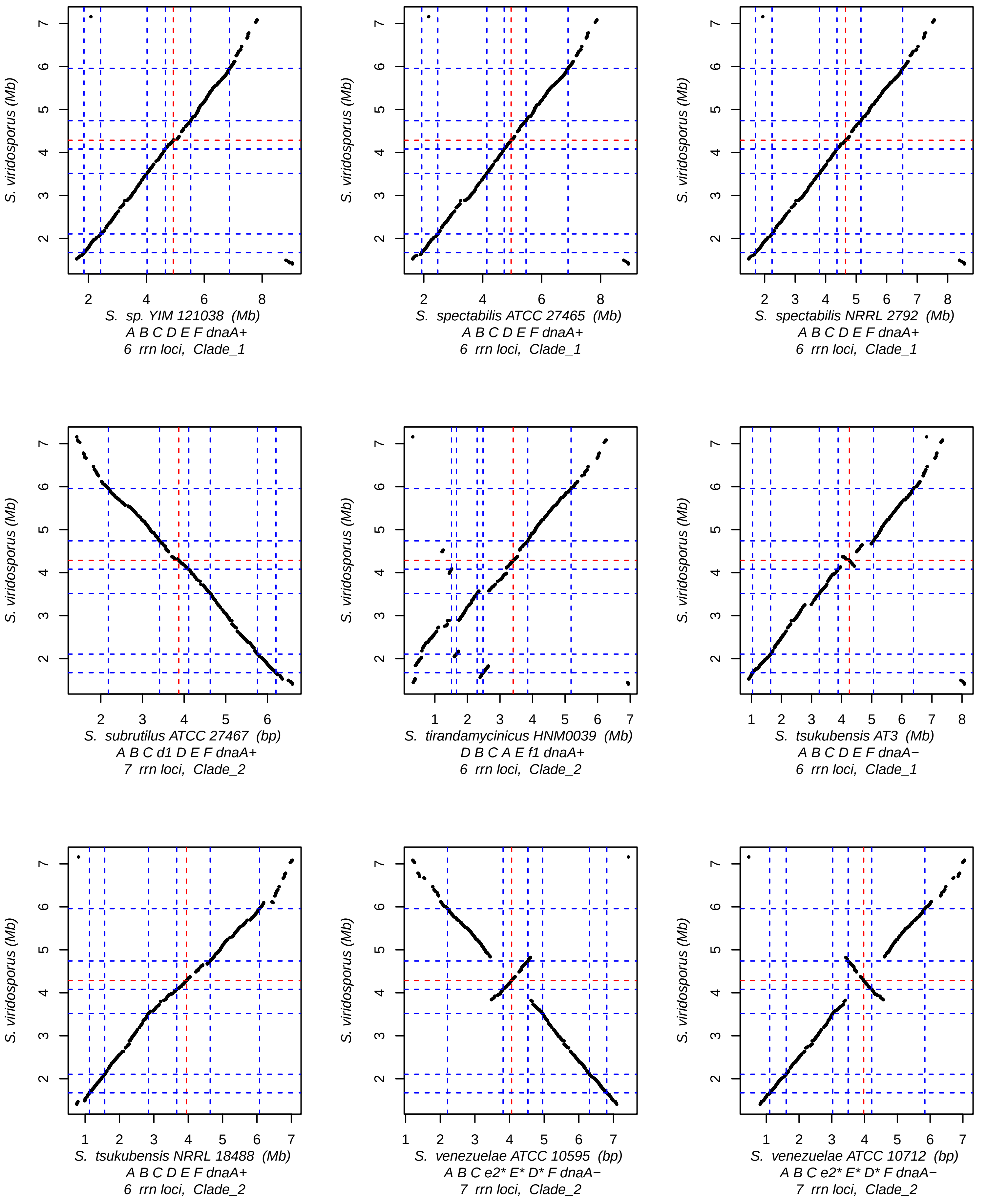
S. viridosporus (Mb)
S. sp. YIM 121038 (Mb)
A B C D E F dnaA+
6 rrn loci, Clade_1
S. spectabilis ATCC 27465 (Mb)
A B C D E F dnaA+
6 rrn loci, Clade_1
S. spectabilis NRRL 2792 (Mb)
A B C D E F dnaA+
6 rrn loci, Clade_1
S. subrutilus ATCC 27467 (bp)
A B C d1 D E F dnaA+
7 rrn loci, Clade_2
S. tirandamycinicus HNM0039 (Mb)
D B C A E f1 dnaA+
6 rrn loci, Clade_2
S. tsukubensis AT3 (Mb)
A B C D E F dnaA−
6 rrn loci, Clade_1
S. tsukubensis NRRL 18488 (Mb)
A B C D E F dnaA+
6 rrn loci, Clade_2
S. venezuelae ATCC 10595 (bp)
A B C e2* E* D* F dnaA−
7 rrn loci, Clade_2
S. venezuelae ATCC 10712 (bp)
A B C e2* E* D* F dnaA−
7 rrn loci, Clade_2

Fig.S1 (14/15)

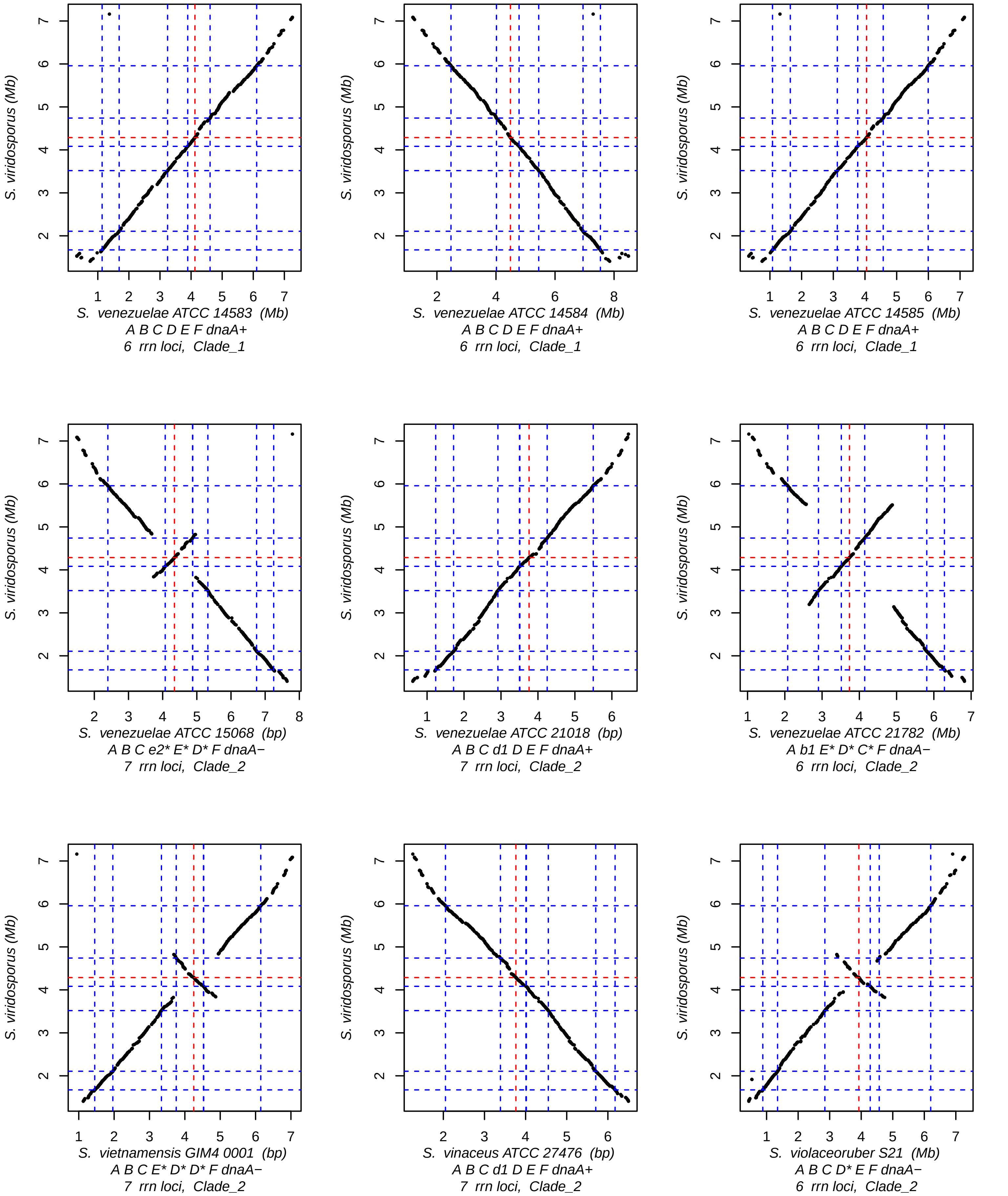
S. viridosporus (Mb)
S. venezuelae ATCC 14583 (Mb)
A B C D E F dnaA+
6 rrn loci, Clade_1
S. viridosporus (Mb)
S. venezuelae ATCC 14584 (Mb)
A B C D E F dnaA+
6 rrn loci, Clade_1
S. viridosporus (Mb)
S. venezuelae ATCC 14585 (Mb)
A B C D E F dnaA+
6 rrn loci, Clade_1
S. viridosporus (Mb)
S. venezuelae ATCC 15068 (bp)
A B C e2* E* D* F dnaA−
7 rrn loci, Clade_2
S. viridosporus (Mb)
S. venezuelae ATCC 21018 (bp)
A B C d1 D E F dnaA+
7 rrn loci, Clade_2
S. viridosporus (Mb)
S. venezuelae ATCC 21782 (Mb)
A b1 E* D* C* F dnaA−
6 rrn loci, Clade_2
S. viridosporus (Mb)
S. vietnamensis GIM4 0001 (bp)
A B C E* D* D* F dnaA−
7 rrn loci, Clade_2
S. viridosporus (Mb)
S. vinaceus ATCC 27476 (bp)
A B C d1 D E F dnaA+
7 rrn loci, Clade_2
S. viridosporus (Mb)
S. violaceoruber S21 (Mb)
A B C D* E F dnaA−
6 rrn loci, Clade_2

**Figure S1: Pairwise comparisons of all the core genomes of the panel with the consensus reference strain, *Streptomyces viridosporus* T7A ATCC 39115**

The *rrn* nomenclature, number of *rrn*, and clade (or group) are indicated for each strain below each graph. The positions of *rrn* operons and of the origin of replication are indicated by blue and red dashed lines, respectively. The presence of an unbroken diagonal indicates that the strains have identically ordered core genomes. When the diagonal starts at the bottom left and ends at the top right, it means that the genome sequence available in the databases is oriented in the same orientation as the consensus used in this study (*Streptomyces viridosporus* T7A ATCC 39115), and that the *rrn* configuration shown below each graph can be directly transposed onto the graph. If not, the sequence is in the opposite direction. In this case, the consensus should be reversed when transposed on the graph (e.g. '*rrn* ABCDEF *dnaA*+' becomes '*rrn* F*E*D*C*B*A* *dnaA*-').
