## Supplementary material for "Ribosomal RNA operons define a central functional compartment in the *Streptomyces* chromosome": Figure S2.pdf

**A**

| *rrn* environment category | *rrn* environment id | Core gene -1[§] | Core gene closest to the *rrn* operon start position[§] | Core gene +1[§] | Number of genomes |
|---|---|---|---|---|---|
| *Aa* | **A** | **Orth_2229 / SCO1389 / SAM23877_1447** | **Orth_2335 / SCO1390 / SAM23877_1448** | **Orth_2240 / SCO1391 / SAM23877_1449** | **125** |
| | a1 | Orth_3457 / SCO1388 / SAM23877_1446 | Orth_2229 | Orth_2335 | 1 |
| | a2 | Orth_3494 / SCO1152 / SAM23877_1235 | Orth_2335 | Orth_2240 | 1 |
| *Bb* | **B** | **Orth_420 / SCO1789 / SAM23877_1854** | **Orth_9302 / SCO1796 / SAM23877_1863** | **Orth_5066 / SCO1797 / SAM23877_1864** | **124** |
| | b1 | Orth_8822 / SCO1788 / SAM23877_1853 | Orth_420 | Orth_9302 | 3 |
| *Cc* | **C** | **Orth_3539 / SCO3013 / SAM23877_3051** | **Orth_9623 / SCO3024 / SAM23877_3064** | **Orth_6428 / SCO3025 / SAM23877_3065** | **121** |
| | c1 | Orth_8414 / SCO3011 / SAM23877_3049 | Orth_3539 | Orth_9623 | 4 |
| | c2 | Orth_4341 / SCO3172 / SAM23877_3221 | Orth_9623 | Orth_6428 | 1 |
| | c3 | Orth_9233 / SCO2966 / SAM23877_3001 | Orth_9623 | Orth_6428 | 1 |
| *Dd* | **D** | **Orth_4139 / SCO4127 / SAM23877_3557** | **Orth_6483 / SCO4122 / SAM23877_3565** | **Orth_2534 / SCO4121 / SAM23877_3566** | **112** |
| | d1 | Orth_7996 / SCO4128 / SAM23877_3556 | Orth_4139 | Orth_6483 | 10 |
| | d2 | Orth_6541 / SCO4137 / SAM23877_3546 | Orth_7996 | Orth_4139 | 1 |
| | d3 | Orth_7996 | Orth_4139 | Orth_1153 / SCO3615 / SAM23877_4004 | 1 |
| *Ee* | **E** | **Orth_8528 / SCO3345 / SAM23877_4216** | **Orth_2277 / SCO3337 / SAM23877_4223** | **Orth_6749 / SCO3330 / SAM23877_4229** | **111** |
| | e1 | Orth_8528 | Orth_2277 | Orth_8774 / SCO4366 / SAM23877_3359 | 1 |
| | e2 | Orth_2277 | Orth_6749 | Orth_4487 / SCO3323 / SAM23877_4236 | 4 |
| *[d/e] recombinants* | [d/e1] | Orth_4139 | Orth_2277 | Orth_8528 | 12 |
| | [d/e2] | Orth_6483 | Orth_2277 | Orth_6749 | 2 |
| | [d/e3] | Orth_4139 | Orth_8528 | Orth_4097 / SCO3347 / SAM23877_4214 | 3 |
| | [d/e4] | Orth_2277 | Orth_6483 | Orth_2534 | 3 |
| | [d/e5] | Orth_6749 | Orth_6483 | Orth_2534 | 12 |
| | [d/e6] | Orth_7996 | Orth_4139 | Orth_2277 | 1 |
| *Ff* | **F** | **Orth_5289 / SCO5740 / SAM23877_5442** | **Orth_6126 / SCO5744 / SAM23877_5446** | **Orth_7379 / SCO5747 / SAM23877_5451** | **126** |
| | f1 | Orth_6126 | Orth_7379 | Orth_6249 / SCO5749 / SAM23877_5453 | 2 |
| *Atypical/ ectopic rrn environments* | g1 | Orth_2159 / SCO2538 / SAM23877_2587 | Orth_2308 / SCO2533 / SAM23877_2584 | Orth_3295 / SCO2547 / SAM23877_2592 | 1 |
| | h1 | Orth_1360 / SCO4979 / SAM23877_4794 | Orth_5738 / SCO5005 / SAM23877_4811 | Orth_7524 / SCO5009 / SAM23877_4815 | 2 |
| | h2 | Orth_7601 / SCO4975 / SAM23877_4786 | Orth_1360 | Orth_5738 | 1 |
| | i1 | Orth_5509 / SCO5291 / SAM23877_5067 | Orth_9438 / SCO5353 / SAM23877_5097 | Orth_7899 / SCO5354 / SAM23877_5098 | 1 |
| | j1 | Orth_75 / SCO5900 / SAM23877_5578 | Orth_9758 / SCO5901 / SAM23877_5579 | Orth_2449 / SCO5998 / SAM23877_5714 | 6 |
| | j2 | Orth_9758 | Orth_2449 | Orth_506 / SCO5999 / SAM23877_5716 | 1 |
| | k1 | Orth_4148 / SCO6060 / SAM23877_5773 | Orth_719 / SCO6061 / SAM23877_5774 | Orth_1203 / SCO6083 / SAM23877_5801 | 4 |

**B**

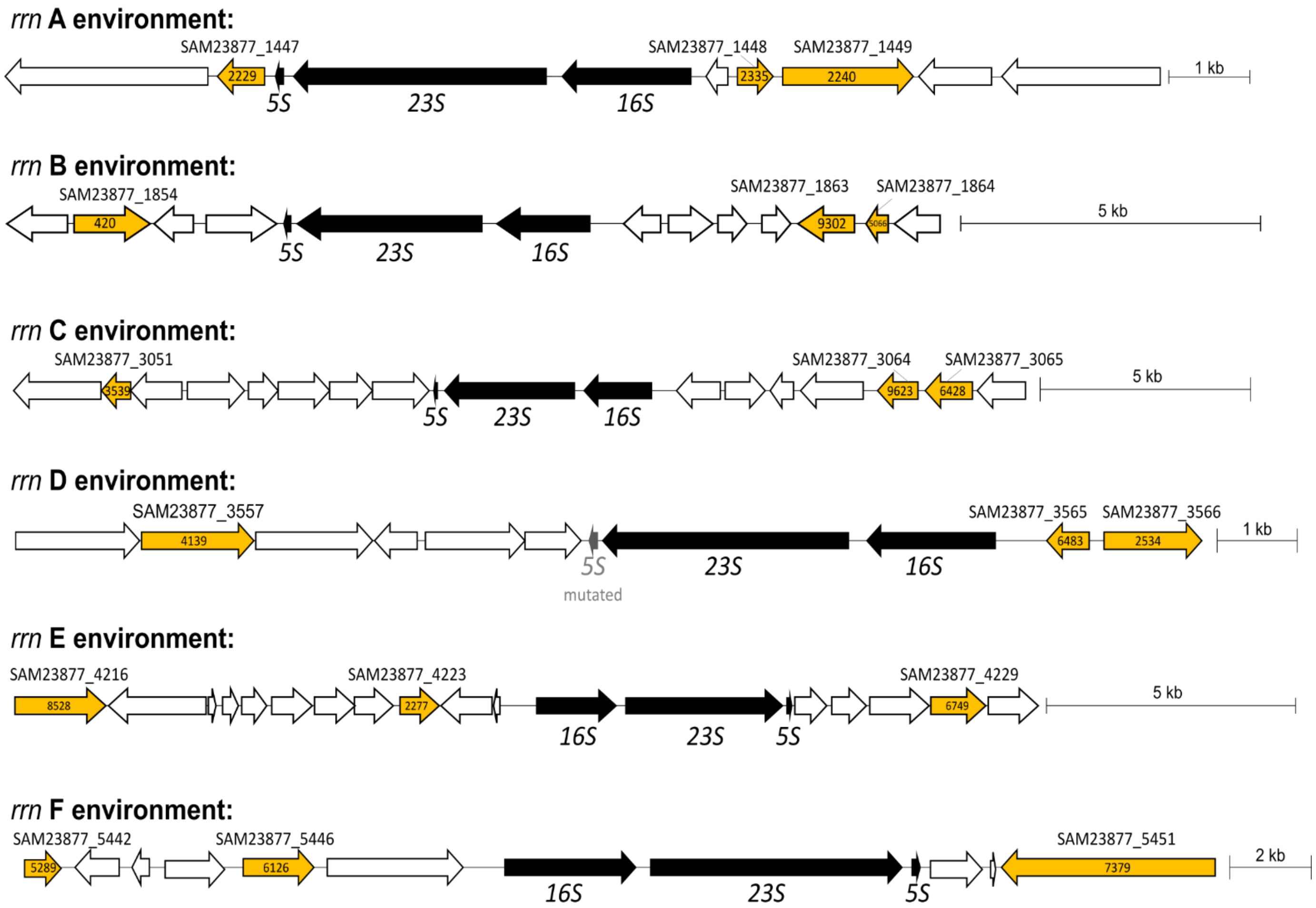


**Figure S2: Proposed *rrn* nomenclature based on core gene environments**

A. **Core gene environments characterized around all the *rrn* of the panel.** Canonical *rrn* neighborhoods are in bold. When first cited, the core gene identifiers are indicated as follows: ortholog group number ('Orth_#') based on the re-annotation performed in this study / *S. coelicolor* ('SCO#') Uniprot identifier / *S. ambofaciens* ATCC 23877 ('SAM23877_#') Uniprot identifier. If already cited, the CDS is then identified solely on the basis of its ortholog group number. These two strains were chosen for reference because *S. coelicolor* A3(2) corresponds to the most documented *Streptomyces* strain, and *S. ambofaciens* ATCC 23877 is another model *Streptomyces*, closest to the ancestral consensus in terms of core gene order. 'Core gene -1' and 'Core gene +1' designate the core genes located upstream and downstream of the core gene closest to the beginning of the *rrn* operon, respectively. The *rrn* environments are oriented according to the consensus order as found in *S. viridosporus* T7A ATCC 39115. An asterisk is used in the nomenclature to indicate that the motif is in the opposite orientation to that shown above.

B. **Schematic representation of the canonical *rrn* environments in *S. ambofaciens* ATCC 23877.** Core genes that are a signature of the *rrn* environment are in orange. The numbers in each arrow correspond to the annotation of the orthologous groups performed in this study (annotation of all genomes in the same manner is available in **Supplementary Table 6**).
