## Supplementary material for "Ribosomal RNA operons define a central functional compartment in the *Streptomyces* chromosome": Figure S3.pdf

**Figure S3 (1/3)**

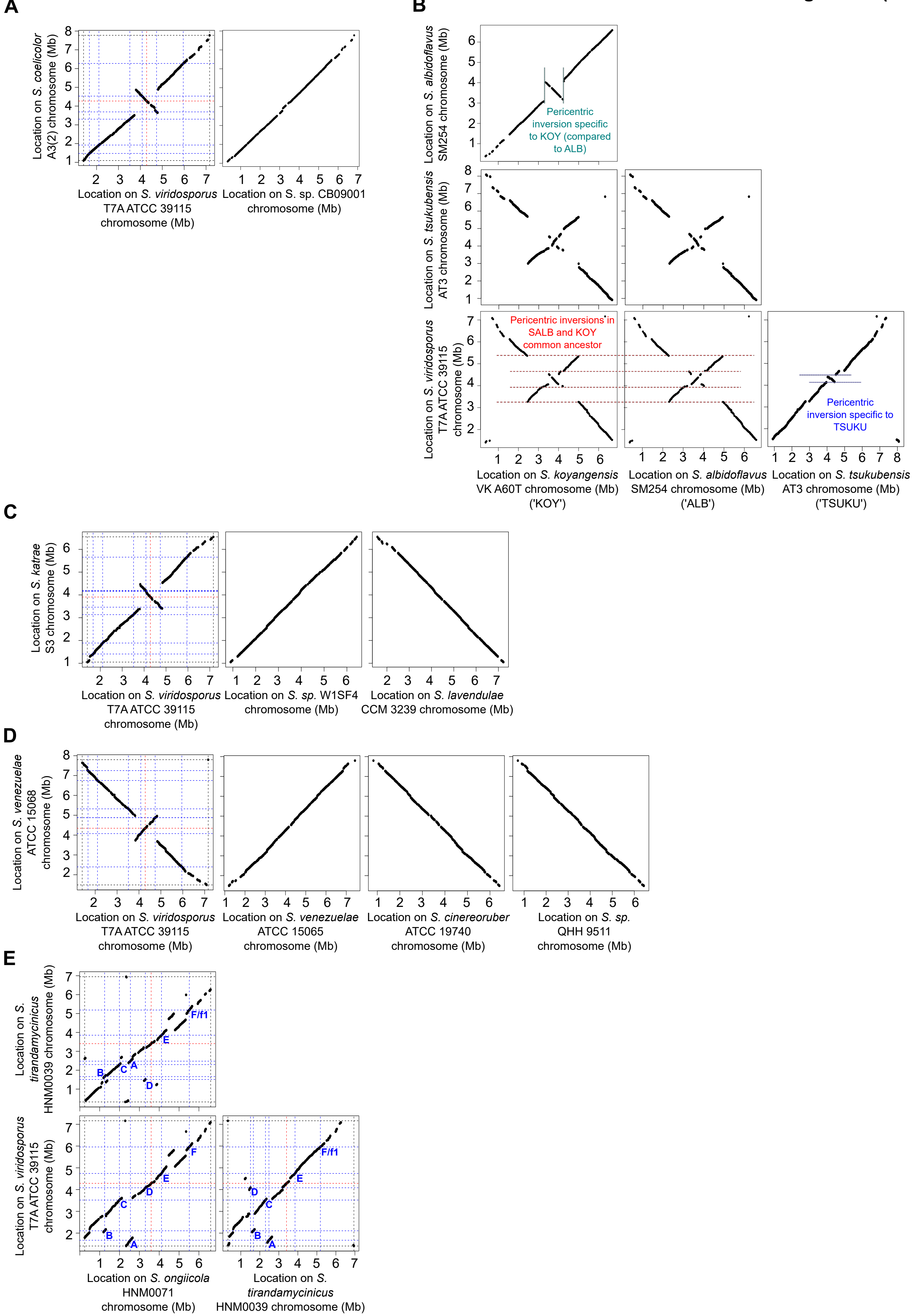

A
Location on S. coelicolor A3(2) chromosome (Mb)
Location on S. viridosporus T7A ATCC 39115 chromosome (Mb)
Location on S. sp. CB09001 chromosome (Mb)
B
Location on S. albidoflavus SM254 chromosome (Mb)
Pericentric inversion specific to KOY (compared to ALB)
Location on S. tsukubensis AT3 chromosome (Mb)
Location on S. viridosporus T7A ATCC 39115 chromosome (Mb)
Pericentric inversions in SALB and KOY common ancestor
Pericentric inversion specific to TSUKU
Location on S. koyangensis VK A60T chromosome (Mb) ('KOY')
Location on S. albidoflavus SM254 chromosome (Mb) ('ALB')
Location on S. tsukubensis AT3 chromosome (Mb) ('TSUKU')
C
Location on S. katrae S3 chromosome (Mb)
Location on S. viridosporus T7A ATCC 39115 chromosome (Mb)
Location on S. sp. W1SF4 chromosome (Mb)
Location on S. lavendulae CCM 3239 chromosome (Mb)
D
Location on S. venezuelae ATCC 15068 chromosome (Mb)
Location on S. viridosporus T7A ATCC 39115 chromosome (Mb)
Location on S. venezuelae ATCC 15065 chromosome (Mb)
Location on S. cinereoruber ATCC 19740 chromosome (Mb)
Location on S. sp. QHH 9511 chromosome (Mb)
E
Location on S. tirandamycinicus HNM0039 chromosome (Mb)
Location on S. viridosporus T7A ATCC 39115 chromosome (Mb)
Location on S. ongiicola HNM0071 chromosome (Mb)
Location on S. tirandamycinicus HNM0039 chromosome (Mb)
A
B
C
D
E
F/f1
F

**Figure S3 (2/3)**

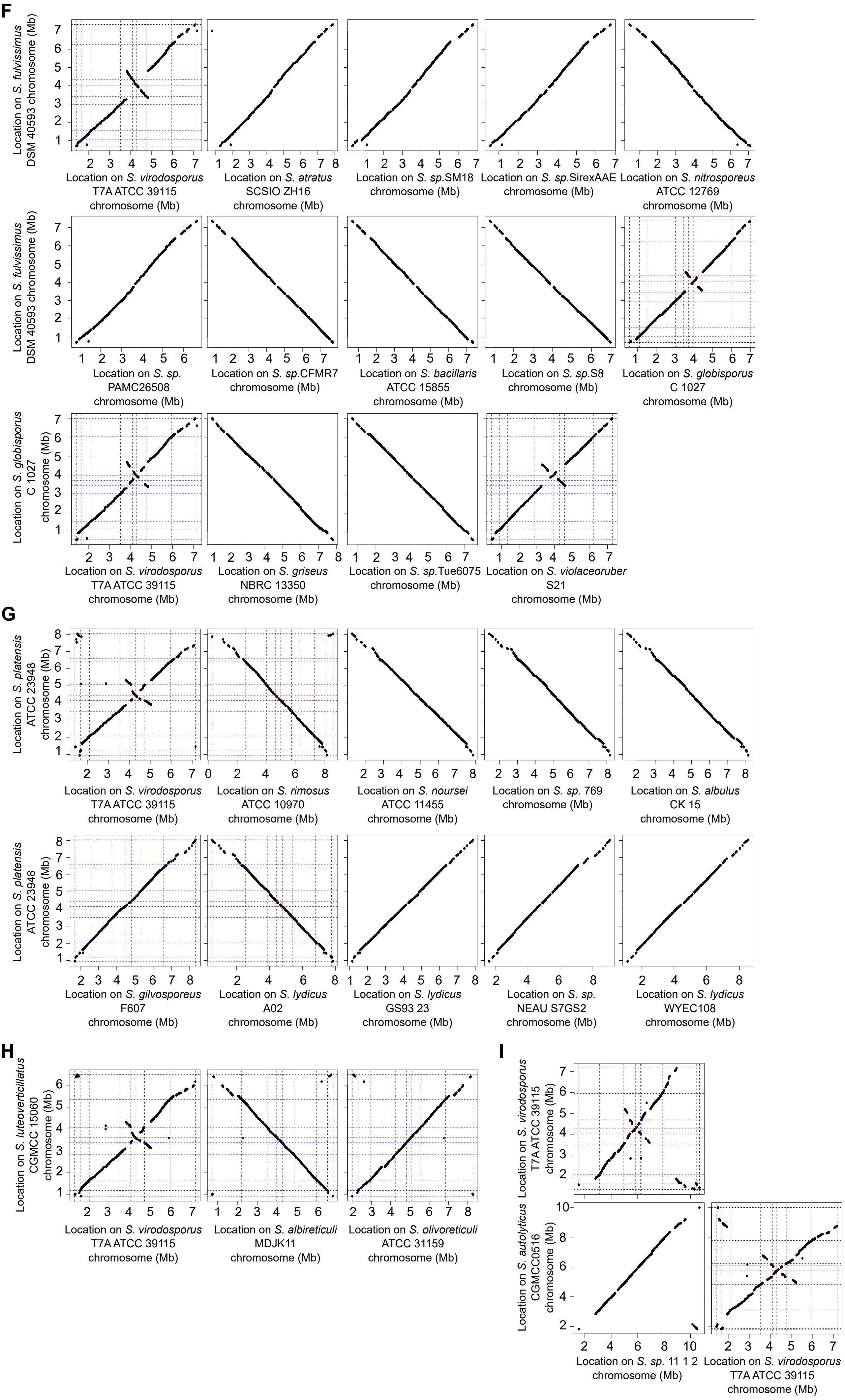
F
Location on S. fulvissimus DSM 40593 chromosome (Mb)
Location on S. virodosporus T7A ATCC 39115 chromosome (Mb)
Location on S. atratus SCSIO ZH16 chromosome (Mb)
Location on S. sp.SM18 chromosome (Mb)
Location on S. sp.SirexAAE chromosome (Mb)
Location on S. nitrosporeus ATCC 12769 chromosome (Mb)
Location on S. fulvissimus DSM 40593 chromosome (Mb)
Location on S. sp. PAMC26508 chromosome (Mb)
Location on S. sp.CFMR7 chromosome (Mb)
Location on S. bacillaris ATCC 15855 chromosome (Mb)
Location on S. sp.S8 chromosome (Mb)
Location on S. globisporus C 1027 chromosome (Mb)
Location on S. globisporus C 1027 chromosome (Mb)
Location on S. virodosporus T7A ATCC 39115 chromosome (Mb)
Location on S. griseus NBRC 13350 chromosome (Mb)
Location on S. sp.Tue6075 chromosome (Mb)
Location on S. violaceoruber S21 chromosome (Mb)
G
Location on S. platensis ATCC 23948 chromosome (Mb)
Location on S. virodosporus T7A ATCC 39115 chromosome (Mb)
Location on S. rimosus ATCC 10970 chromosome (Mb)
Location on S. noursei ATCC 11455 chromosome (Mb)
Location on S. sp. 769 chromosome (Mb)
Location on S. albulus CK 15 chromosome (Mb)
Location on S. platensis ATCC 23948 chromosome (Mb)
Location on S. gilvosporeus F607 chromosome (Mb)
Location on S. lydicus A02 chromosome (Mb)
Location on S. lydicus GS93 23 chromosome (Mb)
Location on S. sp. NEAU S7GS2 chromosome (Mb)
Location on S. lydicus WYEC108 chromosome (Mb)
H
Location on S. luteoverticillatus CGMCC 15060 chromosome (Mb)
Location on S. virodosporus T7A ATCC 39115 chromosome (Mb)
Location on S. albireticuli MDJK11 chromosome (Mb)
Location on S. olivoreticuli ATCC 31159 chromosome (Mb)
I
Location on S. virodosporus T7A ATCC 39115 chromosome (Mb)
Location on S. autolyticus CGMCC0516 chromosome (Mb)
Location on S. sp. 11 1 2 chromosome (Mb)
Location on S. virodosporus T7A ATCC 39115 chromosome (Mb)

### Figure S3: Pairwise comparison of the core genomes that are proposed to share rearrangements that occurred in their common ancestor

Panels A to I present the pairwise comparisons that support the occurrence of intra-chromosomal rearrangement in the ancestor of strains of interest. In each case, a comparison to the consensus strain (*S. viridosporus* T7A ATCC 39115) is included as a reference, as well as the indication of the position of *rrn* operons (blue), the origin of replication (red) and the core genome ends (black) for at least one pairwise comparison. In the case of very complex rearrangements (panels E), the name of the *rrn* according to the nomenclature proposed in this study has been indicated. The presence of an unbroken diagonal indicates that the strains have identically ordered core genomes.
