## Supplementary material for "Ribosomal RNA operons define a central functional compartment in the *Streptomyces* chromosome": Figure S4.pdf

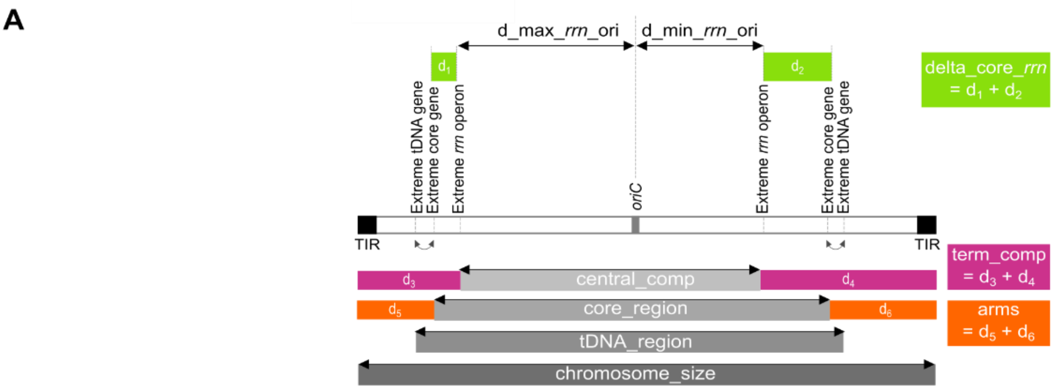


**B**

$$y_i = \beta_0 + \beta_1.x_{i1} + \beta_2.x_{i2} + \beta_3.x_{i3} + \beta_4.x_{i4} + \varepsilon_i$$

where $y_i$ is the core region size (in bp) of the $i^{th}$ genome

| | Estimate | Standard error | *t* value | Pr(>\|t\|) | |
|---|---|---|---|---|---|
| Intercept | $\beta_0$ = 1.016e+06 | 4.974e+05 | 2.042 | 0.043332 | * |
| Maximal distance between *rrn* and *oriC* $(x_{i1})$ | $\beta_1$ = 7.262e-01 | 2.603e-01 | 2.790 | 0.006136 | ** |
| Chromosome size $(x_{i2})$ | $\beta_2$ = 2.398e-01 | 3.970e-02 | 6.041 | 1.77e-08 | *** |
| Central compartment size $(x_{i3})$ | $\beta_3$ = 5.522e-01 | 1.468e-01 | 3.762 | 0.000262 | *** |
| Number of *rrn* genes $(x_{i4})$ | $\beta_3$ = -6.694e+04 | 2.036e+04 | -3.288 | 0.001323 | ** |

| | Min | First quartile | Median | Third quartile | Max |
|---|---|---|---|---|---|
| Residuals $(\varepsilon_i)$ | -892177 | -174499 | -22613 | 164547 | 1663768 |

Residual standard error: 312800 bp on 120 degrees of freedom
Multiple R²: 0.8639, Adjusted R²: 0.8594
F-statistic: 190.4 on 4 and 120 degrees of freedom, *p*-value: < 2.2e-16
AIC: 3524.939, 6 degrees of freedom

**C**

| | Degrees of freedom | Sum of Squares | Mean square | F value | Pr(>F) | |
|---|---|---|---|---|---|---|
| Maximal distance between *rrn* and *oriC* $(x_{i1})$ | 1 | 6.9333e+13 | 6.9333e+13 | 708.736 | < 2.2e-16 | *** |
| Chromosome size $(x_{i2})$ | 1 | 2.9697e+12 | 2.9697e+12 | 30.357 | 2.086e-07 | *** |
| Central compartment size $(x_{i3})$ | 1 | 1.1600e+12 | 1.1600e+12 | 11.858 | 0.0007914 | *** |
| Number of *rrn* genes $(x_{i4})$ | 1 | 1.0578e+12 | 1.0578e+12 | 10.813 | 0.0013229 | ** |
| Residuals $(\varepsilon_i)$ | 120 | 1.1739e+13 | 9.7826e+10 | | | |

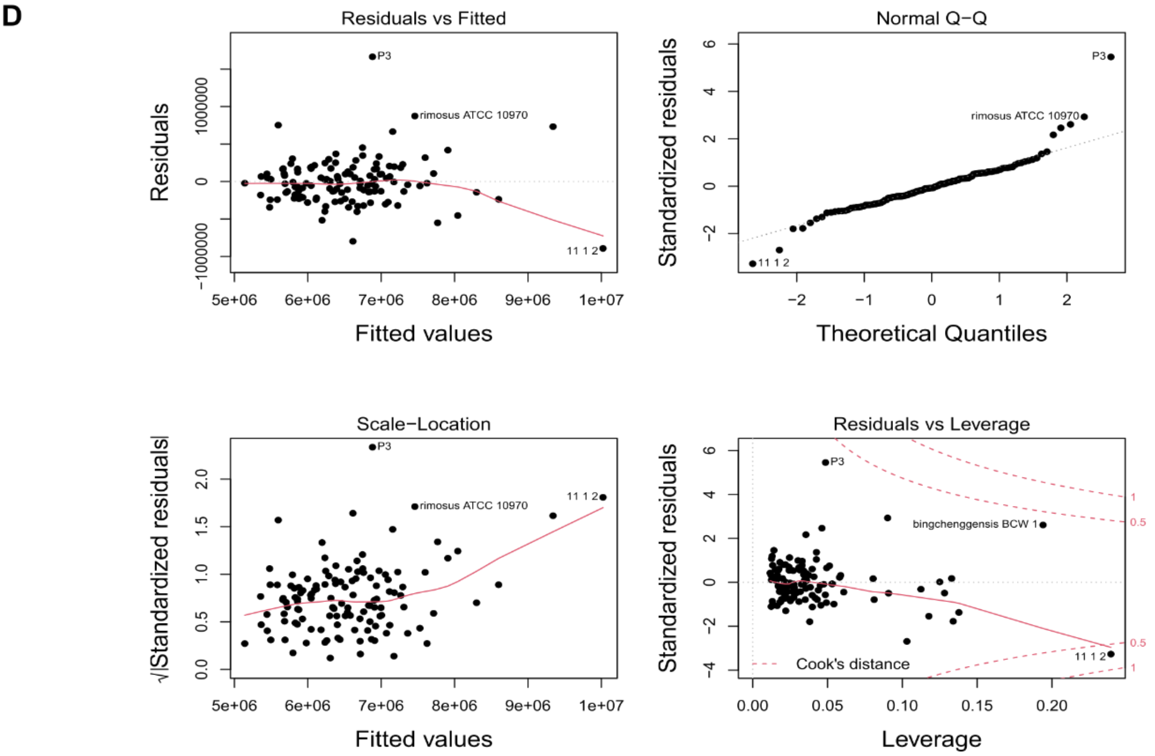


**Figure S4: Generalized linear model of the core region size based on the knowledge of the *rrn* operons (number,locations), the chromosome size and the *oriC* position**
**A. Schematic representation of the *Streptomyces* regions and distances analyzed in the study.** The *rrn* genes are always contained in the core region, while the relative positions of the extreme tDNA and core genes may differ from genome to genome (being thus different

from those represented in this panel). The location of these latter is therefore represented by bijective grey arrows. The cumulative distance between the extreme *rrn* and core genes was further termed ‘delta_core_*rrn*’. The maximal and minimal distances of *oriC* to the most extreme *rrn* (‘d_max_*rrn*_ori’ and ‘d_min_*rrn*_ori’) were also determined. Other abbreviations: ‘term_comp’ = terminal compartments (cumulative size of the left and right compartments), TIR = terminal inverted repeats.

**B. Summary of the ANOVA model based on the sizes of the central compartment and the chromosome, the maximal distance of *rrn* genes to the origin of replication and the number of *rrn* genes.** The explanatory variables are ranked according to their selection order in a forward regression approach. Code: ‘***’ ($p< 0.001$), ‘**’ ($p< 0.01$), ‘*’ ($p< 0.05$), ‘.’ ($p< 0.1$). AIC = Akaike Information Criterion

**C. Type I analysis of variance (ANOVA) sum test on the model**. In this type of test, the *p*-value of each explanatory variable depends on its order in the model. Code: ‘***’ ($p< 0.001$), ‘**’ ($p< 0.01$), ‘*’ ($p< 0.05$), ‘.’ ($p< 0.1$).

**D. Diagnostic scatterplots evaluating the assumptions of linearity, homoscedasticity and bivariate normality as well as the leverage effect.** The plot of the residuals *versus* fitted values addresses both the linearity and homoscedasticity (constant variability) assumptions: If the relationship is linear, the red line in the center of the graph is fairly flat, without distinct patterns; in case of homoscedasticity, the error is relatively constant across all of the fitted values. The assumption of bivariate normality is taken into account with a Quantile-Quantile (Q-Q) normal graph: In case of bivariate normality, the points should follow the straight dashed red line. The scale-location plot also diagnoses the homoscedasticity, a horizontal line with equally spread points being a good indicator. The last graph evaluates the ‘leverage effect’ i.e. allow to identify genomes that may change the regression line. Points presenting a Cook’s distance greater than 1 should be excluded from the model (no case in this study). Based on these analyses, genomes that show a rather unusual pattern are indicated on each graph.
