## Supplementary material for "Ribosomal RNA operons define a central functional compartment in the *Streptomyces* chromosome": Figure S5.pdf

Fig. S5 (1/15)

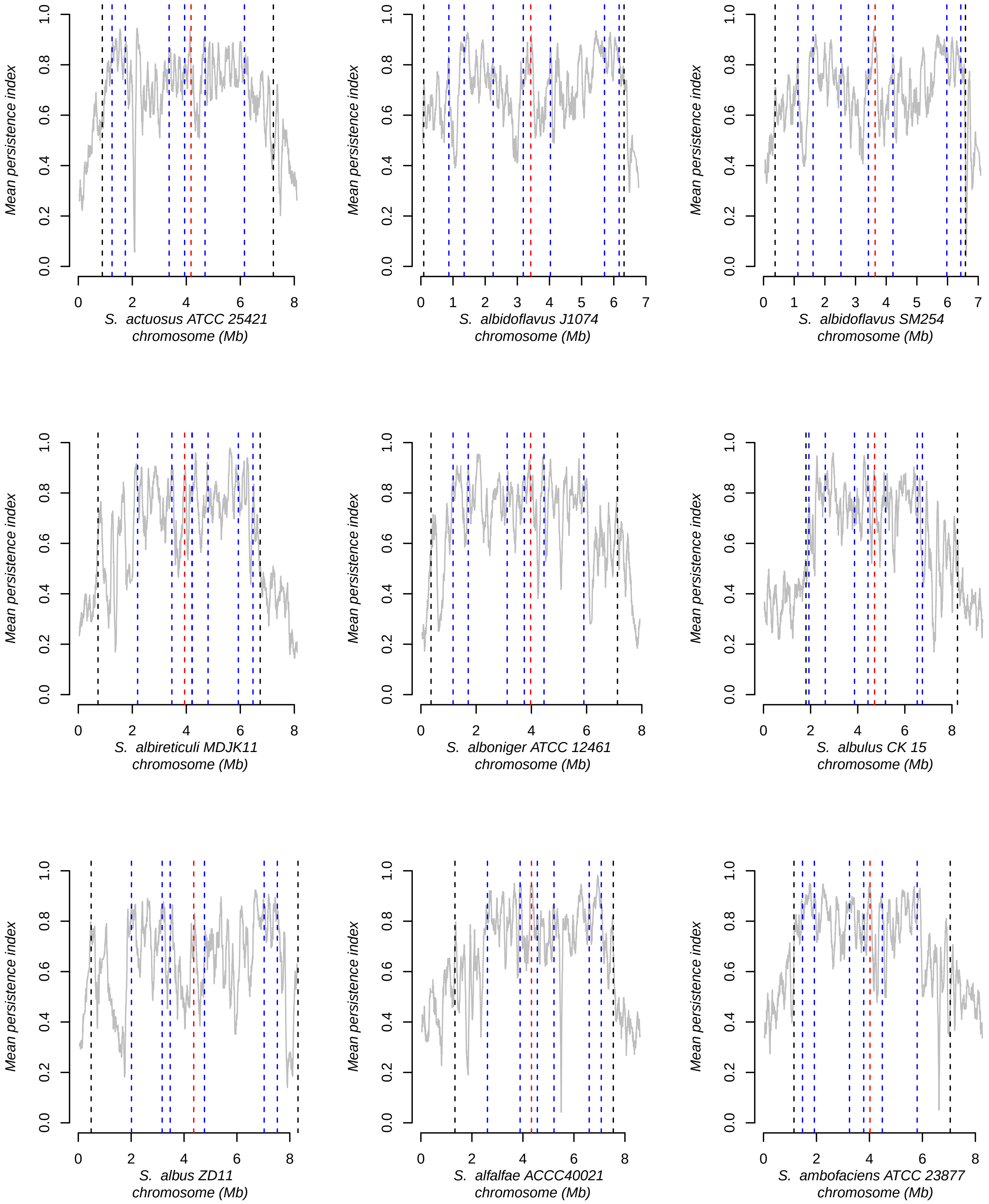
Mean persistence index
0.0
0.2
0.4
0.6
0.8
1.0
S. actuosus ATCC 25421
chromosome (Mb)
S. albidoflavus J1074
chromosome (Mb)
S. albidoflavus SM254
chromosome (Mb)
S. albireticuli MDJK11
chromosome (Mb)
S. alboniger ATCC 12461
chromosome (Mb)
S. albulus CK 15
chromosome (Mb)
S. albus ZD11
chromosome (Mb)
S. alfalfae ACCC40021
chromosome (Mb)
S. ambofaciens ATCC 23877
chromosome (Mb)

Fig. S5 (2/15)

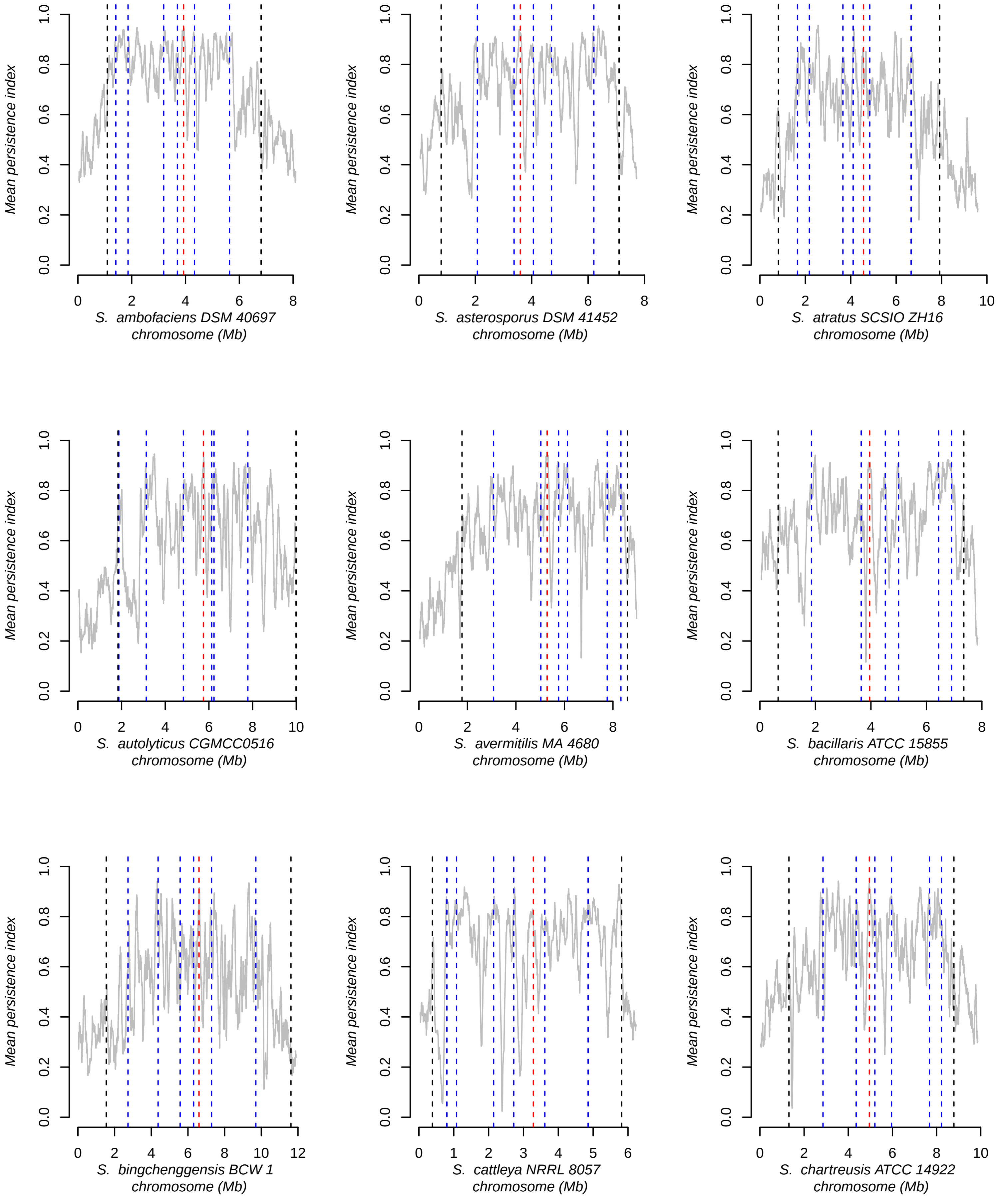
Mean persistence index
S. ambofaciens DSM 40697
chromosome (Mb)
S. asterosporus DSM 41452
chromosome (Mb)
S. atratus SCSIO ZH16
chromosome (Mb)
S. autolyticus CGMCC0516
chromosome (Mb)
S. avermitilis MA 4680
chromosome (Mb)
S. bacillaris ATCC 15855
chromosome (Mb)
S. bingchenggensis BCW 1
chromosome (Mb)
S. cattleya NRRL 8057
chromosome (Mb)
S. chartreusis ATCC 14922
chromosome (Mb)

Fig. S5 (3/15)

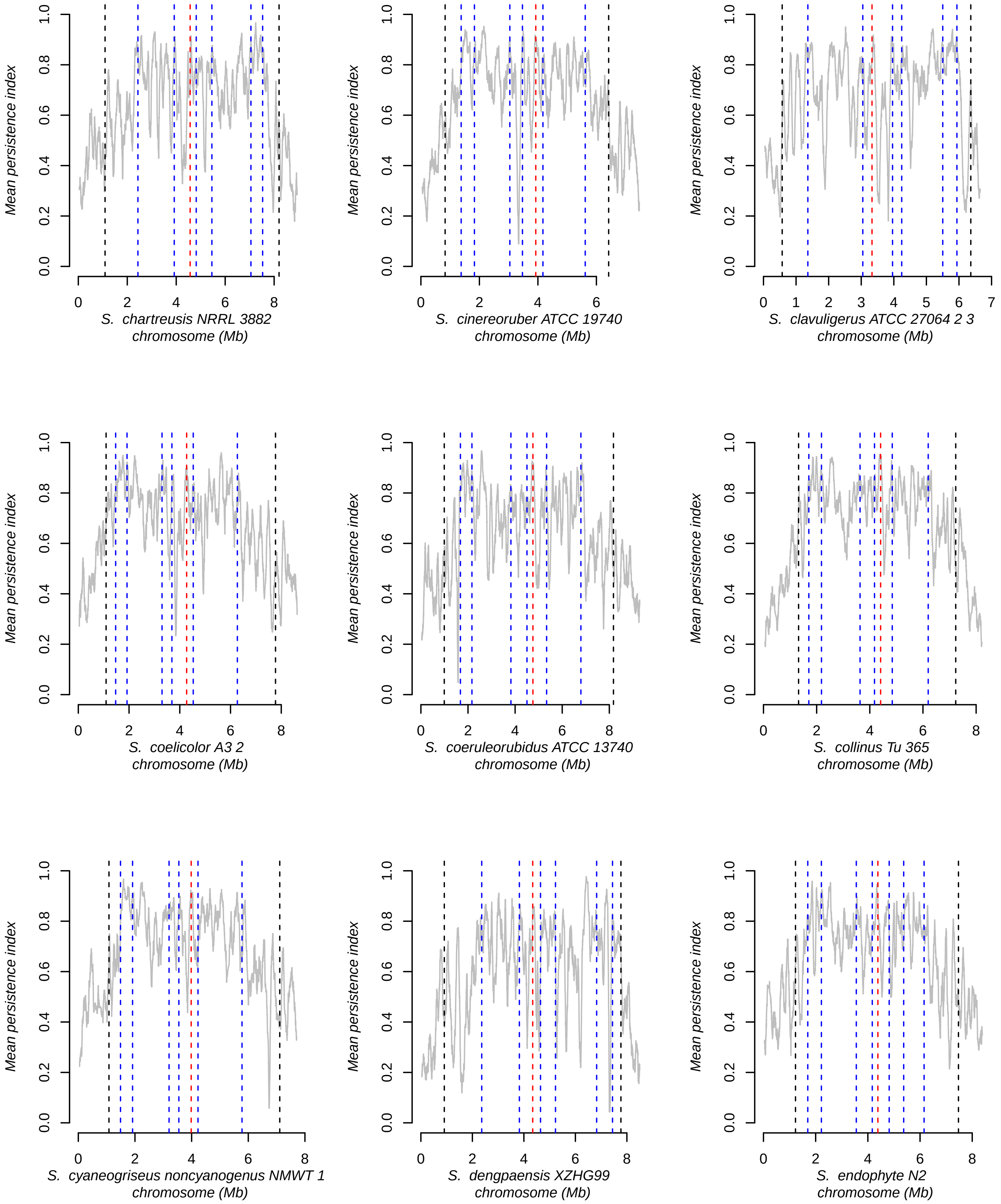

Mean persistence index
S. chartreusis NRRL 3882
chromosome (Mb)
S. cinereoruber ATCC 19740
chromosome (Mb)
S. clavuligerus ATCC 27064 2 3
chromosome (Mb)
S. coelicolor A3 2
chromosome (Mb)
S. coeruleorubidus ATCC 13740
chromosome (Mb)
S. collinus Tu 365
chromosome (Mb)
S. cyaneogriseus noncyanogenus NMWT 1
chromosome (Mb)
S. dengpaensis XZHG99
chromosome (Mb)
S. endophyte N2
chromosome (Mb)

Fig. S5 (4/15)

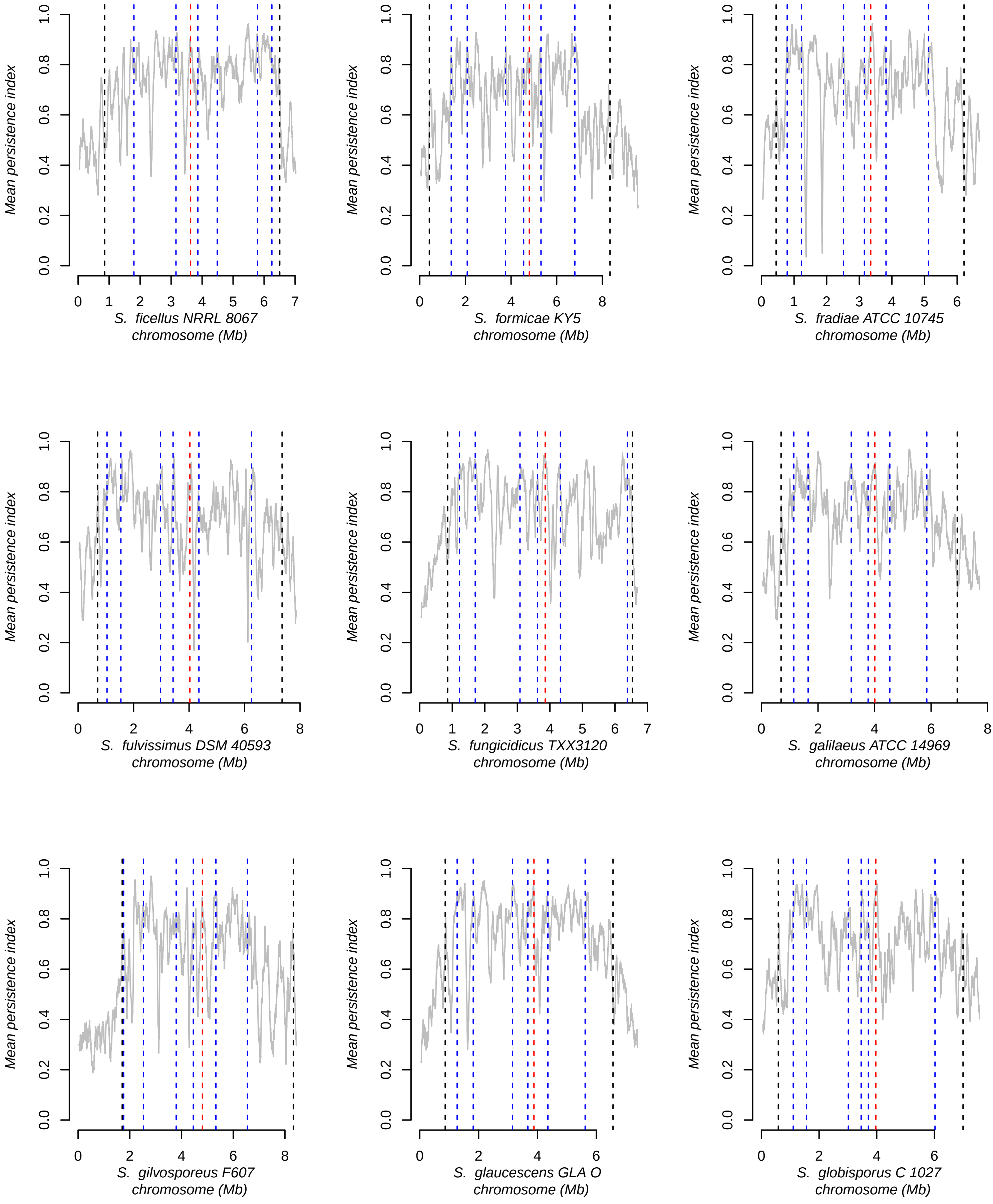
Mean persistence index
S. ficellus NRRL 8067
chromosome (Mb)
S. formicae KY5
chromosome (Mb)
S. fradiae ATCC 10745
chromosome (Mb)
S. fulvissimus DSM 40593
chromosome (Mb)
S. fungicidicus TXX3120
chromosome (Mb)
S. galilaeus ATCC 14969
chromosome (Mb)
S. gilvosporeus F607
chromosome (Mb)
S. glaucescens GLA O
chromosome (Mb)
S. globisporus C 1027
chromosome (Mb)

Fig. S5 (5/15)

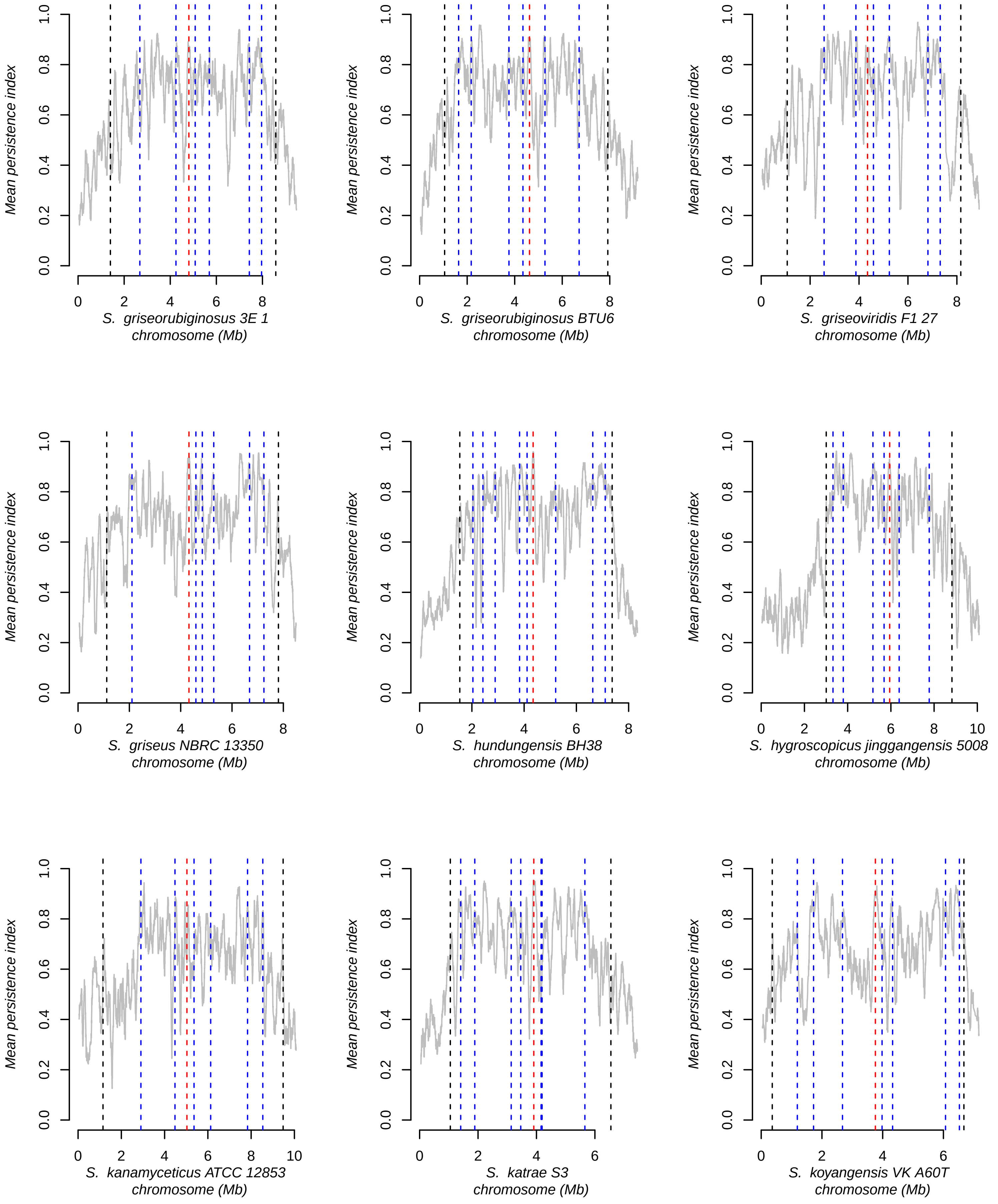
Mean persistence index
0.0
0.2
0.4
0.6
0.8
1.0
S. griseorubiginosus 3E 1
chromosome (Mb)
S. griseorubiginosus BTU6
chromosome (Mb)
S. griseoviridis F1 27
chromosome (Mb)
S. griseus NBRC 13350
chromosome (Mb)
S. hundungensis BH38
chromosome (Mb)
S. hygroscopicus jinggangensis 5008
chromosome (Mb)
S. kanamyceticus ATCC 12853
chromosome (Mb)
S. katrae S3
chromosome (Mb)
S. koyangensis VK A60T
chromosome (Mb)

Fig. S5 (6/15)

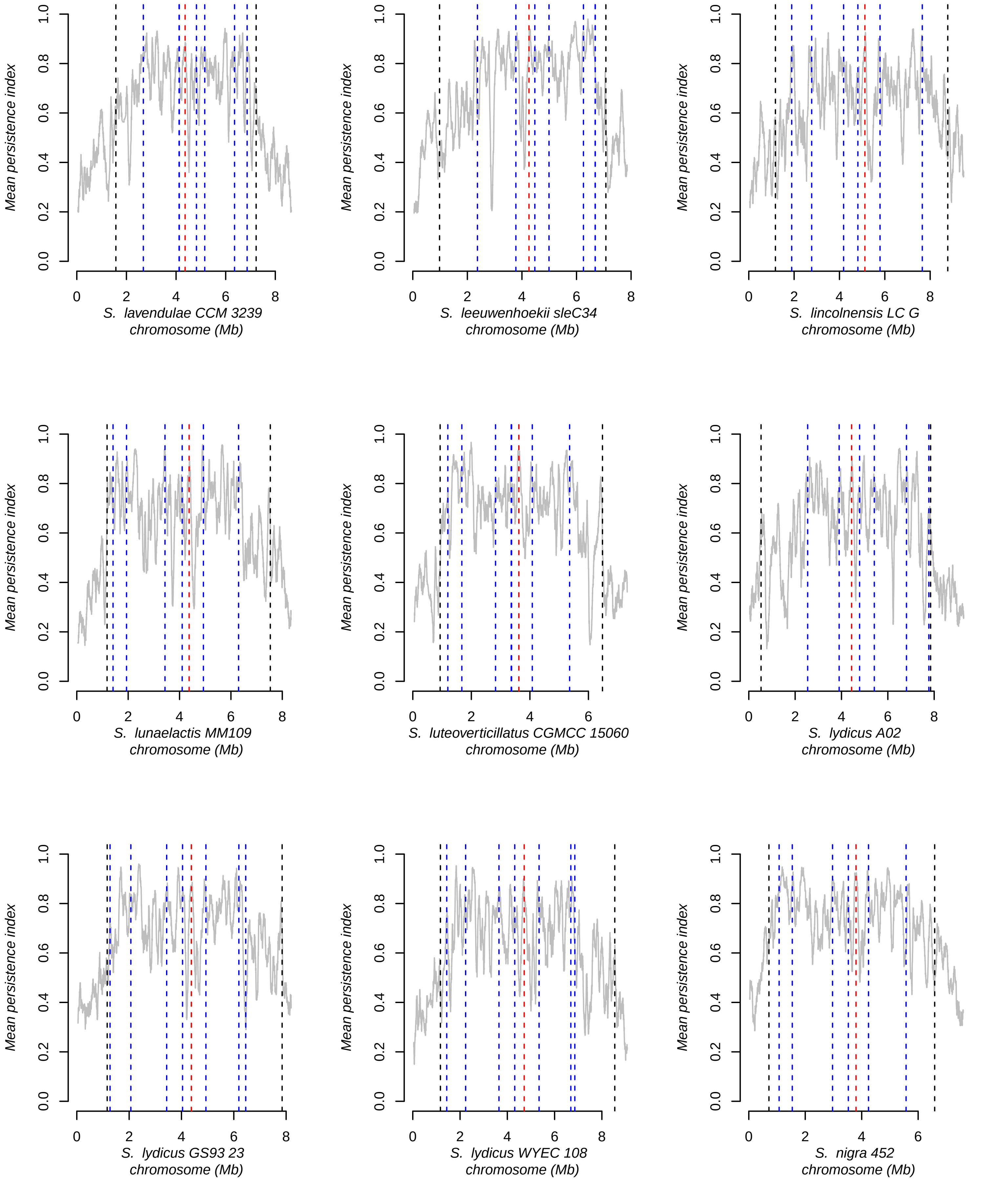
Mean persistence index
S. lavendulae CCM 3239
chromosome (Mb)
S. leeuwenhoekii sleC34
chromosome (Mb)
S. lincolnensis LC G
chromosome (Mb)
S. lunaelactis MM109
chromosome (Mb)
S. luteoverticillatus CGMCC 15060
chromosome (Mb)
S. lydicus A02
chromosome (Mb)
S. lydicus GS93 23
chromosome (Mb)
S. lydicus WYEC 108
chromosome (Mb)
S. nigra 452
chromosome (Mb)
0.0
0.2
0.4
0.6
0.8
1.0
0
2
4
6
8

Fig. S5 (7/15)

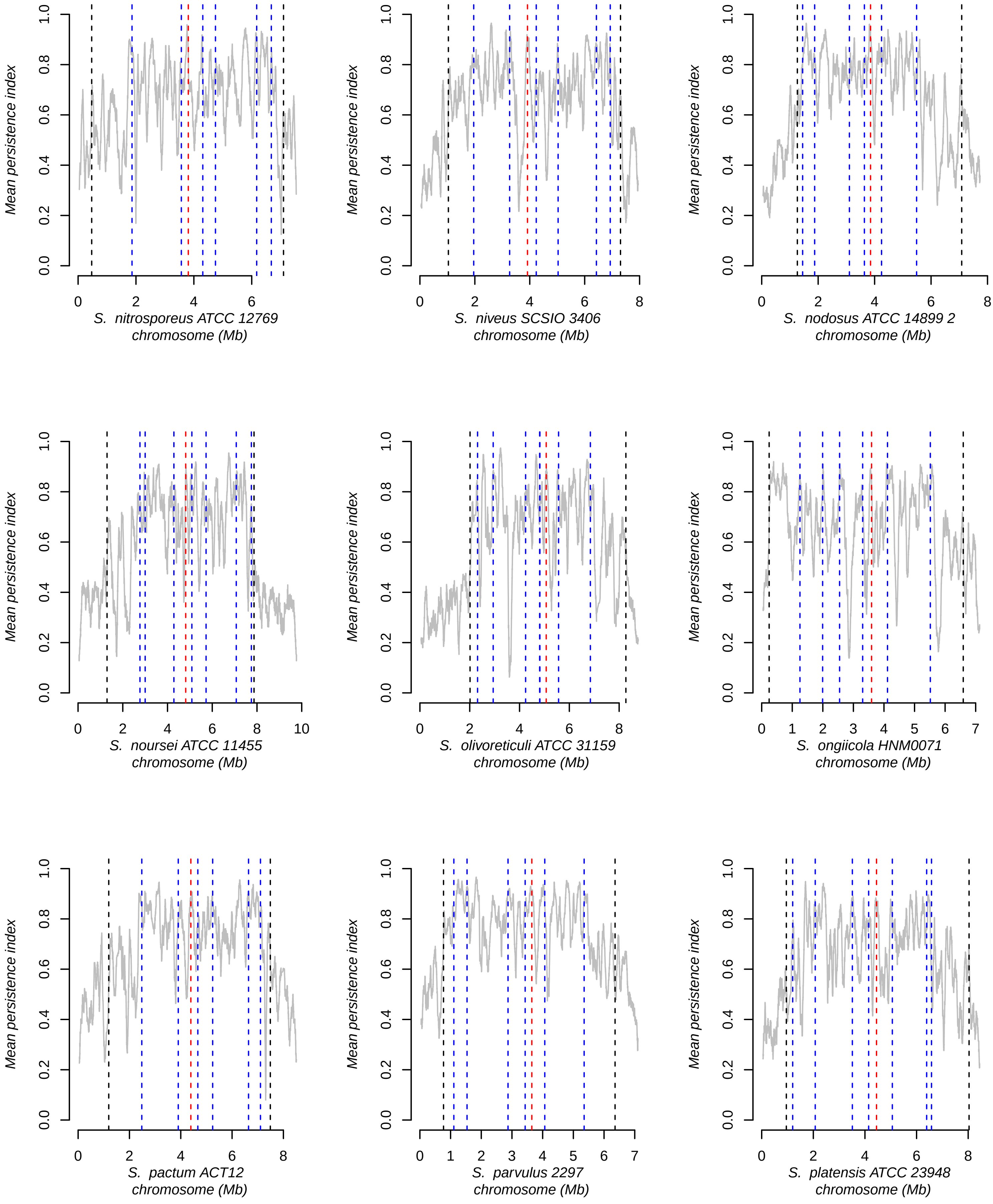
Mean persistence index
S. nitrosporeus ATCC 12769
chromosome (Mb)
S. niveus SCSIO 3406
chromosome (Mb)
S. nodosus ATCC 14899 2
chromosome (Mb)
S. noursei ATCC 11455
chromosome (Mb)
S. olivoreticuli ATCC 31159
chromosome (Mb)
S. ongiicola HNM0071
chromosome (Mb)
S. pactum ACT12
chromosome (Mb)
S. parvulus 2297
chromosome (Mb)
S. platensis ATCC 23948
chromosome (Mb)

Fig. S5 (8/15)

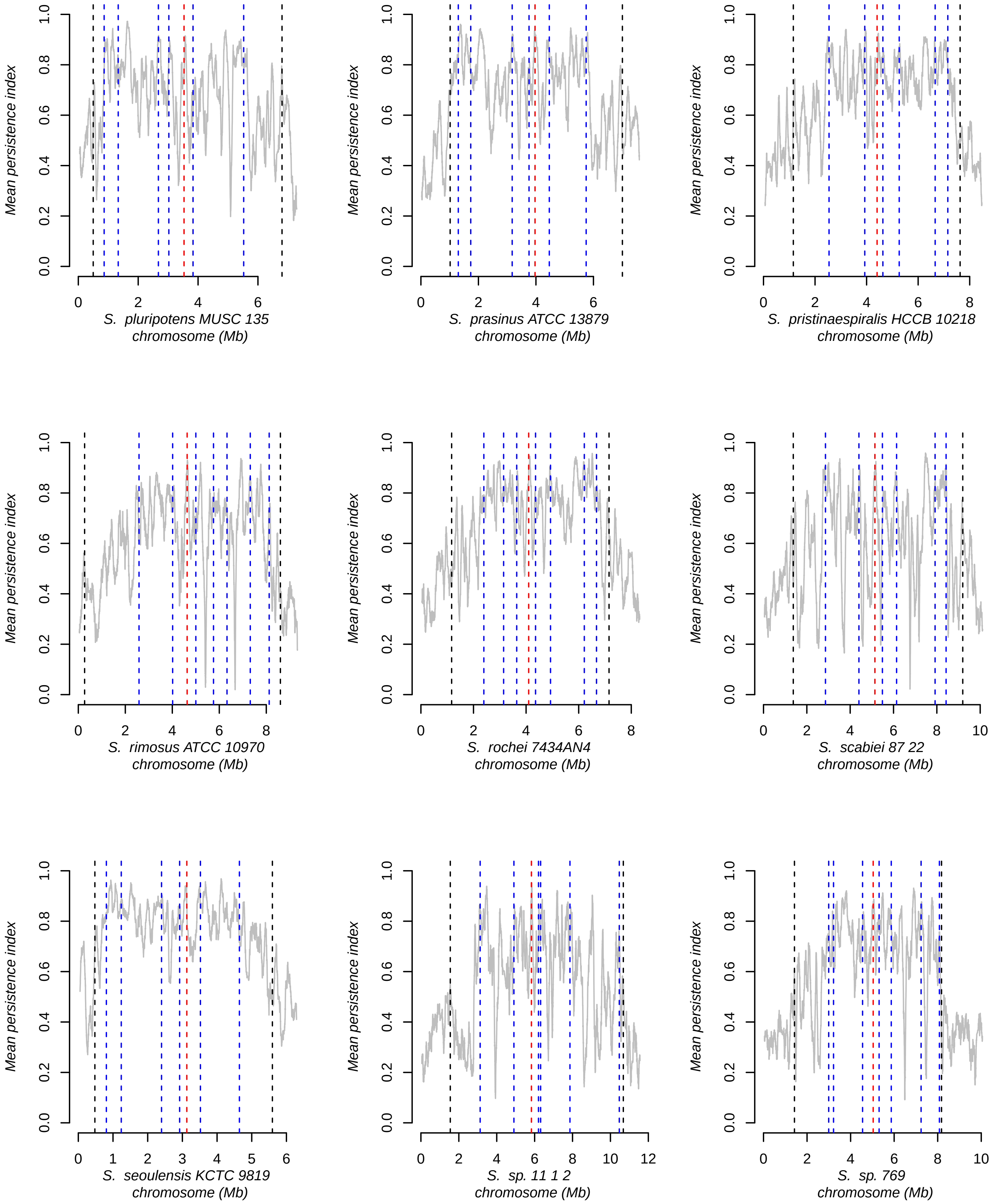

Mean persistence index
S. pluripotens MUSC 135
chromosome (Mb)
S. prasinus ATCC 13879
chromosome (Mb)
S. pristinaespiralis HCCB 10218
chromosome (Mb)
S. rimosus ATCC 10970
chromosome (Mb)
S. rochei 7434AN4
chromosome (Mb)
S. scabiei 87 22
chromosome (Mb)
S. seoulensis KCTC 9819
chromosome (Mb)
S. sp. 11 1 2
chromosome (Mb)
S. sp. 769
chromosome (Mb)

Fig. S5 (9/15)

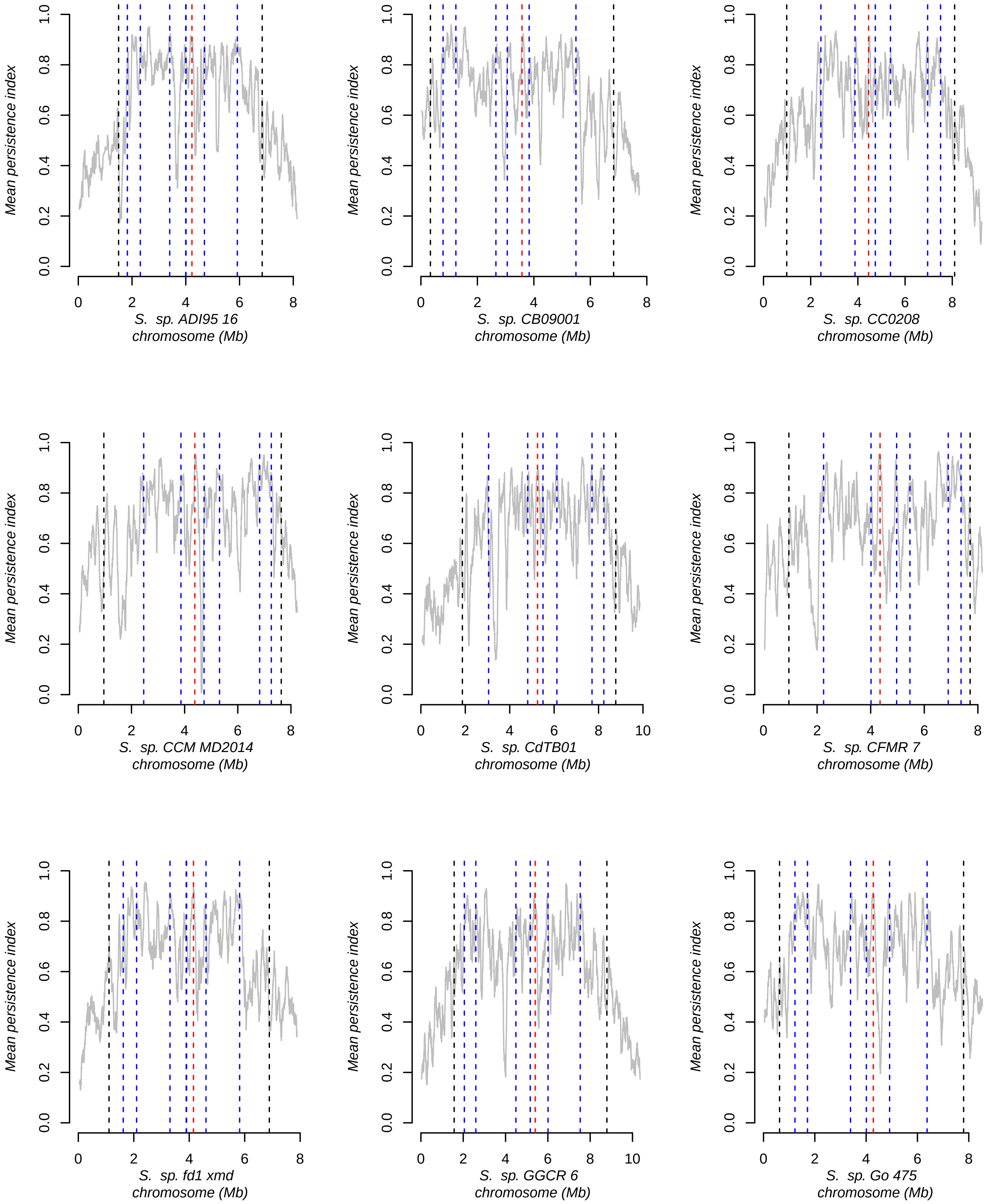

Mean persistence index
S. sp. ADI95 16
chromosome (Mb)
S. sp. CB09001
chromosome (Mb)
S. sp. CC0208
chromosome (Mb)
S. sp. CCM MD2014
chromosome (Mb)
S. sp. CdTB01
chromosome (Mb)
S. sp. CFMR 7
chromosome (Mb)
S. sp. fd1 xmd
chromosome (Mb)
S. sp. GGCR 6
chromosome (Mb)
S. sp. Go 475
chromosome (Mb)

Fig. S5 (10/15)

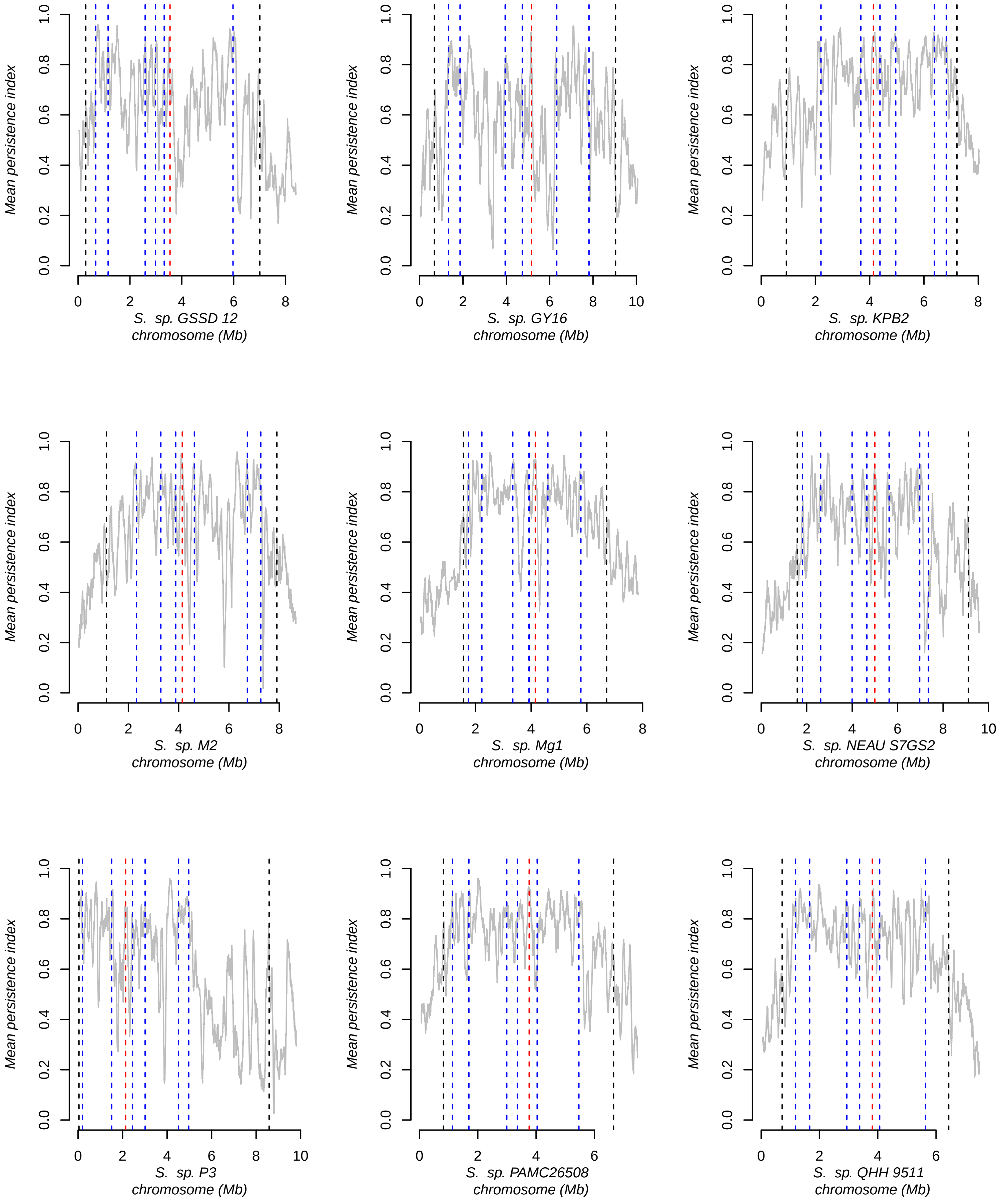
Mean persistence index
0.0
0.2
0.4
0.6
0.8
1.0
S. sp. GSSD 12
chromosome (Mb)
S. sp. GY16
chromosome (Mb)
S. sp. KPB2
chromosome (Mb)
S. sp. M2
chromosome (Mb)
S. sp. Mg1
chromosome (Mb)
S. sp. NEAU S7GS2
chromosome (Mb)
S. sp. P3
chromosome (Mb)
S. sp. PAMC26508
chromosome (Mb)
S. sp. QHH 9511
chromosome (Mb)
0
2
4
6
8
10

Fig. S5 (11/15)

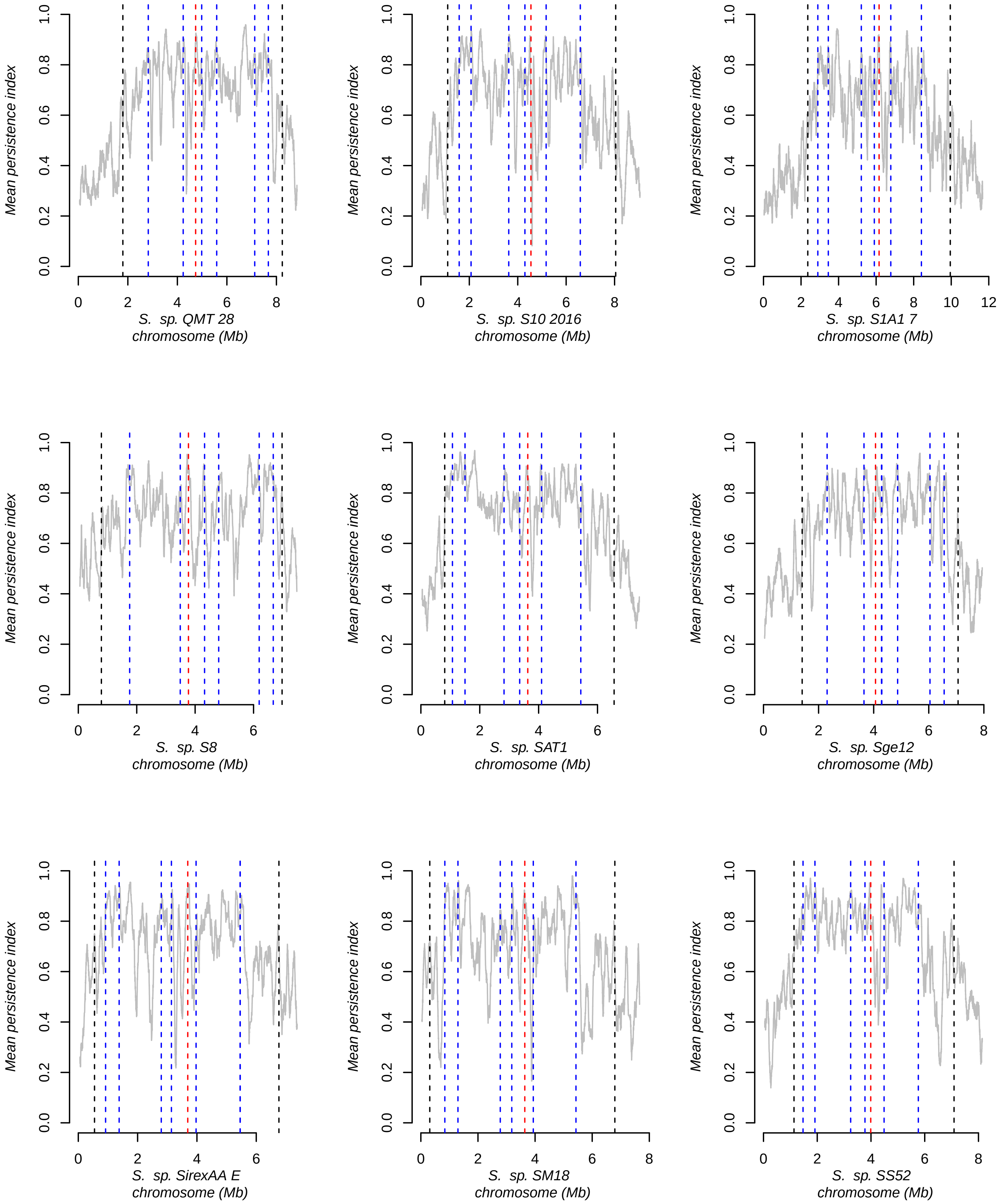
Mean persistence index
0.0
0.2
0.4
0.6
0.8
1.0
0
2
4
6
8
10
12
S. sp. QMT 28
chromosome (Mb)
S. sp. S10 2016
chromosome (Mb)
S. sp. S1A1 7
chromosome (Mb)
S. sp. S8
chromosome (Mb)
S. sp. SAT1
chromosome (Mb)
S. sp. Sge12
chromosome (Mb)
S. sp. SirexAA E
chromosome (Mb)
S. sp. SM18
chromosome (Mb)
S. sp. SS52
chromosome (Mb)

Fig. S5 (12/15)

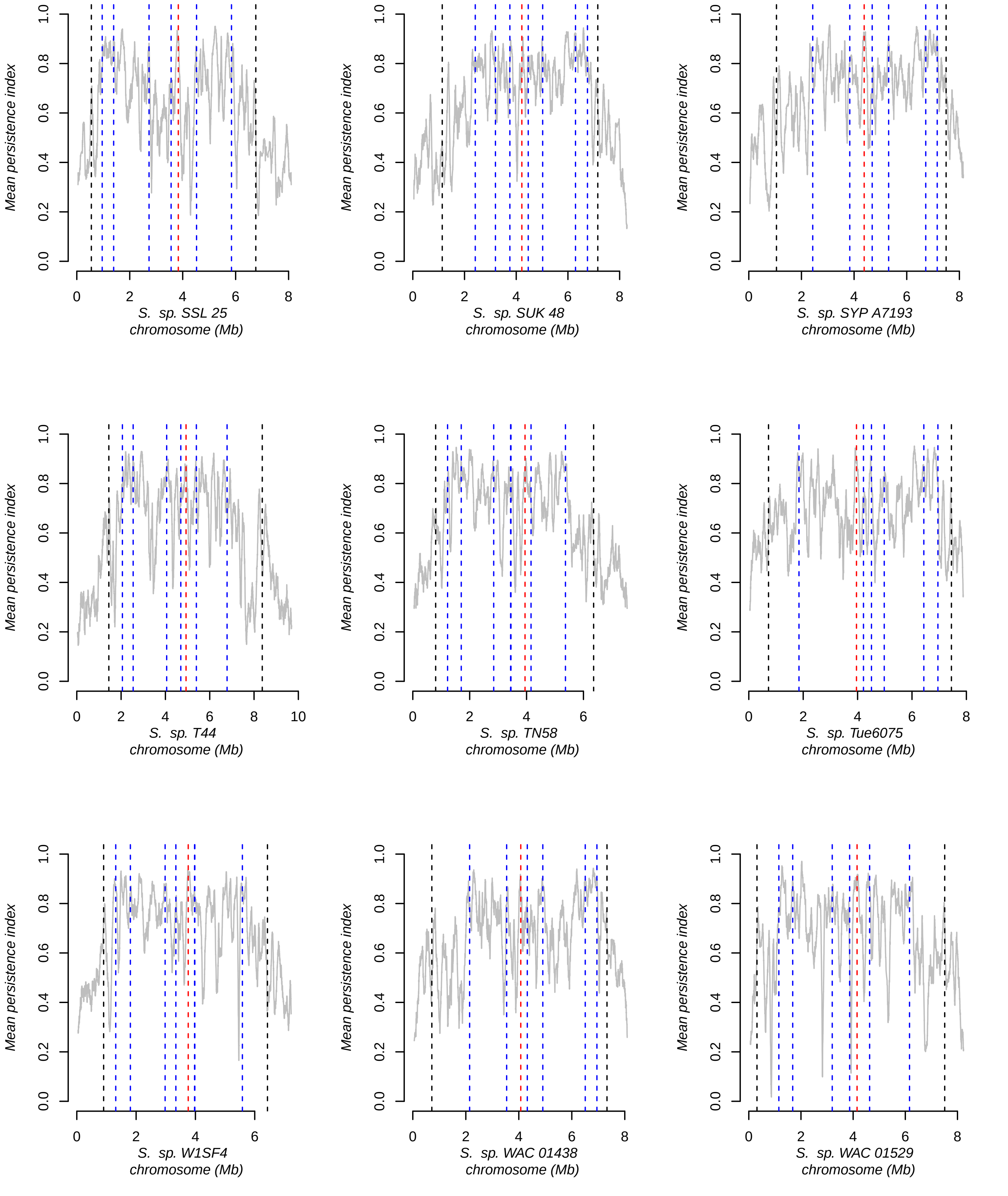
Mean persistence index
S. sp. SSL 25
chromosome (Mb)
S. sp. SUK 48
chromosome (Mb)
S. sp. SYP A7193
chromosome (Mb)
S. sp. T44
chromosome (Mb)
S. sp. TN58
chromosome (Mb)
S. sp. Tue6075
chromosome (Mb)
S. sp. W1SF4
chromosome (Mb)
S. sp. WAC 01438
chromosome (Mb)
S. sp. WAC 01529
chromosome (Mb)

Fig. S5 (13/15)

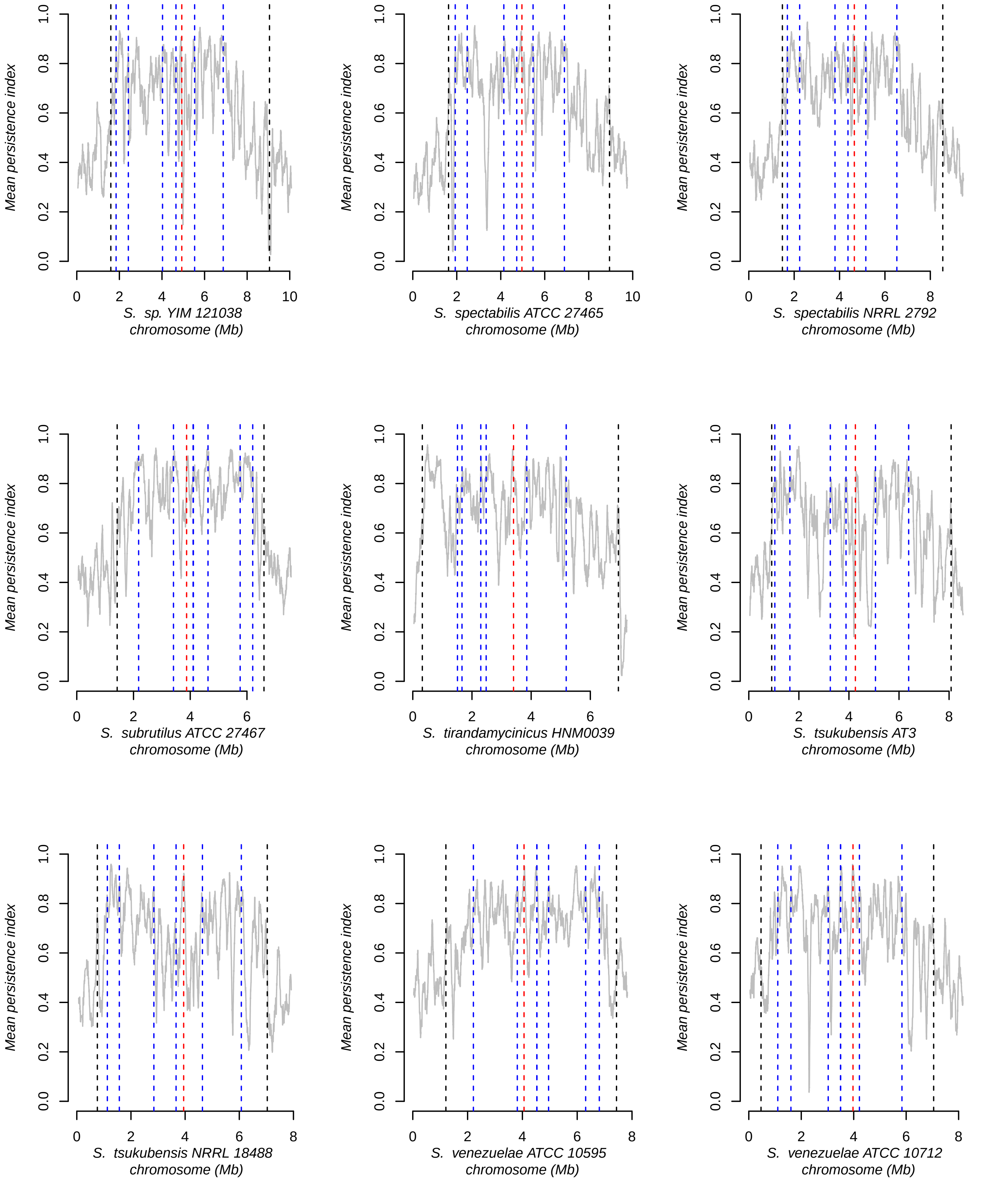
Mean persistence index
0.0
0.2
0.4
0.6
0.8
1.0
0
2
4
6
8
10
S. sp. YIM 121038
chromosome (Mb)
S. spectabilis ATCC 27465
chromosome (Mb)
S. spectabilis NRRL 2792
chromosome (Mb)
S. subrutilus ATCC 27467
chromosome (Mb)
S. tirandamycinicus HNM0039
chromosome (Mb)
S. tsukubensis AT3
chromosome (Mb)
S. tsukubensis NRRL 18488
chromosome (Mb)
S. venezuelae ATCC 10595
chromosome (Mb)
S. venezuelae ATCC 10712
chromosome (Mb)

Fig. S5 (14/15)

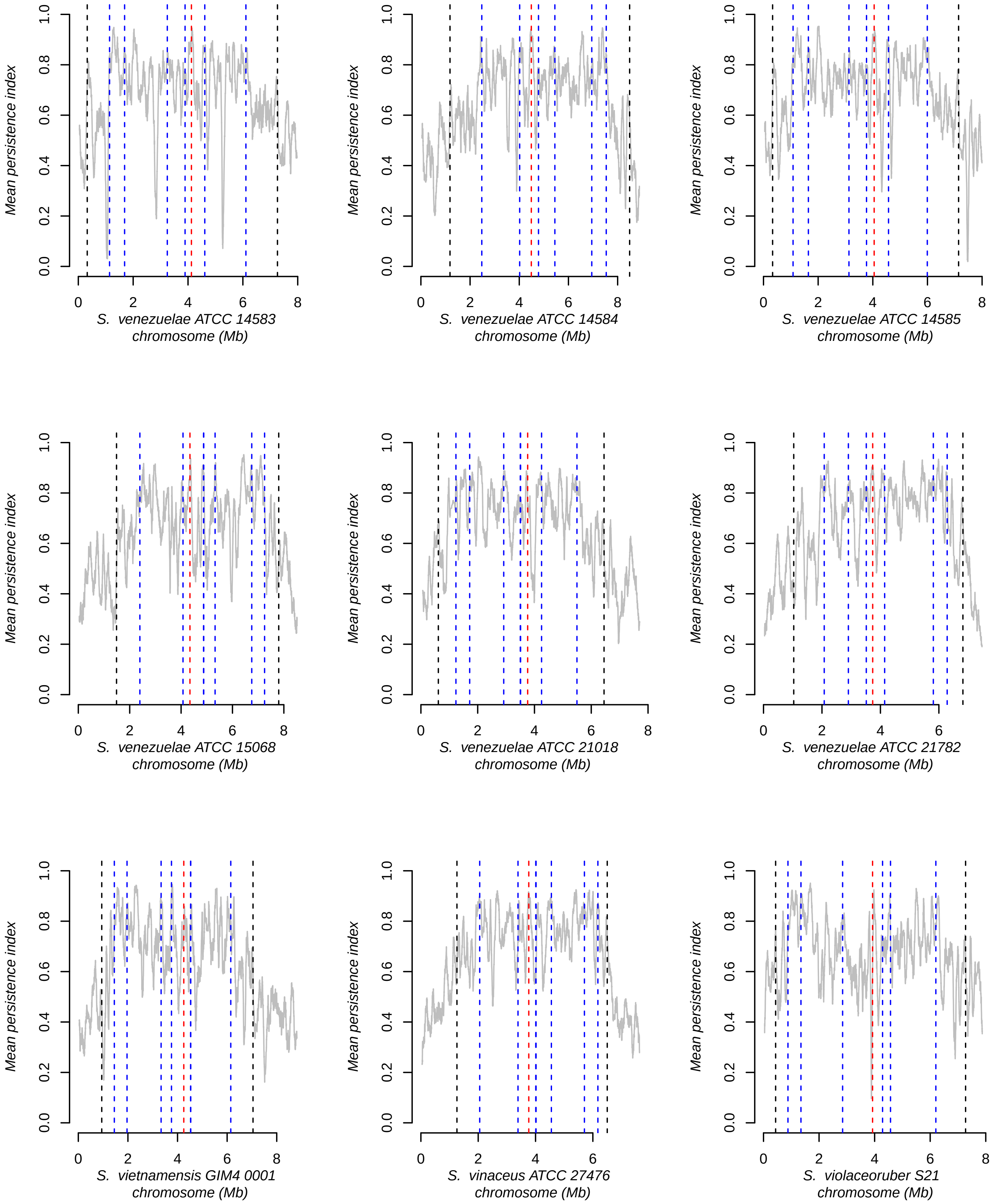
Mean persistence index
S. venezuelae ATCC 14583 chromosome (Mb)
S. venezuelae ATCC 14584 chromosome (Mb)
S. venezuelae ATCC 14585 chromosome (Mb)
S. venezuelae ATCC 15068 chromosome (Mb)
S. venezuelae ATCC 21018 chromosome (Mb)
S. venezuelae ATCC 21782 chromosome (Mb)
S. vietnamensis GIM4 0001 chromosome (Mb)
S. vinaceus ATCC 27476 chromosome (Mb)
S. violaceoruber S21 chromosome (Mb)

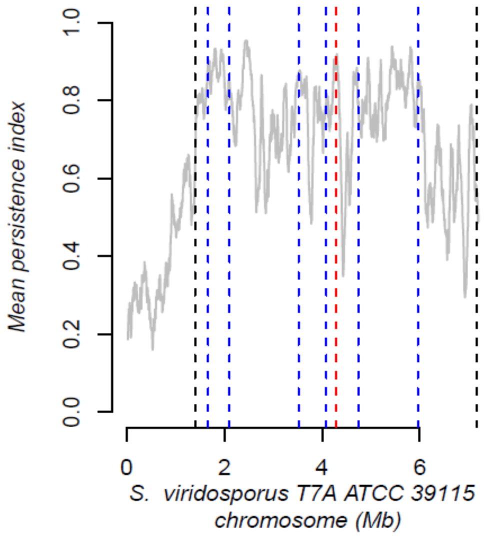


**Figure S5: Gene persistence along the chromosomes of 127 *Streptomyces* strains/species**

The level of persistence (*y*-axis) along the chromosome (*x*-axis) is represented using a sliding window [81 coding sequences (CDSs), with 1 CDS steps]. The positions of the most external core genes, all *rrn* operons and the origin of replication are indicated by black, blue and red dashed lines, respectively. The sequences released from the databases (**Supplemental Table S1**) were directly analyzed, without changing their orientation. To decide whether a sequence from the database is in the same orientation as the reference consensus used in this study (*Streptomyces viridosporus* T7A ATCC 39115), it is necessary to consider the results of pairwise comparison presented in **Supplemental Figure S1**: when the diagonal starts at the bottom left and ends at the top right, it means that the genome sequence available in the databases is oriented as in the consensus, and that the *rrn* configuration shown below each graph can be directly transposed onto the graph. If not, the sequence is in the opposite direction. In this case, the consensus should be reversed when transposed on the graph (e.g. '*rrn* ABCDEF *dnaA*+' becomes '*rrn* F*E*D*C*B*A* *dnaA*-').
