## Supplementary material for "Ribosomal RNA operons define a central functional compartment in the *Streptomyces* chromosome": Figure S6.pdf

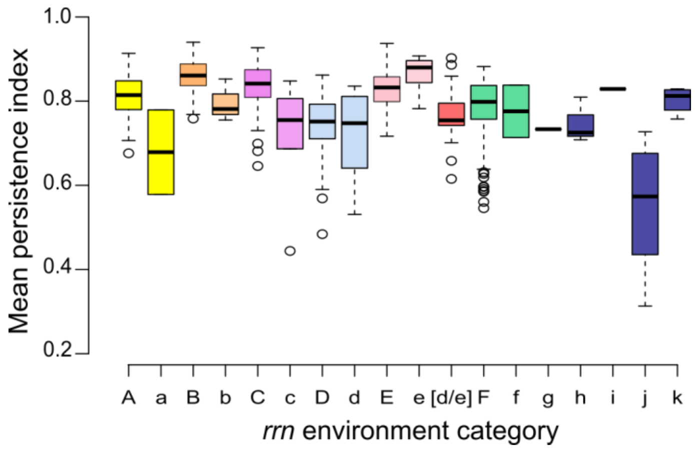


**Figure S6: Mean gene persistence in the regions surrounding the *rrn* operons**

The mean persistence was calculated within a window of 81 CDS centered on the *rrn* operon of interest. The boxplot represents the first quartile, median and third quartile. The upper whisker extends from the hinge to the largest value no further than 1.5 * the inter-quartile range (IQR, i.e. distance between the first and third quartiles) from the hinge. The lower whisker extends from the hinge to the smallest value at most 1.5 * IQR of the hinge. Outliers are represented (dots). The number of *rrn* operons in each category is presented in **Fig.1.C**.
