## Supplementary material for "Ribosomal RNA operons define a central functional compartment in the *Streptomyces* chromosome": Figure S7.pdf

**A**

| Processes | | Adjusted *p* value | # |
|---|---|---|---|
| metabolic process | GO:0008152 | 1,6E-36 | 425 |
| organonitrogen compound metabolic process | GO:1901564 | 7,4E-32 | 243 |
| biosynthetic process | GO:0009058 | 1,2E-31 | 254 |
| organic substance metabolic process | GO:0071704 | 2,5E-30 | 368 |
| cellular metabolic process | GO:0044237 | 2,2E-29 | 281 |
| organonitrogen compound biosynthetic process | GO:1901566 | 4,4E-26 | 123 |
| cellular biosynthetic process | GO:0044249 | 2,2E-24 | 207 |
| carboxylic acid metabolic process | GO:0019752 | 1,1E-23 | 106 |
| primary metabolic process | GO:0044238 | 5,6E-21 | 275 |
| cellular amino acid metabolic process | GO:0006520 | 8,7E-21 | 84 |
| nitrogen compound metabolic process | GO:0006807 | 1,2E-20 | 171 |
| cellular process | GO:0009987 | 2,5E-20 | 363 |
| organic substance biosynthetic process | GO:1901576 | 7,2E-20 | 160 |
| oxoacid metabolic process | GO:0043436 | 1,2E-16 | 89 |
| cellular amino acid biosynthetic process | GO:0008652 | 4,1E-16 | 59 |
| small molecule metabolic process | GO:0044281 | 1,6E-15 | 111 |
| purine nucleotide metabolic process | GO:0006163 | 7,5E-15 | 35 |
| purine nucleotide biosynthetic process | GO:0006164 | 8,5E-15 | 33 |
| alpha-amino acid metabolic process | GO:1901605 | 7,8E-14 | 59 |
| purine ribonucleotide biosynthetic process | GO:0009152 | 4,8E-13 | 30 |
| metal ion binding | GO:0046872 | 1,9E-12 | 159 |
| cation binding | GO:0043169 | 3,5E-11 | 152 |
| alpha-amino acid biosynthetic process | GO:1901607 | 4,5E-11 | 41 |
| rRNA binding | GO:0019843 | 7,3E-11 | 31 |
| cytosolic large ribosomal subunit | GO:0022625 | 1,2E-10 | 17 |
| nucleotide binding | GO:0000166 | 1,6E-10 | 169 |
| nucleotide biosynthetic process | GO:0009165 | 1,9E-10 | 32 |
| macromolecule metabolic process | GO:0043170 | 2,4E-10 | 178 |
| nucleotide metabolic process | GO:0009117 | 3,3E-10 | 38 |
| organic acid biosynthetic process | GO:0016053 | 4,9E-10 | 45 |
| ligase activity | GO:0016874 | 3,0E-09 | 54 |
| RNA binding | GO:0003723 | 3,2E-09 | 70 |
| catalytic activity, acting on RNA | GO:0140098 | 5,7E-09 | 38 |
| catalytic activity | GO:0003824 | 1,0E-08 | 484 |
| small ribosomal subunit | GO:0015935 | 1,8E-08 | 16 |
| translation | GO:0006412 | 2,0E-08 | 68 |
| ribosome | GO:0005840 | 3,4E-08 | 16 |
| tRNA binding | GO:0000049 | 3,7E-08 | 19 |
| cellular macromolecule biosynthetic process | GO:0034645 | 5,5E-08 | 97 |
| peptide biosynthetic process | GO:0043043 | 7,6E-08 | 67 |
| magnesium ion binding | GO:0000287 | 8,3E-08 | 42 |
| cellular aromatic compound metabolic process | GO:0006725 | 1,0E-07 | 79 |
| cytosolic small ribosomal subunit | GO:0022627 | 1,1E-07 | 11 |
| large ribosomal subunit | GO:0015934 | 1,3E-07 | 17 |
| organic acid metabolic process | GO:0006082 | 1,6E-07 | 56 |
| organic cyclic compound biosynthetic process | GO:1901362 | 1,6E-07 | 74 |
| sulfur compound metabolic process | GO:0006790 | 2,0E-07 | 32 |
| protein metabolic process | GO:0019538 | 2,9E-07 | 106 |
| ribonucleoprotein complex | GO:1990904 | 3,4E-07 | 18 |
| ATP metabolic process | GO:0046034 | 1,7E-06 | 18 |
| tRNA metabolic process | GO:0006399 | 1,7E-06 | 23 |
| tRNA aminoacylation | GO:0043039 | 2,4E-06 | 18 |
| nucleoside phosphate metabolic process | GO:0006753 | 3,9E-06 | 29 |
| ATP binding | GO:0005524 | 4,7E-06 | 126 |
| ribonucleoside monophosphate biosynthetic process | GO:0009156 | 4,8E-06 | 16 |
| cellular protein metabolic process | GO:0044267 | 8,1E-06 | 90 |
| nucleobase-containing compound metabolic process | GO:0006139 | 8,2E-06 | 72 |
| NADH dehydrogenase activity | GO:0003954 | 8,7E-06 | 17 |
| ncRNA processing | GO:0034470 | 1,2E-05 | 22 |
| adenyl ribonucleotide binding | GO:0032559 | 1,3E-05 | 126 |

| Processes | | Adjusted *p* value | # |
|---|---|---|---|
| lyase activity | GO:0016829 | 3,2E-05 | 47 |
| aerobic respiration | GO:0009060 | 3,8E-05 | 21 |
| quinone binding | GO:0048038 | 5,6E-05 | 15 |
| tRNA aminoacylation for protein translation | GO:0006418 | 6,1E-05 | 16 |
| RNA processing | GO:0006396 | 7,4E-05 | 20 |
| nucleoside-triphosphatase activity | GO:0017111 | 7,6E-05 | 31 |
| generation of precursor metabolites and energy | GO:0006091 | 7,7E-05 | 28 |
| aminoacyl-tRNA ligase activity | GO:0004812 | 1,1E-04 | 16 |
| IMP metabolic process | GO:0046040 | 1,2E-04 | 12 |
| pyrophosphatase activity | GO:0016462 | 1,7E-04 | 29 |
| oxidoreductase activity, acting on NAD(P)H | GO:0016651 | 1,8E-04 | 19 |
| oxidoreduction-driven active transmembrane transporter activity | GO:0015453 | 1,9E-04 | 17 |
| branched-chain amino acid metabolic process | GO:0009081 | 2,0E-04 | 14 |
| purine ribonucleoside monophosphate biosynthetic process | GO:0009168 | 2,1E-04 | 15 |
| transferase activity | GO:0016740 | 2,2E-04 | 158 |
| hydrolase activity, acting on acid anhydrides, in phosphorus-containing anhydrides | GO:0016818 | 2,2E-04 | 30 |
| RNA modification | GO:0009451 | 2,3E-04 | 19 |
| phosphorus metabolic process | GO:0006793 | 2,8E-04 | 57 |
| ribosome biogenesis | GO:0042254 | 3,6E-04 | 16 |
| electron transfer activity | GO:0009055 | 4,4E-04 | 24 |
| cell cycle | GO:0007049 | 9,1E-04 | 15 |
| NADH dehydrogenase (ubiquinone) activity | GO:0008137 | 1,0E-03 | 13 |
| histidine biosynthetic process | GO:0000105 | 1,2E-03 | 10 |
| electron transport chain | GO:0022900 | 1,6E-03 | 23 |
| histidine metabolic process | GO:0006547 | 1,6E-03 | 11 |
| NADH dehydrogenase (quinone) activity | GO:0050136 | 2,3E-03 | 14 |
| ATP synthesis coupled proton transport | GO:0015986 | 2,3E-03 | 8 |
| proton-transporting ATP synthase activity, rotational mechanism | GO:0046933 | 2,3E-03 | 8 |
| branched-chain amino acid biosynthetic process | GO:0009082 | 2,5E-03 | 12 |
| pyruvate metabolic process | GO:0006090 | 2,6E-03 | 16 |
| 'de novo' IMP biosynthetic process | GO:0006189 | 2,8E-03 | 10 |
| proton-transporting ATP synthase complex | GO:0045259 | 4,1E-03 | 8 |
| tRNA modification | GO:0006400 | 4,7E-03 | 12 |
| proton transmembrane transport | GO:1902600 | 4,7E-03 | 17 |
| cellular respiration | GO:0045333 | 5,0E-03 | 16 |
| tRNA processing | GO:0008033 | 6,2E-03 | 14 |
| cellular amino acid catabolic process | GO:0009063 | 6,9E-03 | 17 |
| phosphate ion transport | GO:0006817 | 7,0E-03 | 8 |
| cell division | GO:0051301 | 7,5E-03 | 18 |
| transition metal ion binding | GO:0046914 | 8,5E-03 | 57 |
| ATP hydrolysis activity | GO:0016887 | 8,6E-03 | 22 |
| ATP biosynthetic process | GO:0006754 | 1,1E-02 | 8 |
| ligase activity, forming carbon-nitrogen bonds | GO:0016879 | 1,6E-02 | 21 |
| channel activity | GO:0015267 | 1,7E-02 | 10 |
| oxidative phosphorylation | GO:0006119 | 1,7E-02 | 10 |
| NAD(P)H dehydrogenase (quinone) activity | GO:0003955 | 1,7E-02 | 10 |
| iron-sulfur cluster assembly | GO:0016226 | 1,7E-02 | 7 |
| pentosyltransferase activity | GO:0016763 | 1,8E-02 | 12 |
| rRNA processing | GO:0006364 | 1,8E-02 | 12 |
| tryptophan metabolic process | GO:0006568 | 2,2E-02 | 9 |
| ATP synthesis coupled electron transport | GO:0042773 | 2,7E-02 | 8 |
| oxidoreductase activity, acting on NAD(P)H, quinone or similar compound as acceptor | GO:0016655 | 2,8E-02 | 11 |
| sulfur amino acid metabolic process | GO:0000096 | 3,7E-02 | 13 |
| protein catabolic process | GO:0030163 | 4,1E-02 | 10 |
| glycolytic process | GO:0006096 | 4,1E-02 | 10 |
| phosphate ion transmembrane transport | GO:0035435 | 4,1E-02 | 5 |
| folic acid-containing compound metabolic process | GO:0006760 | 4,3E-02 | 9 |
| hydro-lyase activity | GO:0016836 | 4,3E-02 | 18 |
| respiratory electron transport chain | GO:0022904 | 4,6E-02 | 11 |
| zinc ion binding | GO:0008270 | 4,7E-02 | 35 |

**B**

| Processes | | Adjusted *p* value | # |
|---|---|---|---|
| phospholipid biosynthetic process | GO:0008654 | 7,7E-04 | 7 |
| phospholipid metabolic process | GO:0006644 | 1,9E-03 | 7 |
| cellular lipid metabolic process | GO:0044255 | 2,5E-03 | 11 |
| catalytic activity | GO:0003824 | 5,6E-03 | 70 |
| glycerophospholipid biosynthetic process | GO:0046474 | 7,9E-03 | 4 |
| glycerophospholipid metabolic process | GO:0006650 | 1,3E-02 | 4 |
| nucleobase-containing compound metabolic process | GO:0006139 | 2,7E-02 | 15 |
| lipid metabolic process | GO:0006629 | 3,0E-02 | 10 |
| lipid biosynthetic process | GO:0008610 | 4,0E-02 | 9 |
| organic substance metabolic process | GO:0071704 | 4,5E-02 | 45 |
| base-excision repair | GO:0097510 | 4,9E-02 | 2 |

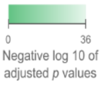


**Figure S7: GO enrichment analysis of the core genes which are mainly located in the central compartment (A) or in the terminal compartments (B)**

The statistically enriched processes were identified by a g:Profiler analysis including 748 genes whose GO annotation was available. The symbol "#" indicates the number of these core genes in each GO category. Some genes belong to several GO categories. The same scale was used for both panels.
