## Supplementary material for "Ribosomal RNA operons define a central functional compartment in the *Streptomyces* chromosome": Figure S8.pdf

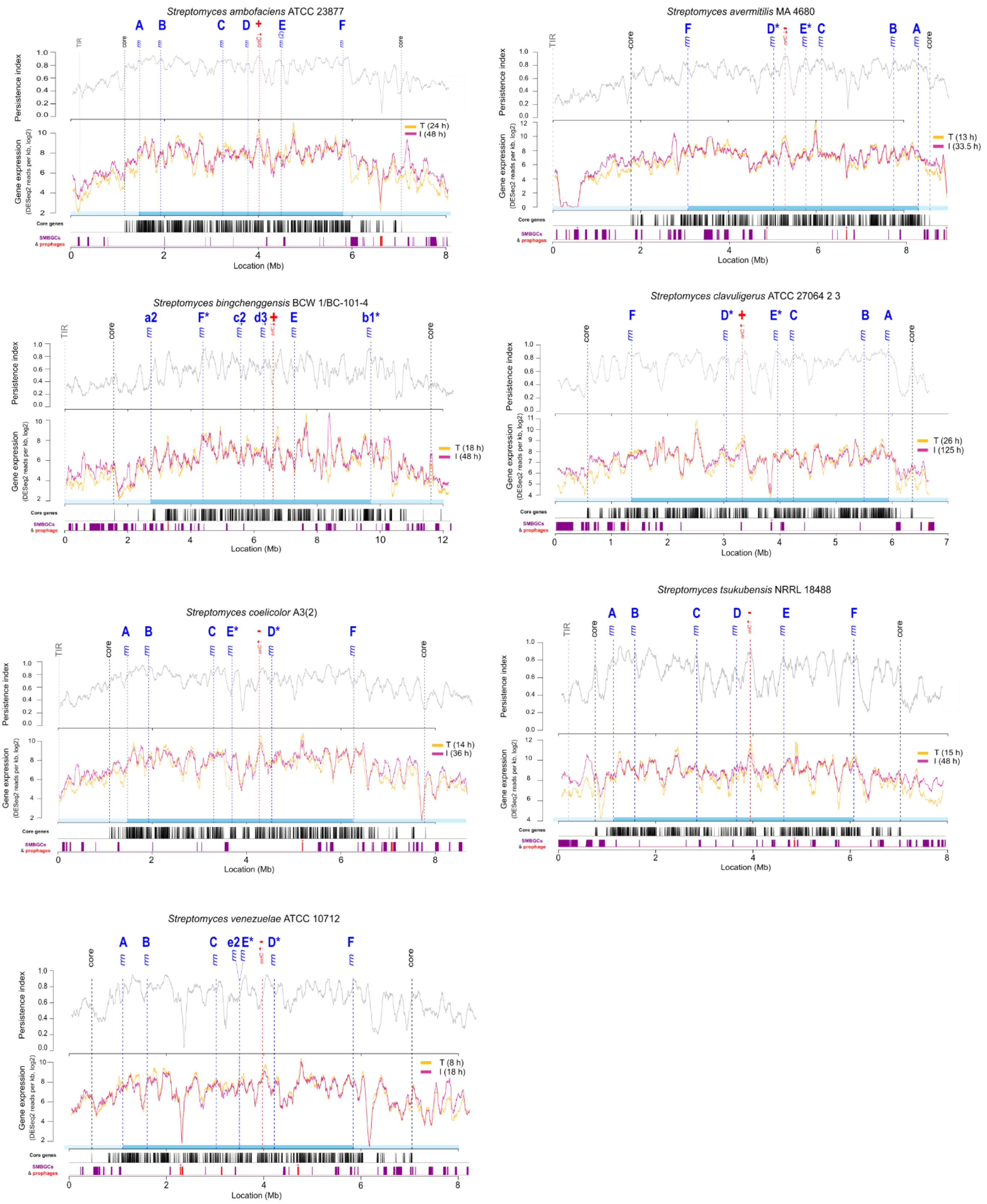


**Supplemental Figure S8: Level of gene persistence and expression along the chromosome of *Streptomyces* species of interest**

The level of gene persistence (*y*-axis, top) and expression (*y*-axis, bottom) along the chromosome (*x*-axis) is represented using a sliding window [81 coding sequences (CDSs), with 1 CDS steps]. Gene expression corresponds to the the DESeq2 normalized counts (log2) measured in cells harvested during the trophophase (T) and the idiophase (I), the time points being indicated for each species (see Methods section for further details). The positions of the most extreme core genes ('core', black lines) and of all *rrn* operon (blue lines) are indicated. The genomic sequences are in the orientation provided by the databases, but the nomenclature used to identify each *rrn* is in reference to an order of the consensus core genome corresponding to '*rrn* ABCDEF *dnaA*+'. The origin of replication (*oriC*) is defined regarding the position of the *dnaA* gene, the red arrow representing the orientation of this gene, and the sign ('+' or '-') its orientation compared to the canonical organization of the chromosome ('*rrn* ABCDEF *dnaA*+'). The density of the core genes as well as the positions of SMBGCs (predicted by AntiSMASH5.0) and prophages (predicted by PHASTER) are indicated under the graph. Incomplete prophages are indicated by an asterisk. Abbreviation: TIR (terminal inverted repeat).
