## Supplementary material for "Ribosomal RNA operons define a central functional compartment in the *Streptomyces* chromosome": Figure S9.pdf

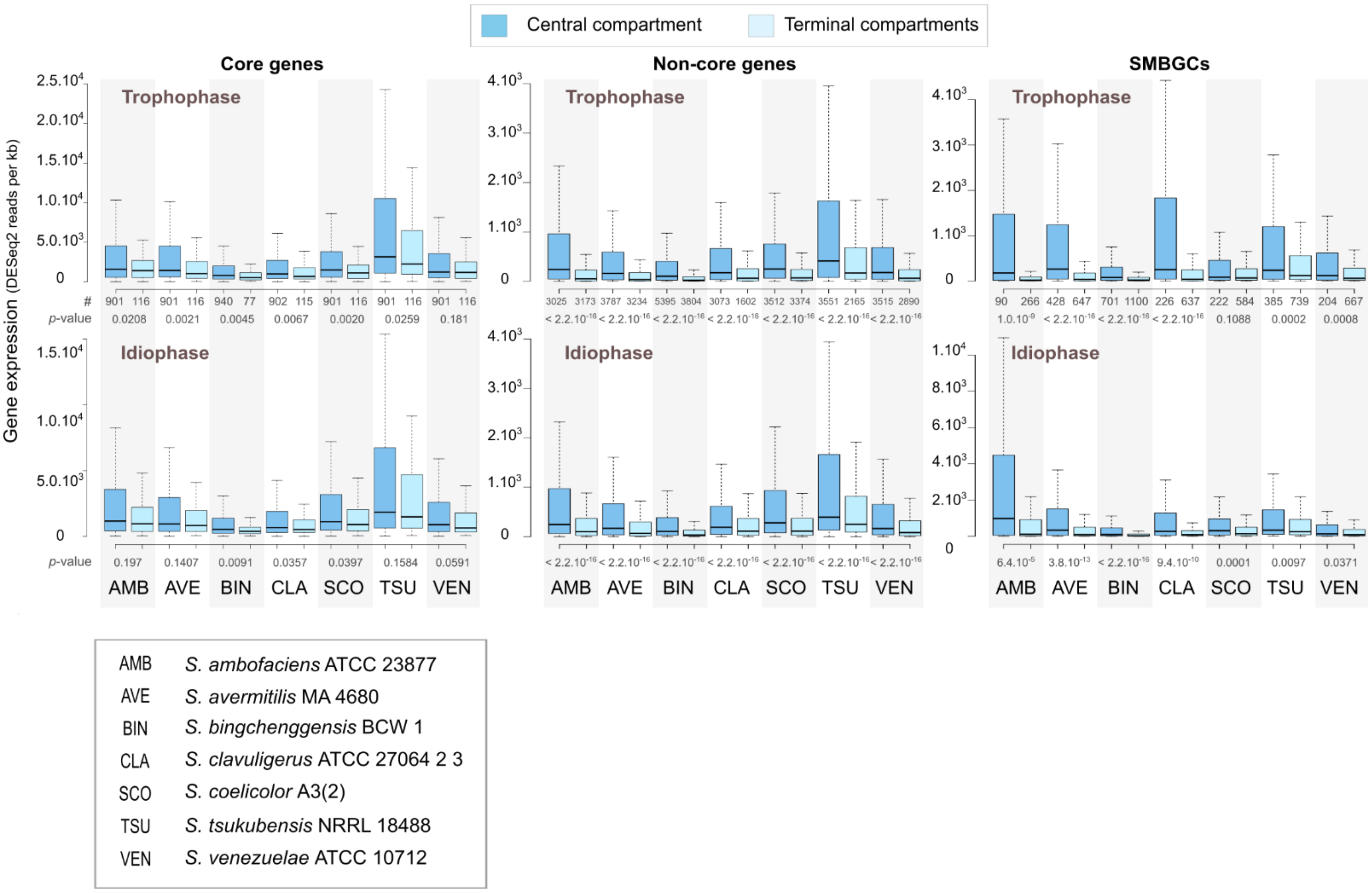


**Supplemental Figure S9: Level of gene expression over growth depending on gene category (core, non-core or SMBGCs) and location inside or outside the central compartment**

The trophophase time points correspond to 24h, 13h, 18h, 26h, 14h, 15h, 8h for *S. ambofaciens* ATCC 23877[1] (AMB), *S. avermitilis* MA 4680[2] (AVE), *S. bingchenggensis* BCW 1/BC-101-4[3] (BIN), *S. clavuligerus* ATCC 27064 2 3[2] (CLA), *S. coelicolor* A3(2) (SCO)[4], *S. tsukubensis* NRRL 18488[2] (TSU) and *S. venezuelae* ATCC 10712[5] (VEN), respectively. The idiophase time points correspond to 48h, 33.5h, 48h, 125h, 36h, 48h, 18h for *S. ambofaciens* ATCC 23877[1] , *S. avermitilis* MA 4680[2], S. *bingchenggensis* BCW 1/BC-101-4[3], *S. clavuligerus* ATCC 27064 2 3[2], *S. coelicolor* A3(2)[4], *S. tsukubensis* NRRL 18488[2] and *S. venezuelae* ATCC 10712[5], respectively. Gene transcription (in sense orientation) corresponds to the number of DESeq2 normalized reads per kb. The boxplot represents the first quartile, median and third quartile. The upper whisker extends from the hinge to the largest value no further than 1.5 * IQR from the hinge. The lower whisker extends from the hinge to the smallest value at most 1.5 * IQR of the hinge.  For clarity, outliers were not represented. Under each bar, the number of genes (#) in each category is indicated as well as the *p*-value of two-sided Wilcoxon rank sum tests with continuity correction (comparing of gene expression in central *versus* terminal compartments).

Transcriptome references:
