## Supplementary material for "Ribosomal RNA operons define a central functional compartment in the *Streptomyces* chromosome": Figure S10.pdf

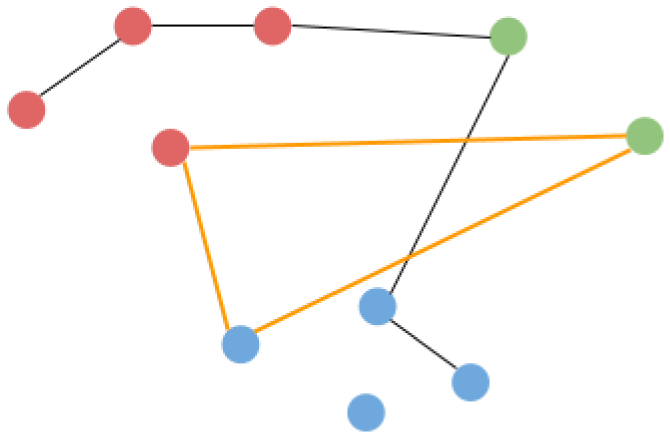

**Supplemental Figure S10: Definition of orthologous groups and core genes**

Each node represents a gene, the color identifies the host species. Edges represent an orthologous relationship. Each set of nodes connected by a path forms an orthologous group/family. The core genome is defined as the set of complete subgraphs (i.e. subsets of fully interconnected nodes, or ‘clique’) with one node in each possible color (example with the orange edges).
