## Supplementary material for "Ribosomal RNA operons define a central functional compartment in the *Streptomyces* chromosome": Supplemental_file_1.pdf

### CODE for rrn-based evolutionary history of Streptomyces

Stéphanie Bury-Moné

20/03/2022

#### Tools and data

Library installations

```
library(readxl)
library(ggplot2)
library("ggpubr")
library("ggcorrplot")
library(corrplot)
library(car)
```

Data import The code below is to be used with the data from the supplemental tables of the article by Lorenzi *et al.*. Remove the sheet dedicated to the legend before importing the tables.

```
ALL <- read_excel("Supplemental_Table_S1.xlsx")
RRNA <- read_excel("Supplemental_Table_S8.xlsx")
CHR<-read_excel("Supplemental_Table_S6.xlsx")
CORE<-subset(CHR, CHR$core=="True")
REARRANG<-read_excel("Supplemental_Table_S5.xlsx")
RNA_seq<-read_excel("Supplemental_Table_S7.xlsx")
SCO_ANNOT<-read_excel("Supplemental_Table_S9.xlsx")
```

#### Figure 1 _ *rrn* operons number and location

Analyzing *rrn* operons number in *Streptomyces* genus

```
ALL$number_rrn_loci<-as.factor(ALL$Number_of_rrn_loci)
ALL$number_rrn_genes<-as.numeric(ALL$Number_of_rrn_genes)
ggplot(ALL, aes(x=number_rrn_genes, fill=number_rrn_loci), binwidth=10) +
  geom_histogram(alpha=0.5) +
  geom_text(aes(label = ..count..), stat = "count", vjust = 0, colour = "blue") +
theme_classic()
```

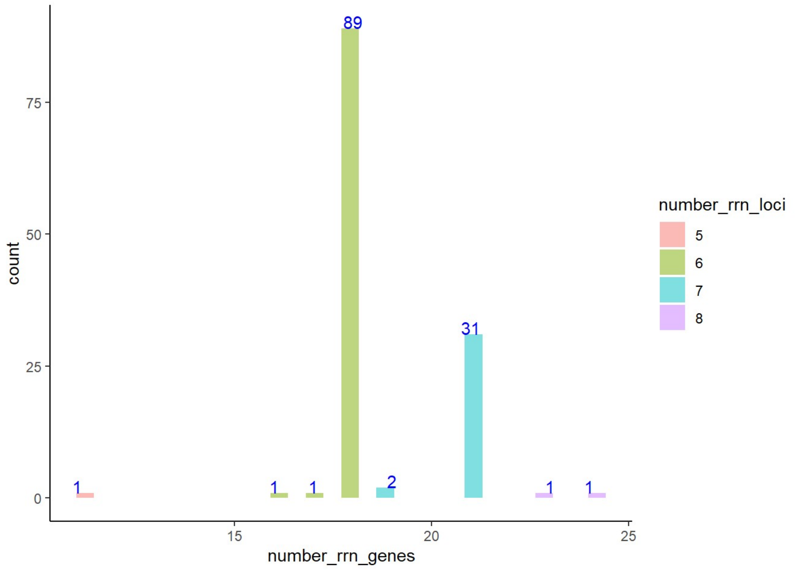


Analyzing *rrn* operons location along the chromosome (defined as the distance to the origin)

```
RRNA$d_ori_bp<-as.numeric(RRNA$d_ori_bp)
RRNA$d_ori_percent_chr_size<-as.numeric(RRNA$d_ori_percent_chr_size)
RRNA$d_ori_percent_central_comp<-as.numeric(RRNA$d_ori_percent_central_comp)
RRNA_nom<-subset(RRNA, RRNA$d_ori_bp!="NA")

ggplot(RRNA_nom, aes(x=d_ori_bp, color=rrn_NOMENCLATURE_simplified, fill = rrn_NOMENCLATURE_simplified)) +    geom_histogram
(position="identity",alpha=0.6, binwidth=20000)+
  labs(y = "Counts", x = "Distance between rrn operons and the oriC in bp")+
  scale_color_manual(values=c( "yellow2", "tan1", "violet", "lightsteelblue1", "pink","springgreen3","navy","firebrick1" ))+
  scale_fill_manual(values=c("yellow2", "tan1", "violet", "lightsteelblue1", "pink","springgreen3","navy","firebrick1" ))+
  theme_classic()
```

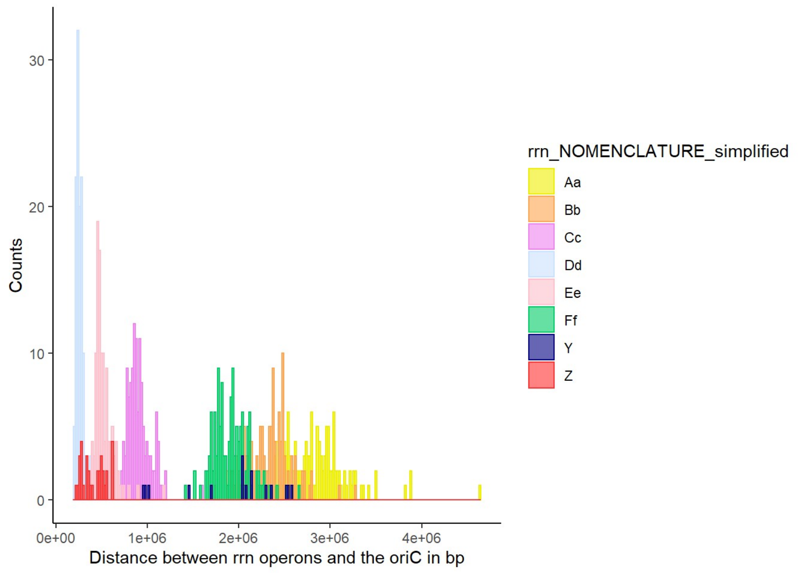


```
ggplot(RRNA_nom, aes(x=d_ori_percent_chr_size, color=rrn_NOMENCLATURE_simplified, fill = rrn_NOMENCLATURE_simplified)) +
geom_histogram(position="identity",alpha=0.6, binwidth=0.2)+
  labs(y = "Counts", x = "Distance between rrn operons and the oriC as the percentage of the chromosome size")+
  scale_color_manual(values=c( "yellow2", "tan1", "violet", "lightsteelblue1", "pink","springgreen3","navy","firebrick1" ))+
  scale_fill_manual(values=c("yellow2", "tan1", "violet", "lightsteelblue1", "pink","springgreen3","navy","firebrick1" ))+
  theme_classic()
```

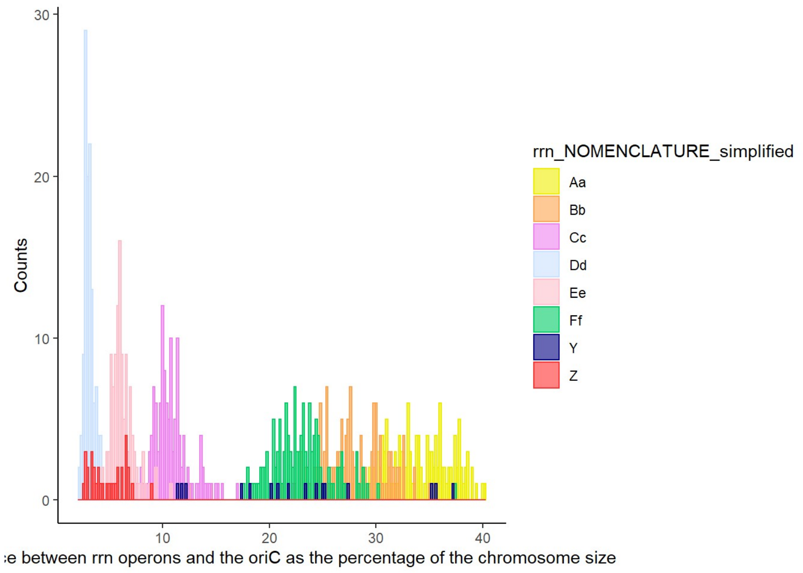


```
ggplot(RRNA_nom, aes(x=d_ori_percent_central_comp, color=rrn_NOMENCLATURE_simplified, fill = rrn_NOMENCLATURE_simplified)) +
geom_histogram(position="identity",alpha=0.6, binwidth=0.2)+
  labs(y = "Counts", x = "Distance between rrn operons and the oriC as the percentage of the central compartment")+
  scale_color_manual(values=c( "yellow2", "tan1", "violet", "lightsteelblue1", "pink","springgreen3","navy","firebrick1" ))+
  scale_fill_manual(values=c("yellow2", "tan1", "violet", "lightsteelblue1", "pink","springgreen3","navy","firebrick1" ))+
  theme_classic()
```

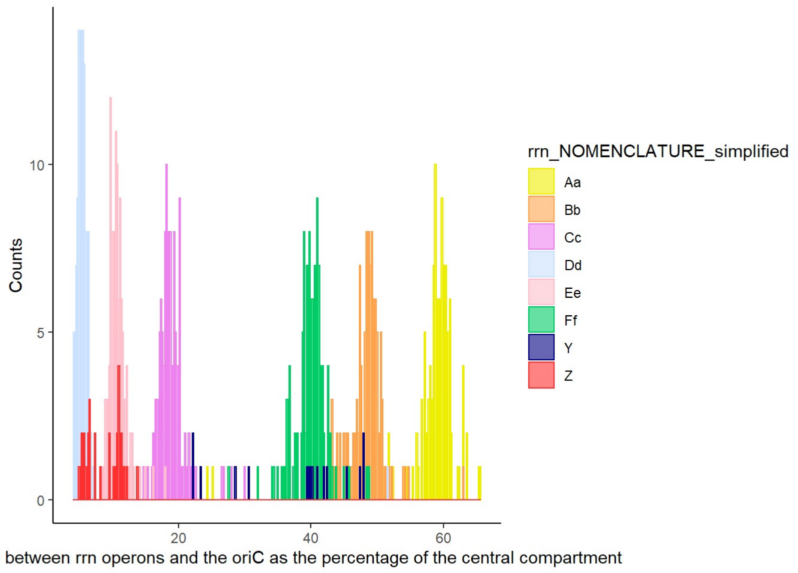


Analyzing the distribution of *rrn* operon environment in *Streptomyces* genus

```
ggplot(data=RRNA_nom, aes(x=rrn_NOMENCLATURE_simplified_2)) +
  geom_bar(stat="Count", width=0.7, fill="steelblue")+
  labs(y = "Number of genomes", x = "rrn environment category")+ coord_flip() +
  theme_classic()
```

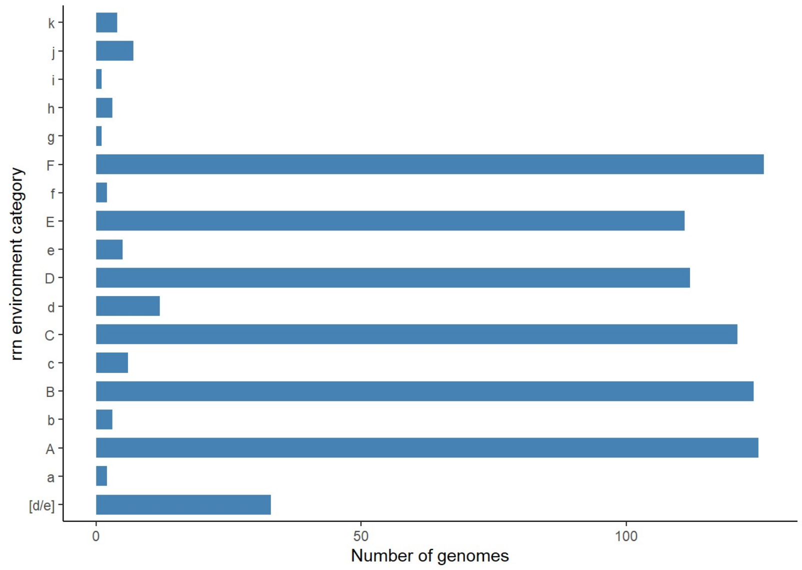


### Ancestral core genome order (Table S3)

```r
#Selection of the strain with the most frequent rrn configuration
ALL_CANON<-subset(ALL, ALL$rrn_configuration=="A B C D E F dnaA+")

#Selection of the core genome of the strains with the most frequent rrn configuration
CORE$CANON<-"NA"
for (i in 1:129145){
  for (j in 1:52){
    if (CORE$genome_id[i]==ALL_CANON$Species[j] ){
      CORE$CANON[i]<-"OK"
    }
  }}
CORE_CANON<-subset(CORE, CORE$CANON=="OK")

#Orientation of the genomes depending on the dnaA orientation released from databases
CORE_CANON$ORIENTATION<-"NA"
for (i in 1:52872){
  for (j in 1:52){
    if (CORE_CANON$genome_id[i]==ALL_CANON$Species[j] ){
      CORE_CANON$ORIENTATION[i]<-ALL_CANON$dnaA_orientation_in_the_available_sequence_genome[j]
    }
  }}
CORE_CANON_SENSE_wo_fungi<-subset(CORE_CANON, CORE_CANON$ORIENTATION=="+" & CORE_CANON$genome_id!="Streptomyces fungicidicus 
TXX3120")
CORE_CANON_SENSE_fungi<-subset(CORE_CANON, CORE_CANON$genome_id=="Streptomyces fungicidicus TXX3120")
CORE_CANON_ANTISENSE<-subset(CORE_CANON, CORE_CANON$ORIENTATION=="-")

#Numbering of genes according to their order in the core genome - the case of Streptomyces fungicidicus TXX3120 is treated s
eparately because there are only 1005 genes on the chromosome (the rest being carried by a plasmid)
CORE_CANON_SENSE_wo_fungi$order<-rep(1:1017, 31)
CORE_CANON_SENSE_fungi$order<-c(1:1005)
CORE_CANON_ANTISENSE$order<-rep(1017:1, 20)
CORE_CANON_ORDER<-rbind(CORE_CANON_SENSE_wo_fungi, CORE_CANON_SENSE_fungi, CORE_CANON_ANTISENSE)

#Counting the possible ranks for each core gene
mytable<-table(CORE_CANON_ORDER$order, CORE_CANON_ORDER$ortholog_group)
tab<-as.data.frame.matrix(mytable)

#Generation of a prototypical subset of core genes to indicate the ancestral order
Order <- c(1:1017)
MAX <- rep("NA", 1017)
Ortholog_group <- rep("NA", 1017)
Ancestral_core_order <- data.frame(Order,MAX, Ortholog_group)

for (i in 1:1017){
  for(j in 1:1017) {
    # checking if the data frame
    # cell value is equal to the maximum
    if(tab[i,j]==Ancestral_core_order$MAX[i]){
      # replacing the second line
      # with the row name which is the rank...
      Ancestral_core_order$Ortholog_group[i]<-colnames(tab[j])}
  }
}
tab<-table(Ancestral_core_order$Ortholog_group)
write.csv2(tab, "tab.csv")
#We notice that one ortholog_group (3790) is present twice, which means that one ortholog_group is missing and 498-499 ranks 
are ambiguous.
#Identification of the missing ortholog_group
VIRO_CORE<-subset(CORE_CANON, CORE_CANON$genome_id=="Streptomyces viridosporus T7A ATCC 39115")
VIRO_CORE$present<-"NA"
for (i in 1:1017){
  for(j in 1:1017) {
    if(VIRO_CORE$ortholog_group[i]==Ancestral_core_order$Ortholog_group[j]){
      VIRO_CORE$present[i]<-"OK"}
  }
}
Absent<-subset(VIRO_CORE, VIRO_CORE$present=="NA")
#The ortholog_group "4240" is missing. Manual analysis reveals that ranks 498 and 499 are ambiguous.
#Analysis of ortholog_group order around "4240" ortholog_group.

ALL_CANON$CDS4240_minus2<-"NA"
ALL_CANON$CDS4240_minus1<-"NA"
ALL_CANON$CDS4240_plus1<-"NA"
ALL_CANON$CDS4240_plus2<-"NA"

for (i in 1:52){
  for (j in 1:31527){
    if (ALL_CANON$Species[i]==CORE_CANON_SENSE_wo_fungi$genome_id[j] & CORE_CANON_SENSE_wo_fungi$ortholog_group[j]=="4240"){
      ALL_CANON$CDS4240_minus2[i]<-CORE_CANON_SENSE_wo_fungi$ortholog_group[j-2]
      ALL_CANON$CDS4240_minus1[i]<-CORE_CANON_SENSE_wo_fungi$ortholog_group[j-1]
      ALL_CANON$CDS4240_plus1[i]<-CORE_CANON_SENSE_wo_fungi$ortholog_group[j+1]
      ALL_CANON$CDS4240_plus2[i]<-CORE_CANON_SENSE_wo_fungi$ortholog_group[j+2]
    }
  }}

for (i in 1:52){
  for (j in 1:20340){
    if (ALL_CANON$Species[i]==CORE_CANON_ANTISENSE$genome_id[j] & CORE_CANON_ANTISENSE$ortholog_group[j]=="4240"){
      ALL_CANON$CDS4240_minus2[i]<-CORE_CANON_ANTISENSE$ortholog_group[j+2]
      ALL_CANON$CDS4240_minus1[i]<-CORE_CANON_ANTISENSE$ortholog_group[j+1]
      ALL_CANON$CDS4240_plus1[i]<-CORE_CANON_ANTISENSE$ortholog_group[j-1]
```

```
      ALL_CANON$CDS4240_plus2[i]<-CORE_CANON_ANTISENSE$ortholog_group[j-2]
    }
  }}


for (i in 1:52){
  for (j in 1:1005){
    if (ALL_CANON$Species[i]==CORE_CANON_SENSE_fungi$genome_id[j] & CORE_CANON_SENSE_fungi$ortholog_group[j]=="4240"){
      ALL_CANON$CDS4240_minus2[i]<-CORE_CANON_SENSE_fungi$ortholog_group[j-2]
      ALL_CANON$CDS4240_minus1[i]<-CORE_CANON_SENSE_fungi$ortholog_group[j-1]
      ALL_CANON$CDS4240_plus1[i]<-CORE_CANON_SENSE_fungi$ortholog_group[j+1]
      ALL_CANON$CDS4240_plus2[i]<-CORE_CANON_SENSE_fungi$ortholog_group[j+2]
    }
  }}
table(ALL_CANON$CDS4240_minus2)
table(ALL_CANON$CDS4240_minus1)
table(ALL_CANON$CDS4240_plus1)
table(ALL_CANON$CDS4240_plus2)

#The most frequent order in this region is so : "3032", "3790", "4240", "7749", "6541".
#The rank n°499 is therefore attributed to "4240".
Ancestral_core_order$Ortholog_group[499]<-"4240"
write.csv2(Ancestral_core_order, "Ancestral_core_order.csv")

#Annotation of all strains to identify which strains are close to the consensus regarding core gene order.

CORE_CANON_SENSE_wo_fungi$ANCESTRAL_RANK<-"NA"
for (i in 1:31527){
  for (j in 1:1017){
    if (CORE_CANON_SENSE_wo_fungi$ortholog_group[i]==Ancestral_core_order$Ortholog_group[j]){
      CORE_CANON_SENSE_wo_fungi$ANCESTRAL_RANK[i]<-Ancestral_core_order$Order[j]
    }
  }}

CORE_CANON_ANTISENSE$ANCESTRAL_RANK<-"NA"
for (i in 1:20340){
  for (j in 1:1017){
    if (CORE_CANON_ANTISENSE$ortholog_group[i]==Ancestral_core_order$Ortholog_group[j]){
      CORE_CANON_ANTISENSE$ANCESTRAL_RANK[i]<-Ancestral_core_order$Order[j]
    }
  }}

CORE_CANON_SENSE_fungi$ANCESTRAL_RANK<-"NA"
for (i in 1:1005){
  for (j in 1:1017){
    if (CORE_CANON_SENSE_fungi$ortholog_group[i]==Ancestral_core_order$Ortholog_group[j]){
      CORE_CANON_SENSE_fungi$ANCESTRAL_RANK[i]<-Ancestral_core_order$Order[j]
    }
  }}

CORE_CANON_ORDER<-rbind(CORE_CANON_SENSE_wo_fungi, CORE_CANON_SENSE_fungi, CORE_CANON_ANTISENSE)
write.csv2(CORE_CANON_ORDER, "CORE_CANON_ORDER.csv")

CORE_CANON_ORDER$order<-as.numeric(CORE_CANON_ORDER$order)
CORE_CANON_ORDER$ANCESTRAL_RANK<-as.numeric(CORE_CANON_ORDER$ANCESTRAL_RANK)
CORE_CANON_ORDER$DIFF<-CORE_CANON_ORDER$order-CORE_CANON_ORDER$ANCESTRAL_RANK

tab<-table(CORE_CANON_ORDER$genome_id,CORE_CANON_ORDER$DIFF)
tab<-as.data.frame.matrix(tab)
strains_as_the_consensus<-subset(tab, tab$`0`=="1017")
row.names(strains_as_the_consensus)
strains_close_to_the_consensus<-subset(tab, tab$`0`=="1015")
row.names(strains_close_to_the_consensus)
```

### Large rearrangements in the central compartment and *rrn* loci (Figure 3, Supplementary Figures 2 and 3)

Plots of pairwise comparisons of the core genomes to that of *Streptomyces viridosporus* T7A ATCC 39115, used as a reference

```
VIRO<-subset(CORE, CORE$genome_id=="Streptomyces viridosporus T7A ATCC 39115")

CORE$start_VIRO<-"NA"
for (i in 1:129145){
  for (j in 1:1017){
    if (CORE$ortholog_group[i]==VIRO$ortholog_group[j]){
      CORE$start_VIRO[i]<-VIRO$start[j]
    }
  }}
write.csv2(CORE, "CORE.csv")

ALL$rrn1<-as.numeric(ALL$rrn1)
ALL$rrn2<-as.numeric(ALL$rrn2)
ALL$rrn3<-as.numeric(ALL$rrn3)
ALL$rrn4<-as.numeric(ALL$rrn4)
ALL$rrn5<-as.numeric(ALL$rrn5)
ALL$rrn6<-as.numeric(ALL$rrn6)
ALL$rrn7<-as.numeric(ALL$rrn7)
ALL$rrn8<-as.numeric(ALL$rrn8)
ALL$dnaA_start_position<-as.numeric(ALL$dnaA_start_position)
CORE$start<-as.numeric(CORE$start)
CORE$start_VIRO<-as.numeric(CORE$start_VIRO)
CORE$start_bis<-CORE$start/1000000
CORE$start_VIRO_bis<-CORE$start_VIRO/1000000

pdf("FigSupp2.pdf",width=8, height=10)
par(mfrow=c(3,3))
for (i in 1:127){
  if (ALL$Number_of_rrn_loci[i]=="6"){
    #For species with 6 rrn loci
    SPECIES<-subset(CORE, CORE$genome_id==ALL$Species[i])
    a<-ALL$rrn1[i]/1000000
    b<-ALL$rrn2[i]/1000000
    c<-ALL$rrn3[i]/1000000
    d<-ALL$rrn4[i]/1000000
    e<-ALL$rrn5[i]/1000000
    f<-ALL$rrn6[i]/1000000
    g<-ALL$dnaA_start_position[i]/1000000
    plot(SPECIES$start_bis, SPECIES$start_VIRO_bis, font.lab = 3, font.sub=3, pch = 20, cex = 0.5, ylab = "S. viridosporus
(Mb)", xlab=paste("S. ",ALL$id[i]," (Mb)","\n",ALL$rrn_configuration[i]), sub=paste(ALL$Number_of_rrn_loci[i]," rrn loci, ",
ALL$Clade[i]))
    #SPECIES
    abline(v=c(a, b, c, d, e, f, g), col=c("blue","blue","blue","blue","blue","blue","red"), lty=2, lwd=1)
    #VIRO
    abline(h=c(1672357/1000000, 2106072/1000000, 3517690/1000000, 4082352/1000000, 4740474/1000000, 5958129/1000000, 4286628
/1000000), col=c("blue","blue","blue","blue","blue","blue","red"), lty=2, lwd=1)
  }else if (ALL$Number_of_rrn_loci[i]=="5"){
    #For species with 5 rrn loci
    SPECIES<-subset(CORE, CORE$genome_id==ALL$Species[i])
    a<-ALL$rrn1[i]/1000000
    b<-ALL$rrn2[i]/1000000
    c<-ALL$rrn3[i]/1000000
    d<-ALL$rrn4[i]/1000000
    e<-ALL$rrn5[i]/1000000
    g<-ALL$dnaA_start_position[i]/1000000
    plot(SPECIES$start_bis, SPECIES$start_VIRO_bis, font.lab = 3, font.sub=3, pch = 20, cex = 0.5, ylab = "S. viridosporus
(Mb)", xlab=paste("S. ",ALL$id[i]," (Mb)","\n",ALL$rrn_configuration[i]), sub=paste(ALL$Number_of_rrn_loci[i]," rrn loci, ",
ALL$Clade[i]))
    #SPECIES
    abline(v=c(a, b, c, d, e, g), col=c("blue","blue","blue","blue","blue","red"), lty=2, lwd=1)
    #VIRO
    abline(h=c(1672357/1000000, 2106072/1000000, 3517690/1000000, 4082352/1000000, 4740474/1000000, 5958129/1000000, 4286628
/1000000), col=c("blue","blue","blue","blue","blue","blue","red"), lty=2, lwd=1)
  }else if (ALL$Number_of_rrn_loci[i]=="7"){
    #For species with 7 rrn loci
    SPECIES<-subset(CORE, CORE$genome_id==ALL$Species[i])
    a<-ALL$rrn1[i]/1000000
    b<-ALL$rrn2[i]/1000000
    c<-ALL$rrn3[i]/1000000
    d<-ALL$rrn4[i]/1000000
    e<-ALL$rrn5[i]/1000000
    f<-ALL$rrn6[i]/1000000
    k<-ALL$rrn7[i]/1000000
    g<-ALL$dnaA_start_position[i]/1000000
    plot(SPECIES$start_bis, SPECIES$start_VIRO_bis, font.lab = 3, font.sub=3, pch = 20, cex = 0.5, ylab = "S. viridosporus
(Mb)", xlab=paste("S. ",ALL$id[i]," (bp)","\n",ALL$rrn_configuration[i]), sub=paste(ALL$Number_of_rrn_loci[i]," rrn loci, ",
ALL$Clade[i]))
    #SPECIES
    abline(v=c(a, b, c, d, e, f, k, g), col=c("blue","blue","blue","blue","blue","blue","blue","red"), lty=2, lwd=1)
    #VIRO
    abline(h=c(1672357/1000000, 2106072/1000000, 3517690/1000000, 4082352/1000000, 4740474/1000000, 5958129/1000000, 4286628
/1000000), col=c("blue","blue","blue","blue","blue","blue","red"), lty=2, lwd=1)
  }else if (ALL$Number_of_rrn_loci[i]=="8"){
    #For species with 8 rrn loci
    SPECIES<-subset(CORE, CORE$genome_id==ALL$Species[i])
    a<-ALL$rrn1[i]/1000000
    b<-ALL$rrn2[i]/1000000
    c<-ALL$rrn3[i]/1000000
    d<-ALL$rrn4[i]/1000000
    e<-ALL$rrn5[i]/1000000
    f<-ALL$rrn6[i]/1000000
```

```
    k<-ALL$rrn7[i]/1000000
    l<-ALL$rrn8[i]/1000000
    g<-ALL$dnaA_start_position[i]/1000000
    plot(SPECIES$start_bis, SPECIES$start_VIRO_bis, font.lab = 3, font.sub=3, pch = 20, cex = 0.5, ylab = "S. viridosporus (Mb)", xlab=paste("S. ",ALL$id[i]," (bp)","\n",ALL$rrn_configuration[i]), sub=paste(ALL$Number_of_rrn_loci[i]," rrn loci, ", ALL$Clade[i]))
    #SPECIES
    abline(v=c(a, b, c, d, e, f, k,l, g), col=c("blue","blue","blue","blue","blue","blue","blue","blue","red"), lty=2, lwd=1)
    #VIRO
    abline(h=c(1672357/1000000, 2106072/1000000, 3517690/1000000, 4082352/1000000, 4740474/1000000, 5958129/1000000, 4286628/1000000), col=c("blue","blue","blue","blue","blue","blue","red"), lty=2, lwd=1)
  }
}


dev.off()
```

Supplementary Figure 3 was constructed following the same logic as the above code, but changing the reference strain.

Analysis of the origin island described by Algora-Gallardo *et al.* (2021)

```
CORE$ORI_ISLAND_SCO_RANK<-"NA"
ORI_ISLAND<-subset(SCO_ANNOT, SCO_ANNOT$ORI_ISLAND_RANK!="NA")

for (i in 1:129145){
  for (j in 1:22){
    if (CORE$ortholog_group[i]==ORI_ISLAND$ortholog_group[j]){
      CORE$ORI_ISLAND_SCO_RANK[i]<-ORI_ISLAND$ORI_ISLAND_RANK[j]
    }
  }}

ORI_ISLAND_CORE<-subset(CORE, CORE$ORI_ISLAND_SCO_RANK!="NA")
ORI_ISLAND_CORE$ORI_ISLAND_SCO_RANK<-as.numeric(ORI_ISLAND_CORE$ORI_ISLAND_SCO_RANK)
ORI_ISLAND_CORE$ORI_ISLAND_RANK_verif<-"NA"
for (i in 1:2793){
    j = i + 1
    a <- ORI_ISLAND_CORE$ORI_ISLAND_SCO_RANK[j]-1
    b <- ORI_ISLAND_CORE$ORI_ISLAND_SCO_RANK[j]+1
    if (ORI_ISLAND_CORE$ORI_ISLAND_SCO_RANK[i]==a){
      ORI_ISLAND_CORE$ORI_ISLAND_RANK_verif[i]<-"ascending order"
    }else if (ORI_ISLAND_CORE$ORI_ISLAND_SCO_RANK[i]==b){
      ORI_ISLAND_CORE$ORI_ISLAND_RANK_verif[i]<-"descending order"
    }}
table(ORI_ISLAND_CORE$ORI_ISLAND_RANK_verif)
```

```
##
##  ascending order descending order               NA
##             1407             1260              127
```

Pairwise comparisons of the core genomes to that of *Streptomyces viridosporus* T7A ATCC 39115, used as a reference, to identify large rearrangements

```
CORE$start<-as.numeric(CORE$start)
CORE$VIRO_start<-as.numeric(CORE$start_VIRO)

#Calculation of the difference in positions of the same core gene between the two strains
CORE$delta_VIRO<-CORE$start-CORE$start_VIRO

#Calculation of the difference in this difference between two successive positions in the genome of the strain of interest -
The purpose of this calculation is to identify an increase or decrease in this difference that would indicate a rearrangemen
t in the genome of the strain compared to the reference

CORE$delta_VIRO_delta<-"NA"
for (i in 2:129145){
  j = i - 1
  if (CORE$genome_id[i]==CORE$genome_id[j]){
    CORE$delta_VIRO_delta[i]<-CORE$delta_VIRO[i]-CORE$delta_VIRO[j]
  }
}
CORE$delta_VIRO_delta<-as.numeric(CORE$delta_VIRO_delta)
write.csv2(CORE, "CORE_delta_VIRO_delta.csv", row.names = F)

#Selection of large rearrangements (>200 kb)
RERRANG_vs_VIRO<-subset(CORE, abs(CORE$delta_VIRO_delta)>200000 & CORE$compartment=="CENTRAL_COMP")

#Filter to eliminate small region "jumps"
RERRANG_vs_VIRO$delta_CDS<-"NA"
for (i in 1:464){
  j = i+1
    if (RERRANG_vs_VIRO$genome_id[i]==RERRANG_vs_VIRO$genome_id[j]){
      RERRANG_vs_VIRO$delta_CDS[i]<-RERRANG_vs_VIRO$start[j]- RERRANG_vs_VIRO$start[i]
    }}

RERRANG_vs_VIRO_2<-subset(RERRANG_vs_VIRO, RERRANG_vs_VIRO$delta_CDS=="NA")
RERRANG_vs_VIRO$delta_CDS<-as.numeric(RERRANG_vs_VIRO$delta_CDS)
RERRANG_vs_VIRO_3<-subset(RERRANG_vs_VIRO, RERRANG_vs_VIRO$delta_CDS>20000)
RERRANG_vs_VIRO_4<-rbind(RERRANG_vs_VIRO_2,RERRANG_vs_VIRO_3)

#To identify later the lines corresponding to the ends of the same rearrangement
RERRANG_vs_VIRO_4$delta_VIRO_delta_ABS<-abs(RERRANG_vs_VIRO_4$delta_VIRO_delta)

#Annotation of each event

RERRANG_vs_VIRO_4$rrn1_start<-"NA"
RERRANG_vs_VIRO_4$rrn2_start<-"NA"
RERRANG_vs_VIRO_4$rrn3_start<-"NA"
RERRANG_vs_VIRO_4$rrn4_start<-"NA"
RERRANG_vs_VIRO_4$rrn5_start<-"NA"
RERRANG_vs_VIRO_4$rrn6_start<-"NA"
RERRANG_vs_VIRO_4$rrn7_start<-"NA"
RERRANG_vs_VIRO_4$rrn8_start<-"NA"

RERRANG_vs_VIRO_4$rrn1_category<-"NA"
RERRANG_vs_VIRO_4$rrn2_category<-"NA"
RERRANG_vs_VIRO_4$rrn3_category<-"NA"
RERRANG_vs_VIRO_4$rrn4_category<-"NA"
RERRANG_vs_VIRO_4$rrn5_category<-"NA"
RERRANG_vs_VIRO_4$rrn6_category<-"NA"
RERRANG_vs_VIRO_4$rrn7_category<-"NA"
RERRANG_vs_VIRO_4$rrn8_category<-"NA"

for (i in 1:400){
  for (j in 1:1016){
    if (RERRANG_vs_VIRO_4$genome_id[i]==RRNA$genome_id[j] & RRNA$rrn_order[j]=="rrn_1"){
      RERRANG_vs_VIRO_4$rrn1_start[i]<-RRNA$rrn_position[j]
      RERRANG_vs_VIRO_4$rrn1_category[i]<-RRNA$rrn_NOMENCLATURE_simplified[j]
    }
  }}

for (i in 1:400){
  for (j in 1:1016){
    if (RERRANG_vs_VIRO_4$genome_id[i]==RRNA$genome_id[j] & RRNA$rrn_order[j]=="rrn_2"){
      RERRANG_vs_VIRO_4$rrn2_start[i]<-RRNA$rrn_position[j]
      RERRANG_vs_VIRO_4$rrn2_category[i]<-RRNA$rrn_NOMENCLATURE_simplified[j]
    }
  }}

for (i in 1:400){
  for (j in 1:1016){
    if (RERRANG_vs_VIRO_4$genome_id[i]==RRNA$genome_id[j] & RRNA$rrn_order[j]=="rrn_3"){
      RERRANG_vs_VIRO_4$rrn3_start[i]<-RRNA$rrn_position[j]
      RERRANG_vs_VIRO_4$rrn3_category[i]<-RRNA$rrn_NOMENCLATURE_simplified[j]
    }
  }}

for (i in 1:400){
  for (j in 1:1016){
    if (RERRANG_vs_VIRO_4$genome_id[i]==RRNA$genome_id[j] & RRNA$rrn_order[j]=="rrn_4"){
      RERRANG_vs_VIRO_4$rrn4_start[i]<-RRNA$rrn_position[j]
      RERRANG_vs_VIRO_4$rrn4_category[i]<-RRNA$rrn_NOMENCLATURE_simplified[j]
    }
  }}
```

```
for (i in 1:400){
  for (j in 1:1016){
    if (RERRANG_vs_VIRO_4$genome_id[i]==RRNA$genome_id[j] & RRNA$rrn_order[j]=="rrn_5"){
      RERRANG_vs_VIRO_4$rrn5_start[i]<-RRNA$rrn_position[j]
      RERRANG_vs_VIRO_4$rrn5_category[i]<-RRNA$rrn_NOMENCLATURE_simplified[j]
    }
  }}

for (i in 1:400){
  for (j in 1:1016){
    if (RERRANG_vs_VIRO_4$genome_id[i]==RRNA$genome_id[j] & RRNA$rrn_order[j]=="rrn_6"){
      RERRANG_vs_VIRO_4$rrn6_start[i]<-RRNA$rrn_position[j]
      RERRANG_vs_VIRO_4$rrn6_category[i]<-RRNA$rrn_NOMENCLATURE_simplified[j]
    }
  }}

for (i in 1:400){
  for (j in 1:1016){
    if (RERRANG_vs_VIRO_4$genome_id[i]==RRNA$genome_id[j] & RRNA$rrn_order[j]=="rrn_7"){
      RERRANG_vs_VIRO_4$rrn7_start[i]<-RRNA$rrn_position[j]
      RERRANG_vs_VIRO_4$rrn7_category[i]<-RRNA$rrn_NOMENCLATURE_simplified[j]
    }
  }}

for (i in 1:400){
  for (j in 1:1016){
    if (RERRANG_vs_VIRO_4$genome_id[i]==RRNA$genome_id[j] & RRNA$rrn_order[j]=="rrn_8"){
      RERRANG_vs_VIRO_4$rrn8_start[i]<-RRNA$rrn_position[j]
      RERRANG_vs_VIRO_4$rrn8_category[i]<-RRNA$rrn_NOMENCLATURE_simplified[j]
    }
  }}
RERRANG_vs_VIRO_4$rrn1_start<-as.numeric(RERRANG_vs_VIRO_4$rrn1_start)
RERRANG_vs_VIRO_4$rrn2_start<-as.numeric(RERRANG_vs_VIRO_4$rrn2_start)
RERRANG_vs_VIRO_4$rrn3_start<-as.numeric(RERRANG_vs_VIRO_4$rrn3_start)
RERRANG_vs_VIRO_4$rrn4_start<-as.numeric(RERRANG_vs_VIRO_4$rrn4_start)
RERRANG_vs_VIRO_4$rrn5_start<-as.numeric(RERRANG_vs_VIRO_4$rrn5_start)
RERRANG_vs_VIRO_4$rrn6_start<-as.numeric(RERRANG_vs_VIRO_4$rrn6_start)
RERRANG_vs_VIRO_4$rrn7_start<-as.numeric(RERRANG_vs_VIRO_4$rrn7_start)
RERRANG_vs_VIRO_4$rrn8_start<-as.numeric(RERRANG_vs_VIRO_4$rrn8_start)

RERRANG_vs_VIRO_4$rrn1_distance<-abs(RERRANG_vs_VIRO_4$start-RERRANG_vs_VIRO_4$rrn1_start)
RERRANG_vs_VIRO_4$rrn2_distance<-abs(RERRANG_vs_VIRO_4$start-RERRANG_vs_VIRO_4$rrn2_start)
RERRANG_vs_VIRO_4$rrn3_distance<-abs(RERRANG_vs_VIRO_4$start-RERRANG_vs_VIRO_4$rrn3_start)
RERRANG_vs_VIRO_4$rrn4_distance<-abs(RERRANG_vs_VIRO_4$start-RERRANG_vs_VIRO_4$rrn4_start)
RERRANG_vs_VIRO_4$rrn5_distance<-abs(RERRANG_vs_VIRO_4$start-RERRANG_vs_VIRO_4$rrn5_start)
RERRANG_vs_VIRO_4$rrn6_distance<-abs(RERRANG_vs_VIRO_4$start-RERRANG_vs_VIRO_4$rrn6_start)
RERRANG_vs_VIRO_4$rrn7_distance<-abs(RERRANG_vs_VIRO_4$start-RERRANG_vs_VIRO_4$rrn7_start)
RERRANG_vs_VIRO_4$rrn8_distance<-abs(RERRANG_vs_VIRO_4$start-RERRANG_vs_VIRO_4$rrn8_start)

for (i in 1:400){
  RERRANG_vs_VIRO_4$rrn_min_distance[i]<-min(RERRANG_vs_VIRO_4$rrn1_distance[i], RERRANG_vs_VIRO_4$rrn2_distance[i], RERRANG
_vs_VIRO_4$rrn3_distance[i], RERRANG_vs_VIRO_4$rrn4_distance[i], RERRANG_vs_VIRO_4$rrn5_distance[i], RERRANG_vs_VIRO_4$rrn6_
distance[i], RERRANG_vs_VIRO_4$rrn7_distance[i], RERRANG_vs_VIRO_4$rrn8_distance[i], na.rm = TRUE)
  }
RERRANG_vs_VIRO_4$rrn_min_distance<-as.numeric(RERRANG_vs_VIRO_4$rrn_min_distance)

RERRANG_vs_VIRO_4$Number_of_rrn_loci<-"NA"
for (i in 1:400){
  for (j in 1:127){
    if (RERRANG_vs_VIRO_4$genome_id[i]==ALL$Species[j]){
      RERRANG_vs_VIRO_4$Number_of_rrn_loci[i]<-ALL$Number_of_rrn_loci[j]
    }
  }}

RERRANG_vs_VIRO_4$rrn_min_category<-"NA"
for (i in 1:400){
  if (RERRANG_vs_VIRO_4$rrn_min_distance[i]==RERRANG_vs_VIRO_4$rrn1_distance[i]){
  RERRANG_vs_VIRO_4$rrn_min_category[i]<-RERRANG_vs_VIRO_4$rrn1_category[i]
  }
}

for (i in 1:400){
  if (RERRANG_vs_VIRO_4$rrn_min_distance[i]==RERRANG_vs_VIRO_4$rrn2_distance[i]){
  RERRANG_vs_VIRO_4$rrn_min_category[i]<-RERRANG_vs_VIRO_4$rrn2_category[i]
  }
}

for (i in 1:400){
  if (RERRANG_vs_VIRO_4$rrn_min_distance[i]==RERRANG_vs_VIRO_4$rrn3_distance[i]){
  RERRANG_vs_VIRO_4$rrn_min_category[i]<-RERRANG_vs_VIRO_4$rrn3_category[i]
  }
}

for (i in 1:400){
  if (RERRANG_vs_VIRO_4$rrn_min_distance[i]==RERRANG_vs_VIRO_4$rrn4_distance[i]){
  RERRANG_vs_VIRO_4$rrn_min_category[i]<-RERRANG_vs_VIRO_4$rrn4_category[i]
  }
}

for (i in 1:400){
  if (RERRANG_vs_VIRO_4$rrn_min_distance[i]==RERRANG_vs_VIRO_4$rrn5_distance[i]){
  RERRANG_vs_VIRO_4$rrn_min_category[i]<-RERRANG_vs_VIRO_4$rrn5_category[i]
```

```
  }
}

for (i in 1:400){
  if (RERRANG_vs_VIRO_4$Number_of_rrn_loci[i]=="6" & RERRANG_vs_VIRO_4$rrn_min_distance[i]==RERRANG_vs_VIRO_4$rrn6_distance
[i]){
  RERRANG_vs_VIRO_4$rrn_min_category[i]<-RERRANG_vs_VIRO_4$rrn6_category[i]
  }
}

for (i in 1:400){
  if (RERRANG_vs_VIRO_4$Number_of_rrn_loci[i]=="7" & RERRANG_vs_VIRO_4$rrn_min_distance[i]==RERRANG_vs_VIRO_4$rrn6_distance
[i]){
  RERRANG_vs_VIRO_4$rrn_min_category[i]<-RERRANG_vs_VIRO_4$rrn6_category[i]
  }
}

for (i in 1:400){
  if (RERRANG_vs_VIRO_4$Number_of_rrn_loci[i]=="7" & RERRANG_vs_VIRO_4$rrn_min_distance[i]==RERRANG_vs_VIRO_4$rrn7_distance
[i]){
  RERRANG_vs_VIRO_4$rrn_min_category[i]<-RERRANG_vs_VIRO_4$rrn7_category[i]
  }
}

for (i in 1:400){
  if (RERRANG_vs_VIRO_4$Number_of_rrn_loci[i]=="8" & RERRANG_vs_VIRO_4$rrn_min_distance[i]==RERRANG_vs_VIRO_4$rrn6_distance
[i]){
  RERRANG_vs_VIRO_4$rrn_min_category[i]<-RERRANG_vs_VIRO_4$rrn6_category[i]
  }
}

for (i in 1:400){
  if (RERRANG_vs_VIRO_4$Number_of_rrn_loci[i]=="8" & RERRANG_vs_VIRO_4$rrn_min_distance[i]==RERRANG_vs_VIRO_4$rrn7_distance
[i]){
  RERRANG_vs_VIRO_4$rrn_min_category[i]<-RERRANG_vs_VIRO_4$rrn7_category[i]
  }
}

for (i in 1:400){
  if (RERRANG_vs_VIRO_4$Number_of_rrn_loci[i]=="8" & RERRANG_vs_VIRO_4$rrn_min_distance[i]==RERRANG_vs_VIRO_4$rrn8_distance
[i]){
  RERRANG_vs_VIRO_4$rrn_min_category[i]<-RERRANG_vs_VIRO_4$rrn8_category[i]
  }
}


write.csv2(RERRANG_vs_VIRO_4, "RERRANG_vs_VIRO_4.csv", row.names = F)
#The remainder of the analysis was manually curated, including comparison to the results of pairwise genome comparisons (Fig
ure S2 and S3), elimination of large insertion events, qualification of rearrangements, and grouping of rearrangements corre
sponding to those present in several strains with a common ancestor. In some cases, the exact position of the rearrangement
was determined by comparison to a closest species (e.g. S. koyakasensis versus S. albidoflavus). Information about rearrange
ments was kept only in cases that were not ambiguous. To avoid introducing biases related to overly complex evolutionary pat
terns, three strains of the group 'O' (S. sp. 11 1 2, S. autolyticus CGMCC0516 and S. bingchenggensis BCW 1) were excluded f
rom this analysis. Thus there were probably more rearrangements than proposed in the scenario (especially in group O). When
the above script had eliminated one of the bounds of the rearrangement, it was searched in the initial data before the appli
cation of filter (CORE, REARRANG_VIRO). This manual curation led to results presented in Supplemental Table 5.
```

```
REARRANG$rrn_min_distance_minimal<-"NA"
REARRANG$rrn_min_distance_maximal<-"NA"
for (i in 1:217){
  j=i+1
  if (REARRANG$genome_id[i]==REARRANG$genome_id[j] & REARRANG$REARRANG_ID[i]==REARRANG$REARRANG_ID[j]){
  REARRANG$rrn_min_distance_minimal[i]<-min(REARRANG$rrn_min_distance[i], REARRANG$rrn_min_distance[j])
  REARRANG$rrn_min_distance_minimal[j]<-min(REARRANG$rrn_min_distance[i], REARRANG$rrn_min_distance[j])
  REARRANG$rrn_min_distance_maximal[i]<-max(REARRANG$rrn_min_distance[i], REARRANG$rrn_min_distance[j])
  REARRANG$rrn_min_distance_maximal[j]<-max(REARRANG$rrn_min_distance[i], REARRANG$rrn_min_distance[j])
  }
}
REARRANG$rrn_min_distance_minimal<-as.numeric(REARRANG$rrn_min_distance_minimal)
REARRANG$rrn_min_distance_maximal<-as.numeric(REARRANG$rrn_min_distance_maximal)

tab1<-aggregate(REARRANG_size~REARRANG_ID, data = REARRANG, FUN=mean)
tab2<-aggregate(rrn_min_distance_minimal~REARRANG_ID, data = REARRANG, FUN=mean)
tab3<-aggregate(rrn_min_distance_maximal~REARRANG_ID, data = REARRANG, FUN=mean)

tab4<-merge(tab1, tab2, by="REARRANG_ID")
tab5<-merge(tab4, tab3, by="REARRANG_ID")
median(tab5$rrn_min_distance_minimal)
```

```
## [1] 92178.25
```

```
write.csv2(tab5, "BILAN_REARRANG.csv")

boxplot(tab5$rrn_min_distance_minimal, tab5$rrn_min_distance_maximal, col = c("yellow", "skyblue2"),names = c("Minimal dista
nce", "Maximal distance"), ylab="Distance to the nearest rrn (bp)",frame=F)
```

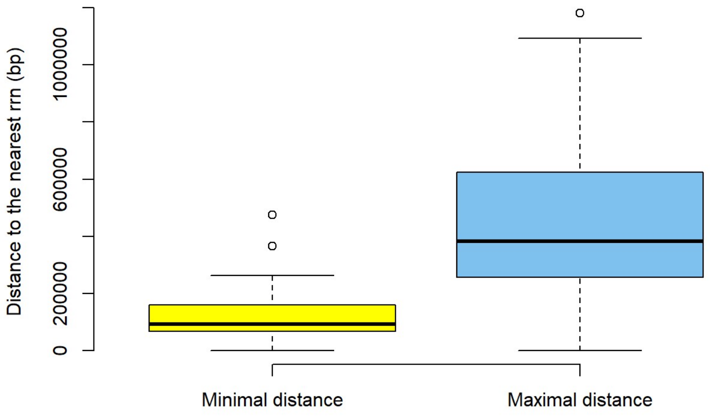


```
REARRANG$rrn_min_category_minimal<-"NA"
for (i in 1:218){
  if (REARRANG$rrn_min_distance[i]==REARRANG$rrn_min_distance_minimal[i]){
  REARRANG$rrn_min_category_minimal[i]<-REARRANG$rrn_min_category[i]
  }
}

REARRANG$rrn_min_category_maximal<-"NA"
for (i in 1:218){
  if (REARRANG$rrn_min_distance[i]==REARRANG$rrn_min_distance_maximal[i]){
  REARRANG$rrn_min_category_maximal[i]<-REARRANG$rrn_min_category[i]
  }
}

#Cleaning to avoid counting twice the [d/e] rrn
for (i in 1:217){
  j=i+1
  if (REARRANG$rrn_min_category_minimal[i]==REARRANG$rrn_min_category_minimal[j] & REARRANG$REARRANG_ID[i]==REARRANG$REARRAN
G_ID[j]){
  REARRANG$rrn_min_category_minimal[i]<-"NA"
  }
}

for (i in 1:217){
  j=i+1
  if (REARRANG$rrn_min_category_maximal[i]==REARRANG$rrn_min_category_maximal[j] & REARRANG$REARRANG_ID[i]==REARRANG$REARRAN
G_ID[j]){
  REARRANG$rrn_min_category_maximal[i]<-"NA"
  }
}

write.csv2(REARRANG, "REARRANG.csv")
```

### Figure 4 _ *rrn* operons operons, as anchors predictive of the core region size

Correlation analysis

```
Clade_1<-subset(ALL, ALL$Clade=="Clade_1")
Clade_2<-subset(ALL, ALL$Clade=="Clade_2")
Group_O<-subset(ALL, ALL$Clade=="Group_O")

ggscatter(ALL, x = "Central_compartment_size", y = "Core_region_size",
          add = "reg.line", conf.int = TRUE,
          cor.coef = TRUE, cor.method = "spearman",
          color = "Clade", palette = c("firebrick3","blue","yellow3"),
          xlab = "Central compartment (bp)", ylab = "Core region size (bp)") + ylim(5000000, 10000000)
```

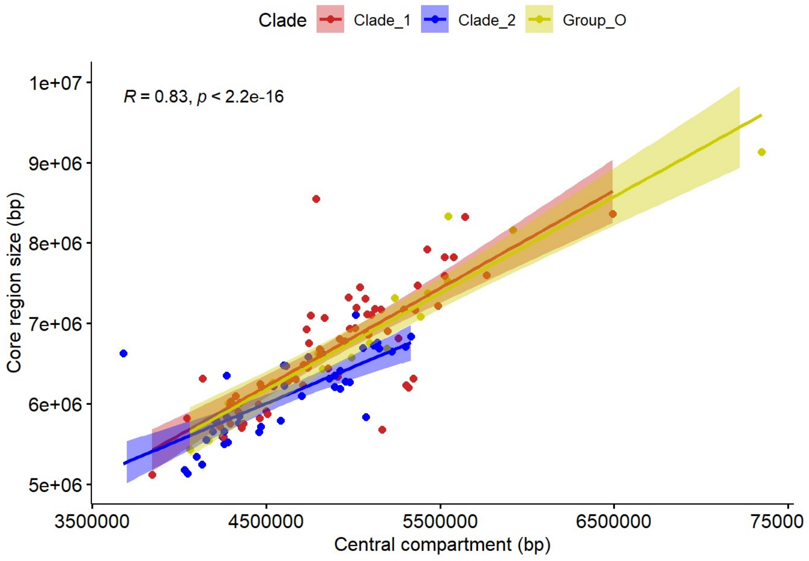


```
cor.test(Clade_1$Central_compartment_size, Clade_1$Core_region_size, method = "spearman")
```

```
##
##  Spearman's rank correlation rho
##
## data:  Clade_1$Central_compartment_size and Clade_1$Core_region_size
## S = 10782, p-value < 2.2e-16
## alternative hypothesis: true rho is not equal to 0
## sample estimates:
##       rho
## 0.7848591
```

```
cor.test(Clade_2$Central_compartment_size, Clade_2$Core_region_size, method = "spearman")
```

```
##
##  Spearman's rank correlation rho
##
## data:  Clade_2$Central_compartment_size and Clade_2$Core_region_size
## S = 2828, p-value = 8.432e-09
## alternative hypothesis: true rho is not equal to 0
## sample estimates:
##       rho
## 0.7864693
```

```
cor.test(Group_O$Central_compartment_size, Group_O$Core_region_size, method = "spearman")
```

```
##
##  Spearman's rank correlation rho
##
## data:  Group_O$Central_compartment_size and Group_O$Core_region_size
## S = 14, p-value = 7.466e-06
## alternative hypothesis: true rho is not equal to 0
## sample estimates:
##       rho
## 0.9828431
```

```
ggscatter(ALL, x = "delta_core_rrn", y = "Core_region_size",
          add = "reg.line", conf.int = TRUE,
          cor.coef = TRUE, cor.method = "spearman",
          color = "Clade", palette = c("firebrick3","blue","yellow3"),
          xlab = "Distance between extreme rrn and core genes (bp)", ylab = "Core region size (bp)") + ylim(5000000, 1000000
0)
```

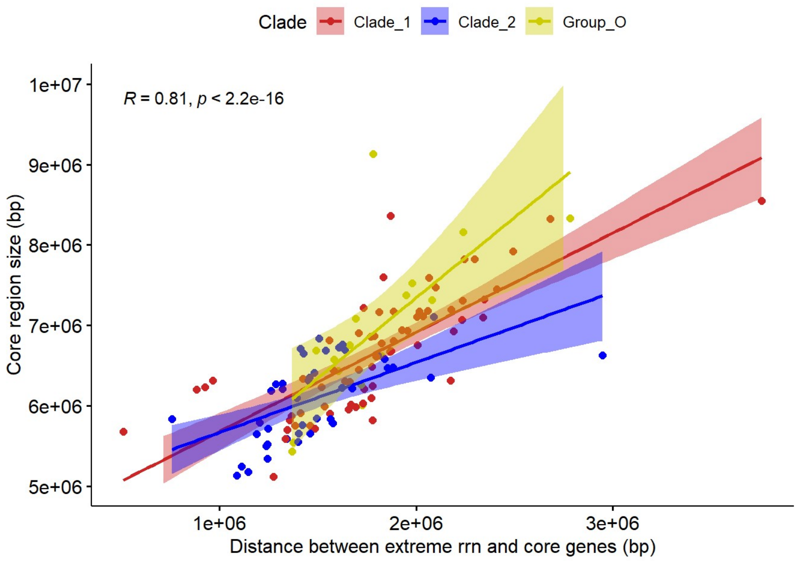


```
cor.test(Clade_1$delta_core_rrn, Clade_1$Core_region_size, method = "spearman")
```

```
##
##  Spearman's rank correlation rho
##
## data:  Clade_1$delta_core_rrn and Clade_1$Core_region_size
## S = 7906, p-value < 2.2e-16
## alternative hypothesis: true rho is not equal to 0
## sample estimates:
##      rho
## 0.842246
```

```
cor.test(Clade_2$delta_core_rrn, Clade_2$Core_region_size, method = "spearman")
```

```
##
##  Spearman's rank correlation rho
##
## data:  Clade_2$delta_core_rrn and Clade_2$Core_region_size
## S = 3804, p-value = 3.322e-07
## alternative hypothesis: true rho is not equal to 0
## sample estimates:
##       rho
## 0.7127756
```

```
cor.test(Group_O$delta_core_rrn, Group_O$Core_region_size, method = "spearman")
```

```
##
##  Spearman's rank correlation rho
##
## data:  Group_O$delta_core_rrn and Group_O$Core_region_size
## S = 162, p-value = 0.0001587
## alternative hypothesis: true rho is not equal to 0
## sample estimates:
##       rho
## 0.8014706
```

```
ggscatter(ALL, x = "Chromosome_size", y = "Core_region_size",
          add = "reg.line", conf.int = TRUE,
          cor.coef = TRUE, cor.method = "spearman",
          color = "Clade", palette = c("firebrick3","blue","yellow3"),
          xlab = "Chromosome size (bp)", ylab = "Core region size (bp)") + ylim(5000000, 10000000)
```

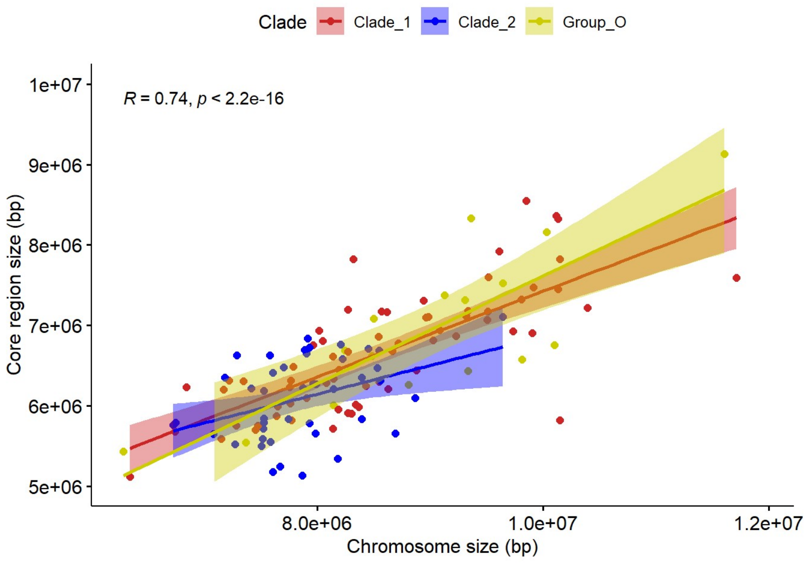


```
cor.test(Clade_1$Chromosome_size, Clade_1$Core_region_size, method = "spearman")
```

```
##
##  Spearman's rank correlation rho
##
## data:  Clade_1$Chromosome_size and Clade_1$Core_region_size
## S = 12274, p-value < 2.2e-16
## alternative hypothesis: true rho is not equal to 0
## sample estimates:
##       rho
## 0.7550882
```

```
cor.test(Clade_2$Chromosome_size, Clade_2$Core_region_size, method = "spearman")
```

```
##
##  Spearman's rank correlation rho
##
## data:  Clade_2$Chromosome_size and Clade_2$Core_region_size
## S = 8088, p-value = 0.01031
## alternative hypothesis: true rho is not equal to 0
## sample estimates:
##       rho
## 0.3893084
```

```
cor.test(Group_O$Chromosome_size, Group_O$Core_region_size, method = "spearman")
```

```
##
##  Spearman's rank correlation rho
##
## data:  Group_O$Chromosome_size and Group_O$Core_region_size
## S = 192, p-value = 0.0005364
## alternative hypothesis: true rho is not equal to 0
## sample estimates:
##       rho
## 0.7647059
```

Modeling_Forward approach

```
DF<-ALL[,c("id", "Core_region_size","Central_compartment_size", "tDNA_region_size", "Chromosome_size", "d_max_rrn_ori", "d_m
in_rrn_ori", "term_comp", "Clade","Number_of_rrn_loci","Number_of_rrn_genes")]
row.names(DF)<-DF$id

predictorlist<-list("Central_compartment_size", "tDNA_region_size", "Chromosome_size", "d_max_rrn_ori", "d_min_rrn_ori", "te
rm_comp", "Clade","Number_of_rrn_loci","Number_of_rrn_genes")

var<-"NA"
res<-"0"
for (i in predictorlist){
  model<-lm(paste("Core_region_size~",i[[1]]), data = DF)
  print(paste("Core_region_size~",i[[1]]))
  print(summary(model)$adj.r.squared)
  var=c(var, paste("Core_region_size~",i[[1]]))
  res=c(res, summary(model)$adj.r.squared)
}
```

```
## [1] "Core_region_size~ Central_compartment_size"
## [1] 0.7326424
## [1] "Core_region_size~ tDNA_region_size"
## [1] 0.1675407
## [1] "Core_region_size~ Chromosome_size"
## [1] 0.6051579
## [1] "Core_region_size~ d_max_rrn_ori"
## [1] 0.7493739
## [1] "Core_region_size~ d_min_rrn_ori"
## [1] 0.4478847
## [1] "Core_region_size~ term_comp"
## [1] 0.1745229
## [1] "Core_region_size~ Clade"
## [1] 0.1659325
## [1] "Core_region_size~ Number_of_rrn_loci"
## [1] 0.07438488
## [1] "Core_region_size~ Number_of_rrn_genes"
## [1] 0.05272926
```

```
result<-data.frame(var, res)
result[which(result$res==max(result$res)),]
```

```
##                                var                res
## 5 Core_region_size~ d_max_rrn_ori 0.749373897423652
```

```
var<-"NA"
res<-"0"
for (i in predictorlist){
  model<-lm(paste("Core_region_size~d_max_rrn_ori+",i[[1]]), data = DF)
  print(paste("Core_region_size~d_max_rrn_ori+",i[[1]]))
  print(summary(model)$adj.r.squared)
  var=c(var, paste("Core_region_size~d_max_rrn_ori+",i[[1]]))
  res=c(res, summary(model)$adj.r.squared)
}
```

```
## [1] "Core_region_size~d_max_rrn_ori+ Central_compartment_size"
## [1] 0.7636316
## [1] "Core_region_size~d_max_rrn_ori+ tDNA_region_size"
## [1] 0.7640709
## [1] "Core_region_size~d_max_rrn_ori+ Chromosome_size"
## [1] 0.7928808
## [1] "Core_region_size~d_max_rrn_ori+ d_max_rrn_ori"
## [1] 0.7493739
## [1] "Core_region_size~d_max_rrn_ori+ d_min_rrn_ori"
## [1] 0.7636316
## [1] "Core_region_size~d_max_rrn_ori+ term_comp"
## [1] 0.7762893
## [1] "Core_region_size~d_max_rrn_ori+ Clade"
## [1] 0.7593171
## [1] "Core_region_size~d_max_rrn_ori+ Number_of_rrn_loci"
## [1] 0.7583629
## [1] "Core_region_size~d_max_rrn_ori+ Number_of_rrn_genes"
## [1] 0.7643517
```

```
result<-data.frame(var, res)
result[which(result$res==max(result$res)),]
```

```
##                                                var                res
## 4 Core_region_size~d_max_rrn_ori+ Chromosome_size 0.792880815347247
```

```
var<-"NA"
res<-"0"
for (i in predictorlist){
  model<-lm(paste("Core_region_size~d_max_rrn_ori+Chromosome_size+",i[[1]]), data = DF)
  print(paste("Core_region_size~d_max_rrn_ori+Chromosome_size+",i[[1]]))
  print(summary(model)$adj.r.squared)
  var=c(var, paste("Core_region_size~d_max_rrn_ori+Chromosome_size+",i[[1]]))
  res=c(res, summary(model)$adj.r.squared)
}
```

```
## [1] "Core_region_size~d_max_rrn_ori+Chromosome_size+ Central_compartment_size"
## [1] 0.8103039
## [1] "Core_region_size~d_max_rrn_ori+Chromosome_size+ tDNA_region_size"
## [1] 0.7914119
## [1] "Core_region_size~d_max_rrn_ori+Chromosome_size+ Chromosome_size"
## [1] 0.7928808
## [1] "Core_region_size~d_max_rrn_ori+Chromosome_size+ d_max_rrn_ori"
## [1] 0.7928808
## [1] "Core_region_size~d_max_rrn_ori+Chromosome_size+ d_min_rrn_ori"
## [1] 0.8103039
## [1] "Core_region_size~d_max_rrn_ori+Chromosome_size+ term_comp"
## [1] 0.8103039
## [1] "Core_region_size~d_max_rrn_ori+Chromosome_size+ Clade"
## [1] 0.7957445
## [1] "Core_region_size~d_max_rrn_ori+Chromosome_size+ Number_of_rrn_loci"
## [1] 0.8062495
## [1] "Core_region_size~d_max_rrn_ori+Chromosome_size+ Number_of_rrn_genes"
## [1] 0.8123109
```

```
result<-data.frame(var, res)
result[which(result$res==max(result$res)),]
```

```
##                                                                      var
## 10 Core_region_size~d_max_rrn_ori+Chromosome_size+ Number_of_rrn_genes
##                  res
## 10 0.81231091999527
```

```
var<-"NA"
res<-"0"
for (i in predictorlist){
  model<-lm(paste("Core_region_size~d_max_rrn_ori+Chromosome_size+Number_of_rrn_genes+",i[[1]]), data = DF)
  print(paste("Core_region_size~d_max_rrn_ori+Chromosome_size+Number_of_rrn_genes+",i[[1]]))
  print(summary(model)$adj.r.squared)
  var=c(var, paste("Core_region_size~d_max_rrn_ori+Chromosome_size+Number_of_rrn_genes+",i[[1]]))
  res=c(res, summary(model)$adj.r.squared)
}
```

```
## [1] "Core_region_size~d_max_rrn_ori+Chromosome_size+Number_of_rrn_genes+ Central_compartment_size"
## [1] 0.8357701
## [1] "Core_region_size~d_max_rrn_ori+Chromosome_size+Number_of_rrn_genes+ tDNA_region_size"
## [1] 0.8107868
## [1] "Core_region_size~d_max_rrn_ori+Chromosome_size+Number_of_rrn_genes+ Chromosome_size"
## [1] 0.8123109
## [1] "Core_region_size~d_max_rrn_ori+Chromosome_size+Number_of_rrn_genes+ d_max_rrn_ori"
## [1] 0.8123109
## [1] "Core_region_size~d_max_rrn_ori+Chromosome_size+Number_of_rrn_genes+ d_min_rrn_ori"
## [1] 0.8357701
## [1] "Core_region_size~d_max_rrn_ori+Chromosome_size+Number_of_rrn_genes+ term_comp"
## [1] 0.8357701
## [1] "Core_region_size~d_max_rrn_ori+Chromosome_size+Number_of_rrn_genes+ Clade"
## [1] 0.8107369
## [1] "Core_region_size~d_max_rrn_ori+Chromosome_size+Number_of_rrn_genes+ Number_of_rrn_loci"
## [1] 0.8138512
## [1] "Core_region_size~d_max_rrn_ori+Chromosome_size+Number_of_rrn_genes+ Number_of_rrn_genes"
## [1] 0.8123109
```

```
result<-data.frame(var, res)
result[which(result$res==max(result$res)),]
```

```
##                                                                                             var
## 2 Core_region_size~d_max_rrn_ori+Chromosome_size+Number_of_rrn_genes+ Central_compartment_size
## 6            Core_region_size~d_max_rrn_ori+Chromosome_size+Number_of_rrn_genes+ d_min_rrn_ori
## 7                Core_region_size~d_max_rrn_ori+Chromosome_size+Number_of_rrn_genes+ term_comp
##                  res
## 2 0.835770121684967
## 6 0.835770121684967
## 7 0.835770121684967
```

```
var<-"NA"
res<-"0"
for (i in predictorlist){
  model<-lm(paste("Core_region_size~d_max_rrn_ori+Chromosome_size+Number_of_rrn_genes+Central_compartment_size+",i[[1]]), da
ta = DF)
  print(paste("Core_region_size~d_max_rrn_ori+Chromosome_size+Number_of_rrn_genes+Central_compartment_size+",i[[1]]))
  print(summary(model)$adj.r.squared)
  var=c(var, paste("Core_region_size~d_max_rrn_ori+Chromosome_size+Number_of_rrn_genes+Central_compartment_size+",i[[1]]))
  res=c(res, summary(model)$adj.r.squared)
}
```

```
## [1] "Core_region_size~d_max_rrn_ori+Chromosome_size+Number_of_rrn_genes+Central_compartment_size+ Central_compartment_siz
e"
## [1] 0.8357701
## [1] "Core_region_size~d_max_rrn_ori+Chromosome_size+Number_of_rrn_genes+Central_compartment_size+ tDNA_region_size"
## [1] 0.8346461
## [1] "Core_region_size~d_max_rrn_ori+Chromosome_size+Number_of_rrn_genes+Central_compartment_size+ Chromosome_size"
## [1] 0.8357701
## [1] "Core_region_size~d_max_rrn_ori+Chromosome_size+Number_of_rrn_genes+Central_compartment_size+ d_max_rrn_ori"
## [1] 0.8357701
## [1] "Core_region_size~d_max_rrn_ori+Chromosome_size+Number_of_rrn_genes+Central_compartment_size+ d_min_rrn_ori"
## [1] 0.8357701
## [1] "Core_region_size~d_max_rrn_ori+Chromosome_size+Number_of_rrn_genes+Central_compartment_size+ term_comp"
## [1] 0.8357701
## [1] "Core_region_size~d_max_rrn_ori+Chromosome_size+Number_of_rrn_genes+Central_compartment_size+ Clade"
## [1] 0.8361823
## [1] "Core_region_size~d_max_rrn_ori+Chromosome_size+Number_of_rrn_genes+Central_compartment_size+ Number_of_rrn_loci"
## [1] 0.8379817
## [1] "Core_region_size~d_max_rrn_ori+Chromosome_size+Number_of_rrn_genes+Central_compartment_size+ Number_of_rrn_genes"
## [1] 0.8357701
```

```
result<-data.frame(var, res)
result[which(result$res==max(result$res)),]
```

```
##                                                                                                             var
## 9 Core_region_size~d_max_rrn_ori+Chromosome_size+Number_of_rrn_genes+Central_compartment_size+ Number_of_rrn_loci
##                 res
## 9 0.837981669435844
```

```
var<-"NA"
res<-"0"
for (i in predictorlist){
  model<-lm(paste("Core_region_size~d_max_rrn_ori+Chromosome_size+Number_of_rrn_genes+Central_compartment_size+Number_of_rrn
_loci+",i[[1]]), data = DF)
  print(paste("Core_region_size~d_max_rrn_ori+Chromosome_size+Number_of_rrn_genes+Central_compartment_size+Number_of_rrn_loc
i+",i[[1]]))
  print(summary(model)$adj.r.squared)
  var=c(var, paste("Core_region_size~d_max_rrn_ori+Chromosome_size+Number_of_rrn_genes+Central_compartment_size+Number_of_rr
n_loci+",i[[1]]))
  res=c(res, summary(model)$adj.r.squared)
}
```

```
## [1] "Core_region_size~d_max_rrn_ori+Chromosome_size+Number_of_rrn_genes+Central_compartment_size+Number_of_rrn_loci+ Cent
ral_compartment_size"
## [1] 0.8379817
## [1] "Core_region_size~d_max_rrn_ori+Chromosome_size+Number_of_rrn_genes+Central_compartment_size+Number_of_rrn_loci+ tDNA
_region_size"
## [1] 0.8369084
## [1] "Core_region_size~d_max_rrn_ori+Chromosome_size+Number_of_rrn_genes+Central_compartment_size+Number_of_rrn_loci+ Chro
mosome_size"
## [1] 0.8379817
## [1] "Core_region_size~d_max_rrn_ori+Chromosome_size+Number_of_rrn_genes+Central_compartment_size+Number_of_rrn_loci+ d_ma
x_rrn_ori"
## [1] 0.8379817
## [1] "Core_region_size~d_max_rrn_ori+Chromosome_size+Number_of_rrn_genes+Central_compartment_size+Number_of_rrn_loci+ d_mi
n_rrn_ori"
## [1] 0.8379817
## [1] "Core_region_size~d_max_rrn_ori+Chromosome_size+Number_of_rrn_genes+Central_compartment_size+Number_of_rrn_loci+ term
_comp"
## [1] 0.8379817
## [1] "Core_region_size~d_max_rrn_ori+Chromosome_size+Number_of_rrn_genes+Central_compartment_size+Number_of_rrn_loci+ Clad
e"
## [1] 0.8379125
## [1] "Core_region_size~d_max_rrn_ori+Chromosome_size+Number_of_rrn_genes+Central_compartment_size+Number_of_rrn_loci+ Numb
er_of_rrn_loci"
## [1] 0.8379817
## [1] "Core_region_size~d_max_rrn_ori+Chromosome_size+Number_of_rrn_genes+Central_compartment_size+Number_of_rrn_loci+ Numb
er_of_rrn_genes"
## [1] 0.8379817
```

```
result<-data.frame(var, res)
result[which(result$res==max(result$res)),]
```

```
##
var
## 2  Core_region_size~d_max_rrn_ori+Chromosome_size+Number_of_rrn_genes+Central_compartment_size+Number_of_rrn_loci+ Centra
l_compartment_size
## 4           Core_region_size~d_max_rrn_ori+Chromosome_size+Number_of_rrn_genes+Central_compartment_size+Number_of_rrn_loc
i+ Chromosome_size
## 5             Core_region_size~d_max_rrn_ori+Chromosome_size+Number_of_rrn_genes+Central_compartment_size+Number_of_rrn_l
oci+ d_max_rrn_ori
## 6             Core_region_size~d_max_rrn_ori+Chromosome_size+Number_of_rrn_genes+Central_compartment_size+Number_of_rrn_l
oci+ d_min_rrn_ori
## 7                 Core_region_size~d_max_rrn_ori+Chromosome_size+Number_of_rrn_genes+Central_compartment_size+Number_of_r
rn_loci+ term_comp
## 9        Core_region_size~d_max_rrn_ori+Chromosome_size+Number_of_rrn_genes+Central_compartment_size+Number_of_rrn_loci+
Number_of_rrn_loci
## 10      Core_region_size~d_max_rrn_ori+Chromosome_size+Number_of_rrn_genes+Central_compartment_size+Number_of_rrn_loci+ N
umber_of_rrn_genes
##                  res
## 2  0.837981669435844
## 4  0.837981669435844
## 5  0.837981669435844
## 6  0.837981669435844
## 7  0.837981669435844
## 9  0.837981669435844
## 10 0.837981669435844
```

```
LM0<-lm(Core_region_size~1, DF)
LM1<-lm(Core_region_size~d_max_rrn_ori, DF)
LM2<-lm(Core_region_size~d_max_rrn_ori+Chromosome_size, DF)
LM3<-lm(Core_region_size~d_max_rrn_ori+Chromosome_size+Number_of_rrn_genes, DF)
LM4<-lm(Core_region_size~d_max_rrn_ori+Chromosome_size+Number_of_rrn_genes+Central_compartment_size, DF)
LM5<-lm(Core_region_size~d_max_rrn_ori+Chromosome_size+Number_of_rrn_genes+Central_compartment_size+Number_of_rrn_loci, DF)

AIC(LM0, LM1, LM2, LM3, LM4,LM5)
```

```
##     df      AIC
## LM0  2 3824.509
## LM1  3 3649.755
## LM2  4 3626.520
## LM3  5 3614.982
## LM4  6 3598.988
## LM5  7 3598.221
```

```
#Based on the AIC, the best model is LM5. Hower, we noticed that S; asterosporus has a too strong leverage effect with this
model, see diagnostic test below:
par(mfrow=c(2,2))
plot(LM5,cex.lab=1.5,cex.main=1.5,pch=16)
```

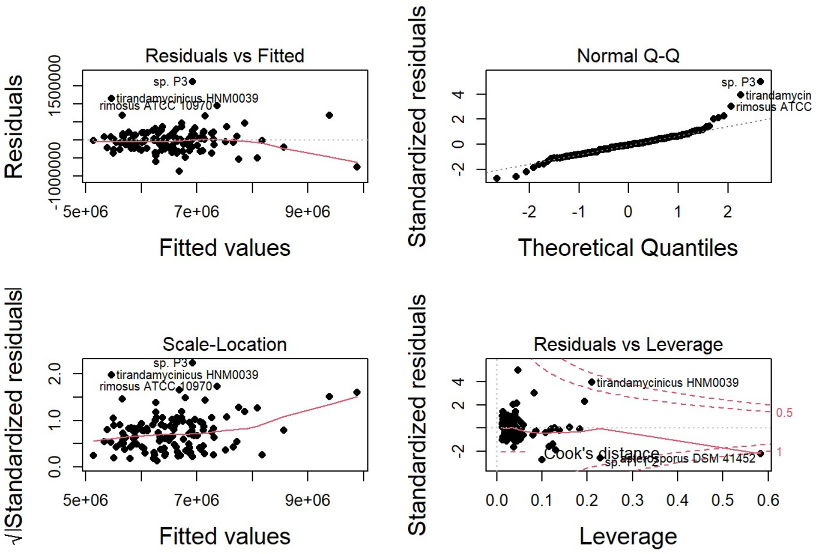

```r
#A second run of analysis without S. asterosporus (too strong leverage effect)
ALL_wo_astero<-subset(ALL, ALL$Species!="Streptomyces asterosporus DSM 41452")
DF<-ALL_wo_astero[,c("id", "Core_region_size","Central_compartment_size", "tDNA_region_size", "Chromosome_size", "d_max_rrn_ori", "d_min_rrn_ori", "term_comp", "Clade","Number_of_rrn_loci","Number_of_rrn_genes")]
row.names(DF)<-DF$id

predictorlist<-list("Central_compartment_size", "tDNA_region_size", "Chromosome_size", "d_max_rrn_ori", "d_min_rrn_ori","term_comp","Clade","Number_of_rrn_loci","Number_of_rrn_genes")

var<-"NA"
res<-"0"
for (i in predictorlist){
  model<-lm(paste("Core_region_size~",i[[1]]), data = DF)
  print(paste("Core_region_size~",i[[1]]))
  print(summary(model)$adj.r.squared)
  var=c(var, paste("Core_region_size~",i[[1]]))
  res=c(res, summary(model)$adj.r.squared)
}
```

```
## [1] "Core_region_size~ Central_compartment_size"
## [1] 0.7374612
## [1] "Core_region_size~ tDNA_region_size"
## [1] 0.1672942
## [1] "Core_region_size~ Chromosome_size"
## [1] 0.6052676
## [1] "Core_region_size~ d_max_rrn_ori"
## [1] 0.7501272
## [1] "Core_region_size~ d_min_rrn_ori"
## [1] 0.4537875
## [1] "Core_region_size~ term_comp"
## [1] 0.1746816
## [1] "Core_region_size~ Clade"
## [1] 0.1668331
## [1] "Core_region_size~ Number_of_rrn_loci"
## [1] 0.08232904
## [1] "Core_region_size~ Number_of_rrn_genes"
## [1] 0.07327507
```

```r
result<-data.frame(var, res)
result[which(result$res==max(result$res)),]
```

```
##                                var               res
## 5 Core_region_size~ d_max_rrn_ori 0.750127160373274
```

```r
var<-"NA"
res<-"0"
for (i in predictorlist){
  model<-lm(paste("Core_region_size~d_max_rrn_ori+",i[[1]]), data = DF)
  print(paste("Core_region_size~d_max_rrn_ori+",i[[1]]))
  print(summary(model)$adj.r.squared)
  var=c(var, paste("Core_region_size~d_max_rrn_ori+",i[[1]]))
  res=c(res, summary(model)$adj.r.squared)
}
```

```
## [1] "Core_region_size~d_max_rrn_ori+ Central_compartment_size"
## [1] 0.7659475
## [1] "Core_region_size~d_max_rrn_ori+ tDNA_region_size"
## [1] 0.7647886
## [1] "Core_region_size~d_max_rrn_ori+ Chromosome_size"
## [1] 0.793849
## [1] "Core_region_size~d_max_rrn_ori+ d_max_rrn_ori"
## [1] 0.7501272
## [1] "Core_region_size~d_max_rrn_ori+ d_min_rrn_ori"
## [1] 0.7659475
## [1] "Core_region_size~d_max_rrn_ori+ term_comp"
## [1] 0.7768112
## [1] "Core_region_size~d_max_rrn_ori+ Clade"
## [1] 0.7595086
## [1] "Core_region_size~d_max_rrn_ori+ Number_of_rrn_loci"
## [1] 0.7582093
## [1] "Core_region_size~d_max_rrn_ori+ Number_of_rrn_genes"
## [1] 0.7651483
```

```r
result<-data.frame(var, res)
result[which(result$res==max(result$res)),]
```

```
##                                                var               res
## 4 Core_region_size~d_max_rrn_ori+ Chromosome_size 0.793849001950542
```

```r
var<-"NA"
res<-"0"
for (i in predictorlist){
  model<-lm(paste("Core_region_size~d_max_rrn_ori+Chromosome_size+",i[[1]]), data = DF)
  print(paste("Core_region_size~d_max_rrn_ori+Chromosome_size+",i[[1]]))
  print(summary(model)$adj.r.squared)
  var=c(var, paste("Core_region_size~d_max_rrn_ori+Chromosome_size+",i[[1]]))
  res=c(res, summary(model)$adj.r.squared)
}
```

```
## [1] "Core_region_size~d_max_rrn_ori+Chromosome_size+ Central_compartment_size"
## [1] 0.813154
## [1] "Core_region_size~d_max_rrn_ori+Chromosome_size+ tDNA_region_size"
## [1] 0.7923932
## [1] "Core_region_size~d_max_rrn_ori+Chromosome_size+ Chromosome_size"
## [1] 0.793849
## [1] "Core_region_size~d_max_rrn_ori+Chromosome_size+ d_max_rrn_ori"
## [1] 0.793849
## [1] "Core_region_size~d_max_rrn_ori+Chromosome_size+ d_min_rrn_ori"
## [1] 0.813154
## [1] "Core_region_size~d_max_rrn_ori+Chromosome_size+ term_comp"
## [1] 0.813154
## [1] "Core_region_size~d_max_rrn_ori+Chromosome_size+ Clade"
## [1] 0.7962647
## [1] "Core_region_size~d_max_rrn_ori+Chromosome_size+ Number_of_rrn_loci"
## [1] 0.8061078
## [1] "Core_region_size~d_max_rrn_ori+Chromosome_size+ Number_of_rrn_genes"
## [1] 0.8133905
```

```r
result<-data.frame(var, res)
result[which(result$res==max(result$res)),]
```

```
##                                                                     var
## 10 Core_region_size~d_max_rrn_ori+Chromosome_size+ Number_of_rrn_genes
##                   res
## 10 0.813390504869708
```

```r
var<-"NA"
res<-"0"
for (i in predictorlist){
  model<-lm(paste("Core_region_size~d_max_rrn_ori+Chromosome_size+Number_of_rrn_genes+",i[[1]]), data = DF)
  print(paste("Core_region_size~d_max_rrn_ori+Chromosome_size+Number_of_rrn_genes+",i[[1]]))
  print(summary(model)$adj.r.squared)
  var=c(var, paste("Core_region_size~d_max_rrn_ori+Chromosome_size+Number_of_rrn_genes+",i[[1]]))
  res=c(res, summary(model)$adj.r.squared)
}
```

```
## [1] "Core_region_size~d_max_rrn_ori+Chromosome_size+Number_of_rrn_genes+ Central_compartment_size"
## [1] 0.8359765
## [1] "Core_region_size~d_max_rrn_ori+Chromosome_size+Number_of_rrn_genes+ tDNA_region_size"
## [1] 0.8118945
## [1] "Core_region_size~d_max_rrn_ori+Chromosome_size+Number_of_rrn_genes+ Chromosome_size"
## [1] 0.8133905
## [1] "Core_region_size~d_max_rrn_ori+Chromosome_size+Number_of_rrn_genes+ d_max_rrn_ori"
## [1] 0.8133905
## [1] "Core_region_size~d_max_rrn_ori+Chromosome_size+Number_of_rrn_genes+ d_min_rrn_ori"
## [1] 0.8359765
## [1] "Core_region_size~d_max_rrn_ori+Chromosome_size+Number_of_rrn_genes+ term_comp"
## [1] 0.8359765
## [1] "Core_region_size~d_max_rrn_ori+Chromosome_size+Number_of_rrn_genes+ Clade"
## [1] 0.8120444
## [1] "Core_region_size~d_max_rrn_ori+Chromosome_size+Number_of_rrn_genes+ Number_of_rrn_loci"
## [1] 0.8236773
## [1] "Core_region_size~d_max_rrn_ori+Chromosome_size+Number_of_rrn_genes+ Number_of_rrn_genes"
## [1] 0.8133905
```

```r
result<-data.frame(var, res)
result[which(result$res==max(result$res)),]
```

```
##                                                                                         var
## 2 Core_region_size~d_max_rrn_ori+Chromosome_size+Number_of_rrn_genes+ Central_compartment_size
## 6            Core_region_size~d_max_rrn_ori+Chromosome_size+Number_of_rrn_genes+ d_min_rrn_ori
## 7                Core_region_size~d_max_rrn_ori+Chromosome_size+Number_of_rrn_genes+ term_comp
##                 res
## 2 0.835976485318349
## 6 0.835976485318349
## 7 0.835976485318349
```

```
var<-"NA"
res<-"0"
for (i in predictorlist){
  model<-lm(paste("Core_region_size~d_max_rrn_ori+Chromosome_size+Number_of_rrn_genes+Central_compartment_size+",i[[1]]), da
ta = DF)
  print(paste("Core_region_size~d_max_rrn_ori+Chromosome_size+Number_of_rrn_genes+Central_compartment_size+",i[[1]]))
  print(summary(model)$adj.r.squared)
  var=c(var, paste("Core_region_size~d_max_rrn_ori+Chromosome_size+Number_of_rrn_genes+Central_compartment_size+",i[[1]]))
  res=c(res, summary(model)$adj.r.squared)
}
```

```
## [1] "Core_region_size~d_max_rrn_ori+Chromosome_size+Number_of_rrn_genes+Central_compartment_size+ Central_compartment_siz
e"
## [1] 0.8359765
## [1] "Core_region_size~d_max_rrn_ori+Chromosome_size+Number_of_rrn_genes+Central_compartment_size+ tDNA_region_size"
## [1] 0.8347919
## [1] "Core_region_size~d_max_rrn_ori+Chromosome_size+Number_of_rrn_genes+Central_compartment_size+ Chromosome_size"
## [1] 0.8359765
## [1] "Core_region_size~d_max_rrn_ori+Chromosome_size+Number_of_rrn_genes+Central_compartment_size+ d_max_rrn_ori"
## [1] 0.8359765
## [1] "Core_region_size~d_max_rrn_ori+Chromosome_size+Number_of_rrn_genes+Central_compartment_size+ d_min_rrn_ori"
## [1] 0.8359765
## [1] "Core_region_size~d_max_rrn_ori+Chromosome_size+Number_of_rrn_genes+Central_compartment_size+ term_comp"
## [1] 0.8359765
## [1] "Core_region_size~d_max_rrn_ori+Chromosome_size+Number_of_rrn_genes+Central_compartment_size+ Clade"
## [1] 0.8367111
## [1] "Core_region_size~d_max_rrn_ori+Chromosome_size+Number_of_rrn_genes+Central_compartment_size+ Number_of_rrn_loci"
## [1] 0.8444919
## [1] "Core_region_size~d_max_rrn_ori+Chromosome_size+Number_of_rrn_genes+Central_compartment_size+ Number_of_rrn_genes"
## [1] 0.8359765
```

```
result<-data.frame(var, res)
result[which(result$res==max(result$res)),]
```

```
##                                                                                                               var
## 9 Core_region_size~d_max_rrn_ori+Chromosome_size+Number_of_rrn_genes+Central_compartment_size+ Number_of_rrn_loci
##                  res
## 9 0.844491890327038
```

```
var<-"NA"
res<-"0"
for (i in predictorlist){
  model<-lm(paste("Core_region_size~d_max_rrn_ori+Chromosome_size+Number_of_rrn_genes+Central_compartment_size+Number_of_rrn
_loci+",i[[1]]), data = DF)
  print(paste("Core_region_size~d_max_rrn_ori+Chromosome_size+Number_of_rrn_genes+Central_compartment_size+Number_of_rrn_loc
i+",i[[1]]))
  print(summary(model)$adj.r.squared)
  var=c(var, paste("Core_region_size~d_max_rrn_ori+Chromosome_size+Number_of_rrn_genes+Central_compartment_size+Number_of_rr
n_loci+",i[[1]]))
  res=c(res, summary(model)$adj.r.squared)
}
```

```
## [1] "Core_region_size~d_max_rrn_ori+Chromosome_size+Number_of_rrn_genes+Central_compartment_size+Number_of_rrn_loci+ Cent
ral_compartment_size"
## [1] 0.8444919
## [1] "Core_region_size~d_max_rrn_ori+Chromosome_size+Number_of_rrn_genes+Central_compartment_size+Number_of_rrn_loci+ tDNA
_region_size"
## [1] 0.8432382
## [1] "Core_region_size~d_max_rrn_ori+Chromosome_size+Number_of_rrn_genes+Central_compartment_size+Number_of_rrn_loci+ Chro
mosome_size"
## [1] 0.8444919
## [1] "Core_region_size~d_max_rrn_ori+Chromosome_size+Number_of_rrn_genes+Central_compartment_size+Number_of_rrn_loci+ d_ma
x_rrn_ori"
## [1] 0.8444919
## [1] "Core_region_size~d_max_rrn_ori+Chromosome_size+Number_of_rrn_genes+Central_compartment_size+Number_of_rrn_loci+ d_mi
n_rrn_ori"
## [1] 0.8444919
## [1] "Core_region_size~d_max_rrn_ori+Chromosome_size+Number_of_rrn_genes+Central_compartment_size+Number_of_rrn_loci+ term
_comp"
## [1] 0.8444919
## [1] "Core_region_size~d_max_rrn_ori+Chromosome_size+Number_of_rrn_genes+Central_compartment_size+Number_of_rrn_loci+ Clad
e"
## [1] 0.8452709
## [1] "Core_region_size~d_max_rrn_ori+Chromosome_size+Number_of_rrn_genes+Central_compartment_size+Number_of_rrn_loci+ Numb
er_of_rrn_loci"
## [1] 0.8444919
## [1] "Core_region_size~d_max_rrn_ori+Chromosome_size+Number_of_rrn_genes+Central_compartment_size+Number_of_rrn_loci+ Numb
er_of_rrn_genes"
## [1] 0.8444919
```

```
result<-data.frame(var, res)
result[which(result$res==max(result$res)),]
```

```
##                                                                                                                    var
## 8 Core_region_size~d_max_rrn_ori+Chromosome_size+Number_of_rrn_genes+Central_compartment_size+Number_of_rrn_loci+ Clade
##                 res
## 8 0.845270946845142
```

```
var<-"NA"
res<-"0"
for (i in predictorlist){
  model<-lm(paste("Core_region_size~d_max_rrn_ori+Chromosome_size+Number_of_rrn_genes+Central_compartment_size+Number_of_rrn
_loci+Clade+",i[[1]]), data = DF)
  print(paste("Core_region_size~d_max_rrn_ori+Chromosome_size+Number_of_rrn_genes+Central_compartment_size+Number_of_rrn_loc
i+Clade+",i[[1]]))
  print(summary(model)$adj.r.squared)
  var=c(var, paste("Core_region_size~d_max_rrn_ori+Chromosome_size+Number_of_rrn_genes+Central_compartment_size+Number_of_rr
n_loci+Clade+",i[[1]]))
  res=c(res, summary(model)$adj.r.squared)
}
```

```
## [1] "Core_region_size~d_max_rrn_ori+Chromosome_size+Number_of_rrn_genes+Central_compartment_size+Number_of_rrn_loci+Clad
e+ Central_compartment_size"
## [1] 0.8452709
## [1] "Core_region_size~d_max_rrn_ori+Chromosome_size+Number_of_rrn_genes+Central_compartment_size+Number_of_rrn_loci+Clad
e+ tDNA_region_size"
## [1] 0.8439611
## [1] "Core_region_size~d_max_rrn_ori+Chromosome_size+Number_of_rrn_genes+Central_compartment_size+Number_of_rrn_loci+Clad
e+ Chromosome_size"
## [1] 0.8452709
## [1] "Core_region_size~d_max_rrn_ori+Chromosome_size+Number_of_rrn_genes+Central_compartment_size+Number_of_rrn_loci+Clad
e+ d_max_rrn_ori"
## [1] 0.8452709
## [1] "Core_region_size~d_max_rrn_ori+Chromosome_size+Number_of_rrn_genes+Central_compartment_size+Number_of_rrn_loci+Clad
e+ d_min_rrn_ori"
## [1] 0.8452709
## [1] "Core_region_size~d_max_rrn_ori+Chromosome_size+Number_of_rrn_genes+Central_compartment_size+Number_of_rrn_loci+Clad
e+ term_comp"
## [1] 0.8452709
## [1] "Core_region_size~d_max_rrn_ori+Chromosome_size+Number_of_rrn_genes+Central_compartment_size+Number_of_rrn_loci+Clad
e+ Clade"
## [1] 0.8452709
## [1] "Core_region_size~d_max_rrn_ori+Chromosome_size+Number_of_rrn_genes+Central_compartment_size+Number_of_rrn_loci+Clad
e+ Number_of_rrn_loci"
## [1] 0.8452709
## [1] "Core_region_size~d_max_rrn_ori+Chromosome_size+Number_of_rrn_genes+Central_compartment_size+Number_of_rrn_loci+Clad
e+ Number_of_rrn_genes"
## [1] 0.8452709
```

```
result<-data.frame(var, res)
result[which(result$res==max(result$res)),]
```

```
##
var
## 2  Core_region_size~d_max_rrn_ori+Chromosome_size+Number_of_rrn_genes+Central_compartment_size+Number_of_rrn_loci+Clade+
Central_compartment_size
## 4           Core_region_size~d_max_rrn_ori+Chromosome_size+Number_of_rrn_genes+Central_compartment_size+Number_of_rrn_loc
i+Clade+ Chromosome_size
## 5             Core_region_size~d_max_rrn_ori+Chromosome_size+Number_of_rrn_genes+Central_compartment_size+Number_of_rrn_l
oci+Clade+ d_max_rrn_ori
## 6             Core_region_size~d_max_rrn_ori+Chromosome_size+Number_of_rrn_genes+Central_compartment_size+Number_of_rrn_l
oci+Clade+ d_min_rrn_ori
## 7                 Core_region_size~d_max_rrn_ori+Chromosome_size+Number_of_rrn_genes+Central_compartment_size+Number_of_r
rn_loci+Clade+ term_comp
## 8                     Core_region_size~d_max_rrn_ori+Chromosome_size+Number_of_rrn_genes+Central_compartment_size+Number_
of_rrn_loci+Clade+ Clade
## 9        Core_region_size~d_max_rrn_ori+Chromosome_size+Number_of_rrn_genes+Central_compartment_size+Number_of_rrn_loci+C
lade+ Number_of_rrn_loci
## 10      Core_region_size~d_max_rrn_ori+Chromosome_size+Number_of_rrn_genes+Central_compartment_size+Number_of_rrn_loci+Cl
ade+ Number_of_rrn_genes
##                  res
## 2  0.845270946845142
## 4  0.845270946845142
## 5  0.845270946845142
## 6  0.845270946845142
## 7  0.845270946845142
## 8  0.845270946845142
## 9  0.845270946845142
## 10 0.845270946845142
```

```
LM0<-lm(Core_region_size~1, DF)
LM1<-lm(Core_region_size~d_max_rrn_ori, DF)
LM2<-lm(Core_region_size~d_max_rrn_ori+Chromosome_size, DF)
LM3<-lm(Core_region_size~d_max_rrn_ori+Chromosome_size+Number_of_rrn_genes, DF)
LM4<-lm(Core_region_size~d_max_rrn_ori+Chromosome_size+Number_of_rrn_genes+Central_compartment_size, DF)
LM5<-lm(Core_region_size~d_max_rrn_ori+Chromosome_size+Number_of_rrn_genes+Central_compartment_size+Number_of_rrn_loci, DF)
LM6<-lm(Core_region_size~d_max_rrn_ori+Chromosome_size+Number_of_rrn_genes+Central_compartment_size+Number_of_rrn_loci+Clad
e, DF)

AIC(LM0, LM1, LM2, LM3, LM4,LM5, LM6)
```

```
##     df      AIC
## LM0  2 3795.348
## LM1  3 3621.599
## LM2  4 3598.343
## LM3  5 3586.766
## LM4  6 3571.474
## LM5  7 3565.711
## LM6  9 3566.960
```

```
#Based on the AIC, LM5 is the best model but S. tirandamycinius has a strong leverage effect.
par(mfrow=c(2,2))
plot(LM5,cex.lab=1.5,cex.main=1.5,pch=16)
```

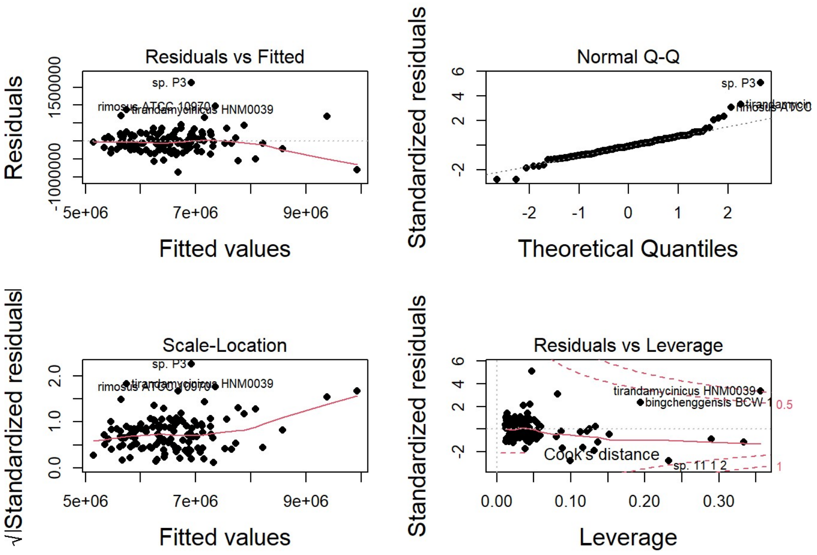


```
#A Third run of analysis without S. asterosporus and S. tirandamycinius(too strong leverage effect)
ALL_wo_astero_wo_tiran<-subset(ALL, ALL$Species!="Streptomyces asterosporus DSM 41452" & ALL$Species!="Streptomyces tirandam
ycinicus HNM0039")
DF<-ALL_wo_astero_wo_tiran[,c("id", "Core_region_size","Central_compartment_size", "tDNA_region_size", "Chromosome_size", "d
_max_rrn_ori", "d_min_rrn_ori", "term_comp", "Clade","Number_of_rrn_loci","Number_of_rrn_genes")]
row.names(DF)<-DF$id

predictorlist<-list("Central_compartment_size", "tDNA_region_size", "Chromosome_size", "d_max_rrn_ori", "d_min_rrn_ori","ter
m_comp","Clade","Number_of_rrn_loci","Number_of_rrn_genes")

var<-"NA"
res<-"0"
for (i in predictorlist){
  model<-lm(paste("Core_region_size~",i[[1]]), data = DF)
  print(paste("Core_region_size~",i[[1]]))
  print(summary(model)$adj.r.squared)
  var=c(var, paste("Core_region_size~",i[[1]]))
  res=c(res, summary(model)$adj.r.squared)
}
```

```
## [1] "Core_region_size~ Central_compartment_size"
## [1] 0.7668689
## [1] "Core_region_size~ tDNA_region_size"
## [1] 0.1685328
## [1] "Core_region_size~ Chromosome_size"
## [1] 0.6125626
## [1] "Core_region_size~ d_max_rrn_ori"
## [1] 0.8021755
## [1] "Core_region_size~ d_min_rrn_ori"
## [1] 0.4566995
## [1] "Core_region_size~ term_comp"
## [1] 0.1746247
## [1] "Core_region_size~ Clade"
## [1] 0.1700044
## [1] "Core_region_size~ Number_of_rrn_loci"
## [1] 0.08223202
## [1] "Core_region_size~ Number_of_rrn_genes"
## [1] 0.07480597
```

```
result<-data.frame(var, res)
result[which(result$res==max(result$res)),]
```

```
##                                 var               res
## 5 Core_region_size~ d_max_rrn_ori 0.802175451864825
```

```
var<-"NA"
res<-"0"
for (i in predictorlist){
  model<-lm(paste("Core_region_size~d_max_rrn_ori+",i[[1]]), data = DF)
  print(paste("Core_region_size~d_max_rrn_ori+",i[[1]]))
  print(summary(model)$adj.r.squared)
  var=c(var, paste("Core_region_size~d_max_rrn_ori+",i[[1]]))
  res=c(res, summary(model)$adj.r.squared)
}
```

```
## [1] "Core_region_size~d_max_rrn_ori+ Central_compartment_size"
## [1] 0.8114092
## [1] "Core_region_size~d_max_rrn_ori+ tDNA_region_size"
## [1] 0.8161216
## [1] "Core_region_size~d_max_rrn_ori+ Chromosome_size"
## [1] 0.8355459
## [1] "Core_region_size~d_max_rrn_ori+ d_max_rrn_ori"
## [1] 0.8021755
## [1] "Core_region_size~d_max_rrn_ori+ d_min_rrn_ori"
## [1] 0.8114092
## [1] "Core_region_size~d_max_rrn_ori+ term_comp"
## [1] 0.8233131
## [1] "Core_region_size~d_max_rrn_ori+ Clade"
## [1] 0.8166469
## [1] "Core_region_size~d_max_rrn_ori+ Number_of_rrn_loci"
## [1] 0.8060654
## [1] "Core_region_size~d_max_rrn_ori+ Number_of_rrn_genes"
## [1] 0.8069957
```

```
result<-data.frame(var, res)
result[which(result$res==max(result$res)),]
```

```
##                                                var                res
## 4 Core_region_size~d_max_rrn_ori+ Chromosome_size 0.835545871766225
```

```
var<-"NA"
res<-"0"
for (i in predictorlist){
  model<-lm(paste("Core_region_size~d_max_rrn_ori+Chromosome_size+",i[[1]]), data = DF)
  print(paste("Core_region_size~d_max_rrn_ori+Chromosome_size+",i[[1]]))
  print(summary(model)$adj.r.squared)
  var=c(var, paste("Core_region_size~d_max_rrn_ori+Chromosome_size+",i[[1]]))
  res=c(res, summary(model)$adj.r.squared)
}
```

```
## [1] "Core_region_size~d_max_rrn_ori+Chromosome_size+ Central_compartment_size"
## [1] 0.8479681
## [1] "Core_region_size~d_max_rrn_ori+Chromosome_size+ tDNA_region_size"
## [1] 0.8341894
## [1] "Core_region_size~d_max_rrn_ori+Chromosome_size+ Chromosome_size"
## [1] 0.8355459
## [1] "Core_region_size~d_max_rrn_ori+Chromosome_size+ d_max_rrn_ori"
## [1] 0.8355459
## [1] "Core_region_size~d_max_rrn_ori+Chromosome_size+ d_min_rrn_ori"
## [1] 0.8479681
## [1] "Core_region_size~d_max_rrn_ori+Chromosome_size+ term_comp"
## [1] 0.8479681
## [1] "Core_region_size~d_max_rrn_ori+Chromosome_size+ Clade"
## [1] 0.8424337
## [1] "Core_region_size~d_max_rrn_ori+Chromosome_size+ Number_of_rrn_loci"
## [1] 0.8427596
## [1] "Core_region_size~d_max_rrn_ori+Chromosome_size+ Number_of_rrn_genes"
## [1] 0.8440841
```

```
result<-data.frame(var, res)
result[which(result$res==max(result$res)),]
```

```
##                                                                         var
## 2 Core_region_size~d_max_rrn_ori+Chromosome_size+ Central_compartment_size
## 6            Core_region_size~d_max_rrn_ori+Chromosome_size+ d_min_rrn_ori
## 7                Core_region_size~d_max_rrn_ori+Chromosome_size+ term_comp
##                 res
## 2 0.847968132811557
## 6 0.847968132811557
## 7 0.847968132811557
```

```
var<-"NA"
res<-"0"
for (i in predictorlist){
  model<-lm(paste("Core_region_size~d_max_rrn_ori+Chromosome_size+Central_compartment_size+",i[[1]]), data = DF)
  print(paste("Core_region_size~d_max_rrn_ori+Chromosome_size+Central_compartment_size+",i[[1]]))
  print(summary(model)$adj.r.squared)
  var=c(var, paste("Core_region_size~d_max_rrn_ori+Chromosome_size+Central_compartment_size+",i[[1]]))
  res=c(res, summary(model)$adj.r.squared)
}
```

```
## [1] "Core_region_size~d_max_rrn_ori+Chromosome_size+Central_compartment_size+ Central_compartment_size"
## [1] 0.8479681
## [1] "Core_region_size~d_max_rrn_ori+Chromosome_size+Central_compartment_size+ tDNA_region_size"
## [1] 0.8469689
## [1] "Core_region_size~d_max_rrn_ori+Chromosome_size+Central_compartment_size+ Chromosome_size"
## [1] 0.8479681
## [1] "Core_region_size~d_max_rrn_ori+Chromosome_size+Central_compartment_size+ d_max_rrn_ori"
## [1] 0.8479681
## [1] "Core_region_size~d_max_rrn_ori+Chromosome_size+Central_compartment_size+ d_min_rrn_ori"
## [1] 0.8479681
## [1] "Core_region_size~d_max_rrn_ori+Chromosome_size+Central_compartment_size+ term_comp"
## [1] 0.8479681
## [1] "Core_region_size~d_max_rrn_ori+Chromosome_size+Central_compartment_size+ Clade"
## [1] 0.8538154
## [1] "Core_region_size~d_max_rrn_ori+Chromosome_size+Central_compartment_size+ Number_of_rrn_loci"
## [1] 0.8577124
## [1] "Core_region_size~d_max_rrn_ori+Chromosome_size+Central_compartment_size+ Number_of_rrn_genes"
## [1] 0.8593727
```

```
result<-data.frame(var, res)
result[which(result$res==max(result$res)),]
```

```
##                                                                                          var
## 10 Core_region_size~d_max_rrn_ori+Chromosome_size+Central_compartment_size+ Number_of_rrn_genes
##                   res
## 10 0.859372686968973
```

```
var<-"NA"
res<-"0"
for (i in predictorlist){
  model<-lm(paste("Core_region_size~d_max_rrn_ori+Chromosome_size+Central_compartment_size+Number_of_rrn_genes+",i[[1]]), da
ta = DF)
  print(paste("Core_region_size~d_max_rrn_ori+Chromosome_size+Central_compartment_size+Number_of_rrn_genes+",i[[1]]))
  print(summary(model)$adj.r.squared)
  var=c(var, paste("Core_region_size~d_max_rrn_ori+Chromosome_size+Central_compartment_size+Number_of_rrn_genes+",i[[1]]))
  res=c(res, summary(model)$adj.r.squared)
}
```

```
## [1] "Core_region_size~d_max_rrn_ori+Chromosome_size+Central_compartment_size+Number_of_rrn_genes+ Central_compartment_siz
e"
## [1] 0.8593727
## [1] "Core_region_size~d_max_rrn_ori+Chromosome_size+Central_compartment_size+Number_of_rrn_genes+ tDNA_region_size"
## [1] 0.858198
## [1] "Core_region_size~d_max_rrn_ori+Chromosome_size+Central_compartment_size+Number_of_rrn_genes+ Chromosome_size"
## [1] 0.8593727
## [1] "Core_region_size~d_max_rrn_ori+Chromosome_size+Central_compartment_size+Number_of_rrn_genes+ d_max_rrn_ori"
## [1] 0.8593727
## [1] "Core_region_size~d_max_rrn_ori+Chromosome_size+Central_compartment_size+Number_of_rrn_genes+ d_min_rrn_ori"
## [1] 0.8593727
## [1] "Core_region_size~d_max_rrn_ori+Chromosome_size+Central_compartment_size+Number_of_rrn_genes+ term_comp"
## [1] 0.8593727
## [1] "Core_region_size~d_max_rrn_ori+Chromosome_size+Central_compartment_size+Number_of_rrn_genes+ Clade"
## [1] 0.860528
## [1] "Core_region_size~d_max_rrn_ori+Chromosome_size+Central_compartment_size+Number_of_rrn_genes+ Number_of_rrn_loci"
## [1] 0.8589775
## [1] "Core_region_size~d_max_rrn_ori+Chromosome_size+Central_compartment_size+Number_of_rrn_genes+ Number_of_rrn_genes"
## [1] 0.8593727
```

```
result<-data.frame(var, res)
result[which(result$res==max(result$res)),]
```

```
##                                                                                               var
## 8 Core_region_size~d_max_rrn_ori+Chromosome_size+Central_compartment_size+Number_of_rrn_genes+ Clade
##                  res
## 8 0.860528011977805
```

```
var<-"NA"
res<-"0"
for (i in predictorlist){
  model<-lm(paste("Core_region_size~d_max_rrn_ori+Chromosome_size+Central_compartment_size+Number_of_rrn_genes+Clade+",i
[[1]]), data = DF)
  print(paste("Core_region_size~d_max_rrn_ori+Chromosome_size+Central_compartment_size+Number_of_rrn_genes+Clade+",i[[1]]))
  print(summary(model)$adj.r.squared)
  var=c(var, paste("Core_region_size~d_max_rrn_ori+Chromosome_size+Central_compartment_size+Number_of_rrn_genes+Clade+",i
[[1]]))
  res=c(res, summary(model)$adj.r.squared)
}
```

```
## [1] "Core_region_size~d_max_rrn_ori+Chromosome_size+Central_compartment_size+Number_of_rrn_genes+Clade+ Central_compartme
nt_size"
## [1] 0.860528
## [1] "Core_region_size~d_max_rrn_ori+Chromosome_size+Central_compartment_size+Number_of_rrn_genes+Clade+ tDNA_region_size"
## [1] 0.8593395
## [1] "Core_region_size~d_max_rrn_ori+Chromosome_size+Central_compartment_size+Number_of_rrn_genes+Clade+ Chromosome_size"
## [1] 0.860528
## [1] "Core_region_size~d_max_rrn_ori+Chromosome_size+Central_compartment_size+Number_of_rrn_genes+Clade+ d_max_rrn_ori"
## [1] 0.860528
## [1] "Core_region_size~d_max_rrn_ori+Chromosome_size+Central_compartment_size+Number_of_rrn_genes+Clade+ d_min_rrn_ori"
## [1] 0.860528
## [1] "Core_region_size~d_max_rrn_ori+Chromosome_size+Central_compartment_size+Number_of_rrn_genes+Clade+ term_comp"
## [1] 0.860528
## [1] "Core_region_size~d_max_rrn_ori+Chromosome_size+Central_compartment_size+Number_of_rrn_genes+Clade+ Clade"
## [1] 0.860528
## [1] "Core_region_size~d_max_rrn_ori+Chromosome_size+Central_compartment_size+Number_of_rrn_genes+Clade+ Number_of_rrn_loc
i"
## [1] 0.8601765
## [1] "Core_region_size~d_max_rrn_ori+Chromosome_size+Central_compartment_size+Number_of_rrn_genes+Clade+ Number_of_rrn_gen
es"
## [1] 0.860528
```

```
result<-data.frame(var, res)
result[which(result$res==max(result$res)),]
```

```
##
var
## 2  Core_region_size~d_max_rrn_ori+Chromosome_size+Central_compartment_size+Number_of_rrn_genes+Clade+ Central_compartment
_size
## 4           Core_region_size~d_max_rrn_ori+Chromosome_size+Central_compartment_size+Number_of_rrn_genes+Clade+ Chromosome
_size
## 5             Core_region_size~d_max_rrn_ori+Chromosome_size+Central_compartment_size+Number_of_rrn_genes+Clade+ d_max_rr
n_ori
## 6             Core_region_size~d_max_rrn_ori+Chromosome_size+Central_compartment_size+Number_of_rrn_genes+Clade+ d_min_rr
n_ori
## 7                 Core_region_size~d_max_rrn_ori+Chromosome_size+Central_compartment_size+Number_of_rrn_genes+Clade+ term
_comp
## 8                     Core_region_size~d_max_rrn_ori+Chromosome_size+Central_compartment_size+Number_of_rrn_genes+Clade+
Clade
## 10      Core_region_size~d_max_rrn_ori+Chromosome_size+Central_compartment_size+Number_of_rrn_genes+Clade+ Number_of_rrn_
genes
##                   res
## 2  0.860528011977805
## 4  0.860528011977805
## 5  0.860528011977805
## 6  0.860528011977805
## 7  0.860528011977805
## 8  0.860528011977805
## 10 0.860528011977805
```

```
LM0<-lm(Core_region_size~1, DF)
LM1<-lm(Core_region_size~d_max_rrn_ori, DF)
LM2<-lm(Core_region_size~d_max_rrn_ori+Chromosome_size, DF)
LM3<-lm(Core_region_size~d_max_rrn_ori+Chromosome_size+Central_compartment_size, DF)
LM4<-lm(Core_region_size~d_max_rrn_ori+Chromosome_size+Central_compartment_size+Number_of_rrn_genes, DF)
LM5<-lm(Core_region_size~d_max_rrn_ori+Chromosome_size+Central_compartment_size+Number_of_rrn_genes+Clade, DF)

AIC(LM0, LM1, LM2, LM3, LM4,LM5)
```

```
##     df      AIC
## LM0  2 3766.243
## LM1  3 3564.684
## LM2  4 3542.570
## LM3  5 3533.723
## LM4  6 3524.939
## LM5  8 3525.807
```

```
#Based on the AIC, LM4 is the best model
plot(LM4,cex.lab=1.5,cex.main=1.5,pch=16)
```

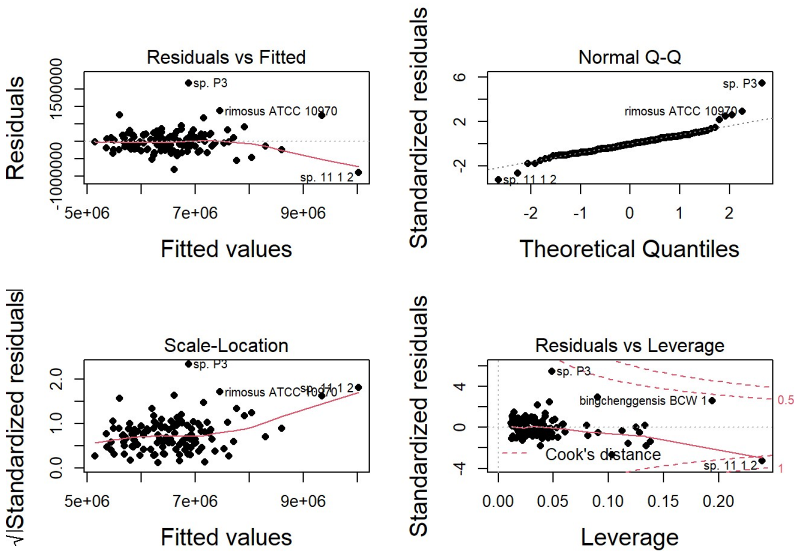


```
#The final model selected correspond to LM4 (lowest AIC and all variable statistically significant, no strain showing a leverage effect).
summary(LM4)
```

```
##
## Call:
## lm(formula = Core_region_size ~ d_max_rrn_ori + Chromosome_size +
##     Central_compartment_size + Number_of_rrn_genes, data = DF)
##
## Residuals:
##     Min      1Q  Median      3Q     Max
## -892177 -174499  -22613  164547 1663768
##
## Coefficients:
##                            Estimate Std. Error t value Pr(>|t|)
## (Intercept)               1.016e+06  4.974e+05   2.042 0.043332 *
## d_max_rrn_ori             7.262e-01  2.603e-01   2.790 0.006136 **
## Chromosome_size           2.398e-01  3.970e-02   6.041 1.77e-08 ***
## Central_compartment_size  5.522e-01  1.468e-01   3.762 0.000262 ***
## Number_of_rrn_genes      -6.694e+04  2.036e+04  -3.288 0.001323 **
## ---
## Signif. codes:  0 '***' 0.001 '**' 0.01 '*' 0.05 '.' 0.1 ' ' 1
##
## Residual standard error: 312800 on 120 degrees of freedom
## Multiple R-squared:  0.8639, Adjusted R-squared:  0.8594
## F-statistic: 190.4 on 4 and 120 DF,  p-value: < 2.2e-16
```

Modeling_Backward approach

```
DF<-ALL[,c("id", "Core_region_size","Central_compartment_size", "tDNA_region_size", "Chromosome_size", "d_max_rrn_ori", "d_min_rrn_ori", "term_comp", "Clade","Number_of_rrn_loci","Number_of_rrn_genes")]
row.names(DF)<-DF$id
```

```
## Warning: Setting row names on a tibble is deprecated.
```

```
predictorlist<-list("Central_compartment_size", "tDNA_region_size", "Chromosome_size", "d_max_rrn_ori", "d_min_rrn_ori","term_comp","Clade","Number_of_rrn_loci","Number_of_rrn_genes")

BLM0<-lm(Core_region_size~1, DF)
BLM1<-lm(Core_region_size~d_max_rrn_ori+Chromosome_size+Central_compartment_size+d_min_rrn_ori+tDNA_region_size+term_comp+Clade+Number_of_rrn_loci+Number_of_rrn_genes, DF)
summary(BLM1)
```

```
##
## Call:
## lm(formula = Core_region_size ~ d_max_rrn_ori + Chromosome_size +
##     Central_compartment_size + d_min_rrn_ori + tDNA_region_size +
##     term_comp + Clade + Number_of_rrn_loci + Number_of_rrn_genes,
##     data = DF)
##
## Residuals:
##     Min      1Q  Median      3Q     Max
## -859689 -179835  -24247  123533 1645011
##
## Coefficients: (2 not defined because of singularities)
##                           Estimate Std. Error t value Pr(>|t|)
## (Intercept)              1.448e+06  5.812e+05   2.491  0.01413 *
## d_max_rrn_ori            3.347e-01  2.885e-01   1.160  0.24831
## Chromosome_size          2.680e-01  5.739e-02   4.669 8.06e-06 ***
## Central_compartment_size  7.026e-01  1.575e-01   4.461 1.87e-05 ***
## d_min_rrn_ori                   NA         NA      NA       NA
## tDNA_region_size        -1.462e-02  3.908e-02  -0.374  0.70904
## term_comp                       NA         NA      NA       NA
## CladeClade_2            -6.086e+04  7.242e+04  -0.840  0.40237
## CladeGroup_O             8.502e+04  1.077e+05   0.789  0.43156
## Number_of_rrn_loci       3.193e+05  2.112e+05   1.512  0.13326
## Number_of_rrn_genes     -1.822e+05  6.424e+04  -2.836  0.00538 **
## ---
## Signif. codes:  0 '***' 0.001 '**' 0.01 '*' 0.05 '.' 0.1 ' ' 1
##
## Residual standard error: 334400 on 118 degrees of freedom
## Multiple R-squared:  0.8471, Adjusted R-squared:  0.8367
## F-statistic: 81.72 on 8 and 118 DF,  p-value: < 2.2e-16
```

```
BLM2<-lm(Core_region_size~d_max_rrn_ori+Chromosome_size+Central_compartment_size+tDNA_region_size+term_comp+Clade+Number_of_
rrn_loci+Number_of_rrn_genes, DF)
summary(BLM2)
```

```
##
## Call:
## lm(formula = Core_region_size ~ d_max_rrn_ori + Chromosome_size +
##     Central_compartment_size + tDNA_region_size + term_comp +
##     Clade + Number_of_rrn_loci + Number_of_rrn_genes, data = DF)
##
## Residuals:
##     Min      1Q  Median      3Q     Max
## -859689 -179835  -24247  123533 1645011
##
## Coefficients: (1 not defined because of singularities)
##                           Estimate Std. Error t value Pr(>|t|)
## (Intercept)              1.448e+06  5.812e+05   2.491  0.01413 *
## d_max_rrn_ori            3.347e-01  2.885e-01   1.160  0.24831
## Chromosome_size          2.680e-01  5.739e-02   4.669 8.06e-06 ***
## Central_compartment_size  7.026e-01  1.575e-01   4.461 1.87e-05 ***
## tDNA_region_size        -1.462e-02  3.908e-02  -0.374  0.70904
## term_comp                       NA         NA      NA       NA
## CladeClade_2            -6.086e+04  7.242e+04  -0.840  0.40237
## CladeGroup_O             8.502e+04  1.077e+05   0.789  0.43156
## Number_of_rrn_loci       3.193e+05  2.112e+05   1.512  0.13326
## Number_of_rrn_genes     -1.822e+05  6.424e+04  -2.836  0.00538 **
## ---
## Signif. codes:  0 '***' 0.001 '**' 0.01 '*' 0.05 '.' 0.1 ' ' 1
##
## Residual standard error: 334400 on 118 degrees of freedom
## Multiple R-squared:  0.8471, Adjusted R-squared:  0.8367
## F-statistic: 81.72 on 8 and 118 DF,  p-value: < 2.2e-16
```

```
BLM3<-lm(Core_region_size~d_max_rrn_ori+Chromosome_size+Central_compartment_size+tDNA_region_size+Clade+Number_of_rrn_loci+N
umber_of_rrn_genes, DF)
summary(BLM3)
```

```
##
## Call:
## lm(formula = Core_region_size ~ d_max_rrn_ori + Chromosome_size +
##     Central_compartment_size + tDNA_region_size + Clade + Number_of_rrn_loci +
##     Number_of_rrn_genes, data = DF)
##
## Residuals:
##     Min      1Q  Median      3Q     Max
## -859689 -179835  -24247  123533 1645011
##
## Coefficients:
##                            Estimate Std. Error t value Pr(>|t|)
## (Intercept)               1.448e+06  5.812e+05   2.491  0.01413 *
## d_max_rrn_ori             3.347e-01  2.885e-01   1.160  0.24831
## Chromosome_size           2.680e-01  5.739e-02   4.669 8.06e-06 ***
## Central_compartment_size  7.026e-01  1.575e-01   4.461 1.87e-05 ***
## tDNA_region_size         -1.462e-02  3.908e-02  -0.374  0.70904
## CladeClade_2             -6.086e+04  7.242e+04  -0.840  0.40237
## CladeGroup_O              8.502e+04  1.077e+05   0.789  0.43156
## Number_of_rrn_loci        3.193e+05  2.112e+05   1.512  0.13326
## Number_of_rrn_genes      -1.822e+05  6.424e+04  -2.836  0.00538 **
## ---
## Signif. codes:  0 '***' 0.001 '**' 0.01 '*' 0.05 '.' 0.1 ' ' 1
##
## Residual standard error: 334400 on 118 degrees of freedom
## Multiple R-squared:  0.8471, Adjusted R-squared:  0.8367
## F-statistic: 81.72 on 8 and 118 DF,  p-value: < 2.2e-16
```

```
BLM4<-lm(Core_region_size~d_max_rrn_ori+Chromosome_size+Central_compartment_size+tDNA_region_size+Number_of_rrn_genes+Number
_of_rrn_loci, DF)
summary(BLM4)
```

```
##
## Call:
## lm(formula = Core_region_size ~ d_max_rrn_ori + Chromosome_size +
##     Central_compartment_size + tDNA_region_size + Number_of_rrn_genes +
##     Number_of_rrn_loci, data = DF)
##
## Residuals:
##     Min      1Q  Median      3Q     Max
## -849931 -204410  -10935  130499 1649976
##
## Coefficients:
##                            Estimate Std. Error t value Pr(>|t|)
## (Intercept)               1.182e+06  5.117e+05   2.311  0.02255 *
## d_max_rrn_ori             4.053e-01  2.757e-01   1.470  0.14411
## Chromosome_size           2.797e-01  5.590e-02   5.003 1.95e-06 ***
## Central_compartment_size  6.805e-01  1.551e-01   4.387 2.49e-05 ***
## tDNA_region_size         -1.759e-02  3.897e-02  -0.451  0.65253
## Number_of_rrn_genes      -1.846e+05  6.418e+04  -2.876  0.00477 **
## Number_of_rrn_loci        3.395e+05  2.075e+05   1.637  0.10434
## ---
## Signif. codes:  0 '***' 0.001 '**' 0.01 '*' 0.05 '.' 0.1 ' ' 1
##
## Residual standard error: 334300 on 120 degrees of freedom
## Multiple R-squared:  0.8447, Adjusted R-squared:  0.8369
## F-statistic: 108.8 on 6 and 120 DF,  p-value: < 2.2e-16
```

```
BLM5<-lm(Core_region_size~d_max_rrn_ori+Chromosome_size+Central_compartment_size+Number_of_rrn_genes+Number_of_rrn_loci, DF)
summary(BLM5)
```

```
##
## Call:
## lm(formula = Core_region_size ~ d_max_rrn_ori + Chromosome_size +
##     Central_compartment_size + Number_of_rrn_genes + Number_of_rrn_loci,
##     data = DF)
##
## Residuals:
##     Min      1Q  Median      3Q     Max
## -867636 -198283  -20376  136742 1624724
##
## Coefficients:
##                            Estimate Std. Error t value Pr(>|t|)
## (Intercept)               1.181e+06  5.100e+05   2.315  0.02231 *
## d_max_rrn_ori             4.345e-01  2.671e-01   1.627  0.10636
## Chromosome_size           2.630e-01  4.185e-02   6.285 5.39e-09 ***
## Central_compartment_size  6.721e-01  1.535e-01   4.378 2.56e-05 ***
## Number_of_rrn_genes      -1.849e+05  6.396e+04  -2.892  0.00455 **
## Number_of_rrn_loci        3.375e+05  2.067e+05   1.633  0.10516
## ---
## Signif. codes:  0 '***' 0.001 '**' 0.01 '*' 0.05 '.' 0.1 ' ' 1
##
## Residual standard error: 333200 on 121 degrees of freedom
## Multiple R-squared:  0.8444, Adjusted R-squared:  0.838
## F-statistic: 131.3 on 5 and 121 DF,  p-value: < 2.2e-16
```

```
BLM6<-lm(Core_region_size~d_max_rrn_ori+Chromosome_size+Central_compartment_size+Number_of_rrn_genes, DF)
summary(BLM6)
```

```
##
## Call:
## lm(formula = Core_region_size ~ d_max_rrn_ori + Chromosome_size +
##     Central_compartment_size + Number_of_rrn_genes, data = DF)
##
## Residuals:
##     Min      1Q  Median      3Q     Max
## -905829 -191258  -22965  138248 1598142
##
## Coefficients:
##                            Estimate Std. Error t value Pr(>|t|)
## (Intercept)               1.541e+06  4.628e+05   3.330  0.00115 **
## d_max_rrn_ori             3.954e-01  2.678e-01   1.477  0.14238
## Chromosome_size           2.667e-01  4.207e-02   6.340 4.05e-09 ***
## Central_compartment_size  6.657e-01  1.545e-01   4.309 3.34e-05 ***
## Number_of_rrn_genes      -8.518e+04  1.901e+04  -4.480 1.69e-05 ***
## ---
## Signif. codes:  0 '***' 0.001 '**' 0.01 '*' 0.05 '.' 0.1 ' ' 1
##
## Residual standard error: 335400 on 122 degrees of freedom
## Multiple R-squared:  0.841,  Adjusted R-squared:  0.8358
## F-statistic: 161.3 on 4 and 122 DF,  p-value: < 2.2e-16
```

```
BLM7<-lm(Core_region_size~Chromosome_size+Central_compartment_size+Number_of_rrn_genes, DF)
summary(BLM7)
```

```
##
## Call:
## lm(formula = Core_region_size ~ Chromosome_size + Central_compartment_size +
##     Number_of_rrn_genes, data = DF)
##
## Residuals:
##     Min      1Q  Median      3Q     Max
## -970167 -209256  -16517  141661 1560863
##
## Coefficients:
##                            Estimate Std. Error t value Pr(>|t|)
## (Intercept)               1.586e+06  4.640e+05   3.417 0.000857 ***
## Chromosome_size           2.910e-01  3.891e-02   7.478 1.23e-11 ***
## Central_compartment_size  8.675e-01  7.244e-02  11.976  < 2e-16 ***
## Number_of_rrn_genes      -9.007e+04  1.881e+04  -4.789 4.73e-06 ***
## ---
## Signif. codes:  0 '***' 0.001 '**' 0.01 '*' 0.05 '.' 0.1 ' ' 1
##
## Residual standard error: 337000 on 123 degrees of freedom
## Multiple R-squared:  0.8381, Adjusted R-squared:  0.8342
## F-statistic: 212.3 on 3 and 123 DF,  p-value: < 2.2e-16
```

```
AIC(BLM0, BLM1, BLM2, BLM3, BLM4,BLM5, BLM6, BLM7)
```

```
##      df      AIC
## BLM0  2 3824.509
## BLM1 10 3602.008
## BLM2 10 3602.008
## BLM3 10 3602.008
## BLM4  8 3600.005
## BLM5  7 3598.221
## BLM6  6 3598.988
## BLM7  5 3599.237
```

```
#The BLM5 is the best model based on AIC criteria. That is the same model than LM4 (obtained with the forward approach - cf. previous part). Taken into account the leverage effect previously mentioned with Streptomyces asterosporus DSM 41452 and Streptomyces tirandamycinicus HNM0039, a new backward approach was conducted without these two strains.

DF<-ALL_wo_astero_wo_tiran[,c("id", "Core_region_size","Central_compartment_size", "tDNA_region_size", "Chromosome_size", "d_max_rrn_ori", "d_min_rrn_ori", "term_comp", "Clade","Number_of_rrn_loci","Number_of_rrn_genes")]
row.names(DF)<-DF$id
```

```
## Warning: Setting row names on a tibble is deprecated.
```

```
predictorlist<-list("Central_compartment_size", "tDNA_region_size", "Chromosome_size", "d_max_rrn_ori", "d_min_rrn_ori","term_comp","Clade","Number_of_rrn_loci","Number_of_rrn_genes")

BLM0<-lm(Core_region_size~1, DF)
BLM1<-lm(Core_region_size~d_max_rrn_ori+Chromosome_size+Central_compartment_size+d_min_rrn_ori+tDNA_region_size+term_comp+Clade+Number_of_rrn_loci+Number_of_rrn_genes, DF)
summary(BLM1)
```

```
##
## Call:
## lm(formula = Core_region_size ~ d_max_rrn_ori + Chromosome_size +
##     Central_compartment_size + d_min_rrn_ori + tDNA_region_size +
##     term_comp + Clade + Number_of_rrn_loci + Number_of_rrn_genes,
##     data = DF)
##
## Residuals:
##     Min      1Q  Median      3Q     Max
## -876177 -160514  -18077  135123 1656259
##
## Coefficients: (2 not defined because of singularities)
##                            Estimate Std. Error t value Pr(>|t|)
## (Intercept)               1.105e+06  5.793e+05   1.908 0.058909 .
## d_max_rrn_ori             7.228e-01  2.853e-01   2.533 0.012631 *
## Chromosome_size           2.205e-01  5.487e-02   4.019 0.000104 ***
## Central_compartment_size  5.555e-01  1.514e-01   3.669 0.000369 ***
## d_min_rrn_ori                    NA         NA      NA       NA
## tDNA_region_size          3.035e-03  3.691e-02   0.082 0.934615
## term_comp                        NA         NA      NA       NA
## CladeClade_2             -1.090e+05  6.889e+04  -1.582 0.116298
## CladeGroup_O              1.785e+04  1.039e+05   0.172 0.863848
## Number_of_rrn_loci        2.557e+05  3.053e+05   0.837 0.404089
## Number_of_rrn_genes      -1.481e+05  1.056e+05  -1.402 0.163487
## ---
## Signif. codes:  0 '***' 0.001 '**' 0.01 '*' 0.05 '.' 0.1 ' ' 1
##
## Residual standard error: 313200 on 116 degrees of freedom
## Multiple R-squared:  0.8681, Adjusted R-squared:  0.859
## F-statistic: 95.41 on 8 and 116 DF,  p-value: < 2.2e-16
```

```
BLM2<-lm(Core_region_size~d_max_rrn_ori+Chromosome_size+Central_compartment_size+tDNA_region_size+Clade+Number_of_rrn_loci+N
umber_of_rrn_genes, DF)
summary(BLM2)
```

```
##
## Call:
## lm(formula = Core_region_size ~ d_max_rrn_ori + Chromosome_size +
##     Central_compartment_size + tDNA_region_size + Clade + Number_of_rrn_loci +
##     Number_of_rrn_genes, data = DF)
##
## Residuals:
##     Min      1Q  Median      3Q     Max
## -876177 -160514  -18077  135123 1656259
##
## Coefficients:
##                            Estimate Std. Error t value Pr(>|t|)
## (Intercept)               1.105e+06  5.793e+05   1.908 0.058909 .
## d_max_rrn_ori             7.228e-01  2.853e-01   2.533 0.012631 *
## Chromosome_size           2.205e-01  5.487e-02   4.019 0.000104 ***
## Central_compartment_size  5.555e-01  1.514e-01   3.669 0.000369 ***
## tDNA_region_size          3.035e-03  3.691e-02   0.082 0.934615
## CladeClade_2             -1.090e+05  6.889e+04  -1.582 0.116298
## CladeGroup_O              1.785e+04  1.039e+05   0.172 0.863848
## Number_of_rrn_loci        2.557e+05  3.053e+05   0.837 0.404089
## Number_of_rrn_genes      -1.481e+05  1.056e+05  -1.402 0.163487
## ---
## Signif. codes:  0 '***' 0.001 '**' 0.01 '*' 0.05 '.' 0.1 ' ' 1
##
## Residual standard error: 313200 on 116 degrees of freedom
## Multiple R-squared:  0.8681, Adjusted R-squared:  0.859
## F-statistic: 95.41 on 8 and 116 DF,  p-value: < 2.2e-16
```

```
BLM3<-lm(Core_region_size~d_max_rrn_ori+Chromosome_size+Central_compartment_size+tDNA_region_size+Clade+Number_of_rrn_loci+N
umber_of_rrn_genes, DF)
summary(BLM3)
```

```
##
## Call:
## lm(formula = Core_region_size ~ d_max_rrn_ori + Chromosome_size +
##     Central_compartment_size + tDNA_region_size + Clade + Number_of_rrn_loci +
##     Number_of_rrn_genes, data = DF)
##
## Residuals:
##     Min      1Q  Median      3Q     Max
## -876177 -160514  -18077  135123 1656259
##
## Coefficients:
##                           Estimate Std. Error t value Pr(>|t|)
## (Intercept)              1.105e+06  5.793e+05   1.908 0.058909 .
## d_max_rrn_ori            7.228e-01  2.853e-01   2.533 0.012631 *
## Chromosome_size          2.205e-01  5.487e-02   4.019 0.000104 ***
## Central_compartment_size  5.555e-01  1.514e-01   3.669 0.000369 ***
## tDNA_region_size         3.035e-03  3.691e-02   0.082 0.934615
## CladeClade_2            -1.090e+05  6.889e+04  -1.582 0.116298
## CladeGroup_O             1.785e+04  1.039e+05   0.172 0.863848
## Number_of_rrn_loci       2.557e+05  3.053e+05   0.837 0.404089
## Number_of_rrn_genes     -1.481e+05  1.056e+05  -1.402 0.163487
## ---
## Signif. codes:  0 '***' 0.001 '**' 0.01 '*' 0.05 '.' 0.1 ' ' 1
##
## Residual standard error: 313200 on 116 degrees of freedom
## Multiple R-squared:  0.8681, Adjusted R-squared:  0.859
## F-statistic: 95.41 on 8 and 116 DF,  p-value: < 2.2e-16
```

```
BLM4<-lm(Core_region_size~d_max_rrn_ori+Chromosome_size+Central_compartment_size+Number_of_rrn_loci+Number_of_rrn_genes, DF)
summary(BLM4)
```

```
##
## Call:
## lm(formula = Core_region_size ~ d_max_rrn_ori + Chromosome_size +
##     Central_compartment_size + Number_of_rrn_loci + Number_of_rrn_genes,
##     data = DF)
##
## Residuals:
##     Min      1Q  Median      3Q     Max
## -904473 -166314  -21170  165352 1667008
##
## Coefficients:
##                           Estimate Std. Error t value Pr(>|t|)
## (Intercept)              9.921e+05  4.989e+05   1.988 0.049063 *
## d_max_rrn_ori            7.316e-01  2.607e-01   2.806 0.005869 **
## Chromosome_size          2.403e-01  3.976e-02   6.042 1.79e-08 ***
## Central_compartment_size  5.542e-01  1.470e-01   3.770 0.000255 ***
## Number_of_rrn_loci       2.485e+05  3.050e+05   0.815 0.416868
## Number_of_rrn_genes     -1.502e+05  1.043e+05  -1.441 0.152208
## ---
## Signif. codes:  0 '***' 0.001 '**' 0.01 '*' 0.05 '.' 0.1 ' ' 1
##
## Residual standard error: 313200 on 119 degrees of freedom
## Multiple R-squared:  0.8647, Adjusted R-squared:  0.859
## F-statistic: 152.1 on 5 and 119 DF,  p-value: < 2.2e-16
```

```
BLM5<-lm(Core_region_size~d_max_rrn_ori+Chromosome_size+Central_compartment_size+Number_of_rrn_genes, DF)
summary(BLM5)
```

```
##
## Call:
## lm(formula = Core_region_size ~ d_max_rrn_ori + Chromosome_size +
##     Central_compartment_size + Number_of_rrn_genes, data = DF)
##
## Residuals:
##     Min      1Q  Median      3Q     Max
## -892177 -174499  -22613  164547 1663768
##
## Coefficients:
##                           Estimate Std. Error t value Pr(>|t|)
## (Intercept)              1.016e+06  4.974e+05   2.042 0.043332 *
## d_max_rrn_ori            7.262e-01  2.603e-01   2.790 0.006136 **
## Chromosome_size          2.398e-01  3.970e-02   6.041 1.77e-08 ***
## Central_compartment_size  5.522e-01  1.468e-01   3.762 0.000262 ***
## Number_of_rrn_genes     -6.694e+04  2.036e+04  -3.288 0.001323 **
## ---
## Signif. codes:  0 '***' 0.001 '**' 0.01 '*' 0.05 '.' 0.1 ' ' 1
##
## Residual standard error: 312800 on 120 degrees of freedom
## Multiple R-squared:  0.8639, Adjusted R-squared:  0.8594
## F-statistic: 190.4 on 4 and 120 DF,  p-value: < 2.2e-16
```

```
AIC(BLM0, BLM1, BLM2, BLM3, BLM4,BLM5)
```

```
##      df      AIC
## BLM0  2 3766.243
## BLM1 10 3529.050
## BLM2 10 3529.050
## BLM3 10 3529.050
## BLM4  7 3526.243
## BLM5  6 3524.939
```

```
#The best model (based on AIC criteria) is BLM5.
```

Final model analysis & Predictions

```
#Final model with all strains
DF<-ALL[,c("id", "Core_region_size","Central_compartment_size", "Chromosome_size", "d_max_rrn_ori","Number_of_rrn_genes")]
row.names(DF)<-DF$id
```

```
## Warning: Setting row names on a tibble is deprecated.
```

```
Model0<-lm(Core_region_size~1, DF)
ModelFinal1<-lm(Core_region_size~d_max_rrn_ori+Chromosome_size+Central_compartment_size+Number_of_rrn_genes, DF)
summary(ModelFinal1)
```

```
##
## Call:
## lm(formula = Core_region_size ~ d_max_rrn_ori + Chromosome_size +
##     Central_compartment_size + Number_of_rrn_genes, data = DF)
##
## Residuals:
##     Min      1Q  Median      3Q     Max
## -905829 -191258  -22965  138248 1598142
##
## Coefficients:
##                            Estimate Std. Error t value Pr(>|t|)
## (Intercept)               1.541e+06  4.628e+05   3.330  0.00115 **
## d_max_rrn_ori             3.954e-01  2.678e-01   1.477  0.14238
## Chromosome_size           2.667e-01  4.207e-02   6.340 4.05e-09 ***
## Central_compartment_size  6.657e-01  1.545e-01   4.309 3.34e-05 ***
## Number_of_rrn_genes      -8.518e+04  1.901e+04  -4.480 1.69e-05 ***
## ---
## Signif. codes:  0 '***' 0.001 '**' 0.01 '*' 0.05 '.' 0.1 ' ' 1
##
## Residual standard error: 335400 on 122 degrees of freedom
## Multiple R-squared:  0.841,  Adjusted R-squared:  0.8358
## F-statistic: 161.3 on 4 and 122 DF,  p-value: < 2.2e-16
```

```
par(mfrow=c(2,2))
plot(ModelFinal1,cex.lab=1.5,cex.main=1.5,pch=16)
```

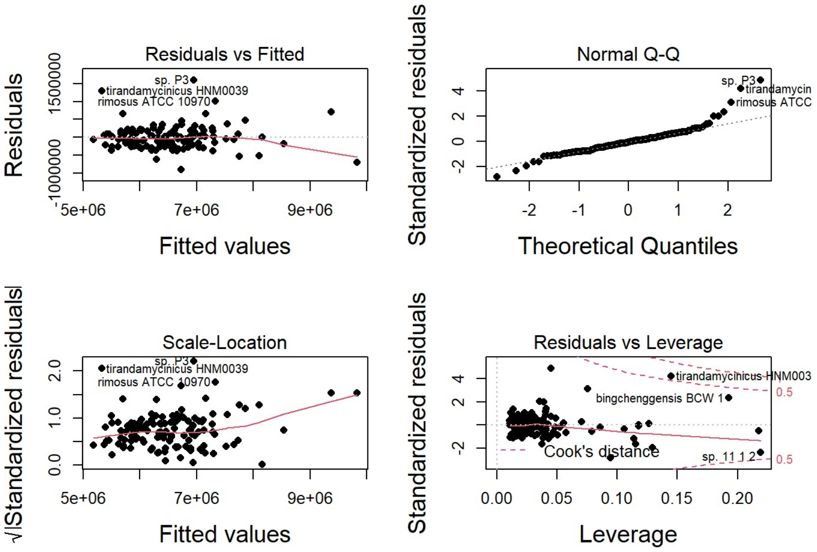


```
anova(ModelFinal1, Model0, test= "Chisq")
```

```
## Analysis of Variance Table
##
## Model 1: Core_region_size ~ d_max_rrn_ori + Chromosome_size + Central_compartment_size +
##     Number_of_rrn_genes
## Model 2: Core_region_size ~ 1
##   Res.Df        RSS Df   Sum of Sq  Pr(>Chi)
## 1    122 1.3726e+13
## 2    126 8.6318e+13 -4 -7.2592e+13 < 2.2e-16 ***
## ---
## Signif. codes:  0 '***' 0.001 '**' 0.01 '*' 0.05 '.' 0.1 ' ' 1
```

```
anova(ModelFinal1)
```

```
## Analysis of Variance Table
##
## Response: Core_region_size
##                           Df     Sum Sq    Mean Sq F value    Pr(>F)
## d_max_rrn_ori              1 6.4856e+13 6.4856e+13 576.459 < 2.2e-16 ***
## Chromosome_size            1 3.8675e+12 3.8675e+12  34.376 3.970e-08 ***
## Central_compartment_size   1 1.6100e+12 1.6100e+12  14.310 0.0002416 ***
## Number_of_rrn_genes        1 2.2584e+12 2.2584e+12  20.073 1.692e-05 ***
## Residuals                122 1.3726e+13 1.1251e+11
## ---
## Signif. codes:  0 '***' 0.001 '**' 0.01 '*' 0.05 '.' 0.1 ' ' 1
```

```
Anova(ModelFinal1)
```

```
## Anova Table (Type II tests)
##
## Response: Core_region_size
##                              Sum Sq  Df F value    Pr(>F)
## d_max_rrn_ori            2.4528e+11   1  2.1802    0.1424
## Chromosome_size          4.5216e+12   1 40.1893 4.049e-09 ***
## Central_compartment_size 2.0892e+12   1 18.5698 3.337e-05 ***
## Number_of_rrn_genes      2.2584e+12   1 20.0729 1.692e-05 ***
## Residuals                1.3726e+13 122
## ---
## Signif. codes:  0 '***' 0.001 '**' 0.01 '*' 0.05 '.' 0.1 ' ' 1
```

```
#We notice that in this model, Streptomyces asterosporus DSM 41452 and Streptomyces tirandamycinicus HNM0039 do not have a s
trong leverage effect.

#Final model without Streptomyces asterosporus DSM 41452 and Streptomyces tirandamycinicus HNM0039
DF<-ALL_wo_astero_wo_tiran[,c("id", "Core_region_size","Central_compartment_size", "Chromosome_size", "d_max_rrn_ori","Numbe
r_of_rrn_genes")]
row.names(DF)<-DF$id
```

```
## Warning: Setting row names on a tibble is deprecated.
```

```
Model0<-lm(Core_region_size~1, DF)
ModelFinal2<-lm(Core_region_size~d_max_rrn_ori+Chromosome_size+Central_compartment_size+Number_of_rrn_genes, DF)
summary(ModelFinal2)
```

```
##
## Call:
## lm(formula = Core_region_size ~ d_max_rrn_ori + Chromosome_size +
##     Central_compartment_size + Number_of_rrn_genes, data = DF)
##
## Residuals:
##     Min      1Q  Median      3Q     Max
## -892177 -174499  -22613  164547 1663768
##
## Coefficients:
##                            Estimate Std. Error t value Pr(>|t|)
## (Intercept)               1.016e+06  4.974e+05   2.042 0.043332 *
## d_max_rrn_ori             7.262e-01  2.603e-01   2.790 0.006136 **
## Chromosome_size           2.398e-01  3.970e-02   6.041 1.77e-08 ***
## Central_compartment_size  5.522e-01  1.468e-01   3.762 0.000262 ***
## Number_of_rrn_genes      -6.694e+04  2.036e+04  -3.288 0.001323 **
## ---
## Signif. codes:  0 '***' 0.001 '**' 0.01 '*' 0.05 '.' 0.1 ' ' 1
##
## Residual standard error: 312800 on 120 degrees of freedom
## Multiple R-squared:  0.8639, Adjusted R-squared:  0.8594
## F-statistic: 190.4 on 4 and 120 DF,  p-value: < 2.2e-16
```

```
par(mfrow=c(2,2))
plot(ModelFinal2,cex.lab=1.5,cex.main=1.5,pch=16)
```

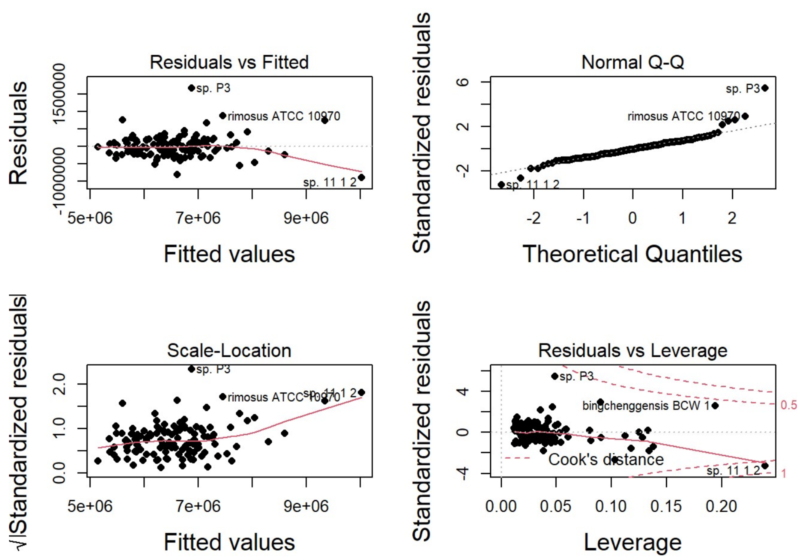


```
anova(ModelFinal2, Model0, test= "Chisq")
```

```
## Analysis of Variance Table
##
## Model 1: Core_region_size ~ d_max_rrn_ori + Chromosome_size + Central_compartment_size +
##     Number_of_rrn_genes
## Model 2: Core_region_size ~ 1
##   Res.Df        RSS Df  Sum of Sq  Pr(>Chi)
## 1    120 1.1739e+13
## 2    124 8.6259e+13 -4 -7.452e+13 < 2.2e-16 ***
## ---
## Signif. codes:  0 '***' 0.001 '**' 0.01 '*' 0.05 '.' 0.1 ' ' 1
```

```
anova(ModelFinal2)
```

```
## Analysis of Variance Table
##
## Response: Core_region_size
##                           Df     Sum Sq    Mean Sq F value    Pr(>F)
## d_max_rrn_ori              1 6.9333e+13 6.9333e+13 708.736 < 2.2e-16 ***
## Chromosome_size            1 2.9697e+12 2.9697e+12  30.357 2.086e-07 ***
## Central_compartment_size   1 1.1600e+12 1.1600e+12  11.858 0.0007914 ***
## Number_of_rrn_genes        1 1.0578e+12 1.0578e+12  10.813 0.0013229 **
## Residuals                120 1.1739e+13 9.7826e+10
## ---
## Signif. codes:  0 '***' 0.001 '**' 0.01 '*' 0.05 '.' 0.1 ' ' 1
```

```
Anova(ModelFinal2)
```

```
## Anova Table (Type II tests)
##
## Response: Core_region_size
##                              Sum Sq  Df F value    Pr(>F)
## d_max_rrn_ori            7.6139e+11   1  7.7831 0.0061361 **
## Chromosome_size          3.5697e+12   1 36.4909 1.771e-08 ***
## Central_compartment_size 1.3847e+12   1 14.1548 0.0002619 ***
## Number_of_rrn_genes      1.0578e+12   1 10.8128 0.0013229 **
## Residuals                1.1739e+13 120
## ---
## Signif. codes:  0 '***' 0.001 '**' 0.01 '*' 0.05 '.' 0.1 ' ' 1
```

```
#Final model without Streptomyces asterosporus DSM 41452
DF<-ALL_wo_astero[,c("id", "Core_region_size","Central_compartment_size", "Chromosome_size", "d_max_rrn_ori","Number_of_rrn_
genes")]
row.names(DF)<-DF$id
```

```
## Warning: Setting row names on a tibble is deprecated.
```

```
Model0<-lm(Core_region_size~1, DF)
ModelFinal3<-lm(Core_region_size~d_max_rrn_ori+Chromosome_size+Central_compartment_size+Number_of_rrn_genes, DF)
summary(ModelFinal2)
```

```
##
## Call:
## lm(formula = Core_region_size ~ d_max_rrn_ori + Chromosome_size +
##     Central_compartment_size + Number_of_rrn_genes, data = DF)
##
## Residuals:
##     Min      1Q  Median      3Q     Max
## -892177 -174499  -22613  164547 1663768
##
## Coefficients:
##                            Estimate Std. Error t value Pr(>|t|)
## (Intercept)               1.016e+06  4.974e+05   2.042 0.043332 *
## d_max_rrn_ori             7.262e-01  2.603e-01   2.790 0.006136 **
## Chromosome_size           2.398e-01  3.970e-02   6.041 1.77e-08 ***
## Central_compartment_size  5.522e-01  1.468e-01   3.762 0.000262 ***
## Number_of_rrn_genes      -6.694e+04  2.036e+04  -3.288 0.001323 **
## ---
## Signif. codes:  0 '***' 0.001 '**' 0.01 '*' 0.05 '.' 0.1 ' ' 1
##
## Residual standard error: 312800 on 120 degrees of freedom
## Multiple R-squared:  0.8639, Adjusted R-squared:  0.8594
## F-statistic: 190.4 on 4 and 120 DF,  p-value: < 2.2e-16
```

```
par(mfrow=c(2,2))
plot(ModelFinal3,cex.lab=1.5,cex.main=1.5,pch=16)
```

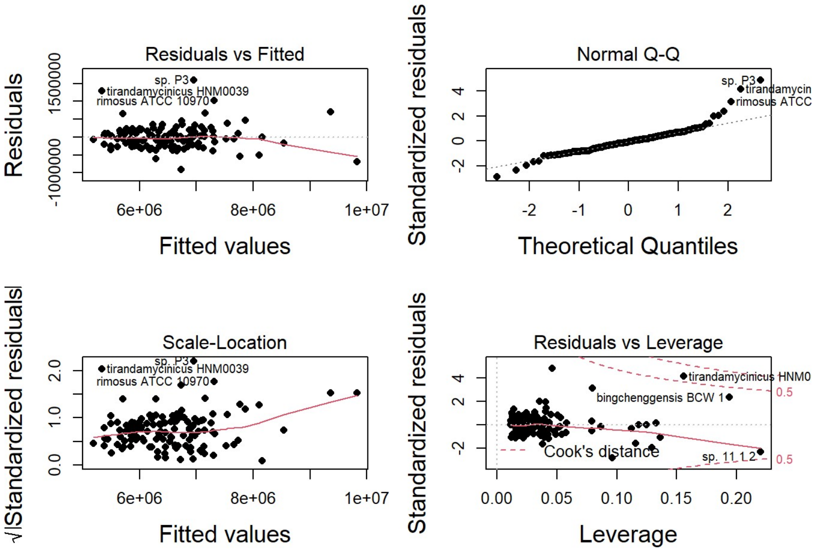


```
anova(ModelFinal3, Model0, test= "Chisq")
```

```
## Analysis of Variance Table
##
## Model 1: Core_region_size ~ d_max_rrn_ori + Chromosome_size + Central_compartment_size +
##     Number_of_rrn_genes
## Model 2: Core_region_size ~ 1
##   Res.Df        RSS Df  Sum of Sq  Pr(>Chi)
## 1    121 1.3697e+13
## 2    125 8.6267e+13 -4 -7.257e+13 < 2.2e-16 ***
## ---
## Signif. codes:  0 '***' 0.001 '**' 0.01 '*' 0.05 '.' 0.1 ' ' 1
```

```
anova(ModelFinal3)
```

```
## Analysis of Variance Table
##
## Response: Core_region_size
##                           Df     Sum Sq    Mean Sq F value    Pr(>F)
## d_max_rrn_ori              1 6.4884e+13 6.4884e+13 573.185 < 2.2e-16 ***
## Chromosome_size            1 3.8838e+12 3.8838e+12  34.310 4.138e-08 ***
## Central_compartment_size   1 1.7677e+12 1.7677e+12  15.616  0.000131 ***
## Number_of_rrn_genes        1 2.0348e+12 2.0348e+12  17.975 4.399e-05 ***
## Residuals                121 1.3697e+13 1.1320e+11
## ---
## Signif. codes:  0 '***' 0.001 '**' 0.01 '*' 0.05 '.' 0.1 ' ' 1
```

```
Anova(ModelFinal3)
```

```
## Anova Table (Type II tests)
##
## Response: Core_region_size
##                              Sum Sq  Df F value    Pr(>F)
## d_max_rrn_ori            2.5228e+11   1  2.2287    0.1381
## Chromosome_size          4.5292e+12   1 40.0113 4.422e-09 ***
## Central_compartment_size 2.0149e+12   1 17.7994 4.768e-05 ***
## Number_of_rrn_genes      2.0348e+12   1 17.9753 4.399e-05 ***
## Residuals                1.3697e+13 121
## ---
## Signif. codes:  0 '***' 0.001 '**' 0.01 '*' 0.05 '.' 0.1 ' ' 1
```

```
#Final model without Streptomyces tirandamycinicus HNM0039
ALL_wo_tiran<-subset(ALL, ALL$Species!="Streptomyces tirandamycinicus HNM0039")
DF<-ALL_wo_tiran[,c("id", "Core_region_size","Central_compartment_size", "Chromosome_size", "d_max_rrn_ori","Number_of_rrn_g
enes")]
row.names(DF)<-DF$id
```

```
## Warning: Setting row names on a tibble is deprecated.
```

```
Model0<-lm(Core_region_size~1, DF)
ModelFinal4<-lm(Core_region_size~d_max_rrn_ori+Chromosome_size+Central_compartment_size+Number_of_rrn_genes, DF)
summary(ModelFinal2)
```

```
##
## Call:
## lm(formula = Core_region_size ~ d_max_rrn_ori + Chromosome_size +
##     Central_compartment_size + Number_of_rrn_genes, data = DF)
##
## Residuals:
##     Min      1Q  Median      3Q     Max
## -892177 -174499  -22613  164547 1663768
##
## Coefficients:
##                            Estimate Std. Error t value Pr(>|t|)
## (Intercept)               1.016e+06  4.974e+05   2.042 0.043332 *
## d_max_rrn_ori             7.262e-01  2.603e-01   2.790 0.006136 **
## Chromosome_size           2.398e-01  3.970e-02   6.041 1.77e-08 ***
## Central_compartment_size  5.522e-01  1.468e-01   3.762 0.000262 ***
## Number_of_rrn_genes      -6.694e+04  2.036e+04  -3.288 0.001323 **
## ---
## Signif. codes:  0 '***' 0.001 '**' 0.01 '*' 0.05 '.' 0.1 ' ' 1
##
## Residual standard error: 312800 on 120 degrees of freedom
## Multiple R-squared:  0.8639, Adjusted R-squared:  0.8594
## F-statistic: 190.4 on 4 and 120 DF,  p-value: < 2.2e-16
```

```
par(mfrow=c(2,2))
plot(ModelFinal4,cex.lab=1.5,cex.main=1.5,pch=16)
```

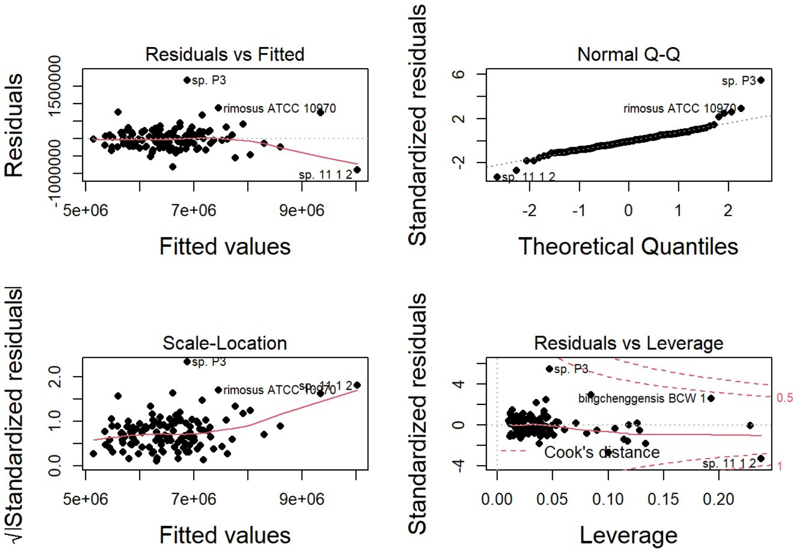


```
anova(ModelFinal4, Model0, test= "Chisq")
```

```
## Analysis of Variance Table
##
## Model 1: Core_region_size ~ d_max_rrn_ori + Chromosome_size + Central_compartment_size +
##     Number_of_rrn_genes
## Model 2: Core_region_size ~ 1
##   Res.Df        RSS Df   Sum of Sq  Pr(>Chi)
## 1    121 1.1739e+13
## 2    125 8.6310e+13 -4 -7.4571e+13 < 2.2e-16 ***
## ---
## Signif. codes:  0 '***' 0.001 '**' 0.01 '*' 0.05 '.' 0.1 ' ' 1
```

```
anova(ModelFinal4)
```

```
## Analysis of Variance Table
##
## Response: Core_region_size
##                           Df     Sum Sq    Mean Sq F value    Pr(>F)
## d_max_rrn_ori              1 6.9274e+13 6.9274e+13 714.032 < 2.2e-16 ***
## Chromosome_size            1 2.9565e+12 2.9565e+12  30.474 1.963e-07 ***
## Central_compartment_size   1 1.0277e+12 1.0277e+12  10.593  0.001472 **
## Number_of_rrn_genes        1 1.3124e+12 1.3124e+12  13.528  0.000352 ***
## Residuals                121 1.1739e+13 9.7018e+10
## ---
## Signif. codes:  0 '***' 0.001 '**' 0.01 '*' 0.05 '.' 0.1 ' ' 1
```

```
Anova(ModelFinal4)
```

```
## Anova Table (Type II tests)
##
## Response: Core_region_size
##                              Sum Sq  Df F value    Pr(>F)
## d_max_rrn_ori            7.6129e+11   1  7.8469 0.0059289 **
## Chromosome_size          3.5716e+12   1 36.8138 1.533e-08 ***
## Central_compartment_size 1.3967e+12   1 14.3967 0.0002327 ***
## Number_of_rrn_genes      1.3124e+12   1 13.5277 0.0003520 ***
## Residuals                1.1739e+13 121
## ---
## Signif. codes:  0 '***' 0.001 '**' 0.01 '*' 0.05 '.' 0.1 ' ' 1
```

```
#Comparison of the predictions (models established with all strains versus (ModelFinal1) without some strains (ModelFinal2, 3 and 4))
ALL$predict_ModelFinal1<-predict(ModelFinal1, ALL)
ALL$predict_ModelFinal2<-predict(ModelFinal2, ALL)
ALL$predict_ModelFinal3<-predict(ModelFinal3, ALL)
ALL$predict_ModelFinal4<-predict(ModelFinal4, ALL)
write.csv2(ALL,"ALL_prediction.csv", row.names = FALSE)

par(mfrow=c(1,1))
ggscatter(ALL, x = "Core_region_size", y = "predict_ModelFinal1",
          add = "reg.line", conf.int = TRUE,
          cor.coef = TRUE, cor.method = "pearson",
          xlab = "Observed size of the core region (bp)", ylab = "Predicted size of the core region (bp)")
```

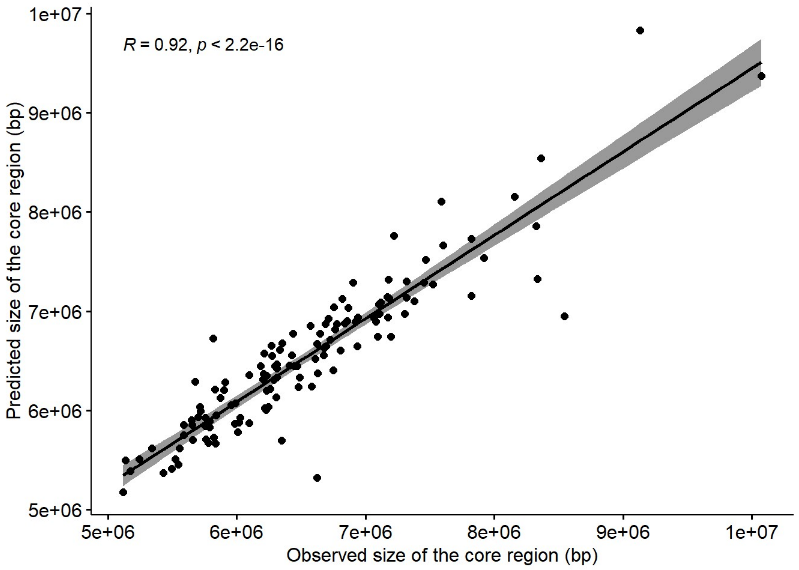


```
ggscatter(ALL, x = "Core_region_size", y = "predict_ModelFinal1",
          add = "reg.line", conf.int = TRUE,
          cor.coef = TRUE, cor.method = "pearson",
          color = "Clade", palette = c("firebrick3","blue","yellow3"),
          xlab = "Observed size of the core region (bp)", ylab = "Predicted size of the core region (bp)")
```

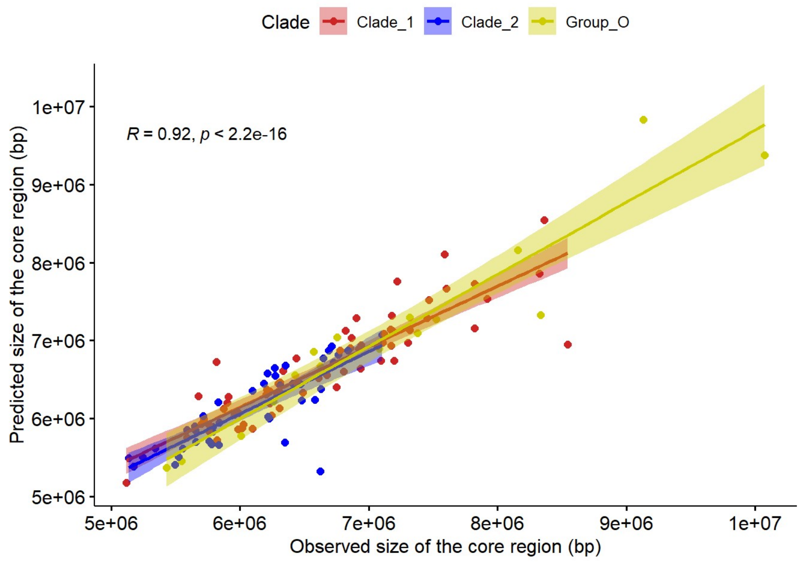


```
ggscatter(ALL, x = "Core_region_size", y = "predict_ModelFinal2",
          add = "reg.line", conf.int = TRUE,
          cor.coef = TRUE, cor.method = "pearson",
          color = "Clade", palette = c("firebrick3","blue","yellow3"),
          xlab = "Observed size of the core region (bp)", ylab = "Predicted size of the core region (bp)")
```

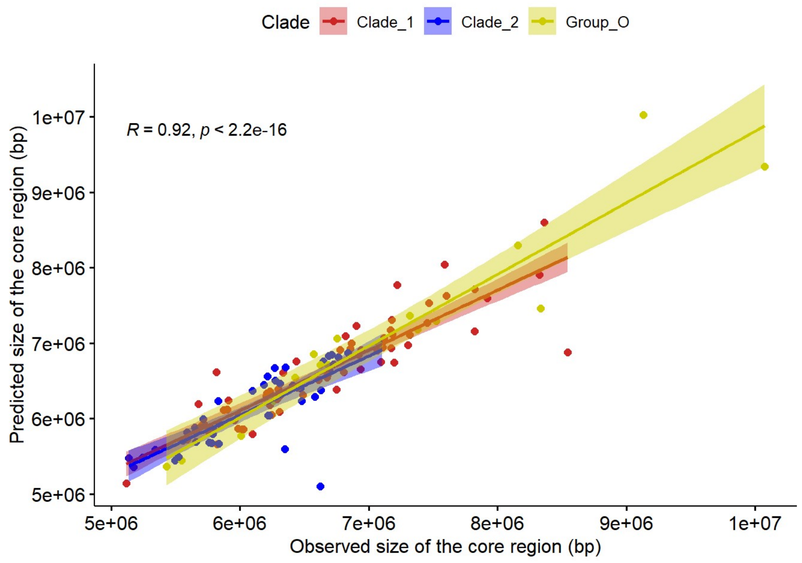


```
ggscatter(ALL, x = "Core_region_size", y = "predict_ModelFinal3",
          add = "reg.line", conf.int = TRUE,
          cor.coef = TRUE, cor.method = "pearson",
          color = "Clade", palette = c("firebrick3","blue","yellow3"),
          xlab = "Observed size of the core region (bp)", ylab = "Predicted size of the core region (bp)")
```

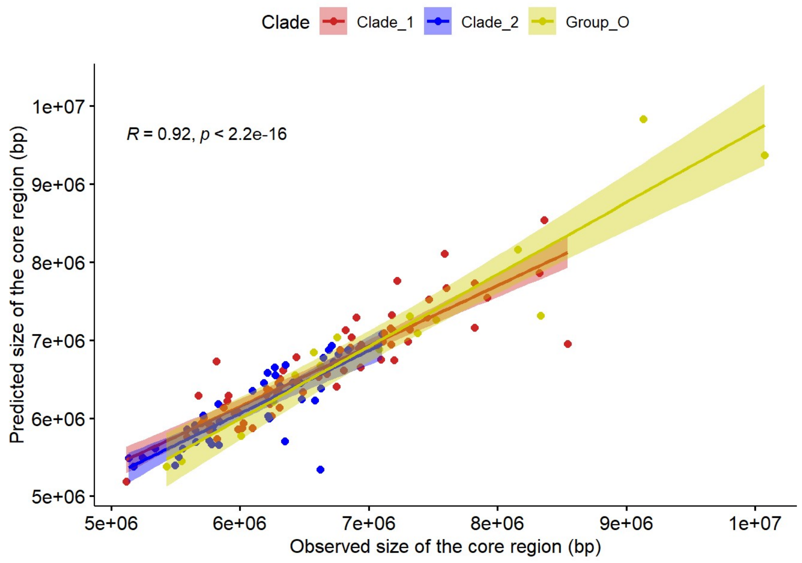

```
ggscatter(ALL, x = "Core_region_size", y = "predict_ModelFinal4",
          add = "reg.line", conf.int = TRUE,
          cor.coef = TRUE, cor.method = "pearson",
          color = "Clade", palette = c("firebrick3","blue","yellow3"),
          xlab = "Observed size of the core region (bp)", ylab = "Predicted size of the core region (bp)")
```

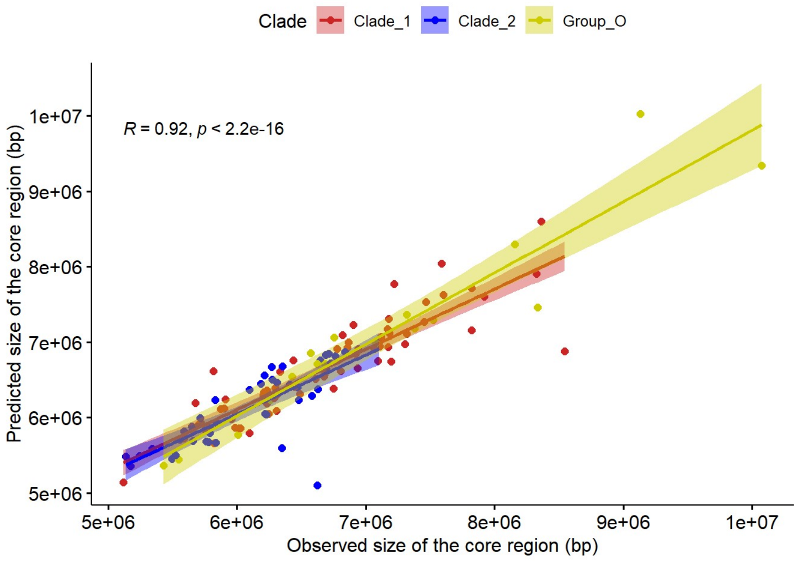


```
Clade_1<-subset(ALL, ALL$Clade=="Clade_1")
Clade_2<-subset(ALL, ALL$Clade=="Clade_2")
Group_O<-subset(ALL, ALL$Clade=="Group_O")

cor.test(ALL$predict_ModelFinal1, ALL$Core_region_size, method = "spearman")
```

```
##
##  Spearman's rank correlation rho
##
## data:  ALL$predict_ModelFinal1 and ALL$Core_region_size
## S = 26444, p-value < 2.2e-16
## alternative hypothesis: true rho is not equal to 0
## sample estimates:
##      rho
## 0.922537
```

```
cor.test(Clade_1$predict_ModelFinal1, Clade_1$Core_region_size, method = "spearman")
```

```
##
##  Spearman's rank correlation rho
##
## data:  Clade_1$predict_ModelFinal1 and Clade_1$Core_region_size
## S = 4122, p-value < 2.2e-16
## alternative hypothesis: true rho is not equal to 0
## sample estimates:
##       rho
## 0.9177508
```

```
cor.test(Clade_2$predict_ModelFinal1, Clade_2$Core_region_size, method = "spearman")
```

```
##
##  Spearman's rank correlation rho
##
## data:  Clade_2$predict_ModelFinal1 and Clade_2$Core_region_size
## S = 2510, p-value < 2.2e-16
## alternative hypothesis: true rho is not equal to 0
## sample estimates:
##       rho
## 0.8104802
```

```
cor.test(Group_O$predict_ModelFinal1, Group_O$Core_region_size, method = "spearman")
```

```
##
##  Spearman's rank correlation rho
##
## data:  Group_O$predict_ModelFinal1 and Group_O$Core_region_size
## S = 20, p-value = 5.41e-06
## alternative hypothesis: true rho is not equal to 0
## sample estimates:
##       rho
## 0.9754902
```

```
cor.test(ALL$predict_ModelFinal2, ALL$Core_region_size, method = "spearman")
```

```
##
##  Spearman's rank correlation rho
##
## data:  ALL$predict_ModelFinal2 and ALL$Core_region_size
## S = 25900, p-value < 2.2e-16
## alternative hypothesis: true rho is not equal to 0
## sample estimates:
##       rho
## 0.9241306
```

```
cor.test(Clade_1$predict_ModelFinal2, Clade_1$Core_region_size, method = "spearman")
```

```
##
##  Spearman's rank correlation rho
##
## data:  Clade_1$predict_ModelFinal2 and Clade_1$Core_region_size
## S = 4244, p-value < 2.2e-16
## alternative hypothesis: true rho is not equal to 0
## sample estimates:
##       rho
## 0.9153165
```

```
cor.test(Clade_2$predict_ModelFinal2, Clade_2$Core_region_size, method = "spearman")
```

```
##
##  Spearman's rank correlation rho
##
## data:  Clade_2$predict_ModelFinal2 and Clade_2$Core_region_size
## S = 2624, p-value < 2.2e-16
## alternative hypothesis: true rho is not equal to 0
## sample estimates:
##       rho
## 0.8018725
```

```
cor.test(Group_O$predict_ModelFinal2, Group_O$Core_region_size, method = "spearman")
```

```
##
##  Spearman's rank correlation rho
##
## data:  Group_O$predict_ModelFinal2 and Group_O$Core_region_size
## S = 18, p-value = 6.127e-06
## alternative hypothesis: true rho is not equal to 0
## sample estimates:
##       rho
## 0.9779412
```

```
cor.test(ALL$predict_ModelFinal3, ALL$Core_region_size, method = "spearman")
```

```
##
##  Spearman's rank correlation rho
##
## data:  ALL$predict_ModelFinal3 and ALL$Core_region_size
## S = 26430, p-value < 2.2e-16
## alternative hypothesis: true rho is not equal to 0
## sample estimates:
##      rho
## 0.922578
```

```
cor.test(Clade_1$predict_ModelFinal3, Clade_1$Core_region_size, method = "spearman")
```

```
##
##  Spearman's rank correlation rho
##
## data:  Clade_1$predict_ModelFinal3 and Clade_1$Core_region_size
## S = 4082, p-value < 2.2e-16
## alternative hypothesis: true rho is not equal to 0
## sample estimates:
##      rho
## 0.918549
```

```
cor.test(Clade_2$predict_ModelFinal3, Clade_2$Core_region_size, method = "spearman")
```

```
##
##  Spearman's rank correlation rho
##
## data:  Clade_2$predict_ModelFinal3 and Clade_2$Core_region_size
## S = 2458, p-value < 2.2e-16
## alternative hypothesis: true rho is not equal to 0
## sample estimates:
##       rho
## 0.8144065
```

```
cor.test(Group_O$predict_ModelFinal3, Group_O$Core_region_size, method = "spearman")
```

```
##
##  Spearman's rank correlation rho
##
## data:  Group_O$predict_ModelFinal3 and Group_O$Core_region_size
## S = 20, p-value = 5.41e-06
## alternative hypothesis: true rho is not equal to 0
## sample estimates:
##       rho
## 0.9754902
```

```
cor.test(ALL$predict_ModelFinal4, ALL$Core_region_size, method = "spearman")
```

```
##
##  Spearman's rank correlation rho
##
## data:  ALL$predict_ModelFinal4 and ALL$Core_region_size
## S = 26014, p-value < 2.2e-16
## alternative hypothesis: true rho is not equal to 0
## sample estimates:
##       rho
## 0.9237966
```

```
cor.test(Clade_1$predict_ModelFinal4, Clade_1$Core_region_size, method = "spearman")
```

```
##
##  Spearman's rank correlation rho
##
## data:  Clade_1$predict_ModelFinal4 and Clade_1$Core_region_size
## S = 4244, p-value < 2.2e-16
## alternative hypothesis: true rho is not equal to 0
## sample estimates:
##       rho
## 0.9153165
```

```
cor.test(Clade_2$predict_ModelFinal4, Clade_2$Core_region_size, method = "spearman")
```

```
##
##  Spearman's rank correlation rho
##
## data:  Clade_2$predict_ModelFinal4 and Clade_2$Core_region_size
## S = 2654, p-value < 2.2e-16
## alternative hypothesis: true rho is not equal to 0
## sample estimates:
##       rho
## 0.7996074
```

```
cor.test(Group_O$predict_ModelFinal4, Group_O$Core_region_size, method = "spearman")
```

```
##
##  Spearman's rank correlation rho
##
## data:  Group_O$predict_ModelFinal4 and Group_O$Core_region_size
## S = 18, p-value = 6.127e-06
## alternative hypothesis: true rho is not equal to 0
## sample estimates:
##       rho
## 0.9779412
```

```
cor.test(ALL$predict_ModelFinal1, ALL$Core_region_size, method = "pearson")
```

```
##
##  Pearson's product-moment correlation
##
## data:  ALL$predict_ModelFinal1 and ALL$Core_region_size
## t = 25.712, df = 125, p-value < 2.2e-16
## alternative hypothesis: true correlation is not equal to 0
## 95 percent confidence interval:
##  0.8840812 0.9409384
## sample estimates:
##       cor
## 0.9170517
```

```
cor.test(Clade_1$predict_ModelFinal1, Clade_1$Core_region_size, method = "pearson")
```

```
##
##  Pearson's product-moment correlation
##
## data:  Clade_1$predict_ModelFinal1 and Clade_1$Core_region_size
## t = 15.903, df = 65, p-value < 2.2e-16
## alternative hypothesis: true correlation is not equal to 0
## 95 percent confidence interval:
##  0.8294301 0.9323795
## sample estimates:
##       cor
## 0.8919332
```

```
cor.test(Clade_2$predict_ModelFinal1, Clade_2$Core_region_size, method = "pearson")
```

```
##
##  Pearson's product-moment correlation
##
## data:  Clade_2$predict_ModelFinal1 and Clade_2$Core_region_size
## t = 9.0613, df = 41, p-value = 2.439e-11
## alternative hypothesis: true correlation is not equal to 0
## 95 percent confidence interval:
##  0.6841358 0.8969986
## sample estimates:
##      cor
## 0.816673
```

```
cor.test(Group_O$predict_ModelFinal1, Group_O$Core_region_size, method = "pearson")
```

```
##
##  Pearson's product-moment correlation
##
## data:  Group_O$predict_ModelFinal1 and Group_O$Core_region_size
## t = 11.994, df = 15, p-value = 4.354e-09
## alternative hypothesis: true correlation is not equal to 0
## 95 percent confidence interval:
##  0.8679698 0.9827575
## sample estimates:
##       cor
## 0.9516149
```

```
cor.test(ALL$predict_ModelFinal2, ALL$Core_region_size, method = "pearson")
```

```
##
##  Pearson's product-moment correlation
##
## data:  ALL$predict_ModelFinal2 and ALL$Core_region_size
## t = 25.433, df = 125, p-value < 2.2e-16
## alternative hypothesis: true correlation is not equal to 0
## 95 percent confidence interval:
##  0.8818789 0.9397818
## sample estimates:
##       cor
## 0.9154477
```

```
cor.test(Clade_1$predict_ModelFinal2, Clade_1$Core_region_size, method = "pearson")
```

```
##
##  Pearson's product-moment correlation
##
## data:  Clade_1$predict_ModelFinal2 and Clade_1$Core_region_size
## t = 16.392, df = 65, p-value < 2.2e-16
## alternative hypothesis: true correlation is not equal to 0
## 95 percent confidence interval:
##  0.8376953 0.9358312
## sample estimates:
##       cor
## 0.8973398
```

```
cor.test(Clade_2$predict_ModelFinal2, Clade_2$Core_region_size, method = "pearson")
```

```
##
##  Pearson's product-moment correlation
##
## data:  Clade_2$predict_ModelFinal2 and Clade_2$Core_region_size
## t = 8.249, df = 41, p-value = 3.023e-10
## alternative hypothesis: true correlation is not equal to 0
## 95 percent confidence interval:
##  0.6418942 0.8812160
## sample estimates:
##       cor
## 0.7899453
```

```
cor.test(Group_O$predict_ModelFinal2, Group_O$Core_region_size, method = "pearson")
```

```
##
##  Pearson's product-moment correlation
##
## data:  Group_O$predict_ModelFinal2 and Group_O$Core_region_size
## t = 11.698, df = 15, p-value = 6.123e-09
## alternative hypothesis: true correlation is not equal to 0
## 95 percent confidence interval:
##  0.8619856 0.9819257
## sample estimates:
##       cor
## 0.9493191
```

```
cor.test(ALL$predict_ModelFinal3, ALL$Core_region_size, method = "pearson")
```

```
##
##  Pearson's product-moment correlation
##
## data:  ALL$predict_ModelFinal3 and ALL$Core_region_size
## t = 25.703, df = 125, p-value < 2.2e-16
## alternative hypothesis: true correlation is not equal to 0
## 95 percent confidence interval:
##  0.8840139 0.9409030
## sample estimates:
##       cor
## 0.9170026
```

```
cor.test(Clade_1$predict_ModelFinal3, Clade_1$Core_region_size, method = "pearson")
```

```
##
##  Pearson's product-moment correlation
##
## data:  Clade_1$predict_ModelFinal3 and Clade_1$Core_region_size
## t = 15.864, df = 65, p-value < 2.2e-16
## alternative hypothesis: true correlation is not equal to 0
## 95 percent confidence interval:
##  0.8287427 0.9320916
## sample estimates:
##       cor
## 0.8914828
```

```
cor.test(Clade_2$predict_ModelFinal3, Clade_2$Core_region_size, method = "pearson")
```

```
##
##  Pearson's product-moment correlation
##
## data:  Clade_2$predict_ModelFinal3 and Clade_2$Core_region_size
## t = 9.1678, df = 41, p-value = 1.763e-11
## alternative hypothesis: true correlation is not equal to 0
## 95 percent confidence interval:
##  0.6891902 0.8988514
## sample estimates:
##       cor
## 0.8198334
```

```
cor.test(Group_O$predict_ModelFinal3, Group_O$Core_region_size, method = "pearson")
```

```
##
##  Pearson's product-moment correlation
##
## data:  Group_O$predict_ModelFinal3 and Group_O$Core_region_size
## t = 11.944, df = 15, p-value = 4.609e-09
## alternative hypothesis: true correlation is not equal to 0
## 95 percent confidence interval:
##  0.8669885 0.9826214
## sample estimates:
##       cor
## 0.9512391
```

```
cor.test(ALL$predict_ModelFinal4, ALL$Core_region_size, method = "pearson")
```

```
##
##  Pearson's product-moment correlation
##
## data:  ALL$predict_ModelFinal4 and ALL$Core_region_size
## t = 25.429, df = 125, p-value < 2.2e-16
## alternative hypothesis: true correlation is not equal to 0
## 95 percent confidence interval:
##  0.8818527 0.9397680
## sample estimates:
##       cor
## 0.9154285
```

```
cor.test(Clade_1$predict_ModelFinal4, Clade_1$Core_region_size, method = "pearson")
```

```
##
##  Pearson's product-moment correlation
##
## data:  Clade_1$predict_ModelFinal4 and Clade_1$Core_region_size
## t = 16.394, df = 65, p-value < 2.2e-16
## alternative hypothesis: true correlation is not equal to 0
## 95 percent confidence interval:
##  0.8377158 0.9358397
## sample estimates:
##       cor
## 0.8973531
```

```
cor.test(Clade_2$predict_ModelFinal4, Clade_2$Core_region_size, method = "pearson")
```

```
##
##  Pearson's product-moment correlation
##
## data:  Clade_2$predict_ModelFinal4 and Clade_2$Core_region_size
## t = 8.242, df = 41, p-value = 3.09e-10
## alternative hypothesis: true correlation is not equal to 0
## 95 percent confidence interval:
##  0.6414995 0.8810660
## sample estimates:
##       cor
## 0.7896929
```

```
cor.test(Group_O$predict_ModelFinal4, Group_O$Core_region_size, method = "pearson")
```

```
##
##  Pearson's product-moment correlation
##
## data:  Group_O$predict_ModelFinal4 and Group_O$Core_region_size
## t = 11.7, df = 15, p-value = 6.106e-09
## alternative hypothesis: true correlation is not equal to 0
## 95 percent confidence interval:
##  0.8620335 0.9819323
## sample estimates:
##       cor
## 0.9493375
```

```
#The results of these tests lead to the choice of Final model 1 (including all strains).
par(mfrow=c(1,1))
plot(ALL$Core_region_size, ALL$predict_ModelFinal1, col=ifelse(ALL$Species=="Streptomyces tirandamycinicus HNM0039", "blac
k", "grey"), pch=20,xlab="Observed size of the core region (bp)", ylab="Predicted size of the core region (bp)")
```

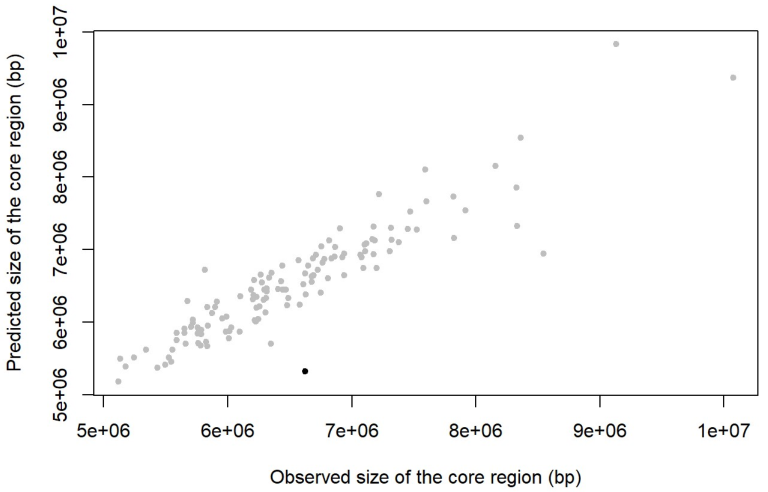


```
plot(ALL$Core_region_size, ALL$predict_ModelFinal1, col=ifelse(ALL$Species=="Streptomyces sp. P3", "black", "grey"), pch=20,
xlab="Observed size of the core region (bp)", ylab="Predicted size of the core region (bp)")
```

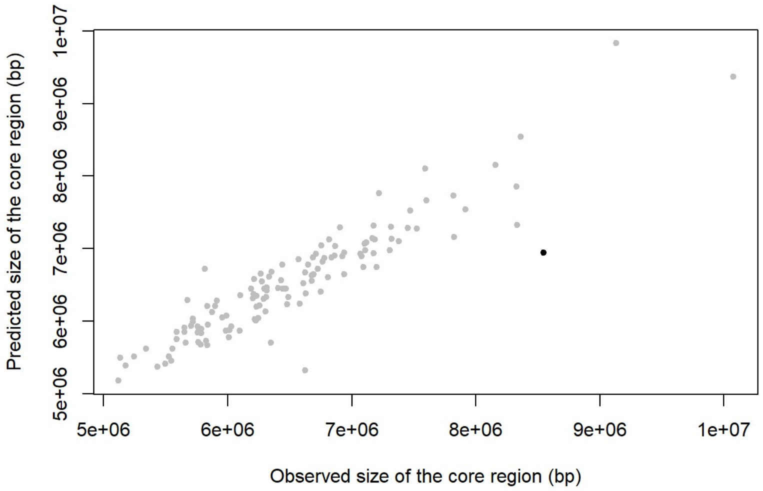


```
plot(ALL$Core_region_size, ALL$predict_ModelFinal1, col=ifelse(ALL$Species=="Streptomyces rimosus ATCC 10970", "black", "gre
y"), pch=20,xlab="Observed size of the core region (bp)", ylab="Predicted size of the core region (bp)")
```

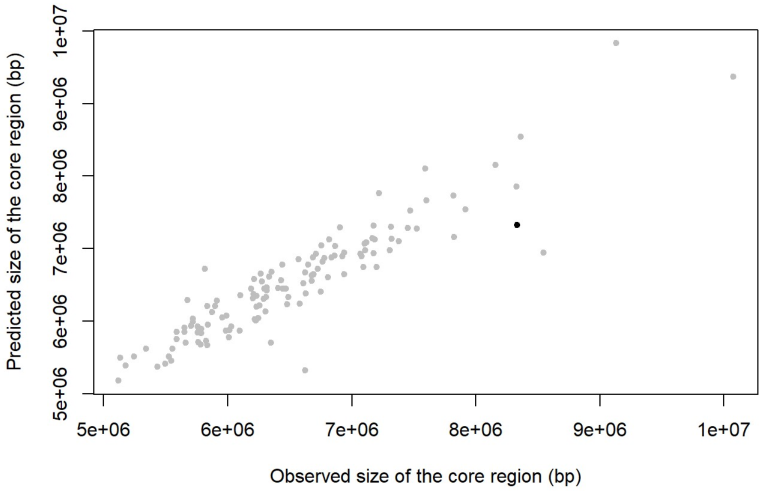


```
plot(ALL$Core_region_size, ALL$predict_ModelFinal1, col=ifelse(ALL$Species=="Streptomyces sp. 11 1 2", "black", "grey"), pch
=20,xlab="Observed size of the core region (bp)", ylab="Predicted size of the core region (bp)")
```

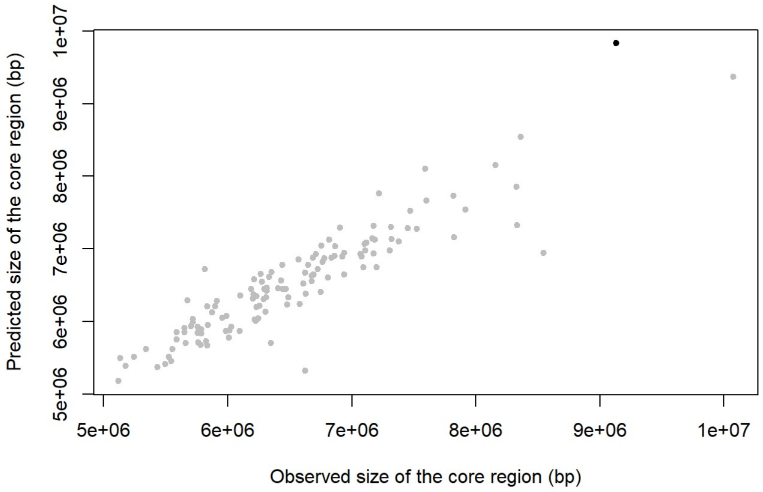


```
plot(ALL$Core_region_size, ALL$predict_ModelFinal1, col=ifelse(ALL$Species=="Streptomyces bingchenggensis BCW 1", "black", "
grey"), pch=20,xlab="Observed size of the core region (bp)", ylab="Predicted size of the core region (bp)")
```

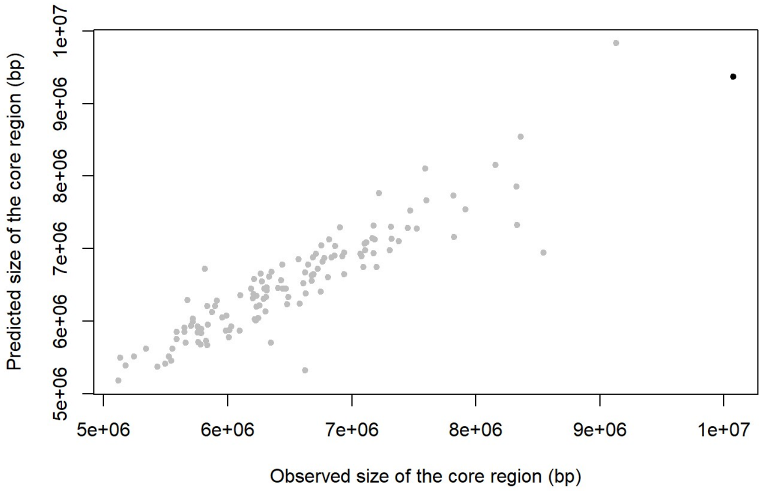


```
plot(ALL$Core_region_size, ALL$predict_ModelFinal1, col=ifelse(ALL$Species=="Streptomyces asterosporus DSM 41452", "black",
"grey"), pch=20,xlab="Observed size of the core region (bp)", ylab="Predicted size of the core region (bp)")
```

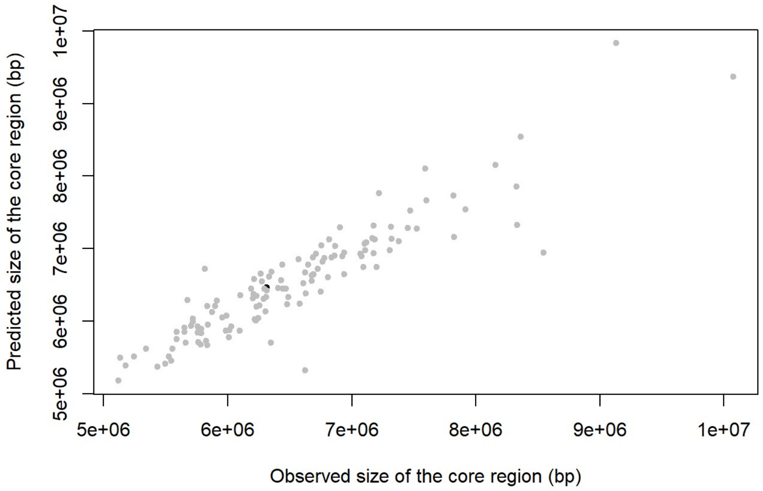


*Streptomyces tirandamycinicus* HNM0039 (in black below) harbors only 16 rDNA genes.

```
par(mfrow=c(1,1))
plot(ALL$Core_region_size, ALL$Number_of_rrn_genes, col=ifelse(ALL$Species=="Streptomyces tirandamycinicus HNM0039", "blac
k", "grey"), xlab="Size of the core region (bp)", ylab="Number of rDNA genes", pch=20)
```

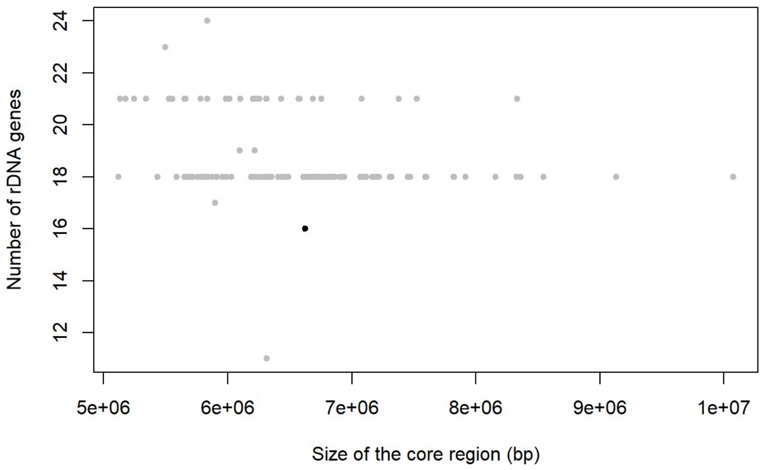


*Streptomyces rimosus* ATCC 10970 (in black below) presents the largest core region of the genomes harboring at least 7 rDNA operons.

```
par(mfrow=c(1,1))
plot(ALL$Core_region_size, ALL$Number_of_rrn_genes, col=ifelse(ALL$Species=="Streptomyces rimosus ATCC 10970", "black", "gre
y"), xlab="Size of the core region (bp)", ylab="Number of rDNA genes", pch=20)
```

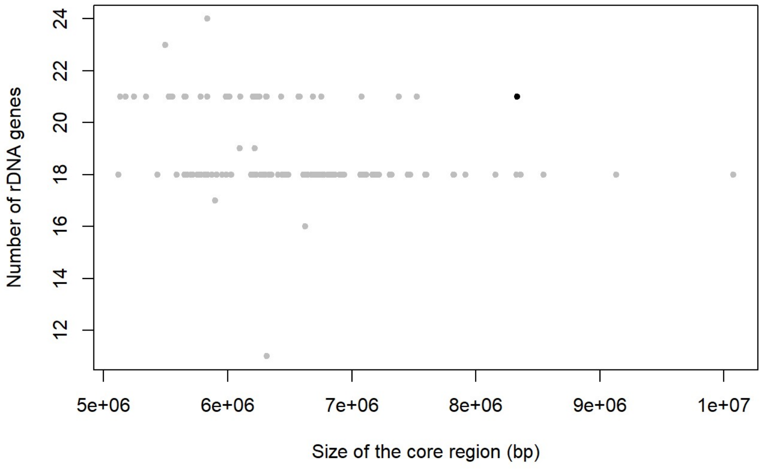


*Streptomyces* sp. P3 (in black below) genome is highly assymetrical.

```
par(mfrow=c(1,1))
plot(ALL$d_max_core_ori, ALL$d_min_core_ori, col=ifelse(ALL$Species=="Streptomyces sp. P3", "black", "grey"), xlab="d_max_co
re_ori", ylab="d_min_core_ori", pch=20)
```

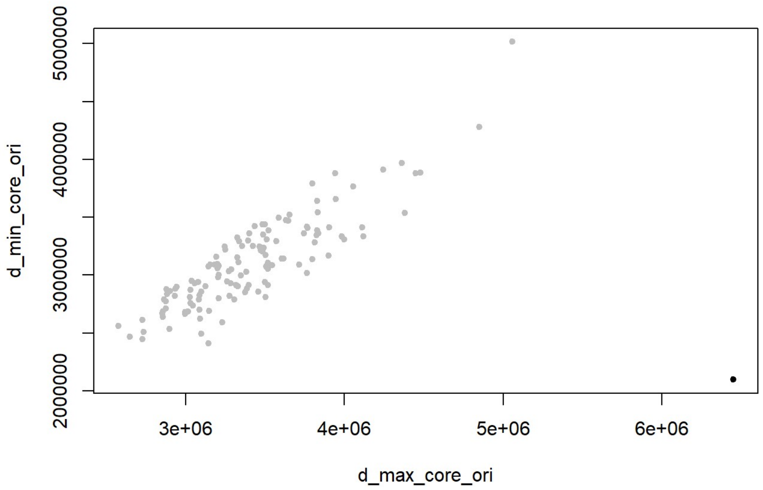


*Streptomyces bingchenggensis* BWC1 (in black below) genome presents the largest core region

```
par(mfrow=c(1,1))
plot(ALL$Core_region_size, ALL$Number_of_rrn_genes, col=ifelse(ALL$Species=="Streptomyces bingchenggensis BCW 1", "black", "
grey"), xlab="Size of the core region (bp)", ylab="Number of rDNA genes", pch=20)
```

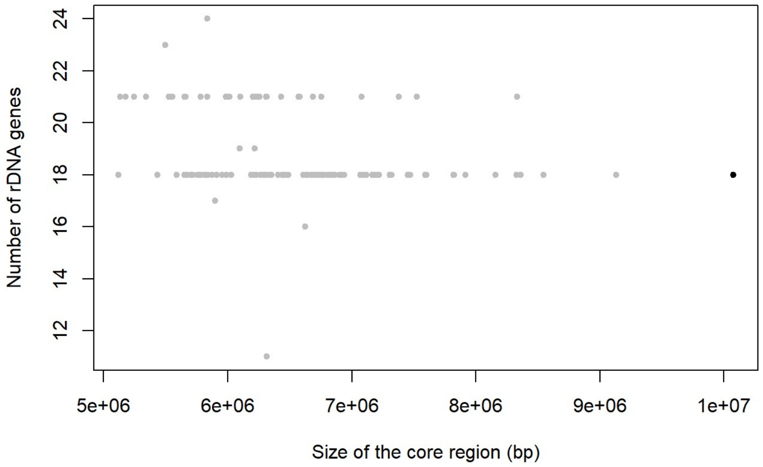


*Streptomyces* sp. 11 1 2 (in black below) genome presents the second largest core region

```
par(mfrow=c(1,1))
plot(ALL$Core_region_size, ALL$Number_of_rrn_genes, col=ifelse(ALL$Species=="Streptomyces sp. 11 1 2", "black", "grey"), xla
b="Size of the core region (bp)", ylab="Number of rDNA genes", pch=20)
```

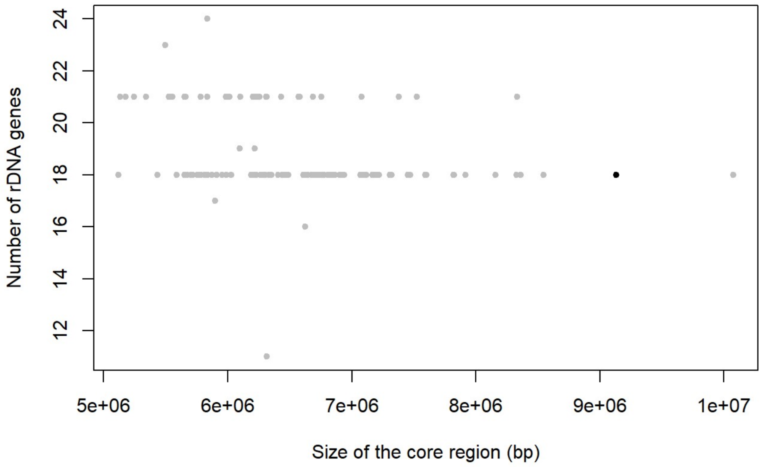


### Core region ‘gene density’ depending on the number of *rrn* genes

The previous model highlights some kind of core region ‘densification’ (more core genes, fewer non-core genes in the core region ) associated with the increase in the number of rrn operons. To evaluate this phenomenon, we analyzed the proportion of core and non-core genes in the core regions, regarding their size and the number of *rrn* operons.

```
mytable<-table(CHR$genome_id,CHR$region_core_vs_arm)
tab<-as.data.frame.matrix(mytable)
write.csv2(tab,"tab_number_genes_within_core.csv")
#These values were reported in Table S1 for further analyses.

ggscatter(ALL, x = "Core_region_size", y = "Number_of_non_core_genes_in_the_core_region",
          add = "reg.line", conf.int = TRUE,
          cor.coef = TRUE, cor.method = "pearson",
          color = "Number_of_rrn_genes",
          xlab = "Core region size (bp)", ylab = "Number of non core genes in the central compartment")+ gradient_color(c("b
lue", "lemonchiffon", "red"))
```

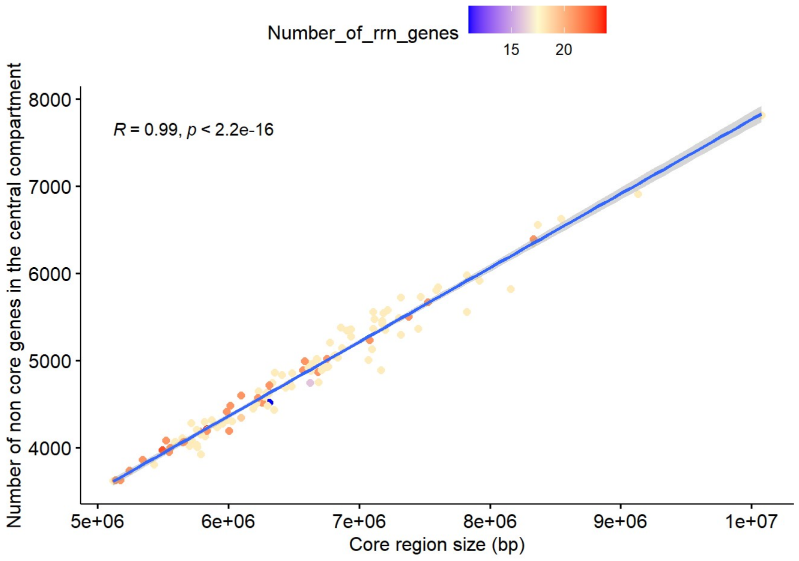


```
boxplot(ALL$Percentage_of_non_core_genes_in_the_core_region~ALL$Number_of_rrn_genes, ylab="Percentage of non core genes in t
he core region", xlab = "Number of rrn genes", frame=F, col="indianred1")
```

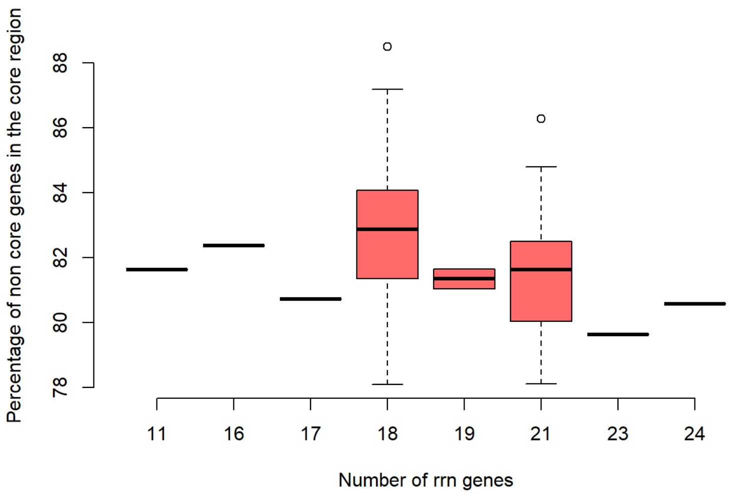


```
boxplot(ALL$Core_region_size~ALL$Number_of_rrn_genes, ylab="Core region size (bp)", xlab = "Number of rrn genes", frame=F, c
ol="indianred1")
```

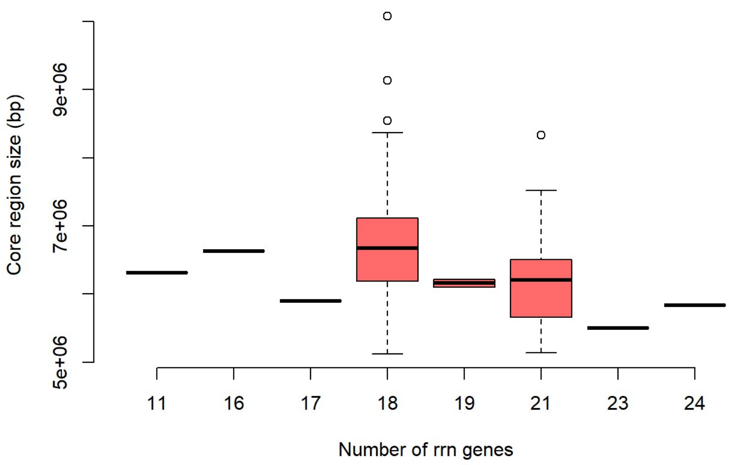

```
genome_18_max<-subset(ALL, ALL$Number_of_rrn_genes<19)
genome_19_min<-subset(ALL, ALL$Number_of_rrn_genes>18)

boxplot(genome_18_max$Percentage_of_non_core_genes_in_the_core_region, genome_19_min$Percentage_of_non_core_genes_in_the_cor
e_region, ylab="Percentage of non core genes in the core region", xlab = "Number of rrn genes", frame=F, col="indianred1", n
ames=c("18 rrn genes maximum", "19 rrn genes minimum"))
```

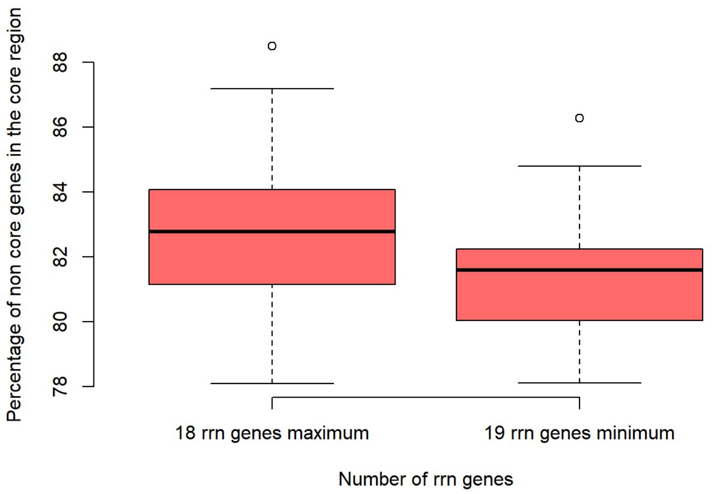


```
wilcox.test(genome_18_max$Percentage_of_non_core_genes_in_the_core_region, genome_19_min$Percentage_of_non_core_genes_in_the
_core_region)
```

```
##
##  Wilcoxon rank sum test with continuity correction
##
## data:  genome_18_max$Percentage_of_non_core_genes_in_the_core_region and genome_19_min$Percentage_of_non_core_genes_in_th
e_core_region
## W = 2243, p-value = 0.0006428
## alternative hypothesis: true location shift is not equal to 0
```

```
boxplot(genome_18_max$Core_region_size, genome_19_min$Core_region_size, ylab="Core region size (bp)", xlab = "Number of rrn
genes", frame=F, col="indianred1", names=c("18 rrn genes maximum", "19 rrn genes minimum"))
```

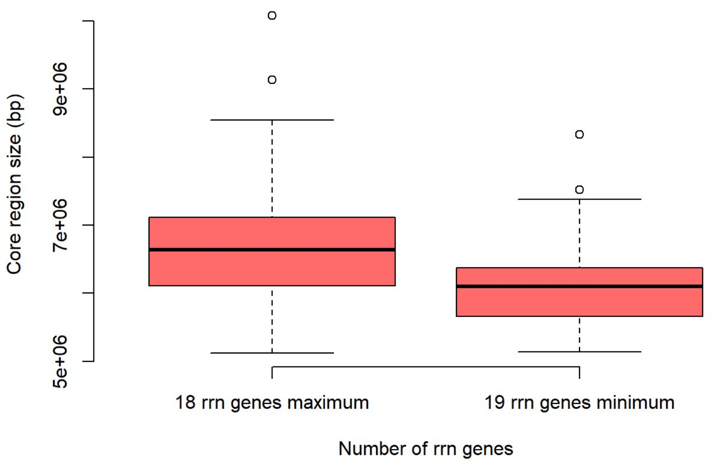


```
wilcox.test(genome_18_max$Core_region_size, genome_19_min$Core_region_size)
```

```
##
##  Wilcoxon rank sum test with continuity correction
##
## data:  genome_18_max$Core_region_size and genome_19_min$Core_region_size
## W = 2283, p-value = 0.0002849
## alternative hypothesis: true location shift is not equal to 0
```

```
ALL$central_comp_density<-ALL$Number_of_CCC_genes/ALL$Central_compartment_size*10000
ALL$delta_core_rrn_density<-ALL$Number_of_TCC_genes/ALL$delta_core_rrn*10000
genome_18_max<-subset(ALL, ALL$Number_of_rrn_genes<19)
genome_19_min<-subset(ALL, ALL$Number_of_rrn_genes>18)

boxplot(genome_18_max$central_comp_density, genome_19_min$central_comp_density, genome_18_max$delta_core_rrn_density, genome_19_min$delta_core_rrn_density, ylab="Core gene density (number of core genes per 10 kb)", frame=F, col="indianred1", names=c("18 max_central", "19 min_central", "18 max_term", "19 min_term"))
```

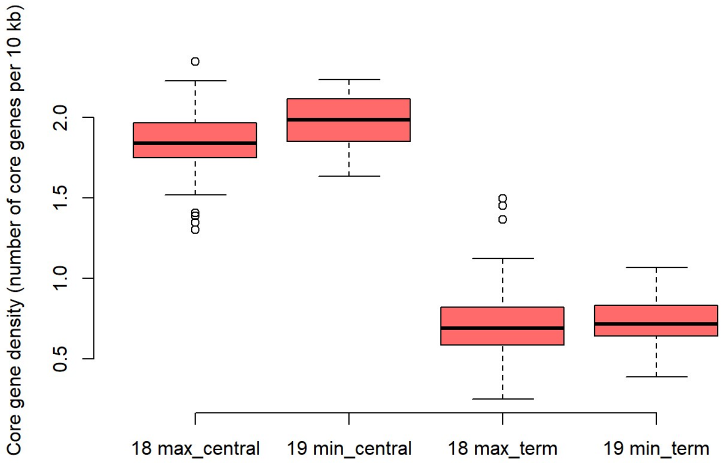


```
wilcox.test(genome_18_max$central_comp_density, genome_19_min$central_comp_density)
```

```
##
##  Wilcoxon rank sum test with continuity correction
##
## data:  genome_18_max$central_comp_density and genome_19_min$central_comp_density
## W = 950, p-value = 0.0003729
## alternative hypothesis: true location shift is not equal to 0
```

```
wilcox.test(genome_18_max$delta_core_rrn_density, genome_19_min$delta_core_rrn_density)
```

```
##
##  Wilcoxon rank sum test with continuity correction
##
## data:  genome_18_max$delta_core_rrn_density and genome_19_min$delta_core_rrn_density
## W = 1507, p-value = 0.5802
## alternative hypothesis: true location shift is not equal to 0
```

```
wilcox.test(genome_18_max$central_comp_density, genome_18_max$delta_core_rrn_density)
```

```
##
##  Wilcoxon rank sum test with continuity correction
##
## data:  genome_18_max$central_comp_density and genome_18_max$delta_core_rrn_density
## W = 8454, p-value < 2.2e-16
## alternative hypothesis: true location shift is not equal to 0
```

```
wilcox.test(genome_19_min$central_comp_density, genome_19_min$delta_core_rrn_density)
```

```
##
##  Wilcoxon rank sum exact test
##
## data:  genome_19_min$central_comp_density and genome_19_min$delta_core_rrn_density
## W = 1225, p-value < 2.2e-16
## alternative hypothesis: true location shift is not equal to 0
```

### Gene persistence around *rrn* operons

The script below allows to calculate the persistence of genes in a window of 81 CDS (it is centered on the gene of interest and/or the one closest to the rrn operon of interest, around which the average persistence is calculated by including this CDS, 40 CDSs before and 40 CDSs after).

```
ALL$persistW81_dnaA_start_position<-"NA"
ALL$persistW81_First_core_gene_position<-"NA"
ALL$persistW81_Last_core_gene_position<-"NA"
ALL$persistW81_First_rrn_position<-"NA"
ALL$persistW81_Last_rrn_position<-"NA"

ALL$dnaA_start_position<-as.numeric(ALL$dnaA_start_position)
ALL$First_core_gene_position<-as.numeric(ALL$First_core_gene_position)
ALL$Last_core_gene_position<-as.numeric(ALL$Last_core_gene_position)
ALL$First_rrn_position<-as.numeric(ALL$First_rrn_position)
ALL$Last_rrn_position<-as.numeric(ALL$Last_rrn_position)

CHR$start<-as.numeric(CHR$start)
CHR$persistence_index<-as.numeric(CHR$persistence_index)

for (i in 1:127){
  for (j in 1:950145){
    k = j+1
    l = j - 40
    m = j + 40
    if (ALL$Species[i]==CHR$genome_id[j] & ALL$dnaA_start_position[i]==CHR$start[j]){
      ALL$persistW81_dnaA_start_position[i]<-mean(CHR$persistence_index[l:m])}
  }
}


for (i in 1:127){
  for (j in 1:950145){
    k = j+1
    l = j - 40
    m = j + 40
    if (ALL$Species[i]==CHR$genome_id[j] & ALL$First_core_gene_position[i]==CHR$start[j]){
      ALL$persistW81_First_core_gene_position[i]=mean(CHR$persistence_index[l:m])}
  }
}

for (i in 1:127){
  for (j in 1:950145){
    k = j+1
    l = j - 40
    m = j + 40
    if (ALL$Species[i]==CHR$genome_id[j] & ALL$Last_core_gene_position[i]==CHR$stop[j]){
      ALL$persistW81_Last_core_gene_position[i]=mean(CHR$persistence_index[l:m])}
  }
}

for (i in 1:127){
  for (j in 1:950145){
    k = j+1
    l = j - 40
    m = j + 40
    if (ALL$Species[i]==CHR$genome_id[j] & ALL$First_rrn_position[i]>CHR$start[j] & ALL$First_rrn_position[i]<CHR$start[k]){
      ALL$persistW81_First_rrn_position[i]=mean(CHR$persistence_index[l:m])}
  }
}

for (i in 1:127){
  for (j in 1:950145){
    k = j+1
    l = j - 40
    m = j + 40
    if (ALL$Species[i]==CHR$genome_id[j] & ALL$Last_rrn_position[i]>CHR$start[j] & ALL$Last_rrn_position[i]<CHR$start[k]){
      ALL$persistW81_Last_rrn_position[i]=mean(CHR$persistence_index[l:m])}
  }
}

write.csv2(ALL,"ALL_PersistW81.csv")
```

Graph comparing the mean persistence in the neighborhood of the ori, the central compartment limits and the core region limits

```
ALL$persistW81_dnaA_start_position<-as.numeric(ALL$persistW81_dnaA_start_position)
ALL$persistW81_First_core_gene_position<-as.numeric(ALL$persistW81_First_core_gene_position)
ALL$persistW81_Last_core_gene_position<-as.numeric(ALL$persistW81_Last_core_gene_position)
ALL$persistW81_First_rrn_position<-as.numeric(ALL$persistW81_First_rrn_position)
ALL$persistW81_Last_rrn_position<-as.numeric(ALL$persistW81_Last_rrn_position)

boxplot(ALL$persistW81_dnaA_start_position, ALL$persistW81_First_rrn_position, ALL$persistW81_Last_rrn_position, ALL$persist
W81_First_core_gene_position, ALL$persistW81_Last_core_gene_position, names = c("ori", "first core", "last core", "first rr
n", "last rrn"), ylab = "mean persistence", frame = F, col = "yellow")
```

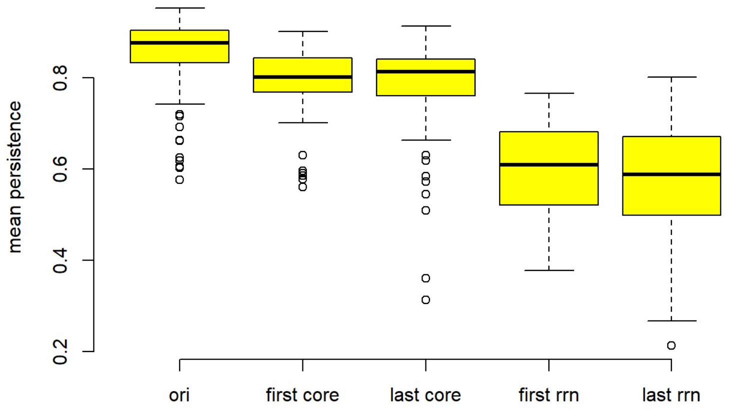


```
persistW81_core_extreme<-rbind(ALL$persistW81_First_core_gene_position, ALL$persistW81_Last_core_gene_position)
persistW81_rrn_extreme<-rbind(ALL$persistW81_First_rrn_position, ALL$persistW81_Last_rrn_position)

boxplot(ALL$persistW81_dnaA_start_position, persistW81_rrn_extreme, persistW81_core_extreme, names = c("ori", "rrn limits",
"core limits"), ylab = "mean persistence", frame = F, col = "yellow", ylim=c(0.1,1))
```

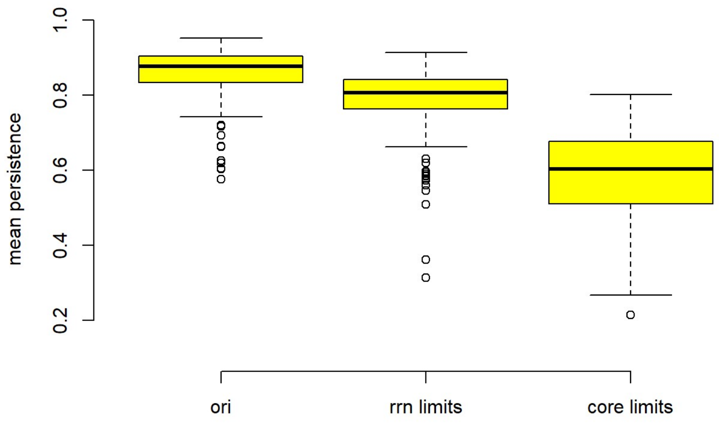


```
wilcox.test(ALL$persistW81_dnaA_start_position, persistW81_rrn_extreme)
```

```
##
##  Wilcoxon rank sum test with continuity correction
##
## data:  ALL$persistW81_dnaA_start_position and persistW81_rrn_extreme
## W = 24972, p-value < 2.2e-16
## alternative hypothesis: true location shift is not equal to 0
```

```
wilcox.test(ALL$persistW81_dnaA_start_position, persistW81_core_extreme)
```

```
##
##  Wilcoxon rank sum test with continuity correction
##
## data:  ALL$persistW81_dnaA_start_position and persistW81_core_extreme
## W = 31333, p-value < 2.2e-16
## alternative hypothesis: true location shift is not equal to 0
```

```
wilcox.test(persistW81_core_extreme, persistW81_rrn_extreme)
```

```
##
##  Wilcoxon rank sum test with continuity correction
##
## data:  persistW81_core_extreme and persistW81_rrn_extreme
## W = 3286, p-value < 2.2e-16
## alternative hypothesis: true location shift is not equal to 0
```

Calculation the mean persistence in the neighborhood of each *rrn* loci

```
RRNA$rrn_position<-as.numeric(RRNA$rrn_position)
CHR$start<-as.numeric(CHR$start)
CHR$persistence_index<-as.numeric(CHR$persistence_index)
RRNA_subset<-subset(RRNA, RRNA$rrn_position!="NA")

RRNA_subset$persistW81_rrn_loci<-"NA"
for (i in 1:798){
  for (j in 1:950145){
    k = j+1
    l = j - 40
    m = j + 40
    if (RRNA_subset$genome_id[i]==CHR$genome_id[j] & RRNA_subset$rrn_position[i]>CHR$start[j] & RRNA_subset$rrn_position
[i]<CHR$start[k]){
      RRNA_subset$persistW81_rrn_loci[i]<-mean(CHR$persistence_index[l:m])}
  }
}
RRNA_subset$persistW81_rrn_loci<-as.numeric(RRNA_subset$persistW81_rrn_loci)
write.csv2(RRNA_subset,"RRNA_PW81.csv")


RRNA$persistW81_rrn_loci<-"NA"
for (i in 1:1016){
  for (j in 1:798){
    if (RRNA$genome_id[i]==RRNA_subset$genome_id[j] & RRNA$rrn_order[i]==RRNA_subset$rrn_order[j]){
      RRNA$persistW81_rrn_loci[i]<-RRNA_subset$persistW81_rrn_loci[j]}
  }
}
}
write.csv2(RRNA,"RRNA_ALL_PW81.csv")
```

Relationship between persistence and rrn category

```
RRNA_subset<-subset(RRNA, RRNA$rrn_position!="NA")
RRNA_subset$persistW81_rrn_loci<-as.numeric(RRNA_subset$persistW81_rrn_loci)
boxplot(RRNA_subset$persistW81_rrn_loci~RRNA_subset$rrn_NOMENCLATURE_simplified_2, ylab = "mean persistence", frame = F, col
= "yellow", ylim=c(0.1,1), xlab="rrn category")
```

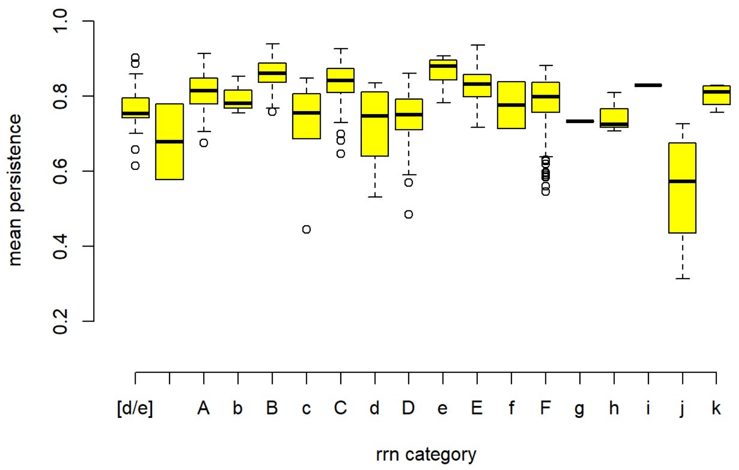


Representation of gene persistence along the chromosome of a species of interest such as *Streptomyces coelicolor* A(3)2

```
SCO<-subset(CHR, CHR$genome_id=="Streptomyces coelicolor A3 2")

ALL$rrn1<-as.numeric(ALL$rrn1)
ALL$rrn2<-as.numeric(ALL$rrn2)
ALL$rrn3<-as.numeric(ALL$rrn3)
ALL$rrn4<-as.numeric(ALL$rrn4)
ALL$rrn5<-as.numeric(ALL$rrn5)
ALL$rrn6<-as.numeric(ALL$rrn6)
```

```
## Warning: NAs introduits lors de la conversion automatique
```

```
ALL$dnaA_start_position<-as.numeric(ALL$dnaA_start_position)
ALL$First_core_gene_position<-as.numeric(ALL$First_core_gene_position)
ALL$Last_core_gene_position<-as.numeric(ALL$Last_core_gene_position)

A<-ALL$rrn1[which(ALL$Species=="Streptomyces coelicolor A3 2")]/1000000
B<-ALL$rrn2[which(ALL$Species=="Streptomyces coelicolor A3 2")]/1000000
C<-ALL$rrn3[which(ALL$Species=="Streptomyces coelicolor A3 2")]/1000000
D<-ALL$rrn4[which(ALL$Species=="Streptomyces coelicolor A3 2")]/1000000
E<-ALL$rrn5[which(ALL$Species=="Streptomyces coelicolor A3 2")]/1000000
F<-ALL$rrn6[which(ALL$Species=="Streptomyces coelicolor A3 2")]/1000000
G<-ALL$dnaA_start_position[which(ALL$Species=="Streptomyces coelicolor A3 2")]/1000000
H<-ALL$First_core_gene_position[which(ALL$Species=="Streptomyces coelicolor A3 2")]/1000000
I<-ALL$Last_core_gene_position[which(ALL$Species=="Streptomyces coelicolor A3 2")]/1000000

SCO$start<-as.numeric(SCO$start)
SCO$persistence_index<-as.numeric(SCO$persistence_index)

SCO$persist_W81<-"NA"
a<-40
b<-a+1
l<-length(SCO$persistence_index)-a
for (i in b:l){
  j = i - a
  k = i + a
  SCO$persist_W81[i]<-mean(SCO$persistence_index[j:k])
}
SCO$persist_W81<-as.numeric(SCO$persist_W81)
```

```
## Warning: NAs introduits lors de la conversion automatique
```

```
plot(SCO$start/1000000, SCO$persist_W81, col = "grey", type = "l", ylab="Mean persistence index",  xlab = "Location (Mb)", y
lim=c(0,1), axes = FALSE)
axis(1, labels=TRUE)
axis(2, labels=TRUE)
abline(v=c(A,B,C,D,E,F,G,H,I), col=c("blue","blue","blue","blue","blue","blue","red","black", "black"), lty=2, lwd=1)
```

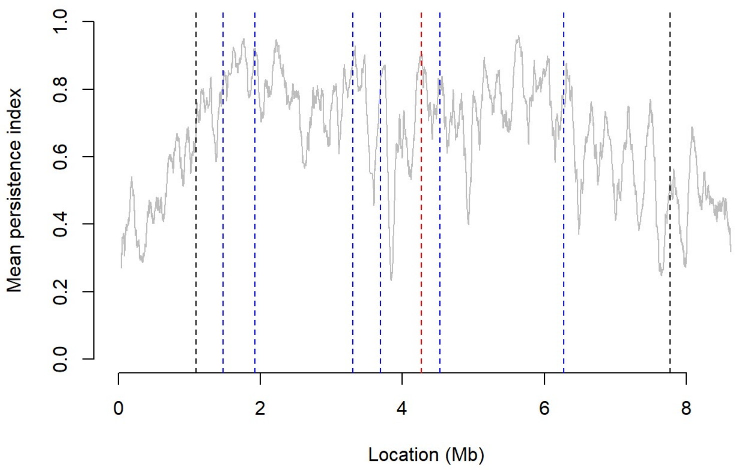


Representation of gene persistence along the chromosome of all species of the panel

```
ALL$rrn1<-as.numeric(ALL$rrn1)
ALL$rrn2<-as.numeric(ALL$rrn2)
ALL$rrn3<-as.numeric(ALL$rrn3)
ALL$rrn4<-as.numeric(ALL$rrn4)
ALL$rrn5<-as.numeric(ALL$rrn5)
ALL$rrn6<-as.numeric(ALL$rrn6)
ALL$rrn7<-as.numeric(ALL$rrn7)
ALL$rrn8<-as.numeric(ALL$rrn8)
ALL$dnaA_start_position<-as.numeric(ALL$dnaA_start_position)
ALL$First_core_gene_position<-as.numeric(ALL$First_core_gene_position)
ALL$Last_core_gene_position<-as.numeric(ALL$Last_core_gene_position)

CHR$persist_W81<-"NA"
a<-40
b<-a+1


pdf("FigSupp5.pdf",width=8, height=10)
par(mfrow=c(3,3))
for (i in 1:127){
  if (ALL$Number_of_rrn_loci[i]=="6"){
    #For species with 6 rrn loci
    SPECIES<-subset(CHR, CHR$genome_id==ALL$Species[i])
    A<-ALL$rrn1[i]/1000000
    B<-ALL$rrn2[i]/1000000
    C<-ALL$rrn3[i]/1000000
    D<-ALL$rrn4[i]/1000000
    E<-ALL$rrn5[i]/1000000
    F<-ALL$rrn6[i]/1000000
    G<-ALL$dnaA_start_position[i]/1000000
    H<-ALL$First_core_gene_position[i]/1000000
    I<-ALL$Last_core_gene_position[i]/1000000

    l<-length(SPECIES$persistence_index)-a
      for (m in b:l){
        j = m - a
        k = m + a
        SPECIES$persist_W81[m]<-mean(SPECIES$persistence_index[j:k])}

    SPECIES$persist_W81<-as.numeric(SPECIES$persist_W81)

    plot(SPECIES$start/1000000, SPECIES$persist_W81, col = "grey", type = "l", ylab="Mean persistence index",  xlab=paste("
S. ",ALL$id[i],"\n","chromosome (Mb)"), font.lab=3, ylim=c(0,1), axes = FALSE)
axis(1, labels=TRUE)
axis(2, labels=TRUE)
abline(v=c(A,B,C,D,E,F,G,H,I), col=c("blue","blue","blue","blue","blue","blue","red","black", "black"), lty=2, lwd=1)

  }else if (ALL$Number_of_rrn_loci[i]=="7"){
      #For species with 7 rrn loci
    SPECIES<-subset(CHR, CHR$genome_id==ALL$Species[i])
    A<-ALL$rrn1[i]/1000000
    B<-ALL$rrn2[i]/1000000
    C<-ALL$rrn3[i]/1000000
    D<-ALL$rrn4[i]/1000000
    E<-ALL$rrn5[i]/1000000
    F<-ALL$rrn6[i]/1000000
    G<-ALL$dnaA_start_position[i]/1000000
    H<-ALL$First_core_gene_position[i]/1000000
    I<-ALL$Last_core_gene_position[i]/1000000
    J<-ALL$rrn7[i]/1000000

    l<-length(SPECIES$persistence_index)-a
      for (m in b:l){
        j = m - a
        k = m + a
        SPECIES$persist_W81[m]<-mean(SPECIES$persistence_index[j:k])}

    SPECIES$persist_W81<-as.numeric(SPECIES$persist_W81)

    plot(SPECIES$start/1000000, SPECIES$persist_W81, col = "grey", type = "l", ylab="Mean persistence index",  xlab=paste("
S. ",ALL$id[i],"\n","chromosome (Mb)"), font.lab=3, ylim=c(0,1), axes = FALSE)
axis(1, labels=TRUE)
axis(2, labels=TRUE)
abline(v=c(A,B,C,D,E,F,J, G,H,I), col=c("blue","blue","blue","blue","blue","blue","blue","red","black", "black"), lty=2, lwd
=1)
}else if (ALL$Number_of_rrn_loci[i]=="5"){
      #For species with 5 rrn loci
    SPECIES<-subset(CHR, CHR$genome_id==ALL$Species[i])
    A<-ALL$rrn1[i]/1000000
    B<-ALL$rrn2[i]/1000000
    C<-ALL$rrn3[i]/1000000
    D<-ALL$rrn4[i]/1000000
    E<-ALL$rrn5[i]/1000000
    G<-ALL$dnaA_start_position[i]/1000000
    H<-ALL$First_core_gene_position[i]/1000000
    I<-ALL$Last_core_gene_position[i]/1000000

    l<-length(SPECIES$persistence_index)-a
      for (m in b:l){
        j = m - a
        k = m + a
```

```
        SPECIES$persist_W81[m]<-mean(SPECIES$persistence_index[j:k])}

    SPECIES$persist_W81<-as.numeric(SPECIES$persist_W81)

    plot(SPECIES$start/1000000, SPECIES$persist_W81, col = "grey", type = "l", ylab="Mean persistence index",  xlab=paste("
S. ",ALL$id[i],"\n","chromosome (Mb)"), font.lab=3, ylim=c(0,1), axes = FALSE)
axis(1, labels=TRUE)
axis(2, labels=TRUE)
abline(v=c(A,B,C,D,E,G,H,I), col=c("blue","blue","blue","blue","blue","red","black", "black"), lty=2, lwd=1)

}else if (ALL$Number_of_rrn_loci[i]=="8"){
      #For species with 8 rrn loci
    SPECIES<-subset(CHR, CHR$genome_id==ALL$Species[i])
    A<-ALL$rrn1[i]/1000000
    B<-ALL$rrn2[i]/1000000
    C<-ALL$rrn3[i]/1000000
    D<-ALL$rrn4[i]/1000000
    E<-ALL$rrn5[i]/1000000
    F<-ALL$rrn6[i]/1000000
    G<-ALL$dnaA_start_position[i]/1000000
    H<-ALL$First_core_gene_position[i]/1000000
    I<-ALL$Last_core_gene_position[i]/1000000
    J<-ALL$rrn7[i]/1000000
    K<-ALL$rrn8[i]/1000000

    l<-length(SPECIES$persistence_index)-a
      for (m in b:l){
        j = m - a
        k = m + a
        SPECIES$persist_W81[m]<-mean(SPECIES$persistence_index[j:k])}

    SPECIES$persist_W81<-as.numeric(SPECIES$persist_W81)

    plot(SPECIES$start/1000000, SPECIES$persist_W81, col = "grey", type = "l", ylab="Mean persistence index",  xlab=paste("
S. ",ALL$id[i],"\n","chromosome (Mb)"), font.lab=3, ylim=c(0,1), axes = FALSE)
axis(1, labels=TRUE)
axis(2, labels=TRUE)
abline(v=c(A,B,C,D,E,F, J, K, G,H,I), col=c("blue","blue","blue","blue","blue","blue","blue","blue","red","black", "black"),
lty=2, lwd=1)
}}

dev.off()
```

Gene content in the central compartment

```
#Location of the core genes
table(CORE$compartment)
```

```
##
##   CENTRAL_COMP CORE_TERM_COMP
##         114347          14798
```

```
#Frequency of occurence of each core ortholog group in the central and terminal compartments
CORE$ortholog_group<-as.factor(CORE$ortholog_group)
TAB1<-table(CORE$ortholog_group,CORE$compartment)
TAB1<-as.data.frame.matrix(TAB1)
TAB1$CENTRAL_COMP<-as.numeric(TAB1$CENTRAL_COMP)
TAB1$CORE_TERM_COMP<-as.numeric(TAB1$CORE_TERM_COMP)
TAB1$ortholog_group<-row.names(TAB1)
TAB1$frequency_central<-TAB1$CENTRAL_COMP/127*100
TAB1$frequency_terminal<-TAB1$CORE_TERM_COMP/127*100

TAB1$TYPE<-"NA"
for (i in 1:1017){
  if (TAB1$frequency_central[i]<30){
  TAB1$TYPE[i]<-"usually TCC"
  }else if (TAB1$frequency_central[i]>30){
  TAB1$TYPE[i]<-"usually CCC"
}}
write.csv2(TAB1,"Frequency_TCC_CCC.csv")

ggplot(TAB1, aes(x=frequency_central, fill=TYPE), binwidth=30)+
  labs(y = "Number of core genes", x = "Frequency of occurence of a given core gene within the central compartment (%)") +
  geom_histogram(alpha=0.5) +
theme_classic()
```

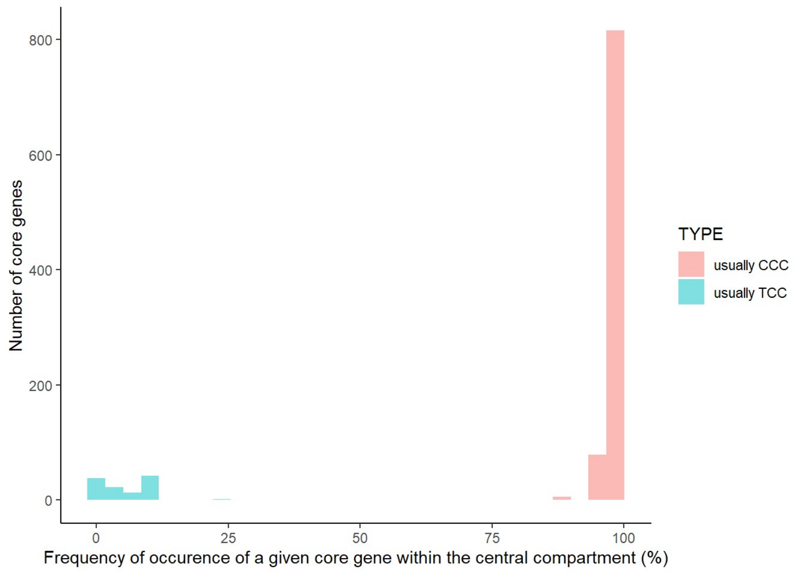


```
#Size of the central compartment
ALL$size_comp_percent<-ALL$Central_compartment_size/ALL$Chromosome_size*100
mean(ALL$size_comp_percent)
```

```
## [1] 57.70409
```

```
sd(ALL$size_comp_percent)
```

```
## [1] 5.716921
```

```
ALL$CCC_percent<-ALL$Number_of_CCC_genes/1017*100
mean(ALL$CCC_percent)
```

```
## [1] 88.53196
```

```
sd(ALL$CCC_percent)
```

```
## [1] 3.055819
```

Gene content in the central compartment of *S. bingchenggensis* BCW 1

```
CORE_BING<-subset(CORE, CORE$genome_id=="Streptomyces bingchenggensis BCW 1")
CORE_VIRIDO<-subset(CORE, CORE$genome_id=="Streptomyces viridosporus T7A ATCC 39115")

CORE_BING$USUAL_LOCATION<-"NA"
for (i in 1:1017){
  for (j in 1:1017){
    if (CORE_BING$ortholog_group[i]==CORE_VIRIDO$ortholog_group[j]){
      CORE_BING$USUAL_LOCATION[i]<-CORE_VIRIDO$compartment[j]}
  }
}

CORE_BING$compartment_type<-"NA"
for (i in 1:1017){
    if (CORE_BING$compartment[i]==CORE_BING$USUAL_LOCATION[i]){
      CORE_BING$compartment_type[i]<-"usual"}
}

for (i in 1:1017){
    if (CORE_BING$compartment[i]!=CORE_BING$USUAL_LOCATION[i] & CORE_BING$compartment[i]=="CENTRAL_COMP"){
      CORE_BING$compartment_type[i]<-"TCC->CCC"}
}

for (i in 1:1017){
    if (CORE_BING$compartment[i]!=CORE_BING$USUAL_LOCATION[i] & CORE_BING$compartment[i]=="CORE_TERM_COMP"){
      CORE_BING$compartment_type[i]<-"CCC->TCC"}
}

table(CORE_BING$compartment_type)
```

```
##
## CCC->TCC TCC->CCC    usual
##       46       85      886
```

### Analysis of gene expression inside and outside the central compartment

Script for the mapping and couting - Example for one strain

```
#In the working directory, place:

#   a subfolder ‘Fastq_files’ containing the fastq files;

#   a subfolder ‘Bam_files’ which will contain the results;

#   a subfolder ‘Genome’ which will contain the .fasta genome

#   a subfolder 'Counts'

#   a subfolder 'Annotation' containing the gff file (e.g. 'SCO.gff')

#INDEX
STAR   --runMode genomeGenerate   --runThreadN 4   --genomeDir ./Genome   --genomeFastaFiles ./Genome/GCF_000203835.1_ASM203
83v1_genomic_ONE_TIR.fna      --genomeSAindexNbases 4

#Import data from SRA
fastq-dump --split-files SRR2043965
fastq-dump --split-files SRR2043964
fastq-dump --split-files SRR2043958
fastq-dump --split-files SRR2043959
fastq-dump --split-files SRR2043960
fastq-dump --split-files SRR2043961
fastq-dump --split-files SRR2043962
fastq-dump --split-files SRR2043963

#Mapping
STAR --genomeDir ./Genome/ --runThreadN 4 --outFileNamePrefix ./Bam_files/SCO_S2. --readFilesIn ./Fastq_files/SRR2043965_1.f
astq --outSAMtype BAM SortedByCoordinate --alignIntronMax 1000 --alignMatesGapMax 10000 --limitBAMsortRAM 1259563133
STAR --genomeDir ./Genome/ --runThreadN 4 --outFileNamePrefix ./Bam_files/SCO_S1. --readFilesIn ./Fastq_files/SRR2043964_1.f
astq --outSAMtype BAM SortedByCoordinate --alignIntronMax 1000 --alignMatesGapMax 10000 --limitBAMsortRAM 1259563133
STAR --genomeDir ./Genome/ --runThreadN 4 --outFileNamePrefix ./Bam_files/SCO_M1. --readFilesIn ./Fastq_files/SRR2043958_1.f
astq --outSAMtype BAM SortedByCoordinate --alignIntronMax 1000 --alignMatesGapMax 10000 --limitBAMsortRAM 1259563133
STAR --genomeDir ./Genome/ --runThreadN 4 --outFileNamePrefix ./Bam_files/SCO_M2. --readFilesIn ./Fastq_files/SRR2043959_1.f
astq --outSAMtype BAM SortedByCoordinate --alignIntronMax 1000 --alignMatesGapMax 10000 --limitBAMsortRAM 1259563133
STAR --genomeDir ./Genome/ --runThreadN 4 --outFileNamePrefix ./Bam_files/SCO_T1. --readFilesIn ./Fastq_files/SRR2043960_1.f
astq --outSAMtype BAM SortedByCoordinate --alignIntronMax 1000 --alignMatesGapMax 10000 --limitBAMsortRAM 1259563133
STAR --genomeDir ./Genome/ --runThreadN 4 --outFileNamePrefix ./Bam_files/SCO_T2. --readFilesIn ./Fastq_files/SRR2043961_1.f
astq --outSAMtype BAM SortedByCoordinate --alignIntronMax 1000 --alignMatesGapMax 10000 --limitBAMsortRAM 1259563133
STAR --genomeDir ./Genome/ --runThreadN 4 --outFileNamePrefix ./Bam_files/SCO_L1. --readFilesIn ./Fastq_files/SRR2043962_1.f
astq --outSAMtype BAM SortedByCoordinate --alignIntronMax 1000 --alignMatesGapMax 10000 --limitBAMsortRAM 1259563133
STAR --genomeDir ./Genome/ --runThreadN 4 --outFileNamePrefix ./Bam_files/SCO_L2. --readFilesIn ./Fastq_files/SRR2043963_1.f
astq --outSAMtype BAM SortedByCoordinate --alignIntronMax 1000 --alignMatesGapMax 10000 --limitBAMsortRAM 1259563133

#Counting
featureCounts -s 2 -t gene -g ID -a ./Annotation/SCO.gff -T 4 -o ./Counts/SCO_DB_M1_counts_S.txt ./Bam_files/SCO_M1.Aligned.
sortedByCoord.out.bam
featureCounts -s 2 -t gene -g ID -a ./Annotation/SCO.gff -T 4 -o ./Counts/SCO_DB_M2_counts_S.txt ./Bam_files/SCO_M2.Aligned.
sortedByCoord.out.bam
featureCounts -s 2 -t gene -g ID -a ./Annotation/SCO.gff -T 4 -o ./Counts/SCO_DB_T1_counts_S.txt ./Bam_files/SCO_T1.Aligned.
sortedByCoord.out.bam
featureCounts -s 2 -t gene -g ID -a ./Annotation/SCO.gff -T 4 -o ./Counts/SCO_DB_T2_counts_S.txt ./Bam_files/SCO_T2.Aligned.
sortedByCoord.out.bam
featureCounts -s 2 -t gene -g ID -a ./Annotation/SCO.gff -T 4 -o ./Counts/SCO_DB_L1_counts_S.txt ./Bam_files/SCO_L1.Aligned.
sortedByCoord.out.bam
featureCounts -s 2 -t gene -g ID -a ./Annotation/SCO.gff -T 4 -o ./Counts/SCO_DB_L2_counts_S.txt ./Bam_files/SCO_L2.Aligned.
sortedByCoord.out.bam
featureCounts -s 2 -t gene -g ID -a ./Annotation/SCO.gff -T 4 -o ./Counts/SCO_DB_S1_counts_S.txt ./Bam_files/SCO_S1.Aligned.
sortedByCoord.out.bam
featureCounts -s 2 -t gene -g ID -a ./Annotation/SCO.gff -T 4 -o ./Counts/SCO_DB_S2_counts_S.txt ./Bam_files/SCO_S2.Aligned.
sortedByCoord.out.bam
```

DESeq2 normalization using SARTools - Example for one strain

```
conda activate sartools
R

rm(list=ls())
workDir <- "/shared/projects/strepthost/DESEQanalyses/TSUKU" # Indicate the appropriate working directory
projectName <- "Streptomyces transcriptomics_TSUKU"
author <- "Stéphanie Bury-Moné" # Indicate the author
targetFile <- "target_TSUKU.txt"
rawDir <- "/shared/projects/strepthost/DESEQanalyses/TSUKU/Counts"# Indicate the path to the counts

featuresToRemove <- c("alignment_not_unique", "ambiguous", "no_feature", "not_aligned", "too_low_aQual")
varInt <- "Condition"
condRef <- "E"
batch <- NULL

fitType <- "parametric"
cooksCutoff <- TRUE
independentFiltering <- TRUE
alpha <- 0.05
pAdjustMethod <- "BH"
typeTrans <- "VST"
locfunc <- "median"
colors <- c(rainbow(4)) #Indicate the number of conditions
forceCairoGraph <- FALSE


################################################################################
###                             running script                               ###
################################################################################
setwd(workDir)
library(SARTools)

checkParameters.DESeq2(projectName=projectName,author=author,targetFile=targetFile, rawDir=rawDir,featuresToRemove=featuresT
oRemove,varInt=varInt, condRef=condRef,batch=batch,fitType=fitType,cooksCutoff=cooksCutoff, independentFiltering=independent
Filtering, alpha=alpha, pAdjustMethod=pAdjustMethod, typeTrans=typeTrans, locfunc=locfunc,colors=colors)
target <- loadTargetFile(targetFile=targetFile, varInt=varInt, condRef=condRef, batch=batch)
counts <- loadCountData(target=target, rawDir=rawDir, featuresToRemove=featuresToRemove)
majSequences <- descriptionPlots(counts=counts, group=target[,varInt], col=colors)
out.DESeq2 <- run.DESeq2(counts=counts, target=target, varInt=varInt, batch=batch, locfunc=locfunc, fitType=fitType, pAdjust
Method=pAdjustMethod, cooksCutoff=cooksCutoff, independentFiltering=independentFiltering, alpha=alpha)
exploreCounts(object=out.DESeq2$dds, group=target[,varInt], typeTrans=typeTrans, col=colors)
summaryResults <- summarizeResults.DESeq2(out.DESeq2, group=target[,varInt], col=colors, independentFiltering=independentFil
tering, cooksCutoff=cooksCutoff, alpha=alpha)
writeReport.DESeq2(target=target, counts=counts, out.DESeq2=out.DESeq2, summaryResults=summaryResults, majSequences=majSeque
nces, workDir=workDir, projectName=projectName, author=author, targetFile=targetFile, rawDir=rawDir, featuresToRemove=featur
esToRemove, varInt=varInt, condRef=condRef, batch=batch, fitType=fitType, cooksCutoff=cooksCutoff, independentFiltering=inde
pendentFiltering, alpha=alpha, pAdjustMethod=pAdjustMethod, typeTrans=typeTrans, locfunc=locfunc, colors=colors)
save.image(file=paste(projectName, ".RData"))
```

Analysis of the whole RNA-seq data set (Supplemental Table S7)

```
RNA_seq$persistence_index<-as.numeric(RNA_seq$persistence_index)
```

```
## Warning: NAs introduits lors de la conversion automatique
```

```
RNA_seq_CDS<-subset(RNA_seq, RNA_seq$ortholog_group!="funcRNA")
CENTRAL<-subset(RNA_seq_CDS, RNA_seq_CDS$region=="CENTRAL")
TERM<-subset(RNA_seq_CDS, RNA_seq_CDS$region=="TERM")
CENTRAL_CORE<-subset(CENTRAL, CENTRAL$CORE_score=="1")
CENTRAL_NONCORE<-subset(CENTRAL, CENTRAL$CORE_score=="0")
TERM_CORE<-subset(TERM, TERM$CORE_score=="1")
TERM_NONCORE<-subset(TERM, TERM$CORE_score=="0")
CORE<-subset(RNA_seq_CDS, RNA_seq_CDS$CORE_score=="1")
NONCORE<-subset(RNA_seq_CDS, RNA_seq_CDS$CORE_score=="0")



#Correlation between gene expression and distance to the origin of replication
ggscatter(CENTRAL_CORE, x = "d_ori", y = "log2_NormRPK_E",
          add = "reg.line", conf.int = TRUE,
          cor.coef = TRUE, cor.method = "spearman",
          color = "genome_id", palette = c("firebrick1", "tan1", "yellow2", "lightsteelblue1", "slateblue1", "slateblue4","s
pringgreen3" ),ylim=c(0, 25),xlim=c(0, 4000000),
          xlab = "Distance to oriC (bp)", ylab = "Transcription (log2)")
```

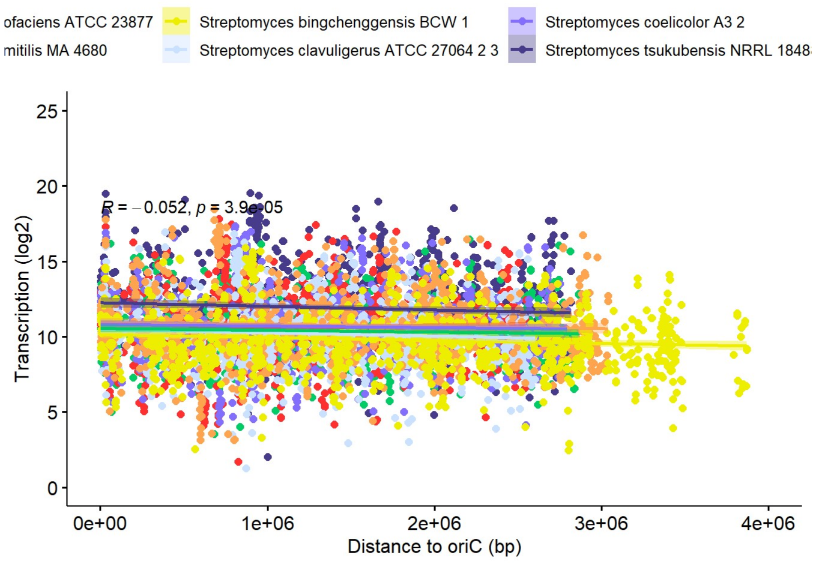


```
ggscatter(CENTRAL_NONCORE, x = "d_ori", y = "log2_NormRPK_E",
          add = "reg.line", conf.int = TRUE,
          cor.coef = TRUE, cor.method = "spearman",
          color = "genome_id", palette = c("firebrick1", "tan1", "yellow2", "lightsteelblue1", "slateblue1", "slateblue4","s
pringgreen3" ),ylim=c(0, 25),xlim=c(0, 4000000),
          xlab = "Distance to oriC (bp)", ylab = "Transcription (log2)")
```

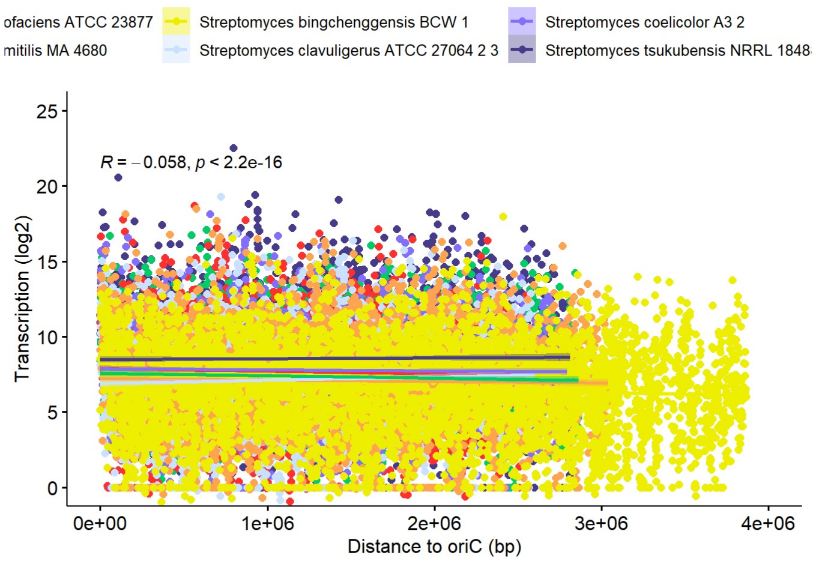


```
ggscatter(TERM_CORE, x = "d_ori", y = "log2_NormRPK_E",
          add = "reg.line", conf.int = TRUE,
          cor.coef = TRUE, cor.method = "spearman",
          color = "genome_id", palette = c("firebrick1", "tan1", "yellow2", "lightsteelblue1", "slateblue1", "slateblue4","s
pringgreen3" ),ylim=c(0, 25),xlim=c(1500000, 7000000),
          xlab = "Distance to oriC (bp)", ylab = "Transcription (log2)")
```

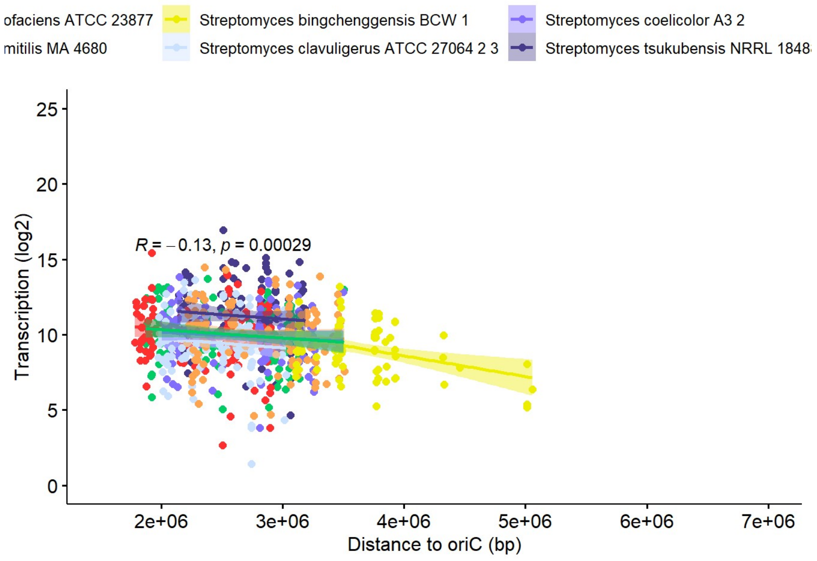


```
ggscatter(TERM_NONCORE, x = "d_ori", y = "log2_NormRPK_E",
          add = "reg.line", conf.int = TRUE,
          cor.coef = TRUE, cor.method = "spearman",
          color = "genome_id", palette = c("firebrick1", "tan1", "yellow2", "lightsteelblue1", "slateblue1", "slateblue4","s
pringgreen3" ),ylim=c(0, 25),xlim=c(1500000, 7000000),
          xlab = "Distance to oriC (bp)", ylab = "Transcription (log2)")
```

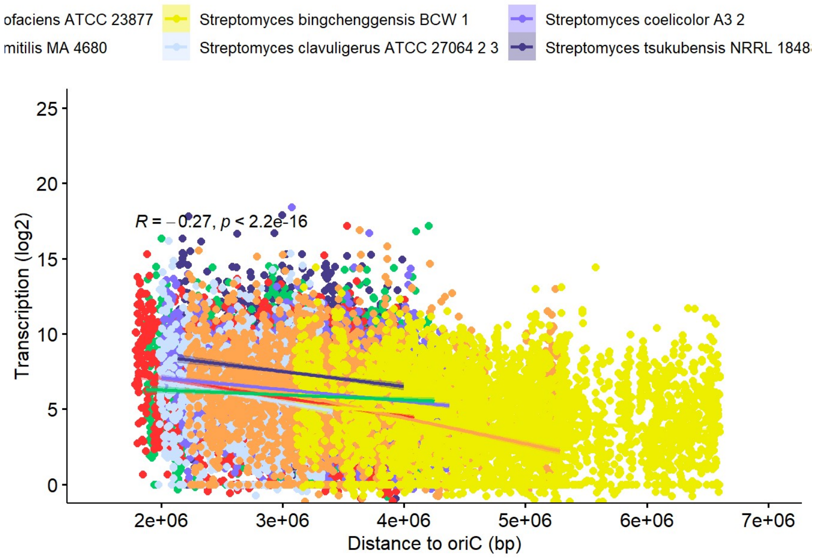


```
#Correlation between gene expression and gene persistence

ggscatter(CENTRAL, x = "persistence_index", y = "log2_NormRPK_E",
          add = "reg.line", conf.int = TRUE,
          cor.coef = TRUE, cor.method = "spearman",
          color = "genome_id", palette = c("firebrick1", "tan1", "yellow2", "lightsteelblue1", "slateblue1", "slateblue4","s
pringgreen3" ),ylim=c(0, 25),
          xlab = "Persistence index", ylab = "transcription (log2)")
```

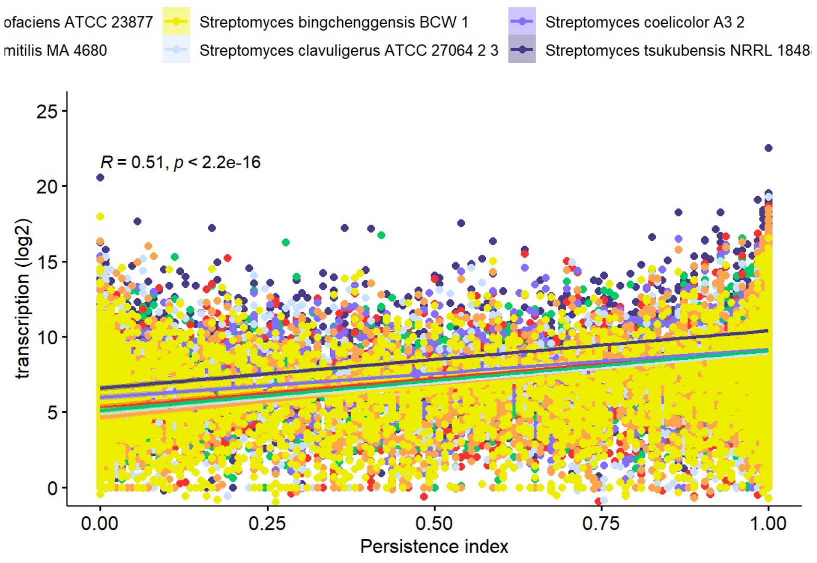


```
ggscatter(TERM, x = "persistence_index", y = "log2_NormRPK_E",
          add = "reg.line", conf.int = TRUE,
          cor.coef = TRUE, cor.method = "spearman",
          color = "genome_id", palette = c("firebrick1", "tan1", "yellow2", "lightsteelblue1", "slateblue1", "slateblue4","s
pringgreen3" ),ylim=c(0, 25),
          xlab = "Persistence index", ylab = "transcription (log2)")
```

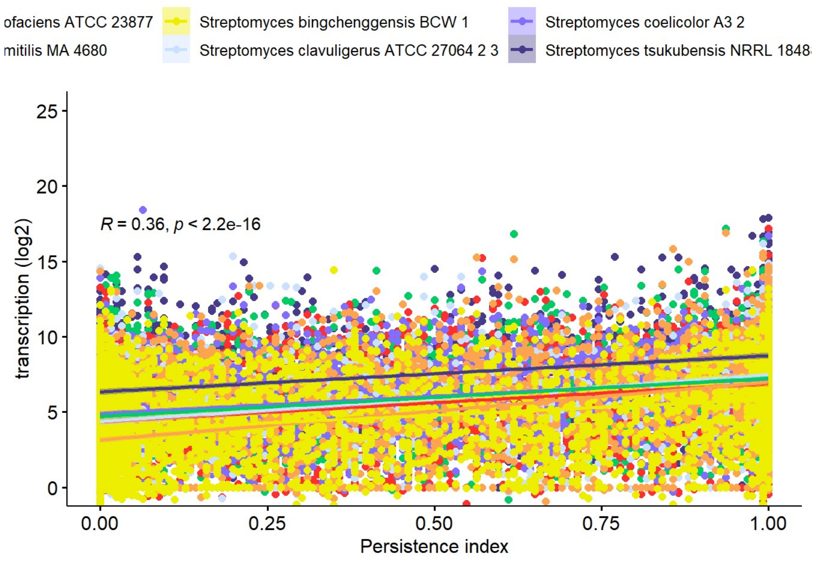


```
#Individual analyses for each strain
AVERM<-subset(RNA_seq_CDS, RNA_seq_CDS$genome_id=="Streptomyces avermitilis MA 4680")
AMBO<-subset(RNA_seq_CDS, RNA_seq_CDS$genome_id=="Streptomyces ambofaciens ATCC 23877")
BING<-subset(RNA_seq_CDS, RNA_seq_CDS$genome_id=="Streptomyces bingchenggensis BCW 1")
CLAV<-subset(RNA_seq_CDS, RNA_seq_CDS$genome_id=="Streptomyces clavuligerus ATCC 27064 2 3")
SCO<-subset(RNA_seq_CDS, RNA_seq_CDS$genome_id=="Streptomyces coelicolor A3 2")
TSUKU<-subset(RNA_seq_CDS, RNA_seq_CDS$genome_id=="Streptomyces tsukubensis NRRL 18488")
VENEZ<-subset(RNA_seq_CDS, RNA_seq_CDS$genome_id=="Streptomyces venezuelae ATCC 10712")

AMBO_CENTRAL<-subset(AMBO, AMBO$region=="CENTRAL")
AMBO_TERM<-subset(AMBO, AMBO$region=="TERM")
res <- cor.test(AMBO_CENTRAL$persistence_index, AMBO_CENTRAL$log2_NormRPK_E, method = "spearman")
```

```
## Warning in cor.test.default(AMBO_CENTRAL$persistence_index,
## AMBO_CENTRAL$log2_NormRPK_E, : Cannot compute exact p-value with ties
```

```
res
```

```
##
##  Spearman's rank correlation rho
##
## data:  AMBO_CENTRAL$persistence_index and AMBO_CENTRAL$log2_NormRPK_E
## S = 5198583651, p-value < 2.2e-16
## alternative hypothesis: true rho is not equal to 0
## sample estimates:
##       rho
## 0.4845513
```

```
res <- cor.test(AMBO_TERM$persistence_index, AMBO_TERM$log2_NormRPK_E, method = "spearman")
```

```
## Warning in cor.test.default(AMBO_TERM$persistence_index,
## AMBO_TERM$log2_NormRPK_E, : Cannot compute exact p-value with ties
```

```
res
```

```
##
##  Spearman's rank correlation rho
##
## data:  AMBO_TERM$persistence_index and AMBO_TERM$log2_NormRPK_E
## S = 4070729242, p-value < 2.2e-16
## alternative hypothesis: true rho is not equal to 0
## sample estimates:
##       rho
## 0.3135137
```

```
AMBO_CENTRAL_CORE<-subset(AMBO, AMBO$region=="CENTRAL" & AMBO$CORE_score=="1")
AMBO_TERM_CORE<-subset(AMBO, AMBO$region=="TERM" & AMBO$CORE_score=="1")
AMBO_CENTRAL_NONCORE<-subset(AMBO, AMBO$region=="CENTRAL" & AMBO$CORE_score=="0")
AMBO_TERM_NONCORE<-subset(AMBO, AMBO$region=="TERM" & AMBO$CORE_score=="0")
res <- cor.test(AMBO_CENTRAL_CORE$d_ori, AMBO_CENTRAL_CORE$log2_NormRPK_E, method = "spearman")
res
```

```
##
##  Spearman's rank correlation rho
##
## data:  AMBO_CENTRAL_CORE$d_ori and AMBO_CENTRAL_CORE$log2_NormRPK_E
## S = 124183956, p-value = 0.5752
## alternative hypothesis: true rho is not equal to 0
## sample estimates:
##         rho
## -0.01869202
```

```
res <- cor.test(AMBO_CENTRAL_NONCORE$d_ori, AMBO_CENTRAL_NONCORE$log2_NormRPK_E, method = "spearman")
```

```
## Warning in cor.test.default(AMBO_CENTRAL_NONCORE$d_ori,
## AMBO_CENTRAL_NONCORE$log2_NormRPK_E, : Cannot compute exact p-value with ties
```

```
res
```

```
##
##  Spearman's rank correlation rho
##
## data:  AMBO_CENTRAL_NONCORE$d_ori and AMBO_CENTRAL_NONCORE$log2_NormRPK_E
## S = 4799679671, p-value = 0.0264
## alternative hypothesis: true rho is not equal to 0
## sample estimates:
##         rho
## -0.04036903
```

```
res <- cor.test(AMBO_TERM_CORE$d_ori, AMBO_TERM_CORE$log2_NormRPK_E, method = "spearman")
res
```

```
##
##  Spearman's rank correlation rho
##
## data:  AMBO_TERM_CORE$d_ori and AMBO_TERM_CORE$log2_NormRPK_E
## S = 284854, p-value = 0.3097
## alternative hypothesis: true rho is not equal to 0
## sample estimates:
##         rho
## -0.09504479
```

```
res <- cor.test(AMBO_TERM_NONCORE$d_ori, AMBO_TERM_NONCORE$log2_NormRPK_E, method = "spearman")
```

```
## Warning in cor.test.default(AMBO_TERM_NONCORE$d_ori,
## AMBO_TERM_NONCORE$log2_NormRPK_E, : Cannot compute exact p-value with ties
```

```
res
```

```
##
##  Spearman's rank correlation rho
##
## data:  AMBO_TERM_NONCORE$d_ori and AMBO_TERM_NONCORE$log2_NormRPK_E
## S = 6501753381, p-value < 2.2e-16
## alternative hypothesis: true rho is not equal to 0
## sample estimates:
##        rho
## -0.2211572
```

```
BING_CENTRAL<-subset(BING, BING$region=="CENTRAL")
BING_TERM<-subset(BING, BING$region=="TERM")
res <- cor.test(BING_CENTRAL$persistence_index, BING_CENTRAL$log2_NormRPK_E, method = "spearman")
```

```
## Warning in cor.test.default(BING_CENTRAL$persistence_index,
## BING_CENTRAL$log2_NormRPK_E, : Cannot compute exact p-value with ties
```

```
res
```

```
##
##  Spearman's rank correlation rho
##
## data:  BING_CENTRAL$persistence_index and BING_CENTRAL$log2_NormRPK_E
## S = 2.2373e+10, p-value < 2.2e-16
## alternative hypothesis: true rho is not equal to 0
## sample estimates:
##       rho
## 0.4719935
```

```
res <- cor.test(BING_TERM$persistence_index, BING_TERM$log2_NormRPK_E, method = "spearman")
```

```
## Warning in cor.test.default(BING_TERM$persistence_index,
## BING_TERM$log2_NormRPK_E, : Cannot compute exact p-value with ties
```

```
res
```

```
##
##  Spearman's rank correlation rho
##
## data:  BING_TERM$persistence_index and BING_TERM$log2_NormRPK_E
## S = 7060741850, p-value < 2.2e-16
## alternative hypothesis: true rho is not equal to 0
## sample estimates:
##       rho
## 0.2752793
```

```
BING_CENTRAL_CORE<-subset(BING, BING$region=="CENTRAL" & BING$CORE_score=="1")
BING_TERM_CORE<-subset(BING, BING$region=="TERM" & BING$CORE_score=="1")
BING_CENTRAL_NONCORE<-subset(BING, BING$region=="CENTRAL" & BING$CORE_score=="0")
BING_TERM_NONCORE<-subset(BING, BING$region=="TERM" & BING$CORE_score=="0")
res <- cor.test(BING_CENTRAL_CORE$d_ori, BING_CENTRAL_CORE$log2_NormRPK_E, method = "spearman")
```

```
## Warning in cor.test.default(BING_CENTRAL_CORE$d_ori,
## BING_CENTRAL_CORE$log2_NormRPK_E, : Cannot compute exact p-value with ties
```

```
res
```

```
##
##  Spearman's rank correlation rho
##
## data:  BING_CENTRAL_CORE$d_ori and BING_CENTRAL_CORE$log2_NormRPK_E
## S = 148762908, p-value = 0.02211
## alternative hypothesis: true rho is not equal to 0
## sample estimates:
##        rho
## -0.0746396
```

```
res <- cor.test(BING_CENTRAL_NONCORE$d_ori, BING_CENTRAL_NONCORE$log2_NormRPK_E, method = "spearman")
```

```
## Warning in cor.test.default(BING_CENTRAL_NONCORE$d_ori,
## BING_CENTRAL_NONCORE$log2_NormRPK_E, : Cannot compute exact p-value with ties
```

```
res
```

```
##
##  Spearman's rank correlation rho
##
## data:  BING_CENTRAL_NONCORE$d_ori and BING_CENTRAL_NONCORE$log2_NormRPK_E
## S = 2.8098e+10, p-value = 6.164e-08
## alternative hypothesis: true rho is not equal to 0
## sample estimates:
##         rho
## -0.07362611
```

```
res <- cor.test(BING_TERM_CORE$d_ori, BING_TERM_CORE$log2_NormRPK_E, method = "spearman")
res
```

```
##
##  Spearman's rank correlation rho
##
## data:  BING_TERM_CORE$d_ori and BING_TERM_CORE$log2_NormRPK_E
## S = 90452, p-value = 0.09974
## alternative hypothesis: true rho is not equal to 0
## sample estimates:
##        rho
## -0.1889689
```

```
res <- cor.test(BING_TERM_NONCORE$d_ori, BING_TERM_NONCORE$log2_NormRPK_E, method = "spearman")
```

```
## Warning in cor.test.default(BING_TERM_NONCORE$d_ori,
## BING_TERM_NONCORE$log2_NormRPK_E, : Cannot compute exact p-value with ties
```

```
res
```

```
##
##  Spearman's rank correlation rho
##
## data:  BING_TERM_NONCORE$d_ori and BING_TERM_NONCORE$log2_NormRPK_E
## S = 1.0696e+10, p-value < 2.2e-16
## alternative hypothesis: true rho is not equal to 0
## sample estimates:
##        rho
## -0.1658435
```

```
AVERM_CENTRAL<-subset(AVERM, AVERM$region=="CENTRAL")
AVERM_TERM<-subset(AVERM, AVERM$region=="TERM")
res <- cor.test(AVERM_CENTRAL$persistence_index, AVERM_CENTRAL$log2_NormRPK_E, method = "spearman")
```

```
## Warning in cor.test.default(AVERM_CENTRAL$persistence_index,
## AVERM_CENTRAL$log2_NormRPK_E, : Cannot compute exact p-value with ties
```

```
res
```

```
##
##  Spearman's rank correlation rho
##
## data:  AVERM_CENTRAL$persistence_index and AVERM_CENTRAL$log2_NormRPK_E
## S = 7746091925, p-value < 2.2e-16
## alternative hypothesis: true rho is not equal to 0
## sample estimates:
##       rho
## 0.5489018
```

```
res <- cor.test(AVERM_TERM$persistence_index, AVERM_TERM$log2_NormRPK_E, method = "spearman")
```

```
## Warning in cor.test.default(AVERM_TERM$persistence_index,
## AVERM_TERM$log2_NormRPK_E, : Cannot compute exact p-value with ties
```

```
res
```

```
##
##  Spearman's rank correlation rho
##
## data:  AVERM_TERM$persistence_index and AVERM_TERM$log2_NormRPK_E
## S = 3538302165, p-value < 2.2e-16
## alternative hypothesis: true rho is not equal to 0
## sample estimates:
##       rho
## 0.4353078
```

```
AVERM_CENTRAL_CORE<-subset(AVERM, AVERM$region=="CENTRAL" & AVERM$CORE_score=="1")
AVERM_TERM_CORE<-subset(AVERM, AVERM$region=="TERM" & AVERM$CORE_score=="1")
AVERM_CENTRAL_NONCORE<-subset(AVERM, AVERM$region=="CENTRAL" & AVERM$CORE_score=="0")
AVERM_TERM_NONCORE<-subset(AVERM, AVERM$region=="TERM" & AVERM$CORE_score=="0")
res <- cor.test(AVERM_CENTRAL_CORE$d_ori, AVERM_CENTRAL_CORE$log2_NormRPK_E, method = "spearman")
```

```
## Warning in cor.test.default(AVERM_CENTRAL_CORE$d_ori,
## AVERM_CENTRAL_CORE$log2_NormRPK_E, : Cannot compute exact p-value with ties
```

```
res
```

```
##
##  Spearman's rank correlation rho
##
## data:  AVERM_CENTRAL_CORE$d_ori and AVERM_CENTRAL_CORE$log2_NormRPK_E
## S = 128544989, p-value = 0.1023
## alternative hypothesis: true rho is not equal to 0
## sample estimates:
##         rho
## -0.05446596
```

```
res <- cor.test(AVERM_CENTRAL_NONCORE$d_ori, AVERM_CENTRAL_NONCORE$log2_NormRPK_E, method = "spearman")
```

```
## Warning in cor.test.default(AVERM_CENTRAL_NONCORE$d_ori,
## AVERM_CENTRAL_NONCORE$log2_NormRPK_E, : Cannot compute exact p-value with ties
```

```
res
```

```
##
##  Spearman's rank correlation rho
##
## data:  AVERM_CENTRAL_NONCORE$d_ori and AVERM_CENTRAL_NONCORE$log2_NormRPK_E
## S = 9215692895, p-value = 0.2653
## alternative hypothesis: true rho is not equal to 0
## sample estimates:
##         rho
## -0.01810685
```

```
res <- cor.test(AVERM_TERM_CORE$d_ori, AVERM_TERM_CORE$log2_NormRPK_E, method = "spearman")
res
```

```
##
##  Spearman's rank correlation rho
##
## data:  AVERM_TERM_CORE$d_ori and AVERM_TERM_CORE$log2_NormRPK_E
## S = 283602, p-value = 0.3349
## alternative hypothesis: true rho is not equal to 0
## sample estimates:
##         rho
## -0.09023181
```

```
res <- cor.test(AVERM_TERM_NONCORE$d_ori, AVERM_TERM_NONCORE$log2_NormRPK_E, method = "spearman")
```

```
## Warning in cor.test.default(AVERM_TERM_NONCORE$d_ori,
## AVERM_TERM_NONCORE$log2_NormRPK_E, : Cannot compute exact p-value with ties
```

```
res
```

```
##
##  Spearman's rank correlation rho
##
## data:  AVERM_TERM_NONCORE$d_ori and AVERM_TERM_NONCORE$log2_NormRPK_E
## S = 7858705233, p-value < 2.2e-16
## alternative hypothesis: true rho is not equal to 0
## sample estimates:
##        rho
## -0.3940625
```

```
CLAV_CENTRAL<-subset(CLAV, CLAV$region=="CENTRAL")
CLAV_TERM<-subset(CLAV, CLAV$region=="TERM")
res <- cor.test(CLAV_CENTRAL$persistence_index, CLAV_CENTRAL$log2_NormRPK_E, method = "spearman")
```

```
## Warning in cor.test.default(CLAV_CENTRAL$persistence_index,
## CLAV_CENTRAL$log2_NormRPK_E, : Cannot compute exact p-value with ties
```

```
res
```

```
##
##  Spearman's rank correlation rho
##
## data:  CLAV_CENTRAL$persistence_index and CLAV_CENTRAL$log2_NormRPK_E
## S = 5127901182, p-value < 2.2e-16
## alternative hypothesis: true rho is not equal to 0
## sample estimates:
##       rho
## 0.5101315
```

```
res <- cor.test(CLAV_TERM$persistence_index, CLAV_TERM$log2_NormRPK_E, method = "spearman")
```

```
## Warning in cor.test.default(CLAV_TERM$persistence_index,
## CLAV_TERM$log2_NormRPK_E, : Cannot compute exact p-value with ties
```

```
res
```

```
##
##  Spearman's rank correlation rho
##
## data:  CLAV_TERM$persistence_index and CLAV_TERM$log2_NormRPK_E
## S = 481317604, p-value < 2.2e-16
## alternative hypothesis: true rho is not equal to 0
## sample estimates:
##       rho
## 0.4294782
```

```
CLAV_CENTRAL_CORE<-subset(CLAV, CLAV$region=="CENTRAL" & CLAV$CORE_score=="1")
CLAV_TERM_CORE<-subset(CLAV, CLAV$region=="TERM" & CLAV$CORE_score=="1")
CLAV_CENTRAL_NONCORE<-subset(CLAV, CLAV$region=="CENTRAL" & CLAV$CORE_score=="0")
CLAV_TERM_NONCORE<-subset(CLAV, CLAV$region=="TERM" & CLAV$CORE_score=="0")
res <- cor.test(CLAV_CENTRAL_CORE$d_ori, CLAV_CENTRAL_CORE$log2_NormRPK_E, method = "spearman")
```

```
## Warning in cor.test.default(CLAV_CENTRAL_CORE$d_ori,
## CLAV_CENTRAL_CORE$log2_NormRPK_E, : Cannot compute exact p-value with ties
```

```
res
```

```
##
##  Spearman's rank correlation rho
##
## data:  CLAV_CENTRAL_CORE$d_ori and CLAV_CENTRAL_CORE$log2_NormRPK_E
## S = 129098264, p-value = 0.09583
## alternative hypothesis: true rho is not equal to 0
## sample estimates:
##         rho
## -0.05548624
```

```
res <- cor.test(CLAV_CENTRAL_NONCORE$d_ori, CLAV_CENTRAL_NONCORE$log2_NormRPK_E, method = "spearman")
```

```
## Warning in cor.test.default(CLAV_CENTRAL_NONCORE$d_ori,
## CLAV_CENTRAL_NONCORE$log2_NormRPK_E, : Cannot compute exact p-value with ties
```

```
res
```

```
##
##  Spearman's rank correlation rho
##
## data:  CLAV_CENTRAL_NONCORE$d_ori and CLAV_CENTRAL_NONCORE$log2_NormRPK_E
## S = 4653598521, p-value = 0.036
## alternative hypothesis: true rho is not equal to 0
## sample estimates:
##        rho
## 0.03782841
```

```
res <- cor.test(CLAV_TERM_CORE$d_ori, CLAV_TERM_CORE$log2_NormRPK_E, method = "spearman")
res
```

```
##
##  Spearman's rank correlation rho
##
## data:  CLAV_TERM_CORE$d_ori and CLAV_TERM_CORE$log2_NormRPK_E
## S = 263504, p-value = 0.6737
## alternative hypothesis: true rho is not equal to 0
## sample estimates:
##         rho
## -0.03962755
```

```
res <- cor.test(CLAV_TERM_NONCORE$d_ori, CLAV_TERM_NONCORE$log2_NormRPK_E, method = "spearman")
```

```
## Warning in cor.test.default(CLAV_TERM_NONCORE$d_ori,
## CLAV_TERM_NONCORE$log2_NormRPK_E, : Cannot compute exact p-value with ties
```

```
res
```

```
##
##  Spearman's rank correlation rho
##
## data:  CLAV_TERM_NONCORE$d_ori and CLAV_TERM_NONCORE$log2_NormRPK_E
## S = 787928146, p-value = 1.659e-09
## alternative hypothesis: true rho is not equal to 0
## sample estimates:
##        rho
## -0.1498746
```

```
SCO_CENTRAL<-subset(SCO, SCO$region=="CENTRAL")
SCO_TERM<-subset(SCO, SCO$region=="TERM")
res <- cor.test(SCO_CENTRAL$persistence_index, SCO_CENTRAL$log2_NormRPK_E, method = "spearman")
```

```
## Warning in cor.test.default(SCO_CENTRAL$persistence_index,
## SCO_CENTRAL$log2_NormRPK_E, : Cannot compute exact p-value with ties
```

```
res
```

```
##
##  Spearman's rank correlation rho
##
## data:  SCO_CENTRAL$persistence_index and SCO_CENTRAL$log2_NormRPK_E
## S = 7320908803, p-value < 2.2e-16
## alternative hypothesis: true rho is not equal to 0
## sample estimates:
##       rho
## 0.4888899
```

```
res <- cor.test(SCO_TERM$persistence_index, SCO_TERM$log2_NormRPK_E, method = "spearman")
```

```
## Warning in cor.test.default(SCO_TERM$persistence_index,
## SCO_TERM$log2_NormRPK_E, : Cannot compute exact p-value with ties
```

```
res
```

```
##
##  Spearman's rank correlation rho
##
## data:  SCO_TERM$persistence_index and SCO_TERM$log2_NormRPK_E
## S = 4877577952, p-value < 2.2e-16
## alternative hypothesis: true rho is not equal to 0
## sample estimates:
##       rho
## 0.3115392
```

```
SCO_CENTRAL_CORE<-subset(SCO, SCO$region=="CENTRAL" & SCO$CORE_score=="1")
SCO_TERM_CORE<-subset(SCO, SCO$region=="TERM" & SCO$CORE_score=="1")
SCO_CENTRAL_NONCORE<-subset(SCO, SCO$region=="CENTRAL" & SCO$CORE_score=="0")
SCO_TERM_NONCORE<-subset(SCO, SCO$region=="TERM" & SCO$CORE_score=="0")
res <- cor.test(SCO_CENTRAL_CORE$d_ori, SCO_CENTRAL_CORE$log2_NormRPK_E, method = "spearman")
```

```
## Warning in cor.test.default(SCO_CENTRAL_CORE$d_ori,
## SCO_CENTRAL_CORE$log2_NormRPK_E, : Cannot compute exact p-value with ties
```

```
res
```

```
##
##  Spearman's rank correlation rho
##
## data:  SCO_CENTRAL_CORE$d_ori and SCO_CENTRAL_CORE$log2_NormRPK_E
## S = 125666480, p-value = 0.3549
## alternative hypothesis: true rho is not equal to 0
## sample estimates:
##         rho
## -0.03085329
```

```
res <- cor.test(SCO_CENTRAL_NONCORE$d_ori, SCO_CENTRAL_NONCORE$log2_NormRPK_E, method = "spearman")
```

```
## Warning in cor.test.default(SCO_CENTRAL_NONCORE$d_ori,
## SCO_CENTRAL_NONCORE$log2_NormRPK_E, : Cannot compute exact p-value with ties
```

```
res
```

```
##
##  Spearman's rank correlation rho
##
## data:  SCO_CENTRAL_NONCORE$d_ori and SCO_CENTRAL_NONCORE$log2_NormRPK_E
## S = 7356376556, p-value = 0.2616
## alternative hypothesis: true rho is not equal to 0
## sample estimates:
##         rho
## -0.01894728
```

```
res <- cor.test(SCO_TERM_CORE$d_ori, SCO_TERM_CORE$log2_NormRPK_E, method = "spearman")
res
```

```
##
##  Spearman's rank correlation rho
##
## data:  SCO_TERM_CORE$d_ori and SCO_TERM_CORE$log2_NormRPK_E
## S = 288062, p-value = 0.2509
## alternative hypothesis: true rho is not equal to 0
## sample estimates:
##        rho
## -0.1073771
```

```
res <- cor.test(SCO_TERM_NONCORE$d_ori, SCO_TERM_NONCORE$log2_NormRPK_E, method = "spearman")
```

```
## Warning in cor.test.default(SCO_TERM_NONCORE$d_ori,
## SCO_TERM_NONCORE$log2_NormRPK_E, : Cannot compute exact p-value with ties
```

```
res
```

```
##
##  Spearman's rank correlation rho
##
## data:  SCO_TERM_NONCORE$d_ori and SCO_TERM_NONCORE$log2_NormRPK_E
## S = 7492616494, p-value < 2.2e-16
## alternative hypothesis: true rho is not equal to 0
## sample estimates:
##        rho
## -0.1704411
```

```
TSUKU_CENTRAL<-subset(TSUKU, TSUKU$region=="CENTRAL")
TSUKU_TERM<-subset(TSUKU, TSUKU$region=="TERM")
res <- cor.test(TSUKU_CENTRAL$persistence_index, TSUKU_CENTRAL$log2_NormRPK_E, method = "spearman")
```

```
## Warning in cor.test.default(TSUKU_CENTRAL$persistence_index,
## TSUKU_CENTRAL$log2_NormRPK_E, : Cannot compute exact p-value with ties
```

```
res
```

```
##
##  Spearman's rank correlation rho
##
## data:  TSUKU_CENTRAL$persistence_index and TSUKU_CENTRAL$log2_NormRPK_E
## S = 7270685253, p-value < 2.2e-16
## alternative hypothesis: true rho is not equal to 0
## sample estimates:
##       rho
## 0.5056197
```

```
res <- cor.test(TSUKU_TERM$persistence_index, TSUKU_TERM$log2_NormRPK_E, method = "spearman")
```

```
## Warning in cor.test.default(TSUKU_TERM$persistence_index,
## TSUKU_TERM$log2_NormRPK_E, : Cannot compute exact p-value with ties
```

```
res
```

```
##
##  Spearman's rank correlation rho
##
## data:  TSUKU_TERM$persistence_index and TSUKU_TERM$log2_NormRPK_E
## S = 1319435721, p-value < 2.2e-16
## alternative hypothesis: true rho is not equal to 0
## sample estimates:
##       rho
## 0.3329418
```

```
TSUKU_CENTRAL_CORE<-subset(TSUKU, TSUKU$region=="CENTRAL" & TSUKU$CORE_score=="1")
TSUKU_TERM_CORE<-subset(TSUKU, TSUKU$region=="TERM" & TSUKU$CORE_score=="1")
TSUKU_CENTRAL_NONCORE<-subset(TSUKU, TSUKU$region=="CENTRAL" & TSUKU$CORE_score=="0")
TSUKU_TERM_NONCORE<-subset(TSUKU, TSUKU$region=="TERM" & TSUKU$CORE_score=="0")
res <- cor.test(TSUKU_CENTRAL_CORE$d_ori, TSUKU_CENTRAL_CORE$log2_NormRPK_E, method = "spearman")
res
```

```
##
##  Spearman's rank correlation rho
##
## data:  TSUKU_CENTRAL_CORE$d_ori and TSUKU_CENTRAL_CORE$log2_NormRPK_E
## S = 128459716, p-value = 0.1068
## alternative hypothesis: true rho is not equal to 0
## sample estimates:
##         rho
## -0.05376646
```

```
res <- cor.test(TSUKU_CENTRAL_NONCORE$d_ori, TSUKU_CENTRAL_NONCORE$log2_NormRPK_E, method = "spearman")
```

```
## Warning in cor.test.default(TSUKU_CENTRAL_NONCORE$d_ori,
## TSUKU_CENTRAL_NONCORE$log2_NormRPK_E, : Cannot compute exact p-value with ties
```

```
res
```

```
##
##  Spearman's rank correlation rho
##
## data:  TSUKU_CENTRAL_NONCORE$d_ori and TSUKU_CENTRAL_NONCORE$log2_NormRPK_E
## S = 7419475163, p-value = 0.7296
## alternative hypothesis: true rho is not equal to 0
## sample estimates:
##         rho
## 0.005802989
```

```
res <- cor.test(TSUKU_TERM_CORE$d_ori, TSUKU_TERM_CORE$log2_NormRPK_E, method = "spearman")
res
```

```
##
##  Spearman's rank correlation rho
##
## data:  TSUKU_TERM_CORE$d_ori and TSUKU_TERM_CORE$log2_NormRPK_E
## S = 292468, p-value = 0.1834
## alternative hypothesis: true rho is not equal to 0
## sample estimates:
##        rho
## -0.1243148
```

```
res <- cor.test(TSUKU_TERM_NONCORE$d_ori, TSUKU_TERM_NONCORE$log2_NormRPK_E, method = "spearman")
```

```
## Warning in cor.test.default(TSUKU_TERM_NONCORE$d_ori,
## TSUKU_TERM_NONCORE$log2_NormRPK_E, : Cannot compute exact p-value with ties
```

```
res
```

```
##
##  Spearman's rank correlation rho
##
## data:  TSUKU_TERM_NONCORE$d_ori and TSUKU_TERM_NONCORE$log2_NormRPK_E
## S = 1950103456, p-value = 8.196e-13
## alternative hypothesis: true rho is not equal to 0
## sample estimates:
##        rho
## -0.1530159
```

```
VENEZ_CENTRAL<-subset(VENEZ, VENEZ$region=="CENTRAL")
VENEZ_TERM<-subset(VENEZ, VENEZ$region=="TERM")
res <- cor.test(VENEZ_CENTRAL$persistence_index, VENEZ_CENTRAL$log2_NormRPK_E, method = "spearman")
```

```
## Warning in cor.test.default(VENEZ_CENTRAL$persistence_index,
## VENEZ_CENTRAL$log2_NormRPK_E, : Cannot compute exact p-value with ties
```

```
res
```

```
##
##  Spearman's rank correlation rho
##
## data:  VENEZ_CENTRAL$persistence_index and VENEZ_CENTRAL$log2_NormRPK_E
## S = 6664469141, p-value < 2.2e-16
## alternative hypothesis: true rho is not equal to 0
## sample estimates:
##       rho
## 0.5356669
```

```
res <- cor.test(VENEZ_TERM$persistence_index, VENEZ_TERM$log2_NormRPK_E, method = "spearman")
```

```
## Warning in cor.test.default(VENEZ_TERM$persistence_index,
## VENEZ_TERM$log2_NormRPK_E, : Cannot compute exact p-value with ties
```

```
res
```

```
##
##  Spearman's rank correlation rho
##
## data:  VENEZ_TERM$persistence_index and VENEZ_TERM$log2_NormRPK_E
## S = 3037925135, p-value < 2.2e-16
## alternative hypothesis: true rho is not equal to 0
## sample estimates:
##       rho
## 0.3289399
```

```
VENEZ_CENTRAL_CORE<-subset(VENEZ, VENEZ$region=="CENTRAL" & VENEZ$CORE_score=="1")
VENEZ_TERM_CORE<-subset(VENEZ, VENEZ$region=="TERM" & VENEZ$CORE_score=="1")
VENEZ_CENTRAL_NONCORE<-subset(VENEZ, VENEZ$region=="CENTRAL" & VENEZ$CORE_score=="0")
VENEZ_TERM_NONCORE<-subset(VENEZ, VENEZ$region=="TERM" & VENEZ$CORE_score=="0")
res <- cor.test(VENEZ_CENTRAL_CORE$d_ori, VENEZ_CENTRAL_CORE$log2_NormRPK_E, method = "spearman")
```

```
## Warning in cor.test.default(VENEZ_CENTRAL_CORE$d_ori,
## VENEZ_CENTRAL_CORE$log2_NormRPK_E, : Cannot compute exact p-value with ties
```

```
res
```

```
##
##  Spearman's rank correlation rho
##
## data:  VENEZ_CENTRAL_CORE$d_ori and VENEZ_CENTRAL_CORE$log2_NormRPK_E
## S = 127566000, p-value = 0.1637
## alternative hypothesis: true rho is not equal to 0
## sample estimates:
##         rho
## -0.04643523
```

```
res <- cor.test(VENEZ_CENTRAL_NONCORE$d_ori, VENEZ_CENTRAL_NONCORE$log2_NormRPK_E, method = "spearman")
```

```
## Warning in cor.test.default(VENEZ_CENTRAL_NONCORE$d_ori,
## VENEZ_CENTRAL_NONCORE$log2_NormRPK_E, : Cannot compute exact p-value with ties
```

```
res
```

```
##
##  Spearman's rank correlation rho
##
## data:  VENEZ_CENTRAL_NONCORE$d_ori and VENEZ_CENTRAL_NONCORE$log2_NormRPK_E
## S = 7553725771, p-value = 0.009721
## alternative hypothesis: true rho is not equal to 0
## sample estimates:
##         rho
## -0.04360587
```

```
res <- cor.test(VENEZ_TERM_CORE$d_ori, VENEZ_TERM_CORE$log2_NormRPK_E, method = "spearman")
res
```

```
##
##  Spearman's rank correlation rho
##
## data:  VENEZ_TERM_CORE$d_ori and VENEZ_TERM_CORE$log2_NormRPK_E
## S = 292670, p-value = 0.1807
## alternative hypothesis: true rho is not equal to 0
## sample estimates:
##        rho
## -0.1250913
```

```
res <- cor.test(VENEZ_TERM_NONCORE$d_ori, VENEZ_TERM_NONCORE$log2_NormRPK_E, method = "spearman")
```

```
## Warning in cor.test.default(VENEZ_TERM_NONCORE$d_ori,
## VENEZ_TERM_NONCORE$log2_NormRPK_E, : Cannot compute exact p-value with ties
```

```
res
```

```
##
##  Spearman's rank correlation rho
##
## data:  VENEZ_TERM_NONCORE$d_ori and VENEZ_TERM_NONCORE$log2_NormRPK_E
## S = 4.321e+09, p-value = 6.681e-05
## alternative hypothesis: true rho is not equal to 0
## sample estimates:
##        rho
## -0.0741016
```

```
AMBO_CCC<-subset(AMBO, AMBO$region=="CENTRAL" & AMBO$CORE_score=="1")
AMBO_TCC<-subset(AMBO, AMBO$region=="TERM" & AMBO$CORE_score=="1")
AMBO_NCCC<-subset(AMBO, AMBO$region=="CENTRAL" & AMBO$CORE_score=="0")
AMBO_NTCC<-subset(AMBO, AMBO$region=="TERM" & AMBO$CORE_score=="0")
AMBO_SMBGCCC<-subset(AMBO, AMBO$region=="CENTRAL" & AMBO$SMBG_score=="1")
AMBO_SMBGCTC<-subset(AMBO, AMBO$region=="TERM" & AMBO$SMBG_score=="1")

AVERM_CCC<-subset(AVERM, AVERM$region=="CENTRAL" & AVERM$CORE_score=="1")
AVERM_TCC<-subset(AVERM, AVERM$region=="TERM" & AVERM$CORE_score=="1")
AVERM_NCCC<-subset(AVERM, AVERM$region=="CENTRAL" & AVERM$CORE_score=="0")
AVERM_NTCC<-subset(AVERM, AVERM$region=="TERM" & AVERM$CORE_score=="0")
AVERM_SMBGCCC<-subset(AVERM, AVERM$region=="CENTRAL" & AVERM$SMBG_score=="1")
AVERM_SMBGCTC<-subset(AVERM, AVERM$region=="TERM" & AVERM$SMBG_score=="1")

BING_CCC<-subset(BING, BING$region=="CENTRAL" & BING$CORE_score=="1")
BING_TCC<-subset(BING, BING$region=="TERM" & BING$CORE_score=="1")
BING_NCCC<-subset(BING, BING$region=="CENTRAL" & BING$CORE_score=="0")
BING_NTCC<-subset(BING, BING$region=="TERM" & BING$CORE_score=="0")
BING_SMBGCCC<-subset(BING, BING$region=="CENTRAL" & BING$SMBG_score=="1")
BING_SMBGCTC<-subset(BING, BING$region=="TERM" & BING$SMBG_score=="1")

CLAV_CCC<-subset(CLAV, CLAV$region=="CENTRAL" & CLAV$CORE_score=="1")
CLAV_TCC<-subset(CLAV, CLAV$region=="TERM" & CLAV$CORE_score=="1")
CLAV_NCCC<-subset(CLAV, CLAV$region=="CENTRAL" & CLAV$CORE_score=="0")
CLAV_NTCC<-subset(CLAV, CLAV$region=="TERM" & CLAV$CORE_score=="0")
CLAV_SMBGCCC<-subset(CLAV, CLAV$region=="CENTRAL" & CLAV$SMBG_score=="1")
CLAV_SMBGCTC<-subset(CLAV, CLAV$region=="TERM" & CLAV$SMBG_score=="1")

SCO_CCC<-subset(SCO, SCO$region=="CENTRAL" & SCO$CORE_score=="1")
SCO_TCC<-subset(SCO, SCO$region=="TERM" & SCO$CORE_score=="1")
SCO_NCCC<-subset(SCO, SCO$region=="CENTRAL" & SCO$CORE_score=="0")
SCO_NTCC<-subset(SCO, SCO$region=="TERM" & SCO$CORE_score=="0")
SCO_SMBGCCC<-subset(SCO, SCO$region=="CENTRAL" & SCO$SMBG_score=="1")
SCO_SMBGCTC<-subset(SCO, SCO$region=="TERM" & SCO$SMBG_score=="1")

TSUKU_CCC<-subset(TSUKU, TSUKU$region=="CENTRAL" & TSUKU$CORE_score=="1")
TSUKU_TCC<-subset(TSUKU, TSUKU$region=="TERM" & TSUKU$CORE_score=="1")
TSUKU_NCCC<-subset(TSUKU, TSUKU$region=="CENTRAL" & TSUKU$CORE_score=="0")
TSUKU_NTCC<-subset(TSUKU, TSUKU$region=="TERM" & TSUKU$CORE_score=="0")
TSUKU_SMBGCCC<-subset(TSUKU, TSUKU$region=="CENTRAL" & TSUKU$SMBG_score=="1")
TSUKU_SMBGCTC<-subset(TSUKU, TSUKU$region=="TERM" & TSUKU$SMBG_score=="1")

VENEZ_CCC<-subset(VENEZ, VENEZ$region=="CENTRAL" & VENEZ$CORE_score=="1")
VENEZ_TCC<-subset(VENEZ, VENEZ$region=="TERM" & VENEZ$CORE_score=="1")
VENEZ_NCCC<-subset(VENEZ, VENEZ$region=="CENTRAL" & VENEZ$CORE_score=="0")
VENEZ_NTCC<-subset(VENEZ, VENEZ$region=="TERM" & VENEZ$CORE_score=="0")
VENEZ_SMBGCCC<-subset(VENEZ, VENEZ$region=="CENTRAL" & VENEZ$SMBG_score=="1")
VENEZ_SMBGCTC<-subset(VENEZ, VENEZ$region=="TERM" & VENEZ$SMBG_score=="1")

boxplot(AMBO_CCC$NormRPK_E, AMBO_TCC$NormRPK_E, AVERM_CCC$NormRPK_E,AVERM_TCC$NormRPK_E,BING_CCC$NormRPK_E,BING_TCC$NormRPK_
E, CLAV_CCC$NormRPK_E,CLAV_TCC$NormRPK_E,SCO_CCC$NormRPK_E,SCO_TCC$NormRPK_E,TSUKU_CCC$NormRPK_E,TSUKU_TCC$NormRPK_E,VENEZ_C
CC$NormRPK_E,VENEZ_TCC$NormRPK_E, outline = FALSE, frame=FALSE, col = c("skyblue2", "lightblue1"), ylab="Transcription (Norm
RPK)", main = "Core genes - Trophophase", names = c("AMB_C", "AMB_T", "AVE_C", "AVE_T","BIN_C","BIN_T","CLA_C", "CLAV_T", "S
CO_C","SCO_T","TSU_C","TSU_T","VEN_C","VEN_T" ), cex.axis = 0.8, las = 2)
```

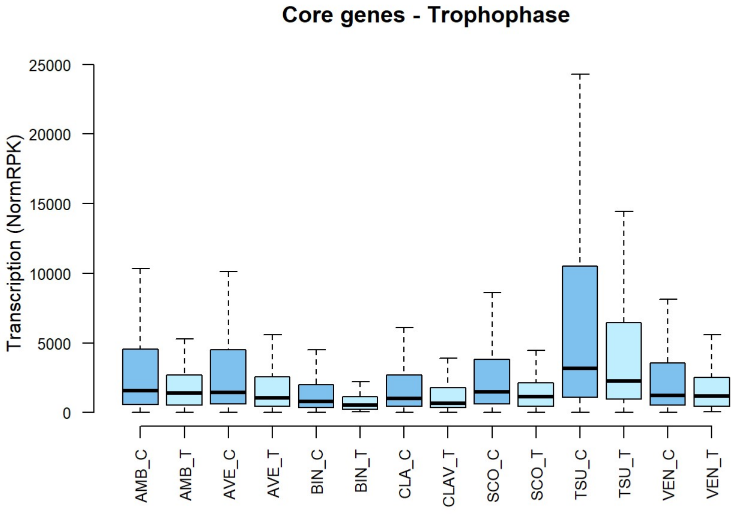


```
wilcox.test(AMBO_CCC$NormRPK_E, AMBO_TCC$NormRPK_E)
```

```
##
##  Wilcoxon rank sum test with continuity correction
##
## data:  AMBO_CCC$NormRPK_E and AMBO_TCC$NormRPK_E
## W = 59139, p-value = 0.02085
## alternative hypothesis: true location shift is not equal to 0
```

```
wilcox.test(AVERM_CCC$NormRPK_E,AVERM_TCC$NormRPK_E)
```

```
##
##  Wilcoxon rank sum test with continuity correction
##
## data:  AVERM_CCC$NormRPK_E and AVERM_TCC$NormRPK_E
## W = 61419, p-value = 0.002095
## alternative hypothesis: true location shift is not equal to 0
```

```
wilcox.test(BING_CCC$NormRPK_E,BING_TCC$NormRPK_E)
```

```
##
##  Wilcoxon rank sum test with continuity correction
##
## data:  BING_CCC$NormRPK_E and BING_TCC$NormRPK_E
## W = 43232, p-value = 0.004491
## alternative hypothesis: true location shift is not equal to 0
```

```
wilcox.test(CLAV_CCC$NormRPK_E,CLAV_TCC$NormRPK_E)
```

```
##
##  Wilcoxon rank sum test with continuity correction
##
## data:  CLAV_CCC$NormRPK_E and CLAV_TCC$NormRPK_E
## W = 59914, p-value = 0.006664
## alternative hypothesis: true location shift is not equal to 0
```

```
wilcox.test(SCO_CCC$NormRPK_E,SCO_TCC$NormRPK_E)
```

```
##
##  Wilcoxon rank sum test with continuity correction
##
## data:  SCO_CCC$NormRPK_E and SCO_TCC$NormRPK_E
## W = 61479, p-value = 0.001959
## alternative hypothesis: true location shift is not equal to 0
```

```
wilcox.test(TSUKU_CCC$NormRPK_E,TSUKU_TCC$NormRPK_E)
```

```
##
##  Wilcoxon rank sum test with continuity correction
##
## data:  TSUKU_CCC$NormRPK_E and TSUKU_TCC$NormRPK_E
## W = 58892, p-value = 0.0259
## alternative hypothesis: true location shift is not equal to 0
```

```
wilcox.test(VENEZ_CCC$NormRPK_E,VENEZ_TCC$NormRPK_E)
```

```
##
##  Wilcoxon rank sum test with continuity correction
##
## data:  VENEZ_CCC$NormRPK_E and VENEZ_TCC$NormRPK_E
## W = 56242, p-value = 0.181
## alternative hypothesis: true location shift is not equal to 0
```

```
boxplot(AMBO_CCC$NormRPK_DIFF_METABO, AMBO_TCC$NormRPK_DIFF_METABO, AVERM_CCC$NormRPK_DIFF_METABO,AVERM_TCC$NormRPK_DIFF_MET
ABO, BING_CCC$NormRPK_DIFF_METABO,BING_TCC$NormRPK_DIFF_METABO, CLAV_CCC$NormRPK_DIFF_METABO,CLAV_TCC$NormRPK_DIFF_METABO,
        SCO_CCC$NormRPK_DIFF_METABO,SCO_TCC$NormRPK_DIFF_METABO,TSUKU_CCC$NormRPK_DIFF_METABO,TSUKU_TCC$NormRPK_DIFF_METABO,
VENEZ_CCC$NormRPK_DIFF_METABO,VENEZ_TCC$NormRPK_DIFF_METABO, col = c("skyblue2", "lightblue1"), ylab="Transcription (NormRP
K)",frame=FALSE, outline = FALSE, main = "Core genes - Idiophase", names = c("AMB_C", "AMB_T", "AVE_C", "AVE_T","BIN_C","BIN
_T","CLA_C", "CLAV_T", "SCO_C","SCO_T","TSU_C","TSU_T","VEN_C","VEN_T" ), cex.axis = 0.8, las = 2)
```

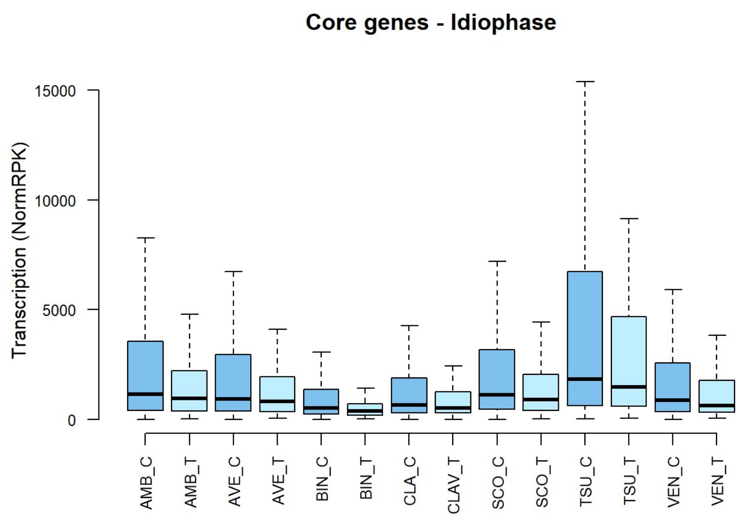


```
wilcox.test(AMBO_CCC$NormRPK_DIFF_METABO, AMBO_TCC$NormRPK_DIFF_METABO)
```

```
##
##  Wilcoxon rank sum test with continuity correction
##
## data:  AMBO_CCC$NormRPK_DIFF_METABO and AMBO_TCC$NormRPK_DIFF_METABO
## W = 56100, p-value = 0.197
## alternative hypothesis: true location shift is not equal to 0
```

```
wilcox.test(AVERM_CCC$NormRPK_DIFF_METABO,AVERM_TCC$NormRPK_DIFF_METABO)
```

```
##
##  Wilcoxon rank sum test with continuity correction
##
## data:  AVERM_CCC$NormRPK_DIFF_METABO and AVERM_TCC$NormRPK_DIFF_METABO
## W = 56645, p-value = 0.1407
## alternative hypothesis: true location shift is not equal to 0
```

```
wilcox.test(BING_CCC$NormRPK_DIFF_METABO,BING_TCC$NormRPK_DIFF_METABO)
```

```
##
##  Wilcoxon rank sum test with continuity correction
##
## data:  BING_CCC$NormRPK_DIFF_METABO and BING_TCC$NormRPK_DIFF_METABO
## W = 42651, p-value = 0.009134
## alternative hypothesis: true location shift is not equal to 0
```

```
wilcox.test(CLAV_CCC$NormRPK_DIFF_METABO,CLAV_TCC$NormRPK_DIFF_METABO)
```

```
##
##  Wilcoxon rank sum test with continuity correction
##
## data:  CLAV_CCC$NormRPK_DIFF_METABO and CLAV_TCC$NormRPK_DIFF_METABO
## W = 58097, p-value = 0.03567
## alternative hypothesis: true location shift is not equal to 0
```

```
wilcox.test(SCO_CCC$NormRPK_DIFF_METABO,SCO_TCC$NormRPK_DIFF_METABO)
```

```
##
##  Wilcoxon rank sum test with continuity correction
##
## data:  SCO_CCC$NormRPK_DIFF_METABO and SCO_TCC$NormRPK_DIFF_METABO
## W = 58383, p-value = 0.0397
## alternative hypothesis: true location shift is not equal to 0
```

```
wilcox.test(TSUKU_CCC$NormRPK_DIFF_METABO,TSUKU_TCC$NormRPK_DIFF_METABO)
```

```
##
##  Wilcoxon rank sum test with continuity correction
##
## data:  TSUKU_CCC$NormRPK_DIFF_METABO and TSUKU_TCC$NormRPK_DIFF_METABO
## W = 56458, p-value = 0.1584
## alternative hypothesis: true location shift is not equal to 0
```

```
wilcox.test(VENEZ_CCC$NormRPK_DIFF_METABO,VENEZ_TCC$NormRPK_DIFF_METABO)
```

```
##
##  Wilcoxon rank sum test with continuity correction
##
## data:  VENEZ_CCC$NormRPK_DIFF_METABO and VENEZ_TCC$NormRPK_DIFF_METABO
## W = 57878, p-value = 0.05913
## alternative hypothesis: true location shift is not equal to 0
```

```
boxplot(AMBO_NCCC$NormRPK_E, AMBO_NTCC$NormRPK_E, AVERM_NCCC$NormRPK_E,AVERM_NTCC$NormRPK_E, BING_NCCC$NormRPK_E,BING_NTCC$N
ormRPK_E, CLAV_NCCC$NormRPK_E,CLAV_NTCC$NormRPK_E,
        SCO_NCCC$NormRPK_E,SCO_NTCC$NormRPK_E,TSUKU_NCCC$NormRPK_E,TSUKU_NTCC$NormRPK_E,VENEZ_NCCC$NormRPK_E,VENEZ_NTCC$Norm
RPK_E, col = c("skyblue2", "lightblue1"), ylab="Transcription (NormRPK)",frame=FALSE, outline = FALSE, main = "Non core gene
s - Trophophase", names = c("AMB_C", "AMB_T", "AVE_C", "AVE_T","BIN_C","BIN_T","CLA_C", "CLAV_T", "SCO_C","SCO_T","TSU_C","T
SU_T","VEN_C","VEN_T" ), cex.axis = 0.8, las = 2)
```

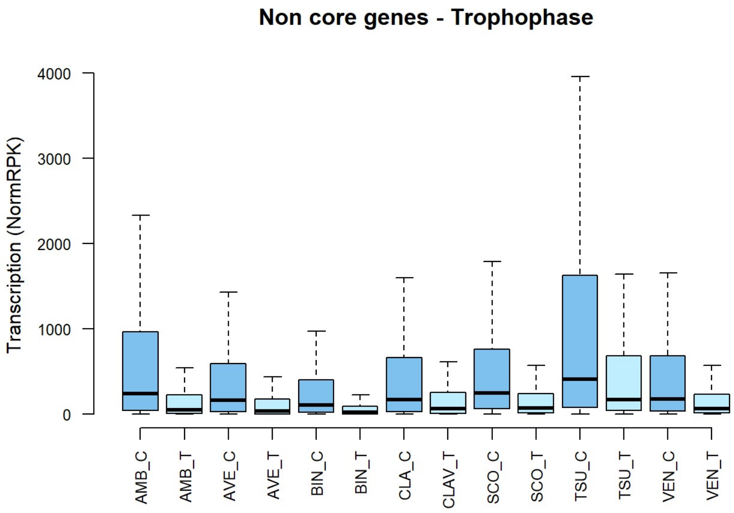


```
wilcox.test(AMBO_NCCC$NormRPK_E, AMBO_NTCC$NormRPK_E)
```

```
##
##  Wilcoxon rank sum test with continuity correction
##
## data:  AMBO_NCCC$NormRPK_E and AMBO_NTCC$NormRPK_E
## W = 6580272, p-value < 2.2e-16
## alternative hypothesis: true location shift is not equal to 0
```

```
wilcox.test(AVERM_NCCC$NormRPK_E,AVERM_NTCC$NormRPK_E)
```

```
##
##  Wilcoxon rank sum test with continuity correction
##
## data:  AVERM_NCCC$NormRPK_E and AVERM_NTCC$NormRPK_E
## W = 8279923, p-value < 2.2e-16
## alternative hypothesis: true location shift is not equal to 0
```

```
wilcox.test(BING_NCCC$NormRPK_E,BING_NTCC$NormRPK_E)
```

```
##
##  Wilcoxon rank sum test with continuity correction
##
## data:  BING_NCCC$NormRPK_E and BING_NTCC$NormRPK_E
## W = 14271539, p-value < 2.2e-16
## alternative hypothesis: true location shift is not equal to 0
```

```
wilcox.test(CLAV_NCCC$NormRPK_E,CLAV_NTCC$NormRPK_E)
```

```
##
##  Wilcoxon rank sum test with continuity correction
##
## data:  CLAV_NCCC$NormRPK_E and CLAV_NTCC$NormRPK_E
## W = 3070609, p-value < 2.2e-16
## alternative hypothesis: true location shift is not equal to 0
```

```
wilcox.test(SCO_NCCC$NormRPK_E,SCO_NTCC$NormRPK_E)
```

```
##
##  Wilcoxon rank sum test with continuity correction
##
## data:  SCO_NCCC$NormRPK_E and SCO_NTCC$NormRPK_E
## W = 8063022, p-value < 2.2e-16
## alternative hypothesis: true location shift is not equal to 0
```

```
wilcox.test(TSUKU_NCCC$NormRPK_E,TSUKU_NTCC$NormRPK_E)
```

```
##
##  Wilcoxon rank sum test with continuity correction
##
## data:  TSUKU_NCCC$NormRPK_E and TSUKU_NTCC$NormRPK_E
## W = 4674836, p-value < 2.2e-16
## alternative hypothesis: true location shift is not equal to 0
```

```
wilcox.test(VENEZ_NCCC$NormRPK_E,VENEZ_NTCC$NormRPK_E)
```

```
##
##  Wilcoxon rank sum test with continuity correction
##
## data:  VENEZ_NCCC$NormRPK_E and VENEZ_NTCC$NormRPK_E
## W = 6505225, p-value < 2.2e-16
## alternative hypothesis: true location shift is not equal to 0
```

```
boxplot(AMBO_NCCC$NormRPK_DIFF_METABO, AMBO_NTCC$NormRPK_DIFF_METABO, AVERM_NCCC$NormRPK_DIFF_METABO,AVERM_NTCC$NormRPK_DIFF
_METABO, BING_NCCC$NormRPK_DIFF_METABO,BING_NTCC$NormRPK_DIFF_METABO, CLAV_NCCC$NormRPK_DIFF_METABO,CLAV_NTCC$NormRPK_DIFF_M
ETABO,
        SCO_NCCC$NormRPK_DIFF_METABO,SCO_NTCC$NormRPK_DIFF_METABO,TSUKU_NCCC$NormRPK_DIFF_METABO,TSUKU_NTCC$NormRPK_DIFF_MET
ABO,VENEZ_NCCC$NormRPK_DIFF_METABO,VENEZ_NTCC$NormRPK_DIFF_METABO, col = c("skyblue2", "lightblue1"), ylab="Transcription (N
ormRPK)",frame=FALSE, outline = FALSE, main = "Non core genes - Idiophase", names = c("AMB_C", "AMB_T", "AVE_C", "AVE_T","BI
N_C","BIN_T","CLA_C", "CLAV_T", "SCO_C","SCO_T","TSU_C","TSU_T","VEN_C","VEN_T" ), cex.axis = 0.8, las = 2)
```

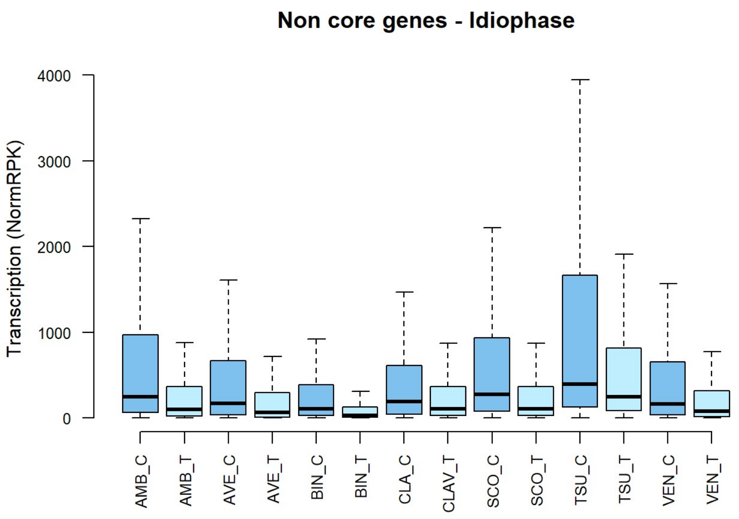


```
wilcox.test(AMBO_NCCC$NormRPK_DIFF_METABO, AMBO_NTCC$NormRPK_DIFF_METABO)
```

```
##
##  Wilcoxon rank sum test with continuity correction
##
## data:  AMBO_NCCC$NormRPK_DIFF_METABO and AMBO_NTCC$NormRPK_DIFF_METABO
## W = 5964700, p-value < 2.2e-16
## alternative hypothesis: true location shift is not equal to 0
```

```
wilcox.test(AVERM_NCCC$NormRPK_DIFF_METABO,AVERM_NTCC$NormRPK_DIFF_METABO)
```

```
##
##  Wilcoxon rank sum test with continuity correction
##
## data:  AVERM_NCCC$NormRPK_DIFF_METABO and AVERM_NTCC$NormRPK_DIFF_METABO
## W = 7637165, p-value < 2.2e-16
## alternative hypothesis: true location shift is not equal to 0
```

```
wilcox.test(BING_NCCC$NormRPK_DIFF_METABO,BING_NTCC$NormRPK_DIFF_METABO)
```

```
##
##  Wilcoxon rank sum test with continuity correction
##
## data:  BING_NCCC$NormRPK_DIFF_METABO and BING_NTCC$NormRPK_DIFF_METABO
## W = 13419324, p-value < 2.2e-16
## alternative hypothesis: true location shift is not equal to 0
```

```
wilcox.test(CLAV_NCCC$NormRPK_DIFF_METABO,CLAV_NTCC$NormRPK_DIFF_METABO)
```

```
##
##  Wilcoxon rank sum test with continuity correction
##
## data:  CLAV_NCCC$NormRPK_DIFF_METABO and CLAV_NTCC$NormRPK_DIFF_METABO
## W = 2821994, p-value < 2.2e-16
## alternative hypothesis: true location shift is not equal to 0
```

```
wilcox.test(SCO_NCCC$NormRPK_DIFF_METABO,SCO_NTCC$NormRPK_DIFF_METABO)
```

```
##
##  Wilcoxon rank sum test with continuity correction
##
## data:  SCO_NCCC$NormRPK_DIFF_METABO and SCO_NTCC$NormRPK_DIFF_METABO
## W = 7500321, p-value < 2.2e-16
## alternative hypothesis: true location shift is not equal to 0
```

```
wilcox.test(TSUKU_NCCC$NormRPK_DIFF_METABO,TSUKU_NTCC$NormRPK_DIFF_METABO)
```

```
##
##  Wilcoxon rank sum test with continuity correction
##
## data:  TSUKU_NCCC$NormRPK_DIFF_METABO and TSUKU_NTCC$NormRPK_DIFF_METABO
## W = 4449643, p-value < 2.2e-16
## alternative hypothesis: true location shift is not equal to 0
```

```
wilcox.test(VENEZ_NCCC$NormRPK_DIFF_METABO,VENEZ_NTCC$NormRPK_DIFF_METABO)
```

```
##
##  Wilcoxon rank sum test with continuity correction
##
## data:  VENEZ_NCCC$NormRPK_DIFF_METABO and VENEZ_NTCC$NormRPK_DIFF_METABO
## W = 6056197, p-value < 2.2e-16
## alternative hypothesis: true location shift is not equal to 0
```

```
boxplot(AMBO_SMBGCCC$NormRPK_E, AMBO_SMBGCTC$NormRPK_E, AVERM_SMBGCCC$NormRPK_E,AVERM_SMBGCTC$NormRPK_E, BING_SMBGCCC$NormRP
K_E,BING_SMBGCTC$NormRPK_E, CLAV_SMBGCCC$NormRPK_E,CLAV_SMBGCTC$NormRPK_E,
        SCO_SMBGCCC$NormRPK_E,SCO_SMBGCTC$NormRPK_E,TSUKU_SMBGCCC$NormRPK_E,TSUKU_SMBGCTC$NormRPK_E,VENEZ_SMBGCCC$NormRPK_E,
VENEZ_SMBGCTC$NormRPK_E, col = c("skyblue2", "lightblue1"), ylab="Transcription (NormRPK)",frame=FALSE, outline = FALSE, mai
n = "SMBGCs - Trophophase", names = c("AMB_C", "AMB_T", "AVE_C", "AVE_T","BIN_C","BIN_T","CLA_C", "CLAV_T", "SCO_C","SCO_
T","TSU_C","TSU_T","VEN_C","VEN_T" ), cex.axis = 0.8, las = 2)
```

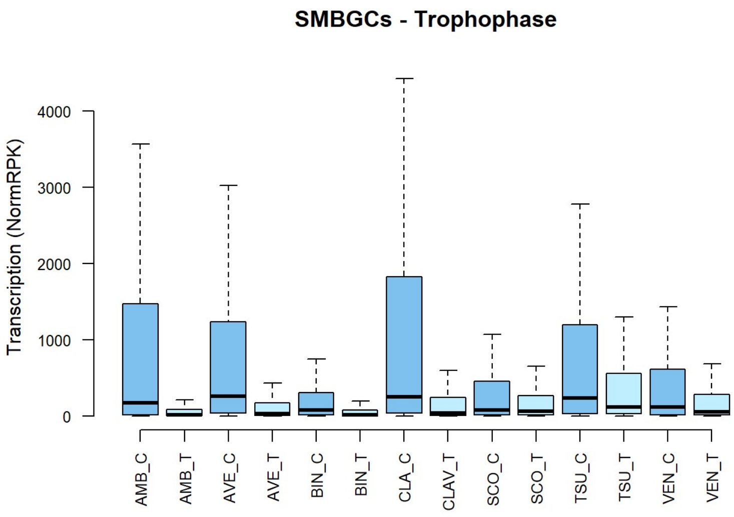


```
wilcox.test(AMBO_SMBGCCC$NormRPK_E, AMBO_SMBGCTC$NormRPK_E)
```

```
##
##  Wilcoxon rank sum test with continuity correction
##
## data:  AMBO_SMBGCCC$NormRPK_E and AMBO_SMBGCTC$NormRPK_E
## W = 17122, p-value = 1.033e-09
## alternative hypothesis: true location shift is not equal to 0
```

```
wilcox.test(AVERM_SMBGCCC$NormRPK_E,AVERM_SMBGCTC$NormRPK_E)
```

```
##
##  Wilcoxon rank sum test with continuity correction
##
## data:  AVERM_SMBGCCC$NormRPK_E and AVERM_SMBGCTC$NormRPK_E
## W = 196686, p-value < 2.2e-16
## alternative hypothesis: true location shift is not equal to 0
```

```
wilcox.test(BING_SMBGCCC$NormRPK_E,BING_SMBGCTC$NormRPK_E)
```

```
##
##  Wilcoxon rank sum test with continuity correction
##
## data:  BING_SMBGCCC$NormRPK_E and BING_SMBGCTC$NormRPK_E
## W = 511296, p-value < 2.2e-16
## alternative hypothesis: true location shift is not equal to 0
```

```
wilcox.test(CLAV_SMBGCCC$NormRPK_E,CLAV_SMBGCTC$NormRPK_E)
```

```
##
##  Wilcoxon rank sum test with continuity correction
##
## data:  CLAV_SMBGCCC$NormRPK_E and CLAV_SMBGCTC$NormRPK_E
## W = 100242, p-value < 2.2e-16
## alternative hypothesis: true location shift is not equal to 0
```

```
wilcox.test(SCO_SMBGCCC$NormRPK_E,SCO_SMBGCTC$NormRPK_E)
```

```
##
##  Wilcoxon rank sum test with continuity correction
##
## data:  SCO_SMBGCCC$NormRPK_E and SCO_SMBGCTC$NormRPK_E
## W = 69559, p-value = 0.1088
## alternative hypothesis: true location shift is not equal to 0
```

```
wilcox.test(TSUKU_SMBGCCC$NormRPK_E,TSUKU_SMBGCTC$NormRPK_E)
```

```
##
##  Wilcoxon rank sum test with continuity correction
##
## data:  TSUKU_SMBGCCC$NormRPK_E and TSUKU_SMBGCTC$NormRPK_E
## W = 161501, p-value = 0.0001947
## alternative hypothesis: true location shift is not equal to 0
```

```
wilcox.test(VENEZ_SMBGCCC$NormRPK_E,VENEZ_SMBGCTC$NormRPK_E)
```

```
##
##  Wilcoxon rank sum test with continuity correction
##
## data:  VENEZ_SMBGCCC$NormRPK_E and VENEZ_SMBGCTC$NormRPK_E
## W = 78548, p-value = 0.0008273
## alternative hypothesis: true location shift is not equal to 0
```

```
boxplot(AMBO_SMBGCCC$NormRPK_DIFF_METABO, AMBO_SMBGCTC$NormRPK_DIFF_METABO, AVERM_SMBGCCC$NormRPK_DIFF_METABO,AVERM_SMBGCT
C$NormRPK_DIFF_METABO, BING_SMBGCCC$NormRPK_DIFF_METABO,BING_SMBGCTC$NormRPK_DIFF_METABO, CLAV_SMBGCCC$NormRPK_DIFF_METABO,C
LAV_SMBGCTC$NormRPK_DIFF_METABO,
        SCO_SMBGCCC$NormRPK_DIFF_METABO,SCO_SMBGCTC$NormRPK_DIFF_METABO,TSUKU_SMBGCCC$NormRPK_DIFF_METABO,TSUKU_SMBGCTC$Norm
RPK_DIFF_METABO,VENEZ_SMBGCCC$NormRPK_DIFF_METABO,VENEZ_SMBGCTC$NormRPK_DIFF_METABO, col = c("skyblue2", "lightblue1"), ylab
="Transcription (NormRPK)",frame=FALSE, outline = FALSE, main = "SMBGCs - Idiophase", names = c("AMB_C", "AMB_T", "AVE_C", "
AVE_T","BIN_C","BIN_T","CLA_C", "CLAV_T", "SCO_C","SCO_T","TSU_C","TSU_T","VEN_C","VEN_T" ), cex.axis = 0.8, las = 2)
```

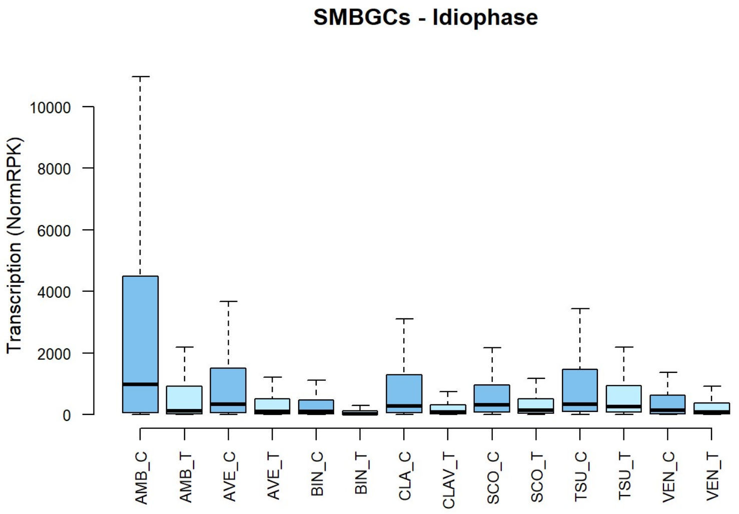


```
wilcox.test(AMBO_SMBGCCC$NormRPK_DIFF_METABO, AMBO_SMBGCTC$NormRPK_DIFF_METABO)
```

```
##
##  Wilcoxon rank sum test with continuity correction
##
## data:  AMBO_SMBGCCC$NormRPK_DIFF_METABO and AMBO_SMBGCTC$NormRPK_DIFF_METABO
## W = 15344, p-value = 6.421e-05
## alternative hypothesis: true location shift is not equal to 0
```

```
wilcox.test(AVERM_SMBGCCC$NormRPK_DIFF_METABO,AVERM_SMBGCTC$NormRPK_DIFF_METABO)
```

```
##
##  Wilcoxon rank sum test with continuity correction
##
## data:  AVERM_SMBGCCC$NormRPK_DIFF_METABO and AVERM_SMBGCTC$NormRPK_DIFF_METABO
## W = 171699, p-value = 3.837e-13
## alternative hypothesis: true location shift is not equal to 0
```

```
wilcox.test(BING_SMBGCCC$NormRPK_DIFF_METABO,BING_SMBGCTC$NormRPK_DIFF_METABO)
```

```
##
##  Wilcoxon rank sum test with continuity correction
##
## data:  BING_SMBGCCC$NormRPK_DIFF_METABO and BING_SMBGCTC$NormRPK_DIFF_METABO
## W = 506218, p-value < 2.2e-16
## alternative hypothesis: true location shift is not equal to 0
```

```
wilcox.test(CLAV_SMBGCCC$NormRPK_DIFF_METABO,CLAV_SMBGCTC$NormRPK_DIFF_METABO)
```

```
##
##  Wilcoxon rank sum test with continuity correction
##
## data:  CLAV_SMBGCCC$NormRPK_DIFF_METABO and CLAV_SMBGCTC$NormRPK_DIFF_METABO
## W = 91685, p-value = 9.352e-10
## alternative hypothesis: true location shift is not equal to 0
```

```
wilcox.test(SCO_SMBGCCC$NormRPK_DIFF_METABO,SCO_SMBGCTC$NormRPK_DIFF_METABO)
```

```
##
##  Wilcoxon rank sum test with continuity correction
##
## data:  SCO_SMBGCCC$NormRPK_DIFF_METABO and SCO_SMBGCTC$NormRPK_DIFF_METABO
## W = 76227, p-value = 0.0001127
## alternative hypothesis: true location shift is not equal to 0
```

```
wilcox.test(TSUKU_SMBGCCC$NormRPK_DIFF_METABO,TSUKU_SMBGCTC$NormRPK_DIFF_METABO)
```

```
##
##  Wilcoxon rank sum test with continuity correction
##
## data:  TSUKU_SMBGCCC$NormRPK_DIFF_METABO and TSUKU_SMBGCTC$NormRPK_DIFF_METABO
## W = 155615, p-value = 0.009703
## alternative hypothesis: true location shift is not equal to 0
```

```
wilcox.test(VENEZ_SMBGCCC$NormRPK_DIFF_METABO,VENEZ_SMBGCTC$NormRPK_DIFF_METABO)
```

```
##
##  Wilcoxon rank sum test with continuity correction
##
## data:  VENEZ_SMBGCCC$NormRPK_DIFF_METABO and VENEZ_SMBGCTC$NormRPK_DIFF_METABO
## W = 74591, p-value = 0.03706
## alternative hypothesis: true location shift is not equal to 0
```
