## Supplementary material for "Ribosomal RNA operons define a central functional compartment in the *Streptomyces* chromosome": Supplemental_Table_S3.pdf

### Table S3: Remarkable species regarding core gene order compared to the *Streptomyces* consensus

| **Species with core genes ordered exactly as in the consensus** |
|---|
| *Streptomyces viridosporus* T7A ATCC 39115<br>*Streptomyces* sp. GGCR 6<br>*Streptomyces* sp. QMT 28<br>*Streptomyces* sp. S10 2016<br>*Streptomyces* sp. S1A1 7 |
| **Species with core genes ordered as in the consensus, except for two genes (local inversion)** |
| *Streptomyces ambofaciens* ATCC 23877<br>*Streptomyces ambofaciens* DSM 40697<br>*Streptomyces chartreusis* NRRL 3882<br>*Streptomyces coeruleorubidus* ATCC 13740<br>*Streptomyces ficellus* NRRL 8067<br>*Streptomyces galilaeus* ATCC 14969<br>*Streptomyces pactum* ACT12<br>*Streptomyces parvulus* 2297<br>*Streptomyces* sp. CCM MD2014<br>*Streptomyces* sp. Go 475<br>*Streptomyces* sp. SS52<br>*Streptomyces* sp. SYP A7193 |
